# The impact of protein architecture on adaptive evolution

**DOI:** 10.1101/560185

**Authors:** Ana Filipa Moutinho, Fernanda Fontes Trancoso, Julien Yann Dutheil

**Affiliations:** Department of Evolutionary Genetics, Max Planck Institute for Evolutionary Biology, 24306 Plön, Germany; Unité Mixte de Recherche 5554 Institut des Sciences de l’Evolution, CNRS, IRD, EPHE, Université de Montpellier, Place E. Bataillon, 34095, Montpellier, France

**Keywords:** protein structure, protein function, adaptation, population genetics, *Drosophila melanogaster*, *Arabidopsis thaliana*

## Abstract

Adaptive mutations play an important role in molecular evolution. However, the frequency and nature of these mutations at the intra-molecular level is poorly understood. To address this, we analysed the impact of protein architecture on the rate of adaptive substitutions, aiming to understand how protein biophysics influences fitness and adaptation. Using *Drosophila melanogaster* and *Arabidopsis thaliana* population genomics data, we fitted models of distribution of fitness effects and estimated the rate of adaptive amino-acid substitutions both at the protein and amino-acid residue level. We performed a comprehensive analysis covering genome, gene and protein structure, by exploring a multitude of factors with a plausible impact on the rate of adaptive evolution, such as intron number, protein length, secondary structure, relative solvent accessibility, intrinsic protein disorder, chaperone affinity, gene expression, protein function and protein-protein interactions. We found that the relative solvent accessibility is a major driver of adaptive evolution, with most adaptive mutations occurring at the surface of proteins. Moreover, we observe that the rate of adaptive substitutions differs between protein functional classes, with genes encoding for protein biosynthesis and degradation signalling exhibiting the fastest rates of protein adaptation. Overall, our results suggest that adaptive evolution in proteins is mainly driven by inter-molecular interactions, with host-pathogen coevolution likely playing a major role.

## Introduction

A long-standing focus in the study of molecular evolution is the role of natural selection in protein evolution (Eyre-Walker 2006). One can measure the strength and direction of selection at the divergence level through the *d*_*N*_/*d*_*S*_ ratio (ω). However, because ω represents a summary statistic across nucleotide sites, it can only provide the average trend, while proteins will typically undergo both negative and positive selection. Because adaptive mutations contribute relatively more to substitution than to polymorphism, we can disentangle the two types of selection by contrasting the number of substitutions to the number of polymorphisms at synonymous and non-synonymous sites, as implemented in the McDonald and Kreitman (MK) test (McDonald and Kreitman 1991). Charlesworth (1994) extended the MK test to estimate the proportion of substitutions that are adaptive (α). Yet, one limitation of this approach was that it didn’t account for the segregation of slightly deleterious mutations, which can either over- or underestimate measurements of α according to the demography of the population (Eyre-Walker 2002; Smith and Eyre-Walker 2002). Recent methods solved this issue by taking into consideration the distribution of fitness effects (DFE) of both slightly deleterious (Fay et al. 2001; Smith and Eyre-Walker 2002; Bierne and Eyre-Walker 2004; Eyre-Walker et al. 2006; Eyre-Walker and Keightley 2009; Stoletzki and Eyre-Walker 2011) and slightly beneficial mutations (Galtier 2016; Tataru et al. 2017). By allowing the estimation of the rate of non-adaptive 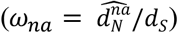 and adaptive (*ω*_*a*_ = ω − *ω*_*na*_) non-synonymous substitutions, in addition to measurements of α (*ω*_*a*_/ω), these methods triggered new insights on the impact of both negative and positive selection on the rate of protein evolution.

Several studies have reported substantial levels of adaptive protein evolution in various animal species, including the fruit fly (Smith and Eyre-Walker 2002; Sawyer et al. 2003; Bierne and Eyre-Walker 2004; Haddrill et al. 2010), the wild mouse (Halligan et al. 2010) and the European rabbit (Carneiro et al. 2012), but also in bacteria (Charlesworth and Eyre-Walker 2006) and in plants (Ingvarsson 2010; Slotte et al. 2010; Strasburg et al. 2011). Whereas for other taxa, such as primates (Boyko et al. 2008; Hvilsom et al. 2012; Galtier 2016), and many other plants (Gossmann et al. 2010), the rate of adaptive mutations was observed to be very low, wherein amino-acid substitutions are expected to be nearly neutral and fixed mainly through random genetic drift (Boyko et al. 2008). Several authors proposed that this across-species variation in the molecular adaptive rate is explained by an effective population size (*N*_*e*_) effect, where higher rates of adaptive evolution are observed for species with larger *N*_*e*_ due to a lower impact of genetic drift (Eyre-Walker 2006; Eyre-Walker and Keightley 2009; Gossmann et al. 2012). Galtier (2016), however, reported that *N*_*e*_ had an impact on α and *ω*_*na*_ but not *ω*_*a*_. Hence, he proposed that the relationship with *N*_*e*_ is mainly explained by deleterious effects, wherein slightly deleterious non-synonymous substitutions accumulate at lower rates in large-*N*_*e*_ species due to a higher efficiency of purifying selection, thus decreasing ω and consequently inflating α.

The rate of adaptive substitutions, however, was observed to vary extensively along the genome. On a genome-wide scale, it was reported that *ω*_*a*_ correlates positively with both the recombination and mutation rates, but negatively with gene density (Campos et al. 2014; Castellano et al. 2016). When looking at the gene level, previous studies have demonstrated the role of protein function in the rate of adaptive evolution, wherein genes involved in immune defence mechanisms appear with higher rates of adaptive mutations in Drosophila (Sackton et al. 2007; Obbard et al. 2009), humans and chimpanzees (Nielsen et al. 2005). In Drosophila, sex-related genes also display higher levels of adaptive evolution, being directly linked with species differentiation (Pröschel et al. 2006; Haerty et al. 2007). At the intra-genic level, however, the frequency and nature of adaptive mutations remain poorly understood.

There are several structural factors that have been reported to influence the rate of protein evolution but have not been investigated at the population level. Molecular evolution studies of protein families revealed that protein structure, for instance, significantly impacts the rate of amino-acid substitutions, with exposed residues evolving faster than buried ones (Liberles et al. 2012). As a stable conformation is often required to ensure proper protein function, mutations that impair the stability or the structural conformation of the folded protein are more likely to be counter-selected. Moreover, distinct sites in a protein sequence differ in the extent of conformational change they endure upon mutation, a pattern generally well predicted by the relative solvent accessibility of a residue (Goldman et al. 1998; Mirny and Shakhnovich 1999; Franzosa and Xia 2009). In this way, residues at the core of proteins evolve slower than the ones at the surface due to their role in maintaining a stable protein structure (Perutz et al. 1965; Overington et al. 1992; Goldman et al. 1998; Bustamante et al. 2000; Dean et al. 2002; Choi et al. 2006; Lin et al. 2007; Conant and Stadler 2009; Franzosa and Xia 2009; Ramsey et al. 2011). Inter-specific comparative sequence analyses also revealed that positively selected sites are often found at the surface of proteins (Proux et al. 2009; Adams et al. 2017). Hence, exploring the role that these structural elements play in shaping the rate of adaptive evolution is crucial in order to fully understand what are the main drivers of adaptation within proteomes. Inter-specific comparisons, however, do not permit to disentangle the effect of relaxed purifying selection from adaptation-driven increase in substitutions rates. Such insights can only be obtained by contrasting patterns of polymorphisms and divergence.

Our study addresses protein adaptive evolution at a fine scale by analysing the impact of several functional variables among protein-coding regions at the population level. To further assess the potential generality of the inferred effects, we carried our comparison on two model species with distinct life-history traits: the dipter *Drosophila melanogaster* and the brassicaceae *Arabidopsis thaliana*. We fitted models of DFE and estimated the rate of adaptive substitutions, both at the protein and amino-acid residue scale, across several variables and found that solvent exposure is the most significant factor influencing protein adaptation, with exposed residues undergoing ten times faster *ω*_*a*_ than buried ones. Moreover, we observed that the functional class of proteins has also a strong impact on the rate of protein adaptation, with genes encoding for processes of protein regulation and signaling pathways exhibiting the highest *ω*_*a*_ values. We therefore hypothesized that inter-molecular interactions are the main drivers of adaptive substitutions in proteins. This hypothesis is consistent with the proposal that, at the inter-organism level, coevolution with pathogens constitute a so-far under-assessed component of protein evolution (Sackton et al. 2007; Obbard et al. 2009; Enard et al. 2016; Mauch-Mani et al. 2017).

## Results and Discussion

In order to identify the genomic and structural variants driving protein adaptive evolution we looked at 10,318 protein-coding genes for *Drosophila melanogaster*, analyzing polymorphism data from an admixed sub-Saharan population (DPGP2, Pool et al. 2012) and divergence out to *D. simulans*; and 18,669 protein-coding genes for *Arabidopsis thaliana*, with polymorphism data from a Spanish population (1001 Genomes Project, Weigel and Mott 2009) and divergence to *A. lyrata*. The rate of adaptive evolution was estimated with the Grapes program (Galtier 2016). The Grapes method extends the approach pioneered by the DoFE program (Fay et al. 2001; Smith and Eyre-Walker 2002; Bierne and Eyre-Walker 2004; Eyre-Walker et al. 2006; Eyre-Walker and Keightley 2009; Stoletzki and Eyre-Walker 2011), by explicitly accounting for mutations with slightly advantageous effects. Grapes estimates the rate of non-adaptive non-synonymous substitutions (*ω*_*na*_), which is then used to estimate the rate of adaptive non-synonymous substitutions (*ω*_*a*_) and the proportion of adaptive non-synonymous substitutions (α). A high α can be potentially explained both by a higher *ω*_*a*_ or a lower *ω*_*na*_, and therefore does not allow to disentangle the two effects. Thus, we explored how, and if, *ω*_*a*_ and *ω*_*na*_, as well as the total ω, depend on the different functional variables analysed here.

Results from model comparison of DFE showed that the Gamma-Exponential model is the one that best fits our data according to Akaike’s information criterion (Akaike 1973) (Table S1 in supplementary File S1). This model combines a Gamma distribution of deleterious mutations with an exponential distribution of beneficial mutations. In agreement with previous surveys within animal species, this model suggests the existence of slightly deleterious, as well as slightly beneficial segregating mutations in *D. melanogaster* and *A. thaliana* genomes (Galtier 2016). Genome-wide estimates of *ω*_*a*_ for *A. thaliana* and *D. melanogaster* are 0.05 and 0.09, respectively, and are in the range of previously reported estimates for these species (Bierne and Eyre-Walker 2004; Gossmann et al. 2012; Smith and Eyre-Walker 2002).

We then categorized genes and amino-acid residues in order to investigate the main drivers of protein adaptive evolution. We distinguished two types of analyses: gene-based and site-based, where we looked into how the molecular adaptive rate varies across different categories of genes and amino-acid residues, respectively. Gene-based analyses allowed us to explore the impact of the background recombination rate, number of introns and mean expression levels. At the protein level, we investigated the effect of binding affinity to the molecular chaperone *DnaK*, protein length, cellular localization of proteins, protein functional class and number of protein-protein interactions. Finally, site-based analyses enabled us to study the effect of the secondary structure of the protein, by comparing residues present in β-sheets, α-helices and loops; the tertiary structure, by considering the relative solvent accessibility of a residue (RSA) and the residue intrinsic disorder; and whether an amino-acid residue participated or not in an annotated active site.

### The impact of gene and genome architecture on adaptive evolution

To study the impact of gene and genome architecture on the rate of adaptive evolution we looked at recombination rate and the number of introns. Recombination rate was previously reported to favour the fixation of adaptive mutations in Drosophila by breaking down linkage disequilibrium (Marais and Charlesworth 2003; Hill and Robertson 2008; Castellano et al. 2016). We analysed this by looking at 50 and 30 categories of recombination rate for *A. thaliana* and *D. melanogaster*, respectively. For Drosophila, we found a significant positive correlation in estimates of *ω*_*a*_ with increasing levels of recombination rate (Kendall’s τ = 0.384, *p* = 0.0029; Figure S1 and File S2), which is consistent with previous studies (Castellano et al. 2016). This was also observed in *A. thaliana* (Kendall’s τ = 0.206, *p* = 0.0343), thus corroborating the effect of recombination in the rate of adaptive evolution.

Previous studies proposed that genes containing more introns are under stronger selective constraints due to the high cost of transcription, especially in highly expressed genes (Castillo-Davis et al. 2002). Hence, we would expect regions with more introns to be under stronger purifying selection. Conversely, by increasing the total gene length, introns might also effectively increase the intra-genic recombination rate, which could in turn increase the efficacy of positive selection and have a positive impact on *ω*_*a*_. To disentangle the two effects, we categorized genes based on their intron content, into 13 and 10 categories for *A. thaliana* and *D. melanogaster* respectively. Results of *ω*_*na*_ showed significant negative correlations in *D. melanogaster* (Kendall’s τ = −0.359, *p* = 0.0876 in *A. thaliana*; τ = −0.867, *p* = 0.0005 in *D. melanogaster*; Figure S2), but we did not find any significant correlation between the number of introns and *ω*_*a*_ (Kendall’s τ = −0.154, *p* = 0.4641 in *A. thaliana*, τ = −0.333, *p* = 0.1797 in *D. melanogaster*; Figure S2 and supplementary File S2). These findings suggest that the effect of the intron content on the rate of protein evolution is essentially due to stronger purifying selection, while having a negligible influence on the rate of adaptive substitutions.

### The impact of protein structure on adaptive evolution

We further explored the impact of three different levels of protein structure (*i.e.* primary, secondary and tertiary) on the rate of adaptive evolution. We first looked at the primary structure by categorizing proteins according to their length. Former studies correlating gene length and *d*_*N*_/*d*_*S*_ have shown that smaller genes evolve more rapidly (Zhang 2000; Lipman et al. 2002; Liao et al. 2006). Here, we investigated if this faster evolution is followed by a higher rate of adaptive substitutions. We performed the analysis with 30 and 50 categories for *A. thaliana* and *D. melanogaster*, respectively. Results show significant negative correlations with protein length for values of ω and *ω*_*na*_ in both species (ω: Kendall’s τ = −0.678, *p* = 1.42e-07 in *A. thaliana*; τ = −0.776, *p* = 1.79e-15 in *D. melanogaster*; *ω*_*na*_: τ = −0.673, *p* = 1.72e-07 in *A. thaliana*; τ = −0.696, *p* = 9.66e-13 in *D. melanogaster*; Figure S3). The same trend was observed for *ω*_*a*_, although it was only significant in *D. melanogaster* (Kendall’s τ = −0.131, *p* = 0.3092 in *A. thaliana*; τ = −0.477, *p* = 9.91e-07 in *D. melanogaster*; Figure S3 and supplementary File S2). These findings suggest that smaller protein-coding regions are indeed under relaxed purifying selection but might also evolve, in some cases, under a higher rate of adaptive substitutions.

The analysis at the secondary structural level showed significant differences in the evolutionary rate between the structural motifs, with loops demonstrating the highest values of ω, followed by α-helices and β-sheets (pairwise comparisons, see Material and Methods; loops/α-helices*: p* = 0.0303, all other estimates: *p* = 0.0101 in *A. thaliana*; *p* = 0.0104 in *D. melanogaster*; Figure 1). When considering adaptive and non-adaptive substitutions separately, this effect was only observed between β-sheets and the two other structural motifs in *ω*_*na*_ in *A. thaliana* (pairwise comparisons; β-sheets/loops: *p* = 0.0101, β-sheets/α-helices: *p* = 0.0303, α-helices/loops: *p* = 0.5960 in *A. thaliana*; β-sheets/loops: *p* = 0.4062, β-sheets/α-helices: *p* = 0.8229, α-helices/loops: *p* = 0.4271 in *D. melanogaster*; Figure 1) and among β-sheets and α-helices in *ω*_*a*_ in both species (pairwise comparisons; β-sheets/loops: *p* = 0.0707, β-sheets/α-helices: *p* = 0.0707, α-helices/loops: *p* = 0.9191 in *A. thaliana*; β-sheets/loops: *p* = 0.0729, β-sheets/α-helices: *p* = 0.0521, α-helices/loops: *p* = 0.9687 in *D. melanogaster*; Figure 1 and supplementary File S3). This implies that the structural motif has an impact on the selective constraints in *A. thaliana* and also contributes to the rate of adaptation in the two species. Previous studies investigating protein tolerance to amino-acid change have similarly shown that loops and turns are the most mutable, followed by α-helices and β-sheets (Goldman et al. 1998; Guo et al. 2004; Choi et al. 2006). Some authors posed this relationship as an outcome of residue exposure (Goldman et al. 1998; Guo et al. 2004), while others associate it to the degree of structural disorder, where ordered proteins are under stronger selective constraint (Choi et al. 2006). In order to clarify this, we further look into the impact of tertiary structure, by exploring the relationship between residue exposure to solvent and intrinsic protein disorder with the rate of adaptive evolution.

**Figure 1.**
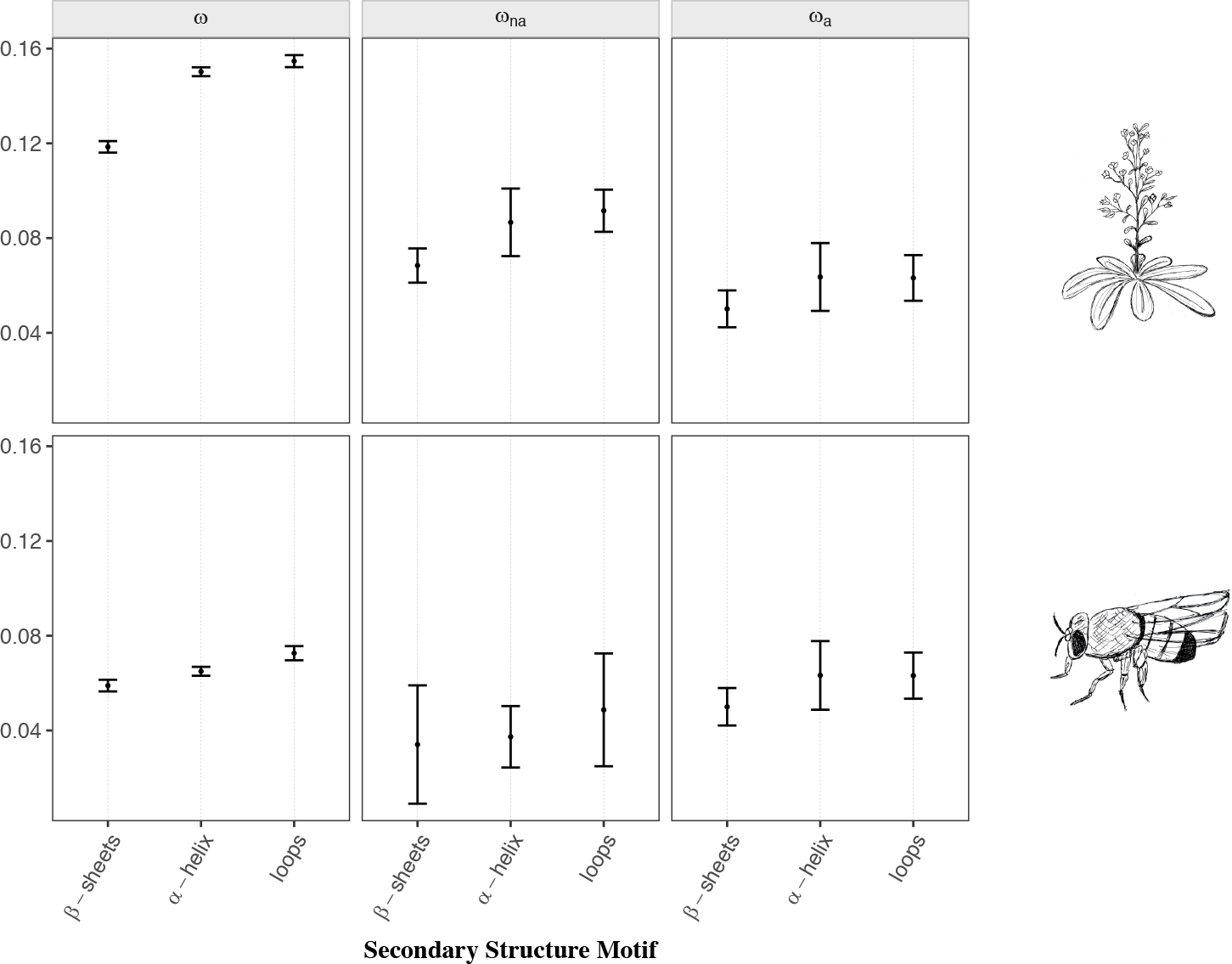
Estimates of the rate of protein evolution (ω), non-adaptive non-synonymous substitutions (*ω*_*na*_) and adaptive non-synonymous substitutions (*ω*_*a*_) for each of the secondary structural motif (β-sheets, α-helices and loops) in *A. thaliana* (top) and *D. melanogaster* (bottom). Mean values of ω, *ω*_*na*_ and *ω*_*a*_ for each motif are represented with the black points. Error bars denote for the 95% confidence interval for each category, computed over 100 bootstrap replicates. The hand-drawings of *A. thaliana* and *D. melanogaster* were made by AFM.

Considering the relative solvent accessibility, several studies previously demonstrated that residues at the surface of proteins evolve faster than the ones at the core (e.g. Goldman et al. 1998; Choi et al. 2007; Lin et al. 2007; Franzosa and Xia 2009). This higher rate of substitution can be either due to a reduced selective constraint at exposed residues and/or to an increased rate of adaptive substitutions. To disentangle the two effects, we compared the site frequency spectra across 28 and 19 categories of RSA in *A. thaliana* and *D. melanogaster* respectively. By using polymorphism data, our results recapitulate those of previous studies on divergence and demonstrate a significant positive correlation with solvent exposure for values of ω (Kendall’s τ = 0.984, *p* = 1.99e-13 in *A. thaliana*; τ = 0.977, *p* = 5.14e-09 in *D. melanogaster*; Figure 2a). Moreover, we demonstrate that both a relaxation of the selective constraints (*ω*_*na*_: Kendall’s τ = 0.846, *p* = 2.58e-10 in *A. thaliana*; τ = 0.579, *p* = 5.33e-04 in *D. melanogaster*; Figure 2a) and a higher rate of adaptive non-synonymous substitutions (*ω*_*a*_: Kendall’s τ = 0.751, *p* = 2.01e-08 in *A. thaliana*; τ = 0.813, *p* = 1.16e-06 in *D. melanogaster*; Figure 2a and supplementary File S2) explain the higher evolutionary rate at the surface of proteins.

**Figure 2.**
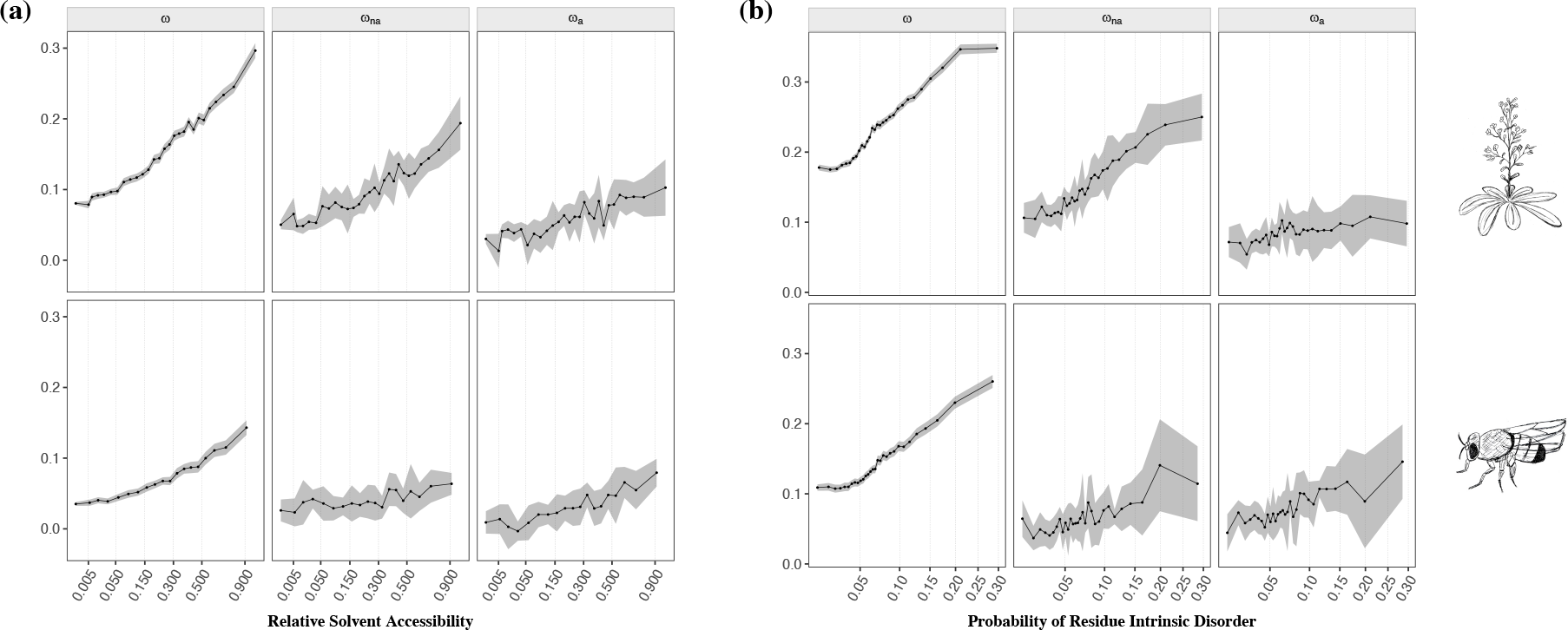
Relationship between ω, *ω*_*na*_ and *ω*_*a*_ with **(a)** the relative solvent accessibility (RSA) and **(b)** the probability of residue intrinsic disorder for *A. thaliana* (top) and *D. melanogaster* (bottom). The x axis is scaled using a squared root function. Mean values of each estimate for each category are represented with connected black dots. The shaded area represents the 95% confidence interval of each category, computed over 100 bootstrap replicates.

Intrinsically disordered proteins are defined by lacking a well-defined three-dimensional fold (Dunker et al. 2002; Dyson and Wright 2005), more specifically, proteins with a higher degree of loop dynamics (“hotloops”) (Linding et al. 2003). As these structures are more flexible we expect them to be under less structural constraint and to accumulate more substitutions (Guo et al. 2004; Wilke et al. 2005; Choi et al. 2006), either deleterious and/or beneficial. To test this hypothesis, we asked two different questions: (1) Are intrinsically disordered protein regions more likely to respond to adaptation? (2) Are proteins with more disordered regions undergoing more adaptive substitutions? For the first question, we divided amino-acid residues into 30 categories based on their predicted value of intrinsic disorder. We report a significant positive correlation with ω, *ω*_*a*_ and *ω*_*na*_ with residue intrinsic disorder for both species (ω: Kendall’s τ = 0.977, *p* = 3.39e-14 in *A. thaliana*; τ = 0.954, *p* = 1.32e-13 in *D. melanogaster*; *ω*_*na*_: τ = 0.917, *p* = 1.09e-12 in *A. thaliana*; τ = 0.670, *p* = 2.08e-07 in *D. melanogaster*; *ω*_*a*_: τ = 0.60, *p* = 3.22e-06 in *A. thaliana*; τ = 0.706, *p* = 4.32e-08 in *D. melanogaster*; Figure 2b and supplementary File S2). For the second question, proteins were categorized according to their proportion of disordered residues (see Material and Methods). Analyses were performed with 30 categories. Results exhibit a significant positive correlation of protein disorder with ω in both species, *ω*_*na*_ in *A. thaliana* and *ω*_*a*_ in *melanogaster* (ω: Kendall’s τ = 0.752, *p* = 5.41e-09 in *A. thaliana*; τ = 0.568, *p* = 4.58e-04 in *D. melanogaster*; *ω*_*na*_: τ = 0.733, *p* = 1.26e-08 in *A. thaliana;* τ = 0.063, *p* = 6.97e-01 in *D. melanogaster*; *ω*_*a*_: τ = 0.191, *p* = 1.38e-01 in *A. thaliana*; τ = 0.726, *p* = 7.56e-06 in *D. melanogaster*; Figure S4). Our findings propose that, at the residue level, intrinsically disordered regions are more likely to respond to adaptation and are also under less selective constraint in both species. However, when considering the whole protein, we observe that intrinsically disordered proteins have different effects between species. In particular, they contribute to the relaxation of purifying selection in *A. thaliana* and to a higher rate of adaptation in *D. melanogaster*. The reason for the difference between species is unclear and will require further analyses.

Finally, we tested whether the rate of adaptive substitutions is affected by the binding affinity of proteins to molecular chaperones. It has been suggested that binding to a chaperone leads to a higher evolutionary rate due to the buffering effect for slightly deleterious mutations (Bogumil and Dagan 2010; Kadibalban et al. 2016). Here, we investigate whether binding to the chaperone *DnaK* could also favour the fixation of adaptive mutations. In agreement with previous studies, we find an effect on ω (pairwise comparisons; *p* = 0.0109 *in A. thaliana*; *p* = 0.0172 in *D. melanogaster*; Figure S5) and *ω*_*na*_ in *D. melanogaster* (pairwise comparisons; *p* = 0.3152 in *A. thaliana*; *p* = 0.0172 in *D. melanogaster*; Figure S5), but no effect on *ω*_*a*_ (pairwise comparisons; *p* > 0.670, FigureS5 and supplementary File S3), suggesting that the interaction with a molecular chaperone does not influence the fixation of beneficial mutations.

### Protein function and adaptive evolution

We further explored the impact of protein function on sequence evolution. To do so, we analysed the effect of gene expression, protein location and protein functional class on the rate of adaptive substitutions. Several studies on both Eukaryote (Pal et al. 2001; Subramanian and Kumar 2004; Wright et al. 2004; Lemos et al. 2005) and Prokaryote (Rocha and Danchin 2004) organisms have shown that highly expressed genes have lower rates of protein sequence evolution. Here we investigated if the lower evolutionary rate is followed by a reduced rate of adaptive substitutions. To do so, we divided genes into 30 and 15 categories for *A. thaliana* and *D. melanogaster* respectively. Our results support previous findings by displaying a significant negative correlation of mean gene expression with estimates of ω and *ω*_*na*_ in both species (ω: Kendall’s τ = −0.894, *p* = 3.92e-12 in *A. thaliana*; τ = −0.771, *p* = 6.11e-05 in *D. melanogaster*; *ω*_*na*_: τ = −0.816, *p* = 2.39e-10 in *A. thaliana*; τ = −0.619, *p* = 1.30e-03 in *D. melanogaster*; Figure 3). Besides, we find that mean gene expression is also significantly negatively correlated with *ω*_*a*_ in *D. melanogaster* (Kendall’s τ = −0.159, *p* = 0.2183 in *A. thaliana*; τ = −0.505, *p* = 8.72e-03 in *D. melanogaster*; Figure 3 and supplementary File S2), suggesting that gene expression also constrains the rate of adaptation, in addition to the well-known effect on purifying selection. It has been hypothesized that the higher selective constraint in highly expressed genes could be driven by the reduced probability of protein misfolding, wherein selection acts by favouring protein sequences that accumulate less translational missense errors (Drummond et al. 2005). Hence, the higher selective pressure to increase stability in highly expressed proteins could also be hampering the fixation of adaptive mutations.

**Figure 3.**
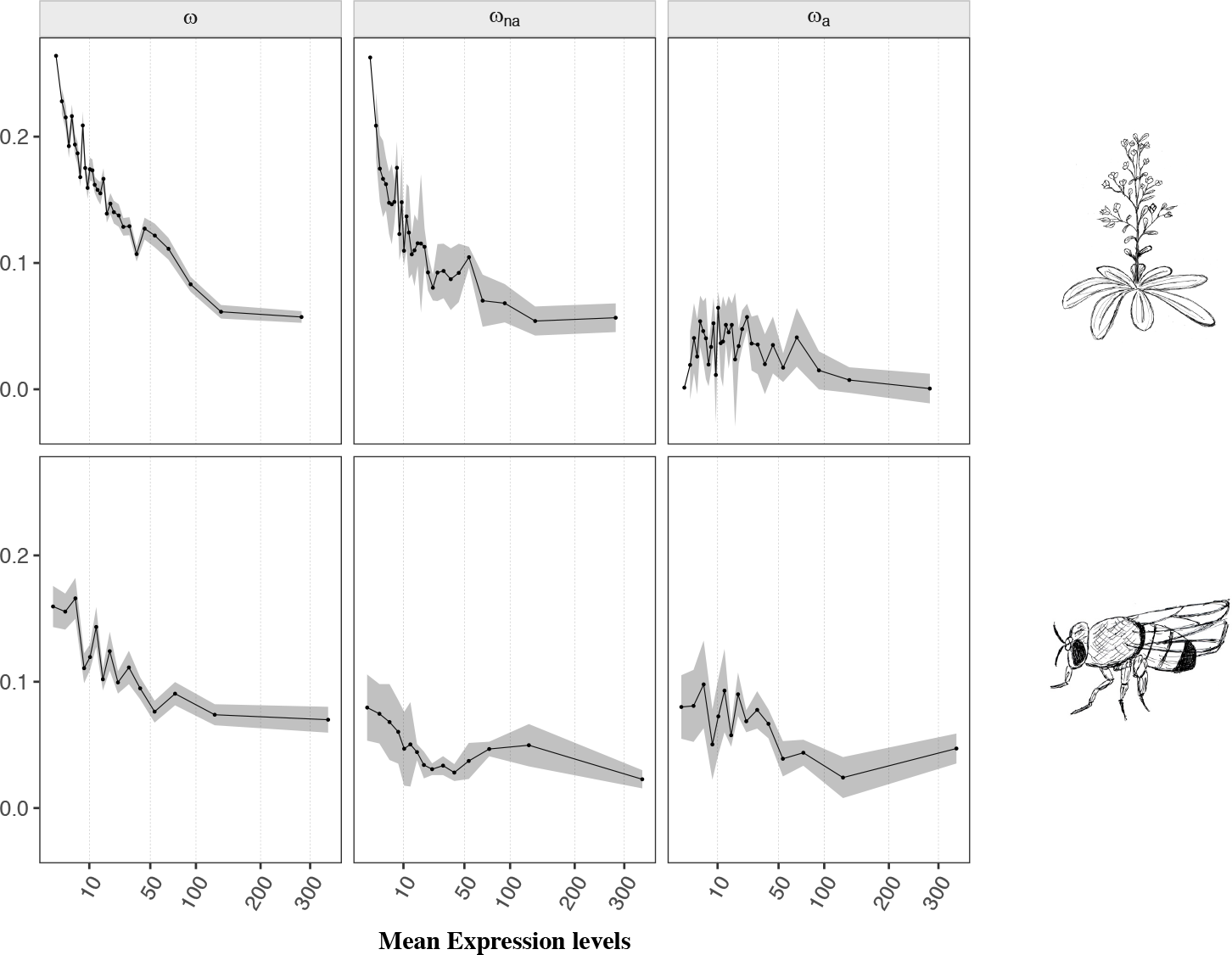
Correlation of ω, *ω*_*na*_ and *ω*_*a*_ with mean gene expression levels for *A. thaliana* (top) and *D. melanogaster* (bottom). The x axis is scaled using a squared root function. Legend as in Figure 2.

In order to assess the impact of protein location we classified genes into the following cellular categories: cytoplasmic, endomembrane system, mitochondrial, nuclear, plasma membrane and secreted proteins (Tables S2 and S3 in supplementary File S1). Results show significantly higher rates of protein evolution in nuclear and secreted proteins, with the lowest values observed in the mitochondria, plasma membrane and endomembrane system (pairwise comparisons; *p* = 0.0128 in *A. thaliana*; *p* = 0.0104 in *melanogaster*; Figure S6). However, this result seems to be explained by a reduced purifying selection, with significantly higher values of *ω*_*na*_ observed in cytoplasmic, nuclear and secreted proteins (pairwise comparisons; *p* = 0.0128 in *A. thaliana*; *p* > 0.0729 in *D. melanogaster*; Figure S6), and not by a higher rate of adaptive substitutions, since no significant differences were found between the categories in the estimates of *ω*_*a*_ (Figure S6 and supplementary File S3).

By analyzing the different categories of protein functional class (Tables S2 and S3 in supplementary File S1), we observe that genes involved in protein biosynthesis (*i.e.* ribosome biogenesis and transcription machinery) and signaling for protein degradation (ubiquitin system) exhibit the highest rates of adaptive substitutions (Figure 4 and supplementary File S4), functions coded mostly by nuclear and cytoplasmic proteins. Signal transduction pathways also appear to play a role in adaptation, since protein phosphatases also present high rates of adaptive mutations (Hunter 1995). Moreover, in *A. thaliana*, cytochrome P450 proteins are also in the top categories of *ω*_*a*_ (Figure 4 and supplementary File S4). We fitted a linear model to the *ω*_*a*_ values of the shared categories (21 categories in total) to see if results were consistent between the two species and found a positive correlation (Kendall’s τ = 0.257, p = 0.1101; Figure S7a), which is stronger after discarding the two outliers, mRNA biogenesis and glycosyltransferases (Kendall’s τ = 0.333, p = 0.0490; Figure S7b). Our findings therefore suggest that adaptive mutations occur mainly through processes of protein regulation and signaling pathways.

**Figure 4.**
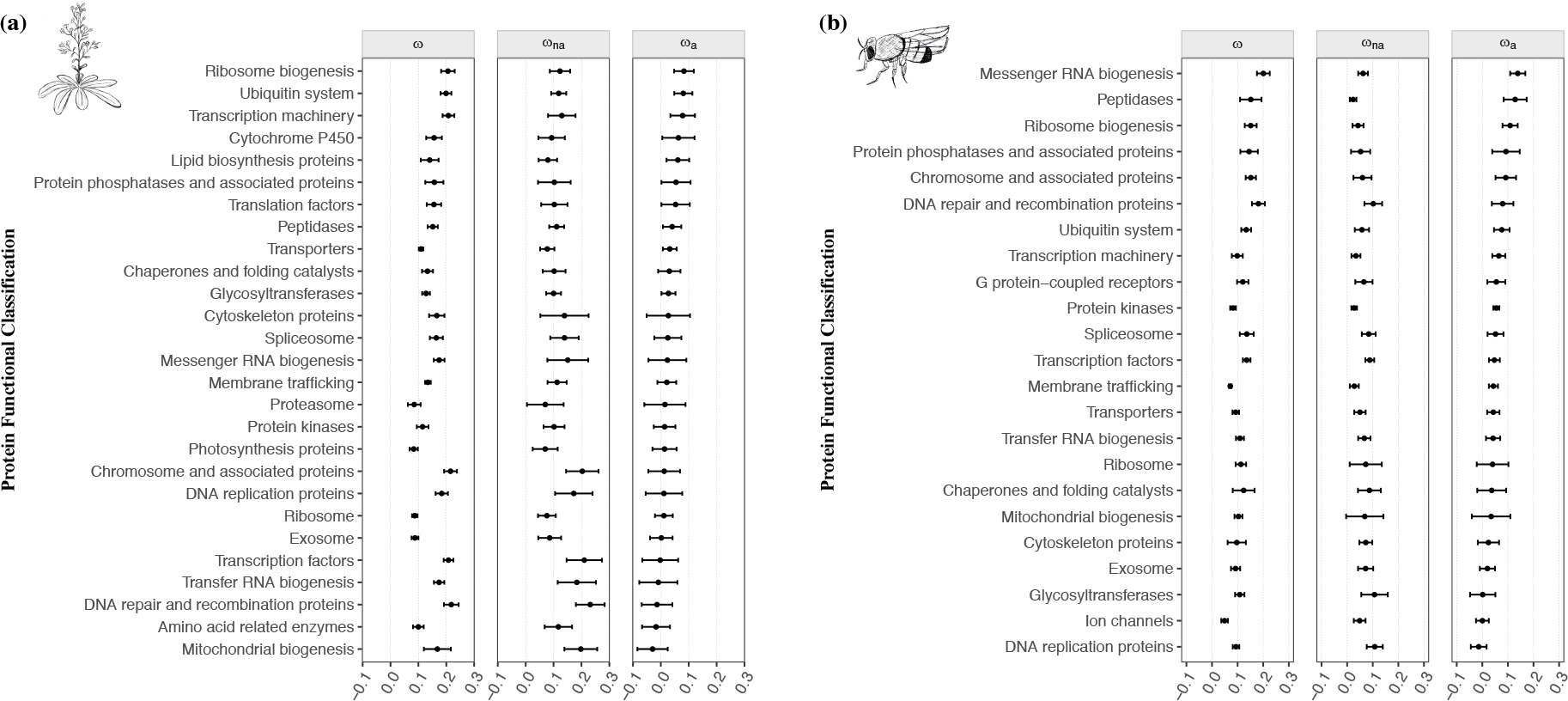
Estimates of ω, *ω*_*na*_ and *ω*_*a*_ for each category of protein functional class in **(a)** *A. thaliana* and **(b)** *D. melanogaster*. Categories are ordered according to the values of *ω*_*a*_. Mean values of ω, *ω*_*na*_ and *ω*_*a*_ for each class are represented with the black points. Error bars denote the 95% confidence interval for each category, computed over 100 bootstrap replicates.

### What are the major drivers of adaptive evolution along the genome?

Overall, we found multiple factors influencing protein adaptive evolution, specifically recombination rate (positive correlation), protein length (negative correlation), secondary structural motif (lower values observed for β-sheets), relative solvent accessibility (positive correlation), protein intrinsic disorder (positive correlation), gene expression levels (negative correlation) and protein functional class. Since some of these variables are intrinsically correlated we next asked whether some of the inferred effects are spurious. First of all, it is known that protein length and gene expression are negatively correlated, wherein highly expressed genes tend to be shorter, as previously reported for vertebrates (Subramanian and Kumar 2004), yeast (Coghlan and Wolfe 2000; Akashi 2003) and observed in this study (Kendall’s τ = −0.015, *p* = 1.22e-02 in *A. thaliana*; τ = −0.093, *p* = 1.70e-28 in *D. melanogaster*; Figure S8). Since highly expressed genes have lower rates of adaptive substitutions and shorter genes have higher rates of adaptive evolution, we may conclude that these two variables independently impact the rate of adaptation in proteins. Protein length is also negatively correlated with the proportion of exposed residues (Kendall’s τ = −0.310, *p* = 0.00 in *A. thaliana*; τ = −0.404, *p* = 1.03e-223 in *D. melanogaster*; Figure S9), as the surface / volume ratio of globular proteins decrease when protein length increases (Janin 1979). By estimating the rate of adaptive mutations of buried and exposed sites separately, we observe that the effect of protein length is no longer significant (*ω*_*a*_-buried: Kendall’s τ = −0.422, *p* = 0.0892 in *A. thaliana*; τ = −0.067, *p* = 0.7884 in *D. melanogaster*; *ω*_*a*_-exposed: τ = −0.289, *p* = 0.2449 in *A. thaliana*; τ = 0.333, *p* = 0.1797 in *D. melanogaster*; Figure 5a and supplementary File S5). This suggests that the effect of protein length on the rate of adaptive substitutions is a by-product of the effect of the residue’s solvent exposure. Furthermore, mean gene expression is positively correlated with solvent exposure (Kendall’s τ = 0.016, *p* = 0.1037 in *A. thaliana*; τ = 0.327, *p* = 4.50e-45 in *D. melanogaster*; Figure S10), as expected since highly expressed genes are shorter and shorter genes have a greater proportion of exposed residues (Figures S8 and S9). These two variables, however, have opposite effects on *ω*_*a*_, and we therefore conclude that gene expression is acting independently from solvent exposure on the rate of adaptive protein evolution.

**Figure 5.**
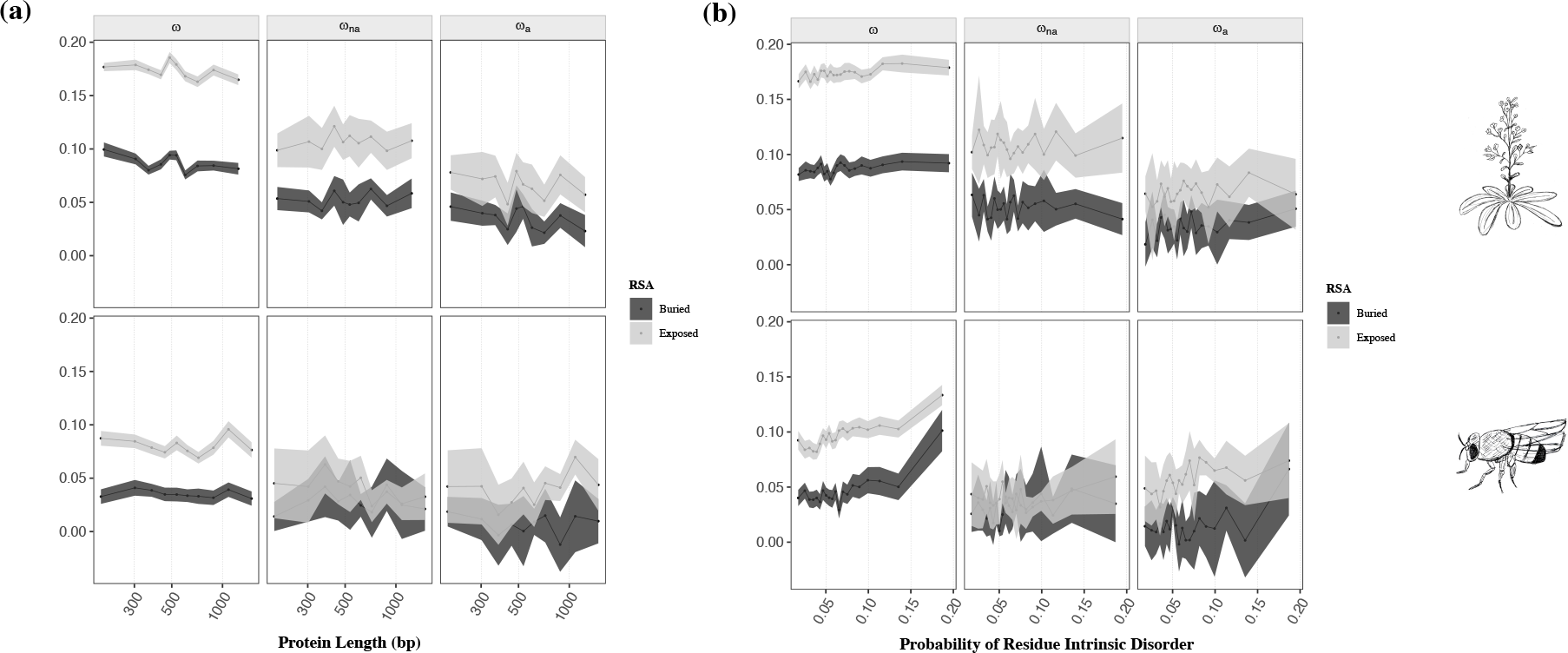
Estimates of ω, *ω*_*na*_ and *ω*_*a*_ for the correlation between **(a)** relative solvent accessibility and protein length and **(b)** relative solvent accessibility and probability of residue intrinsic disorder in *A. thaliana* (top) and *D. melanogaster* (bottom). The x axis is log scaled. Analyses were performed by comparing buried (RSA < 0.05) and exposed (RSA >= 0.05) residues across 10 categories of protein length in (a) and 20 categories of intrinsic disorder in (b) for both species. Legend as in Figure 2.

We further note that the secondary structure motif is intrinsically correlated with the degree of intrinsic disorder, where loops and turns represent the most flexible motifs (Figure S11), consistent with previous studies (Choi et al. 2006). When analysing different degrees of protein disorder across the structural motifs, we observe that secondary structure has only an impact on estimates of ω, while intrinsic protein disorder is significantly positively correlated with ω within the three motifs in both species, and *ω*_*a*_ within β-sheets in *A. thaliana* and within α-helices in *D. melanogaster* (Figure S12 and supplementary File S5). Moreover, we report that the secondary structure motif is correlated with solvent exposure (Figure S13), β-sheets being mostly found at the core of proteins, while α-helices and loops have, on average, higher solvent exposure (Bowie et al. 1990; Guo et al. 2004). By estimating the rate of adaptive substitutions in buried and exposed residues across the three motifs, the impact of secondary structure is no longer noticeable on estimates of *ω*_*a*_ (p > 0.1562; Figure S14 and supplementary File S5), thus suggesting that the effect of secondary structure motif is also a by-product of solvent exposure. When looking at the tertiary structure level, in agreement with Choi et al. (2006), we report that structures with more exposed residues tend to be more flexible (Kendall’s τ = 0.001, *p* = 0.5115 in *A. thaliana*; τ = 0.015, *p* = 0.0222 in *D. melanogaster*; Figure S15). Estimation of the rate of adaptive mutations in buried and exposed sites across different levels of residue intrinsic disorder shows that solvent exposure plays the main role in protein adaptive evolution, with significant impact of protein disorder only observed in values of ω in both species and *ω*_*a*_ in exposed residues for *D. melanogaster* (ω-buried: Kendall’s τ = 0.453, *p* = 5.27e-03 in *A. thaliana;* τ = 0.505, *p* = 1.84e-03 in *D. melanogaster*; ω-exposed: τ = 0.368, *p* = 2.31e-02 in *A. thaliana*; τ = 0.716, *p* = 1.02e-05 in *D. melanogaster*; *ω*_*na*_-buried: τ = −0.063, *p* = 6.97e-01 in *A. thaliana;* τ = 0.295, *p* = 6.92e-02 in *D. melanogaster*; *ω*_*na*_-exposed: τ = −0.0211, *p* = 8.97e-01 in *A. thaliana*; τ = −0.032, *p* = 8.45e-01 in *D. melanogaster*; *ω*_*a*_-buried: τ = 0.211, *p* = 1.94e-01 in *A. thaliana;* τ = 0.084, p = 6.03e-01 in *D. melanogaster*; *ω*_*a*_-exposed: τ = 0.211, *p* = 1.94e-01 in *A. thaliana*; τ = 0.537, *p* = 9.35e-04 in *D. melanogaster*; Figure 5b and supplementary File S5). These findings suggest that the level of exposure of a residue in the protein structure is the main driver of adaptive evolution, and that structural flexibility potentially constitutes a comparatively small, if any, effect to protein adaptation. By comparing the level of exposure of the residues across the different classes of protein function, no differences are observed (Figure S16), thus suggesting that these two variables independently affect the rate of protein adaptation.

Summarizing, our results reveal that besides population genetic processes such as recombination and mutation rate (Marais and Charlesworth 2003; Hill and Robertson 2008; Castellano et al. 2016), three major functional factors significantly impact the rate of protein adaptive evolution: gene expression, relative solvent accessibility and the protein functional class. When looking at the magnitude effect of each of these variables, we observe that exposed residues have a ten-fold higher rate of adaptive substitutions when compared to completely buried sites (Figure 2a and supplementary File S2). The effect of gene expression seems to be of lower magnitude, wherein less expressed genes have a two-fold higher rate of adaptive substitutions with a significant negative correlation observed only in *D. melanogaster* (Figure 3 and supplementary File S2). As a comparison, genes in highly recombining regions have up to a ten-fold higher rate of adaptive substitutions compared to genes within regions with the lowest recombination rates (Figure S1 and supplementary File S2), being therefore similar to that observed with solvent exposure. Previous studies reported that the type of amino-acid change also plays an important role in protein adaptive evolution, where more similar amino-acids present higher rates of adaptive substitutions (Grantham 1974; Miyata et al. 1979; Bergman and Eyre-Walker, personal communication). In order to evaluate a potential bias on the type of amino-acid at the surface and at the core of proteins, we computed the proportion of conservative and radical residue changes, according to volume and polarity indices, as defined by Grantham (Grantham 1974). We found similar frequencies of conserved and radical changes in buried and exposed residues, thus suggesting that our results at the structural level are not influenced by the type of amino-acid mutation (97% of conservative and 3% changes on buried residues; 96% of conservative and 4% changes on exposed sites). Our findings therefore suggest that protein architecture strongly influences the rate of adaptive protein evolution, wherein selection acts by favouring a greater accumulation of adaptive mutations at the surface of proteins.

### What drives protein adaptive evolution?

Our results show that solvent exposure is the functional factor with the highest impact on the rate of adaptive substitutions. To explain this effect, we discuss three hypotheses in which protein adaptive evolution occurs through (1) the acquisition of new biochemical activities at the surface of proteins, (2) the emergence of new functions via network rewiring at the level of protein-protein interactions, and (3) inter-molecule interactions between organisms, as a consequence of host-pathogen coevolution.

We first hypothesized that protein adaptation results from new catalytic activities, wherein adaptive mutations arise within active sites. Barlett et al (2002) reported that active sites are mostly present in more intrinsically disordered regions of the protein. Moreover, they proposed that apoenzymes, which are not yet bound to the substrate or cofactor, present a greater residue flexibility and more exposed catalytic residues, which could favour a higher rate of adaptive substitutions. In order to test this, we estimated the rate of adaptive substitutions on active and non-active sites, controlling for solvent exposure, and observed only significant differences in ω within buried residues in *A. thaliana* (*p* = 0.0309; all other estimates with *p* > 0.1122; Figure S17 and supplementary File S5), although with higher values observed for non-active sites. While the non-significant differences in the rate of adaptive mutations could result from incomplete annotations, which tend to be biased towards motifs highly conserved across species (De Castro et al. 2006), this suggests that being present in an active site does not influence the rate of adaptation. Active sites, however, are rather mobile, presenting different levels of solvent exposure and residue flexibility according to the stage of the enzymatic reaction (Bartlett et al. 2002). Therefore, it may be arbitrary to assign them a certain solvent exposure class based on the phase the enzymes were crystallized, limiting our capacity to test their role on adaptive evolution.

Several studies discussed the impact of protein-protein interactions (PPI) on the rate of protein evolution. Valdar and Thornton (2001) and Caffrey (2004) proposed that PPI may be acting as an inhibitor of protein evolution by enhancing the efficiency of purifying selection due to a higher degree of protein connectivity, typically associated with more complex functions. Mintseris and Weng (2005) supported this assumption but proposed that the proteins evolving slowly are the ones involved in obligate interactions, while proteins involved in transient interactions evolve at faster rates due to a higher interface plasticity. Here, we ask whether the higher rate of adaptive mutations at the surface of proteins could have arisen through inter-molecular interactions at the protein network level. We addressed this question by estimating the rate of adaptive mutations in genes with different degrees of PPI. This was only possible in *D. melanogaster* since there was no data available for *A. thaliana*. We report a negative correlation between the number of PPI and ω, *ω*_*na*_ and *ω*_*a*_, respectively, with only significant values observed for ω (ω: Kendall’s τ = −0.368, *p* = 0.0275; *ω*_*na*_: τ = −0.111, *p* = 0.5062; *ω*_*a*_: τ = −0.309, *p* = 0.0637; Figure S18 and supplementary File S2). These findings suggest that a higher degree of protein connectivity leads to lower rates of protein sequence evolution, but prevent us to assess with confidence whether this effect is due to a stronger purifying selection and/or a slower rate of adaptive substitutions. A potential limitation of this analysis is the low number of genes with PPI information available and the noise associated with the BioGRID annotations. As a physical interaction does not necessarily imply a functional link, we might lack statistical power to detect any putative effect of PPI on *ω*_*a*_ (Chatr-Aryamontri et al. 2017).

In support to our third hypothesis, several studies have described the role of the immune and defense responses in molecular evolution across taxa (Sackton et al. 2007; Obbard et al. 2009; Enard et al. 2016; Mauch-Mani et al. 2017). These studies suggest that pathogens could be key drivers of protein adaptation, by acting as a powerful selective pressure through the coevolutionary arms race between hosts and parasites. This could be driving the higher rate of adaptive mutations in protein biosynthesis enzymes (Figure 4), which are the ones typically hijacked by pathogens during host infection (Dangl and Jones 2001; Enard et al. 2016). Moreover, one of the fastest evolving protein class is the ubiquitin system (Figure 4), which is known to be involved in the defense mechanism, both by the host, through processes like the activation of innate immune responses and degradation signaling of pathogenic proteins; and by the pathogen, which inhibits and/or uses this system in order to modulate host responses (Loureiro and Ploegh 2006; Collins and Brown 2010; Dielen et al. 2010; Trujillo and Shirasu 2010; Hiroshi et al. 2014). Likewise, in *A. thaliana*, cytochrome P450 proteins present a high rate of adaptive mutations (Figure 4), which have been reported to play a crucial role in the defence response in plants (Schuler and Werck-Reichhart 2003). Besides, the reduced selective pressure on nuclear and secreted proteins (Figure S6) may be also a consequence of their role in disease and pathogen immunity (*i.e.* Motion et al. 2015; Mosmann et al. 2016), as observed in yeast (Julenius and Pedersen 2006), insects (Sackton et al. 2007; Obbard et al. 2009) and primates (Nielsen et al. 2005).

Our findings therefore support the hypothesis that coevolutionary arms race of the host-pathogen interactions are a major driver of adaptation in proteins. While we do not rule out that protein-protein interactions and the acquisition of new biochemical functions could also have an impact, more and better annotation data is required to further evaluate their role. In conclusion, our study reveals that, in addition to genome architecture, protein structure has a substantial impact on adaptive evolution consistent between *D. melanogaster* and *A. thaliana*, unraveling the potential generality of such effect. Our study further emphasizes that the rate of adaptation not only varies substantially between genes, but also at the intra-genic scale, and we posit that accounting for a fine-scale, intra-molecular evolution is necessary to fully understand the patterns of molecular adaptation at the species level.

## Materials and Methods

### Population Genomic Data and Data filtering

The *D. melanogaster* data set included alignments of 114 genomes for one chromosome arm of the two large autosomes (2L, 2R, 3L and 3R) and one sex chromosome (X) pooled from 22 sub-Saharan populations with negligible amount of population structure (*F*_*ST*_ = 0.05; DPGP2, Pool et al. 2012). Release 5 of the Berkeley Drosophila Genome Project (BDGP5, http://www.fruitfly.org/sequence/release5genomic.shtml, last updated June 2018) was used as the reference genome. Estimations of divergence were performed with *D. simulans*, for which genome alignments with the reference genome were available (http://www.johnpool.net/genomes.html). For *A. thaliana*, analyses were carried out with 110 genomes for the 5 chromosomes of the Spanish population from the 1001 Genomes Project (Weigel and Mott 2009), using the release 10 from The Arabidopsis Information Resource (TAIR10, ftp://ftp.ensemblgenomes.org/pub/plants/release-40/fasta/arabidopsis_thaliana/dna/) as reference genome. Divergence estimates were made with *A. lyrata* as an outgroup species, for which a pairwise alignment with the reference genome was available (ftp://ftp.ensemblgenomes.org/pub/plants/release-38/maf).

### Estimation of the population genetic parameters and model selection

Coding DNA sequences (CDS) were extracted from the alignments with MafFilter (Dutheil et al. 2014) according to the General Feature Format (GFF) file of the reference genome of both species. First, a cleaning and filtering process was performed to keep only non-overlapping genes with the longest transcript, in cases of multiple transcripts per gene. At this stage, 12,801 and 27,072 genes, for *D. melanogaster* and *A. thaliana* respectively, were kept for further analysis. CDS sequences were then concatenated in order to obtain the full coding region per gene. For the analysis with *A. thaliana*, the alignment of *A. lyrata* with the reference sequence was re-aligned with each gene alignment of the ingroup using MAFFT v7.38 (Katoh and Standley 2013) with the options *add* and *keeplength* so that no gaps were included in the ingroup. Genes with incomplete CDS sequences due to missing information in the outgroup were kept to increase the number of sites for the analysis and gaps were discarded. CDS alignments with premature stop codons were excluded. Final datasets included 10,318 genes for *D. melanogaster/D. simulans* and 18,669 genes for *A. thaliana/A. lyrata*. These datasets were then used to infer both the synonymous and non-synonymous unfolded and folded site frequency spectra (SFS), and synonymous and non-synonymous divergence based on the rate of synonymous and non-synonymous substitutions. All calculations were performed using the BppPopStats program from the Bio++ Program Suite (Guéguen et al. 2013). The Grapes program was then used to compute a genome-wide estimate of the rate of non-adaptive (*ω*_*na*_) and adaptive non-synonymous substitutions (*ω*_*a*_) (Galtier 2016). This method assumes that all sites were sampled in the same number of chromosomes and since some sites were not successfully sampled in all individuals, the original dataset was reduced to 110 and 105 individuals for *D. melanogaster* and *A. thaliana* respectively, by randomly down-sampling polymorphic alleles at each site. The following models were fitted and compared using Akaike’s information criterion: Neutral, Gamma, Gamma-Exponential, Displaced Gamma, Scaled Beta and Bessel K. A model selection procedure was conducted on the two datasets using the complete set of genes for comparison (see Table S1 in supplementary File S1). Following analyses comprised the categorization of genes (see Tables S2 and S3 in supplementary File S1 for detailed information on the genes used for each variable as well as the population genetic parameters estimated per gene for *A. thaliana* and *D. melanogaster* respectively) and amino-acid residues (see Tables S4 and S5 in supplementary File S1 for detailed information on the amino-acid residues used for each category as well as the population genetic parameters estimated per site for *A. thaliana* and *D. melanogaster* respectively) for the different variables analysed and are described below.

### Categorization of gene and genome architecture

Recombination rates were obtained with the R package “MareyMap” (Rezvoy et al. 2007), by using the cubic splines interpolation method. Hereafter we computed the mean recombination rate in cM/Mb units for each gene. Discretization of the observed distribution of recombination rate was performed in 50 and 30 categories with around 350 and 280 genes each for *D. melanogaster* and *A. thaliana* respectively. Intronic information was obtained using the GenomeTools from a GFF with exon annotation and the option *addintrons* (Gremme et al. 2013). Genes were discretized into 10 and 13 categories according to their intron content, for *D. melanogaster* and *A. thaliana* respectively.

### Categorization of protein structure

Genes were discretized according to the total size of the coding region, for which 50 and 30 categories with around 650 and 350 genes each were made for *D. melanogaster* and *A. thaliana* respectively.

In order to obtain structural information for each protein sequence, blastp (Schaffer 2001) was first used to assign each protein sequence to a PDB structure by using the “pdbaa” library and an e-value threshold of 1e-10, allowing only one match per protein sequence. This resulted in 5,008 genes for which a PDB structure was available, making a total of 3,834 PDB structures for *D. melanogaster* and 9,121 genes with a total of 3,832 PDB structures for *A. thaliana*. The corresponding PDB structures were then downloaded and further processed to only keep one chain per polymer. PDB manipulation and analysis were carried on using the R package “bio3d” (Grant et al. 2006). Values for secondary structure (SS) and solvent accessibility (SA) per residue were obtained using the “dssp” program with default options, and were successfully retrieved for 3,613 PDB files corresponding to 4,944 genes for *D. melanogaster* and 3,806 PDB files for a total of 9,106 genes for *A. thaliana*. Subsequently, to map SS and SA values to each residue of the protein sequence a pairwise alignment between each protein and the respective PDB sequence was performed with MAFFT, allowing gaps in both sequences in order to increase the block size of sites aligned. The final data set comprised a total of 1,397,885 sites with SS and SA information, out of 4,821,113 total codon sites obtained with BppPopStats for the complete set of genes of *D. melanogaster*, and 2,585,468 sites mapped with SS and SA information out of 7,479,808 codon sites of *A. thaliana*. We computed the relative solvent accessibility (RSA) by dividing SA by the amino-acid’s solvent accessible area (Tien et al. 2013).

Categorization of secondary structure was performed by comparing 460,702, 975,934 and 523,880 amino-acid residues in β-sheets, α-helices and loops respectively in *A. thaliana*, and 258,898, 516,356 and 282,588 sites in β-sheets, α-helices and loops respectively in *D. melanogaster*. RSA values were analysed with 28 categories with around 85,000 sites each, with the exception of the totally buried residues (RSA = 0) category containing 299,684 sites in *A. thaliana*; and 19 categories with approximately 69,000 residues each, except for 151,417 completely buried residues in *D. melanogaster*. For the analysis of correlation between variables two categories of RSA were considered, comparing buried (RSA < 0.05) and exposed (RSA >= 0.05) residues, following Miller et al (Miller et al. 1987).

Estimates of intrinsic protein disorder were acquired via the software DisEMBL (Linding et al. 2003), wherein intrinsic disorder was estimated per site and classified according to the degree of “hot loops”, meaning loops with a high degree of mobility. This analysis was successfully achieved for a total of 7,479,807 out of 7,479,808 sites for *A. thaliana* and 3,952,602 out of 4,821,113 sites for *D. melanogaster*. Amino-acid residues were divided into 30 categories with an average of 249,000 and 131,000 sites in *A. thaliana* and *D. melanogaster* respectively. For the proportion of disordered regions per protein, we considered a residue “disordered” if it was in the top 25% of the measured probabilities of disorder across the proteomes of each species. Analyses were performed with 30 categories with around 620 and 420 genes for *A. thaliana* and *D. melanogaster* respectively.

### Identification of proteins binding to a molecular chaperone

Prediction of the molecular chaperone *DnaK* binding sites in the protein sequence was estimated with the LIMBO software using the default option *Best overall prediction*. This setting implies 99% specificity and 77.2% sensitivity (Van Durme et al. 2009). Genes were categorized according to this prediction setting, which suggests that every peptide scoring above 11.08 is a predicted *DnaK* binder. Genes scoring below that value were not consider as possible binders.

### Categorization of gene expression

Gene expression data was obtained from the database Expression Atlas (http://www.ebi.ac.uk/gxa; Petryszak et al. 2016), wherein one baseline experiment was used for each species (*D. melanogaster*, E-MTAB-4723; *A. thaliana*, E-GEOD-53197). For both analysis, gene expression levels were averaged across samples and tissues for each gene, ending up with 15 and 30 categories with around 430 and 350 genes each for *A. thaliana* and *D. melanogaster* respectively.

### Protein cellular localization and protein functional class

Cellular localization of each protein sequence was predicted with the software ProtComp (from Softberry, http://www.softberry.com/) with the default options and genes were classified into the following cellular categories: cytoplasmic, endomembrane system, mitochondrial, nuclear, peroxisome, plasma membrane and secreted proteins. The category peroxisome was excluded from further analysis due to the small number of annotated genes (114 and 250 genes in *D. melanogaster* and *A. thaliana* respectively; detailed information in Tables S2 and S3 in supplementary File S1). Protein functional classes were obtained with the Bioconductor package for R “KEGGREST”, using the KEGG BRITE database (Kanehisa et al. 2002). Analysis were carried out with 2,950 and 3,780 genes for *D. melanogaster* and *A. thaliana* respectively, discretized into the highest levels of each of the three top categories of protein classification: metabolism, genetic information processing and signalling and cellular processes (see Tables S2 and S3 in supplementary File S1).

### Enzymatic active sites and protein-protein interactions

In order to check whether a residue was present in an active site, we used the ScanProsite software (De Castro et al. 2006). Datasets included 1,061,876 and 1,870,166 active sites for *D. melanogaster* and *A. thaliana* respectively. All sites that were not predicted by the program were considered as non-active (see Tables S4 and S5 in supplementary File S1). Data on the degree of protein-protein interactions was obtained with the BioGRID database (Chatr-Aryamontri et al. 2017). This was only possible for *D. melanogaster* since there was no corresponding data available for *A. thaliana*. Analyses were carried out with 5,630 genes divided into 19 categories, with 1,114 genes in the first category, and the others ranging from 700 to 130 according to the respective number of interactions (see Tables S2 and S3 in supplementary File S1).

### Estimation of the adaptive and non-adaptive rate of non-synonymous substitutions

For all gene and amino-acid sets, 100 bootstrap replicates were generated by randomly sampling genes or sites in each category. The Grapes program was then run on each category and replicate with the Gamma-Exponential distribution of fitness effects (Galtier 2016). The first step included the removal of replicates for which the distribution of fitness effects parameters was not successfully fitted. For this purpose, we discarded 1% in the maximum and minimum values for the mean and shape parameters of the DFE (see supplementary files for detailed R scripts). Results for ω, *ω*_*na*_ and *ω*_*a*_ were plotted using the R package “ggplot2” (Wickham 2017) (see supplementary files).

### Statistical analyses

Significance for all continuous variables, including protein length, number of introns, gene expression, intrinsic residue disorder, proportion of disordered regions, recombination rate, number of protein-protein interactions and RSA, was assessed through Kendall’s correlation tests. For this, the mean value of the 100 bootstrap replicates was taken for each category (see detailed script as well as all statistical results in supplementary File S2; note: p-values indicated as 0 in this file correspond to a *p* < 1.0e-8). Significance values for discrete variables, comprising binding affinity to *DnaK*, protein location, protein functional class and secondary structure motif, were achieved by estimating the differences between each pair of the categories analysed, by randomly subtracting each bootstrap replicate. Following steps included counting the number of times the differences between categories were below and above 0, which by taking the minimum of those values gives us a statistic that we call k. The two-tailed p was then estimated by applying the following equation: p = (2k + 1)/(N + 1), where N in the number of bootstrap replicates used. For variables comparing more than two categories we corrected the p for multiple testing using the FDR method (Benjamini and Hochberg 1995) as implemented in R (R Core Team 2015) (see detailed script and all statistical results in supplementary Files S3 and S4). Analyses on the correlations between variables are described in supplementary Files S5 and S6.

## Acknowledgements and funding information

The authors thank Adam Eyre-Walker, Guy Sella, Luis-Miguel Chevin and Hinrich Schulenburg for constructive discussions regarding this work. We also thank Joel Alves for his writing suggestions on the manuscript, Joost Schymkowitz and Floor Stam for their help with LIMBO software, David Castellano for sharing the recombination rate data used for comparison and Nicolas Galtier for helping with the Grapes software. JYD acknowledges funding from the Max Planck Society.

## Supplementary material

Supplementary figures S1-S18 and respective legends are attached separately. Supplementary file S1 includes the raw data tables (Tables S2-S5), Table S1 and the R markdowns used to analyse the data (this file will be made public in a gitlab repository after publication). Supplementary files S2-S6 include all scripts, plots and statistical results for all variables analysed. Description of each file is detailed in Material and Methods.

## References

Adams J, Mansfield MJ, Richard DJ, Doxey AC. 2017. Lineage-specific mutational clustering in protein structures predicts evolutionary shifts in function. Bioinformatics. 33(9):1338–1345.

Akaike H. 1973. Maximum likelihood identification of Gaussian autoregressive moving average models. Biometrika 60(2):255–265.

Akashi H. 2003. Translational selection and yeast proteome evolution. Genetics. 164(4):1291–1303.

Bartlett GJ, Porter CT, Borkakoti N, Thornton JM. 2002. Analysis of catalytic residues in enzyme active sites. J Mol Biol. 324(1):105–121.

Benjamini Y, Hochberg Y. 1995. Controlling the False Discovery Rate: A Practical and Powerful Approach to Multiple Testing. J R Stat Soc. 57(1):289–300.

Bergman J, Eyre-Walker A, unpublished data, https://www.biorxiv.org/content/early/2018/07/28/379073, last accessed October 4, 2018.

Bierne N, Eyre-Walker A. 2004. The genomic rate of adaptive amino acid substitution in Drosophila. Mol Biol Evol. 21(7):1350–1360.

Bogumil D, Dagan T. 2010. Chaperonin-dependent accelerated substitution rates in prokaryotes. Genome Biol Evol. 2(1):602–608.

Bowie JU, Reidhaar-Olson JF, Lim WA, Sauer RT. 1990. Deciphering the message in protein sequences: Tolerance to amino acid substitutions. Science 247(4948):1306–1310.

Boyko AR, Williamson SH, Indap AR, Degenhardt JD, Hernandez RD, Lohmueller KE, Adams MD, Schmidt S, Sninsky JJ, Sunyaev SR, et al. 2008. Assessing the evolutionary impact of amino acid mutations in the human genome. PLoS Genet. 4(5): e1000083.

Bustamante CD, Townsend JP, Hartl DL. 2000. Solvent accessibility and purifying selection within proteins of Escherichia coli and Salmonella enterica. Mol Biol Evol. 17(2):301–308.

Caffrey DR. 2004. Are protein-protein interfaces more conserved in sequence than the rest of the protein surface? Protein Sci. 13(1):190–202.

Campos JL, Halligan DL, Haddrill PR, Charlesworth B. 2014. The relation between recombination rate and patterns of molecular evolution and variation in drosophila melanogaster. Mol Biol Evol. 31(4):1010–1028.

Carneiro M, Albert FW, Melo-Ferreira J, Galtier N, Gayral P, Blanco-Aguiar JA, Villafuerte R, Nachman MW, Ferrand N. 2012. Evidence for widespread positive and purifying selection across the european rabbit (oryctolagus cuniculus) genome. Mol Biol Evol. 29(7):1837–1849.

Castellano D, Coronado-Zamora M, Campos JL, Barbadilla A, Eyre-Walker A. 2016. Adaptive evolution is substantially impeded by hill-Robertson interference in drosophila. Mol Biol Evol. 33(2):442–455.

Castillo-Davis CI, Mekhedov SL, Hartl DL, Koonin EV, Kondrashov FA. 2002. Selection for short introns in highly expressed genes. Nat Genet. 31(4):415–418.

Charlesworth B. 1994. The effect of background selection against deleterious mutations on weakly selected, linked variants. Genet Res. 63(3):213–227.

Charlesworth J, Eyre-Walker A. 2006. The rate of adaptive evolution in enteric bacteria. Mol Biol Evol. 23(7):1348–1356.

Chatr-Aryamontri A, Oughtred R, Boucher L, Rust J, Chang C, Kolas NK, O’Donnell L, Oster S, Theesfeld C, Sellam A, et al. 2017. The BioGRID interaction database: 2017 update. Nucleic Acids Res. 45(D1):D369–D379.

Choi SC, Hobolth A, Robinson DM, Kishino H, Thorne JL. 2007. Quantifying the impact of protein tertiary structure on molecular evolution. Mol Biol Evol. 24(8):1769–1782.

Choi SS, Vallender EJ, Lahn BT. 2006. Systematically assessing the influence of 3-dimensional structural context on the molecular evolution of mammalian proteomes. Mol Biol Evol. 23(11):2131–2133.

Coghlan A, Wolfe KH. 2000. Relationship of codon bias to mRNA and concentration protein length in Saccharomyces cerevisiae. Yeast. 16(12):1131–1145.

Collins CA, Brown EJ. 2010. Cytosol as battleground: Ubiquitin as a weapon for both host and pathogen. Trends Cell Biol. 20(4):205–213.

Conant GC, Stadler PF. 2009. Solvent exposure imparts similar selective pressures across a range of yeast proteins. Mol Biol Evol. 26(5):1155–1161.

Dangl JL, Jones JD. 2001. Plant pathogens and integrated defence responses to infection. Nature. 411(6839):826–833.

De Castro E, Sigrist CJA, Gattiker A, Bulliard V, Langendijk-Genevaux PS, Gasteiger E, Bairoch A, Hulo N. 2006. ScanProsite: Detection of PROSITE signature matches and ProRule-associated functional and structural residues in proteins. Nucleic Acids Res. 34(2): W362–W365.

Dean AM, Neuhauser C, Grenier E, Golding GB. 2002. The pattern of amino acid replacements in α/β-barrels. Mol Biol Evol. 19(11):1846–1864.

Dielen AS, Badaoui S, Candresse T, German-Retana S. 2010. The ubiquitin/26S proteasome system in plant-pathogen interactions: A never-ending hide-and-seek game. Mol Plant Pathol. 11(2):293–308.

Drummond DA, Bloom JD, Adami C, Wilke CO, Arnold FH. 2005. Why highly expressed proteins evolve slowly. Proc Natl Acad Sci. 102(40):14338–14343.

Dunker AK, Brown CJ, Lawson JD, Iakoucheva LM, Obradović Z. 2002. Intrinsic disorder and protein function. Biochemistry. 41(21):6573–6582.

Dutheil JY, Gaillard S, Stukenbrock EH. 2014. MafFilter: A highly flexible and extensible multiple genome alignment files processor. BMC Genomics. 15(1):53.

Dyson HJ, Wright PE. 2005. Intrinsically unstructured proteins and their functions. Nat Rev Mol Cell Biol. 6(3):197–208.

Enard D, Cai L, Gwennap C, Petrov DA. 2016. Viruses are a dominant driver of protein adaptation in mammals. Elife. 5: e12469.

Eyre-Walker A. 2002. Changing effective population size and the McDonald-Kreitman test. Genetics. 162(4):2017–2024.

Eyre-Walker A. 2006. The genomic rate of adaptive evolution. Trends Ecol Evol. 21(10):569–75.

Eyre-Walker A, Keightley PD. 2009. Estimating the rate of adaptive molecular evolution in the presence of slightly deleterious mutations and population size change. Mol Biol Evol. 26(9):2097–2108.

Eyre-Walker A, Woolfit M, Phelps T. 2006. The distribution of fitness effects of new deleterious amino acid mutations in humans. Genetics. 173(2):891–900.

Fay JC, Wyckoff GJ, Wu CI. 2001. Positive and negative selection on the human genome. Genetics. 158(3):1227–34.

Franzosa EA, Xia Y. 2009. Structural determinants of protein evolution are context-sensitive at the residue level. Mol Biol Evol. 26(10):2387–2395.

Galtier N. 2016. Adaptive Protein Evolution in Animals and the Effective Population Size Hypothesis. PLoS Genet. 12(1): e1005774.

Goldman N, Thorne JL, Jones DT. 1998. Assessing the Impact of Secondary Structure and Solvent Accessibility on Protein Evolution. Genetics. 149(1):445–458.

Gossmann TI, Keightley PD, Eyre-Walker A. 2012. The effect of variation in the effective population size on the rate of adaptive molecular evolution in eukaryotes. Genome Biol Evol. 4(5):658–667.

Gossmann TI, Song BH, Windsor AJ, Mitchell-Olds T, Dixon CJ, Kapralov M V., Filatov DA, Eyre-Walker A. 2010. Genome wide analyses reveal little evidence for adaptive evolution in many plant species. Mol Biol Evol. 27(8):1822–1832.

Grant BJ, Rodrigues APC, ElSawy KM, McCammon JA, Caves LSD. 2006. Bio3d: An R package for the comparative analysis of protein structures. Bioinformatics. 22(21):2695–2696.

Grantham R. 1974. Amino Acid Difference Formula to Help Explain Protein Evolution. Science. 185(4154):862–4.

Gremme G, Steinbiss S, Kurtz S. 2013. Genome tools: A comprehensive software library for efficient processing of structured genome annotations. IEEE/ACM Trans Comput Biol Bioinforma. 10(3):645–656.

Guéguen L, Gaillard S, Boussau B, Gouy M, Groussin M, Rochette NC, Bigot T, Fournier D, Pouyet F, Cahais V, et al. 2013. Bio++: Efficient extensible libraries and tools for computational molecular evolution. Mol Biol Evol. 30(8):1745–1750.

Guo HH, Choe J, Loeb LA. 2004. Protein tolerance to random amino acid change. Proc Natl Acad Sci. 101(25):9205–9210.

Haddrill PR, Loewe L, Charlesworth B. 2010. Estimating the parameters of selection on nonsynonymous mutations in Drosophila pseudoobscura and D. miranda. Genetics. 185(4):1381–1396.

Haerty W, Jagadeeshan S, Kulathinal RJ, Wong A, Ram KR, Sirot LK, Levesque L, Artieri CG, Wolfner MF, Civetta A, et al. 2007. Evolution in the fast lane: Rapidly evolving sex-related genes in Drosophila. Genetics. 177(3):1321–1335.

Halligan DL, Oliver F, Eyre-Walker A, Harr B, Keightley PD. 2010. Evidence for pervasive adaptive protein evolution in wild mice. PLoS Genet. 6(1): e1000825.

Hill WG, Robertson A. 2008. The effect of linkage on limits to artificial selection. Genetics Research 8(3):269–294.

Hiroshi A, Minsoo K, Chihiro S. 2014. Exploitation of the host ubiquitin system by human bacterial pathogens. Nat Rev Microbiol. 12(1):399–413.

Hunter T. 1995. Protein Kinases and Phosphatases: The Yin and Yang of Protein Phosphorylation and Signaling. Cell. 80:225–236.

Hvilsom C, Qian Y, Bataillon T, Li Y, Mailund T, Salle B, Carlsen F, Li R, Zheng H, Jiang T, et al. 2012. Extensive X-linked adaptive evolution in central chimpanzees. Proc Natl Acad Sci. 109(6):2054–2059.

Ingvarsson PK. 2010. Natural Selection on Synonymous and Nonsynonymous Mutations Shapes Patterns of Polymorphism in Populus tremula. Mol Biol Evol. 27(3):650–660.

Janin J. 1979. Surface and inside volumes in globular proteins. Nature 277(5696):491.

Julenius K, Pedersen AG. 2006. Protein evolution is faster outside the cell. Mol Biol Evol. 23(11):2039–2048.

Kadibalban AS, Bogumil D, Landan G, Dagan T. 2016. DnaK-Dependent Accelerated Evolutionary Rate in Prokaryotes. Genome Biol Evol. 8(5):1590–1599.

Kanehisa M, Goto S, Kawashima S, Nakaya A. 2002. The KEGG databases at GenomeNet. Nucleic Acids Res. 30(1):42–46.

Katoh K, Standley DM. 2013. MAFFT multiple sequence alignment software version 7: Improvements in performance and usability. Mol Biol Evol. 30(4):772–780.

Lemos B, Bettencourt BR, Meiklejohn CD, Hartl DL. 2005. Evolution of proteins and gene expression levels are coupled in Drosophila and are independently associated with mRNA abundance, protein length, and number of protein-protein interactions. Mol Biol Evol. 22(5):1345–1354.

Liao BY, Scott NM, Zhang J. 2006. Impacts of gene essentiality, expression pattern, and gene compactness on the evolutionary rate of mammalian proteins. Mol Biol Evol. 23(11):2072–2080.

Liberles D, Teichmann S, Bahar I, Bastolla U, Bloom J, Bornberg-Bauer E, Colwell LJ, De Koning APJ, Dokholyan NV, Echave J, et al. 2012. The interface of protein structure, protein biophysics, and molecular evolution. Protein Sci. 21(6):769–785.

Lin YS, Hsu WL, Hwang JK, Li WH. 2007. Proportion of solvent-exposed amino acids in a protein and rate of protein evolution. Mol Biol Evol. 24(4):1005–1011.

Linding R, Jensen LJ, Diella F, Bork P, Gibson TJ, Russell RB. 2003. Protein disorder prediction: Implications for structural proteomics. Structure 11(11):1453–1459.

Lipman DJ, Souvorov A, Koonin EV, Panchenko AR, Tatusova TA. 2002. The relationship of protein conservation and sequence length. BMC Evol Biol. 2:1–10.

Loureiro J, Ploegh HL. 2006. Antigen Presentation and the Ubiquitin-Proteasome System in Host–Pathogen Interactions. Adv Immunol. 92:225–305.

Marais G, Charlesworth B. 2003. Genome evolution: Recombination speeds up adaptive evolution. Curr Biol. 13(2):68–70.

Mauch-Mani B, Baccelli I, Luna E, Flors V. 2017. Defense Priming: An Adaptive Part of Induced Resistance. Annu Rev Plant Biol. 68(1):485–512.

McDonald JH, Kreitman M. 1991. Adaptive protein evolution ate the Adh locus in Drosophila. Nature 351(6328):652–4.

Miller S, Lesk AM, Janin J, Chothia C. 1987. The accessible surface area and stability of oligomeric proteins. Nature. 328(6133):834.

Mintseris J, Weng Z. 2005. Structure, function, and evolution of transient and obligate protein-protein interactions. Proc Natl Acad Sci. 102(31):10930–10935.

Mirny LA, Shakhnovich EI. 1999. Universally conserved positions in protein folds: reading evolutionary signals about stability, folding kinetics and function. J Mol Biol. 291(1):177–196.

Miyata T, Miyazawa S, Yasunaga T. 1979. Two Types of Amino Acid Substitutions in Protein Evolution. J Mol Evol. 12(3):219–236.

Mosmann VR, Cherwinski H, Bond MW, Giedlin MA, Coffman RL. 2016. Two types of murine helper T cell clone. I. Definition according to profiles of lymphokine activities and secreted proteins. J Immunol. 136(7):2348–2357.

Motion GB, Amaro T, Kulagina N, Huitema E. 2015. Nuclear processes associated with plant immunity and pathogen susceptibility. Brief Funct Genomics. 14(4):243–252.

Nielsen R, Bustamante C, Clark AG, Glanowski S, Sackton TB, Hubisz MJ, Fledel-Alon A, Tanenbaum DM, Civello D, White TJ, et al. 2005. A scan for positively selected genes in the genomes of humans and chimpanzees. PLoS Biol. 3(6):0976–0985.

Obbard DJ, Welch JJ, Kim KW, Jiggins FM. 2009. Quantifying adaptive evolution in the Drosophila immune system. PLoS Genet. 5(10): e1000698.

Overington J, Donnelly D, Johnson MS, Sali A, Blundell TL. 1992. Environment-specific amino acid substitution tables: Tertiary templates and prediction of protein folds. Protein Sci. 1:216–226.

Pal C, Papp B, Hurst LD. 2001. Highly Expressed Genes in Yeast Evolve Slowly. Genetics. 158(1998):927–931.

Perutz MF, Kendrew JC, Watson HC. 1965. Structure and function of haemoglobin: II. Some relations between polypeptide chain configuration and amino acid sequence. J Mol Biol. 13(3):669–678.

Petryszak R, Keays M, Tang YA, Fonseca NA, Barrera E, Burdett T, Füllgrabe A, Fuentes AMP, Jupp S, Koskinen S, et al. 2016. Expression Atlas update - An integrated database of gene and protein expression in humans, animals and plants. Nucleic Acids Res. 44(D1):D746–D752.

Pool JE, Corbett-Detig RB, Sugino RP, Stevens KA, Cardeno CM, Crepeau MW, Duchen P, Emerson JJ, Saelao P, Begun DJ, et al. 2012. Population Genomics of Sub-Saharan Drosophila melanogaster: African Diversity and Non-African Admixture. PLoS Genet. 8(12): e1003080.

Pröschel M, Zhang Z, Parsch J. 2006. Widespread adaptive evolution of Drosophila genes with sex-biased expression. Genetics. 174(2):893–900.

Proux E, Studer RA, Moretti S, Robinson-Rechavi M. 2009. Selectome: A database of positive selection. Nucleic Acids Res. 37(1):404–407.

Ramsey DC, Scherrer MP, Zhou T, Wilke CO. 2011. The relationship between relative solvent accessibility and evolutionary rate in protein evolution. Genetics. 188(2):479–488.

Rezvoy C, Charif D, Guéguen L, Marais GAB. 2007. MareyMap: An R-based tool with graphical interface for estimating recombination rates. Bioinformatics 23(16):2188–2189.

Rocha EPC, Danchin A. 2004. An Analysis of Determinants of Amino Acids Substitution Rates in Bacterial Proteins. Mol Biol Evol. 21(1):108–116.

Sackton TB, Lazzaro BP, Schlenke TA, Evans JD, Hultmark D, Clark AG. 2007. Dynamic evolution of the innate immune system in Drosophila. Nat Genet. 39(12):1461–1468.

Sawyer SA, Kulathinal RJ, Bustamante CD, Hartl DL. 2003. Bayesian Analysis Suggests that Most Amino Acid Replacements in Drosophila Are Driven by Positive Selection. J Mol Evol. 57(1): S154–S164.

Schaffer AA. 2001. Improving the accuracy of PSI-BLAST protein database searches with composition-based statistics and other refinements. Nucleic Acids Res. 29(14):2994–3005.

Schuler MA, Werck-Reichhart D. 2003. Functional Genomics of P450S. Annu Rev Plant Biol. 54(1):629–667.

Slotte T, Foxe JP, Hazzouri KM, Wright SI. 2010. Genome-wide evidence for efficient positive and purifying selection in capsella grandiflora, a plant species with a large effective population size. Mol Biol Evol. 27(8):1813–1821.

Smith NGC, Eyre-Walker a. 2002. Adaptive protein evolution in Drosophila. Nature. 415(6875):1022.

Stoletzki N, Eyre-Walker A. 2011. Estimation of the neutrality index. Mol Biol Evol. 28(1):63–70.

Strasburg JL, Kane NC, Raduski AR, Bonin A, Michelmore R, Rieseberg LH. 2011. Effective population size is positively correlated with levels of adaptive divergence among annual sunflowers. Mol Biol Evol. 28(5):1569–1580.

Subramanian S, Kumar S. 2004. Gene expression intensity shapes evolutionary rates of the proteins encoded by the vertebrate genome. Genetics. 168(1):373–381.

Tataru P, Mollion M, Glémin S, Bataillon T. 2017. Inference of Distribution of Fitness Effects and. Genetics. 207(3):1103–1119.

Tien MZ, Meyer AG, Sydykova DK, Spielman SJ, Wilke CO. 2013. Maximum allowed solvent accessibilites of residues in proteins. PLoS One. 8(11): e80635.

Trujillo M, Shirasu K. 2010. Ubiquitination in plant immunity. Curr Opin Plant Biol. 13(4):402–408.

Valdar WSJ, Thornton JM. 2001. Protein-protein interfaces: Analysis of amino acid conservation in homodimers. Proteins Struct Funct Genet. 42(1):108–124.

Van Durme J, Maurer-Stroh S, Gallardo R, Wilkinson H, Rousseau F, Schymkowitz J. 2009. Accurate prediction of DnaK-peptide binding via homology modelling and experimental data. PLoS Comput Biol. 5(8): e1000475.

Weigel D, Mott R. 2009. The 1001 Genomes Project for Arabidopsis thaliana. Genome Biol. 8(12): e1003080.

Wickham, H. 2017. ggplot2: elegant graphics for data analysis (2nd Edition). J. Stat. Softw. 35(1):65–88.

Wilke CO, Bloom JD, Drummond DA, Raval A. 2005. Predicting the tolerance of proteins to random amino acid substitution. Biophys J. 89(6):3714–20.

Wright SI, Yau CBK, Looseley M, Meyers BC. 2004. Effects of gene expression on molecular evolution in Arabidopsis thaliana and Arabidopsis lyrata. Mol Biol Evol. 21(9):1719–1726.

Zhang J. 2000. Protein-length distributions for the three domains of life. Trends Genet. 16(3):107–109.

