## supplementary File S6 for "The impact of protein architecture on adaptive evolution"

### Relationship between variables

This notebook describes the R scripts used to analyze the relationship between variables. It will include the relationships between:

Continuous variables:
files.continuous[1] <- Protein length/Gene expression
files.continuous[2] <- RSA/Residue intrinsic disorder
files.continuous[3] <- RSA/Gene Expression
files.continuous[4] <- RSA/Protein length

Continuous/Discrete:
files.discrete[1] <- Protein Functional Class/RSA
files.discrete[2] <- Intrinsic Disorder/Secondary structure
files.discrete[3] <- Secondary Structure/RSA

The first chunk describes the analyses performed for comparing continuous variables.

```
setwd("/Users/moutinho/Dropbox/Data/VariablesRelationship/Continuous/")

# Libraries
library(plyr)
library(dplyr)
library(data.table)
library(ggplot2)
library(reshape2)
library(stringr)
library(knitr)
library(kableExtra)
#

# calling all output tables
files.continuous <- list.files(".", ".csv")

# reading each table into a list
tbl.list <- lapply(files.continuous, fread, header = TRUE)
```

```
# now plotting the relationship
# theme of the plot
theme.plot <- function(x) {
  theme(axis.title = element_text(face = "bold", color = "black", size=12,
                                  family = "Times"),
        text = element_text(size=12),
        axis.title.x = element_text(margin = margin(t = 9, r = 10, b = 0, l = 0)),
        axis.title.y = element_text(margin = margin(t = 9, r = 10, b = 0, l = 0)),
        panel.grid.minor=element_blank(),
        panel.grid.major = element_line(colour = "grey", linetype = "dashed", size = 0.1),
        panel.grid.major.y=element_blank(),
        strip.text.y = element_blank(),
        axis.text.x = element_text(angle = 60, hjust = 1),
        strip.background.x = element_rect(colour = "grey", fill = "gray92"),
        legend.text = element_text(size = 7),
        legend.title = element_text(size = 9, face = "bold"))
}
```

```r
# will do the plots seperately due to the different scales
# proten length/gene expression
plot.len.exp <- ggplot(tbl.list[[1]], aes(med.var1, med.var2)) +
  geom_line(col = "black", size = 0.4) +
  geom_ribbon(aes(ymax=thirdQ, ymin=firstQ), alpha=0.2) +
  geom_point(size=.6) +
  geom_quantile(aes(var1, var2), quantiles = 0.5) +
  facet_grid(rows = vars(species)) +
  xlab(as.character(unique(str_split_fixed(tbl.list[[1]]$var, "\\-", 2)[,2]))) +
  ylab(as.character(unique(str_split_fixed(tbl.list[[1]]$var, "\\-", 2)[,1]))) +
  scale_x_log10(limits = c(200, 15000)) +
  theme_bw() +
  theme.plot()

plot.len.exp
```

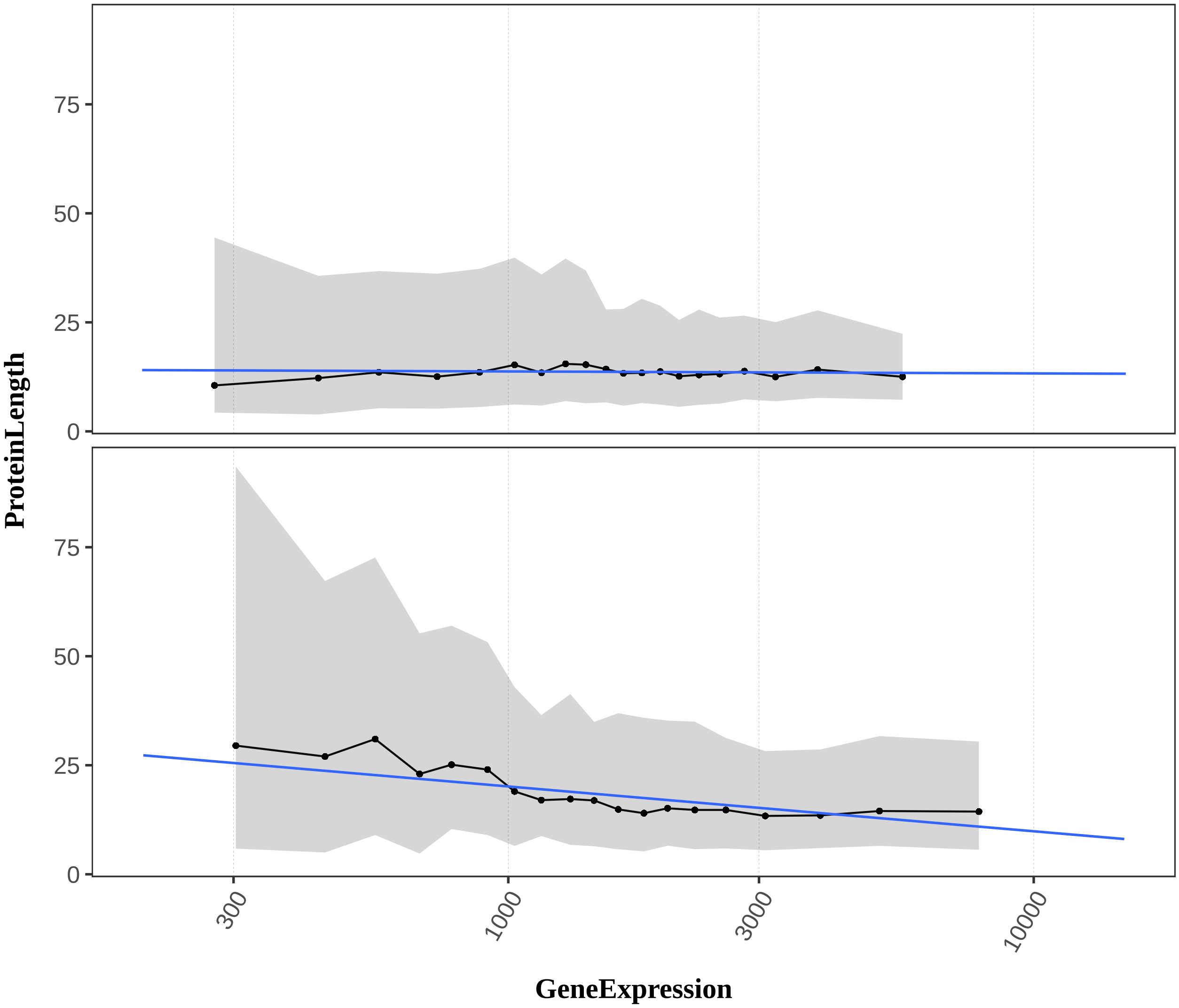


```r
# residue intrinsic disorder/rsa
# sampled random 10000 rows to plot the linear model
```

```
sub.dis.rsa <- tbl.list[[2]][sample(nrow(tbl.list[[2]]), 10000), ]
plot.dis.rsa <- ggplot(tbl.list[[2]], aes(med.var1, med.var2)) +
  geom_line(col = "black", size = 0.4) +
  geom_ribbon(aes(ymax=thirdQ, ymin=firstQ), alpha=0.2) +
  geom_point(size=.6) +
  geom_quantile(data = sub.dis.rsa,
                aes(var1, var2),
                quantiles = 0.5) +
  facet_grid(rows = vars(species)) +
  xlab(as.character(unique(str_split_fixed(tbl.list[[2]]$var, "\\-", 2)[,2]))) +
  ylab(as.character(unique(str_split_fixed(tbl.list[[2]]$var, "\\-", 2)[,1]))) +
  scale_x_log10(limits = c(0.015, 0.22)) +
  theme_bw() +
  theme.plot()

plot.dis.rsa
```

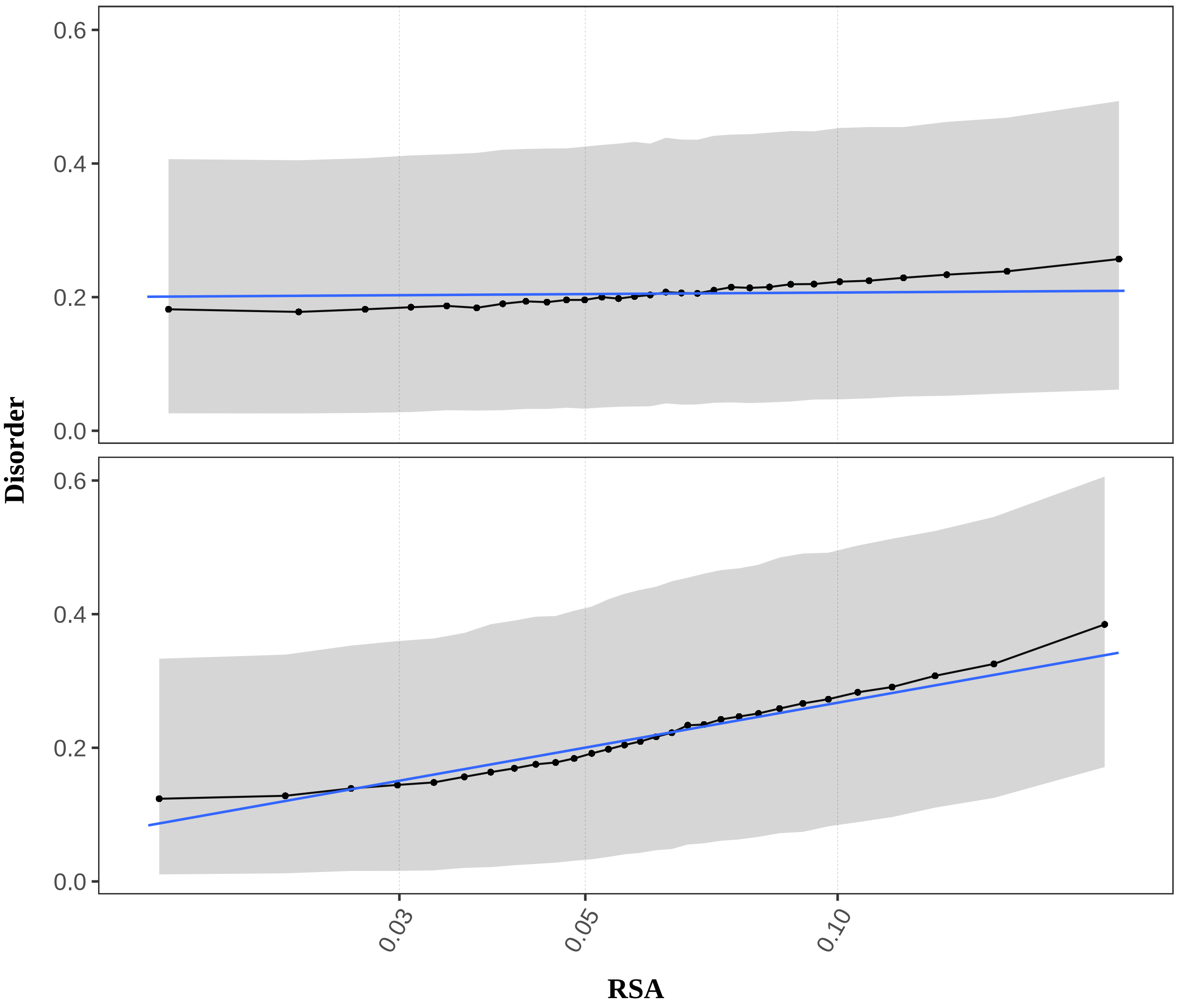


```
# mean gene expression/solvent exposure
plot.rsa.exp <- ggplot(tbl.list[[3]], aes(med.var1, med.var2)) +
  geom_line(col = "black", size = 0.4) +
```

```
  geom_ribbon(aes(ymax=thirdQ, ymin=firstQ), alpha=0.2) +
  geom_point(size=.6) +
  geom_quantile(aes(var1, var2),
               quantiles = 0.5) +
  facet_grid(rows = vars(species)) +
  xlab(as.character(unique(str_split_fixed(tbl.list[[3]]$var, "\\-", 2)[,1]))) +
  ylab(as.character(unique(str_split_fixed(tbl.list[[3]]$var, "\\-", 2)[,2]))) +
  scale_x_continuous(limits = c(0, 180)) +
  theme_bw() +
  theme.plot()

plot.rsa.exp
```

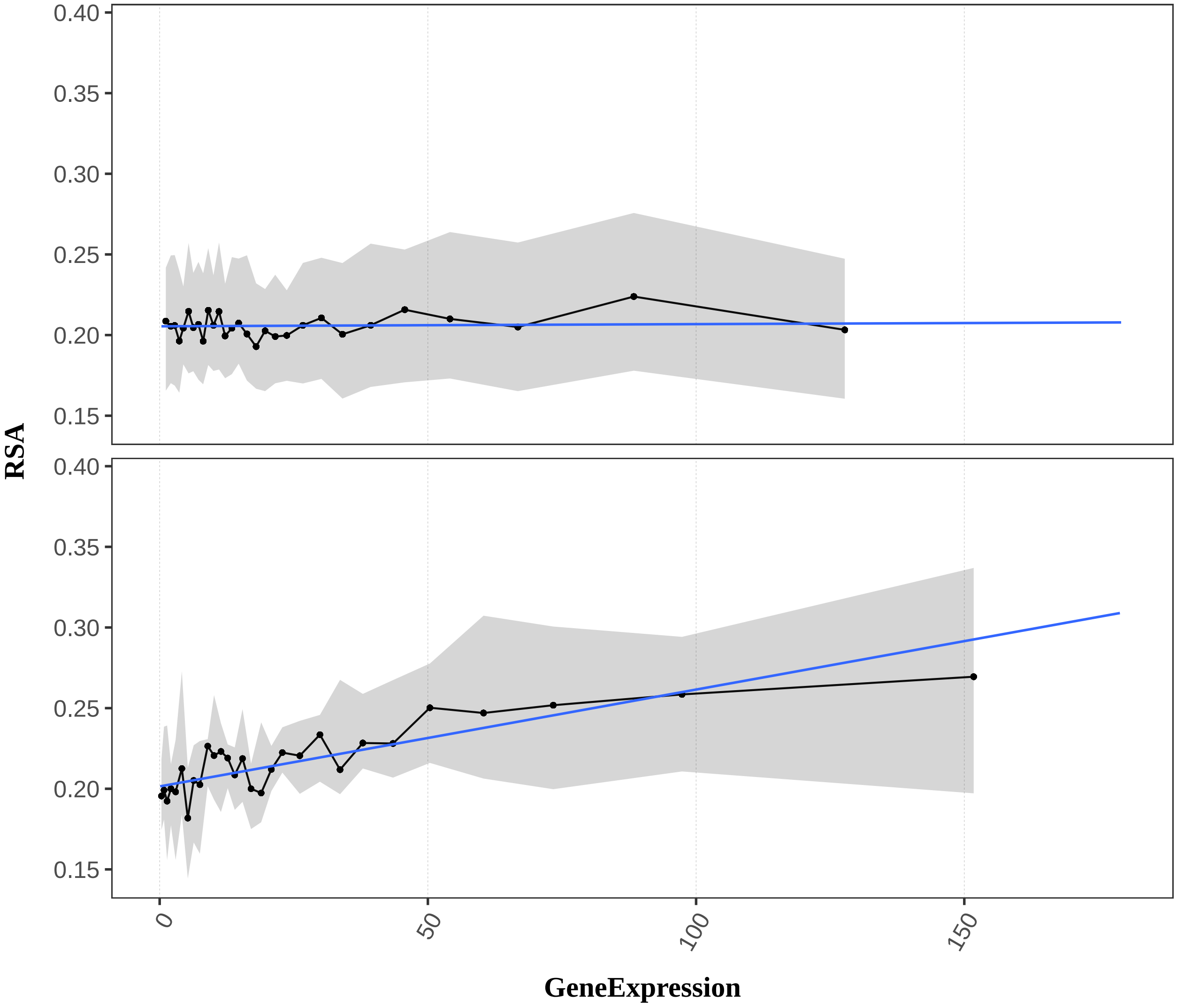


```
# protein length/proportion of exposed residues
plot.rsa.length <- ggplot(tbl.list[[4]], aes(med.var1, med.var2)) +
  geom_line(col = "black", size = 0.4) +
  geom_ribbon(aes(ymax=thirdQ, ymin=firstQ), alpha=0.2) +
  geom_point(size=.6) +
  geom_quantile(aes(var1, var2),
               quantiles = 0.5) +
```

```
  facet_grid(rows = vars(species)) +
  xlab(as.character(unique(str_split_fixed(tbl.list[[4]]$var, "\\-", 2)[,1]))) +
  ylab(as.character(unique(str_split_fixed(tbl.list[[4]]$var, "\\-", 2)[,2]))) +
  scale_x_log10(limits = c(100, 2000)) +
  theme_bw() +
  theme.plot()

plot.rsa.length
```

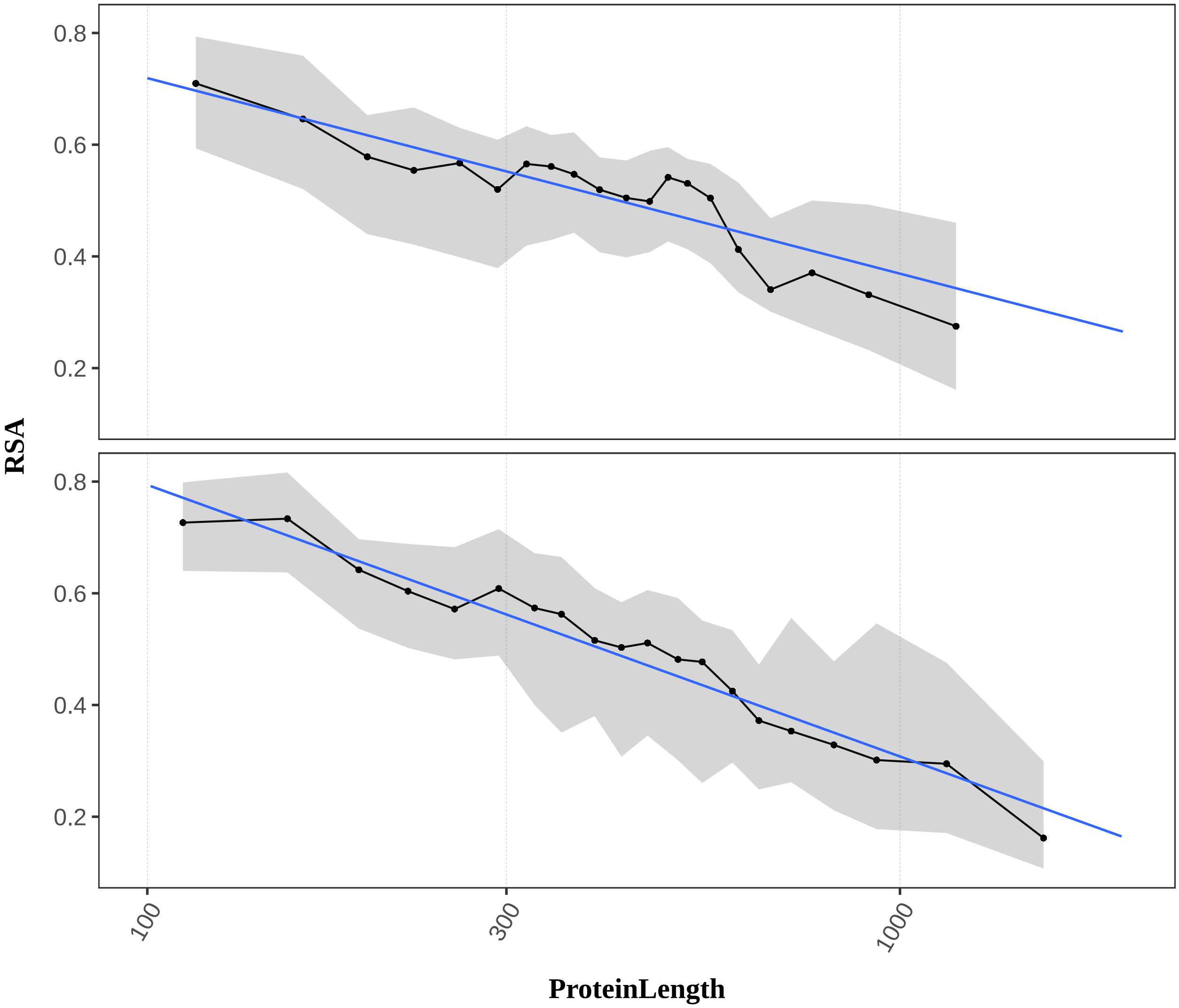


The next chunk presents the R script to estimate the statistics for every correlation.

```
# Protein Length/Gene Expression
tbls1 <- ddply(tbl.list[[1]], c("var","species"), function(x) {
  var <- as.numeric(factor(x$var1))
  variable.value <- as.numeric(factor(x$var2))
  corr = cor.test(var, variable.value, method = "kendall", exact = FALSE)
  Kendall.tau = corr$estimate
  p.value = corr$p.value
  dat = data.frame(Kendall.tau, p.value)
})
```

Table 1: Statistics for Protein Length/Gene Expression

| var | species | Kendall.tau | p.value |
|---|---|---|---|
| ProteinLength-GeneExpression | Arabidopsis | -0.0147082 | 0.0122551 |
| ProteinLength-GeneExpression | Drosophila | -0.0935642 | 0.0000000 |

Table 2: Statistics for Intrinsic Disorder/RSA

| species | kendall.tau | p.value |
|---|---|---|
| Arabidopsis | 0.0015173 | 0.5115756 |
| Drosophila | 0.0154635 | 0.0222795 |

```
kable(tbls1, caption = "Statistics for Protein Length/Gene Expression")
```

```
# Residue disorder/RSA --> need to do the statistics by sampling points several times
rep.dis.rsa <- list()
nboots <- 1000
for (j in 1:nboots) {
  rep.dis.rsa[[j]] <- ddply(tbl.list[[2]], "species", function(x) {
    x[sample.int(10000, replace = TRUE), ]
 })}

# then compute the statistics for each sampling point
tbls2 <- lapply(rep.dis.rsa, function(x) {
  ddply(x, c("species"), function(y) {
    var <- as.numeric(factor(y$var1))
    variable.value <- as.numeric(factor(y$var2))
    corr <- cor.test(var, variable.value, method = "kendall", exact = FALSE)
    Kendall.tau <- corr$estimate
    p.value <- corr$p.value
    data.frame(Kendall.tau, p.value)
  })
})
stat.dis.rsa <- rbindlist(tbls2)

# take the median values
stats.tbl2 <- ddply(stat.dis.rsa, "species", function(x) {
  kendall.tau <- median(x$Kendall.tau)
  p.value <- median(x$p.value)
  data.frame(kendall.tau, p.value)
})
kable(stats.tbl2, caption = "Statistics for Intrinsic Disorder/RSA")
```

```
# gene expression/RSA
tbls3 <- ddply(tbl.list[[3]], c("species", "var"), function(x) {
  var <- as.numeric(factor(x$var1))
  variable.value <- as.numeric(factor(x$var2))
  corr = cor.test(var, variable.value, method = "kendall", exact = FALSE)
  Kendall.tau = corr$estimate
  p.value = corr$p.value
  dat = data.frame(Kendall.tau, p.value)
})
```

Table 3: Statistics for Gene Expression/RSA

| species | var | Kendall.tau | p.value |
|---|---|---|---|
| Arabidopsis | GeneExpression-RSA | 0.0162997 | 0.1037291 |
| Drosophila | GeneExpression-RSA | 0.3274048 | 0.0000000 |

Table 4: Statistics for Protein Length/Solvent Exposure

| species | var | Kendall.tau | p.value |
|---|---|---|---|
| Arabidopsis | ProteinLength-RSA | -0.3102155 | 0 |
| Drosophila | ProteinLength-RSA | -0.4040298 | 0 |

```
kable(tbls3, caption = "Statistics for Gene Expression/RSA")
```

```
# protein length/proportion of exposed residues
tbls4 <- ddply(tbl.list[[4]], c("species", "var"), function(x) {
  var <- as.numeric(factor(x$var1))
  variable.value <- as.numeric(factor(x$var2))
  corr = cor.test(var, variable.value, method = "kendall", exact = FALSE)
  Kendall.tau = corr$estimate
  p.value = corr$p.value
  dat = data.frame(Kendall.tau, p.value)
})

kable(tbls4, caption = "Statistics for Protein Length/Solvent Exposure")
```

The next chunk describes the analyses between discrete and continuous variables, specifically: files.discrete[1] <- Disorder/Secondary structure files.discrete[2] <- RSA/Secondary structure

```
setwd("/Users/moutinho/Dropbox/Data/VariablesRelationship/Discrete/")

# calling all output tables
files.discrete <- list.files(".", ".csv")

# reading each table into a list
tbl.discrete <- lapply(files.discrete, fread, header = TRUE)
```

The next chunck describes the R scripts used to plot these relationships.

```
# plotting the relationship between secondary structure and relative solvent accessibility
tbl.discrete[[2]] <- unique(tbl.discrete[[2]])
tbl.discrete[[3]] <- unique(tbl.discrete[[3]])
plot.discrete2 <- lapply(tbl.discrete[2:3], function(x) {
  ggplot(x, aes(cat.var1, med.var2)) +
    geom_point(size=.6, col= "black") +
    geom_errorbar(aes(ymin=firstQ, ymax=thirdQ), width=.2) +
    facet_grid(rows = vars(species)) +
    xlab(as.character(unique(str_split_fixed(x$var, "\\-", 2)[,1]))) +
    ylab(as.character(unique(str_split_fixed(x$var, "\\-", 2)[,2]))) +
    theme_bw() +
    theme.plot()
})
```

```
plot.discrete2
```

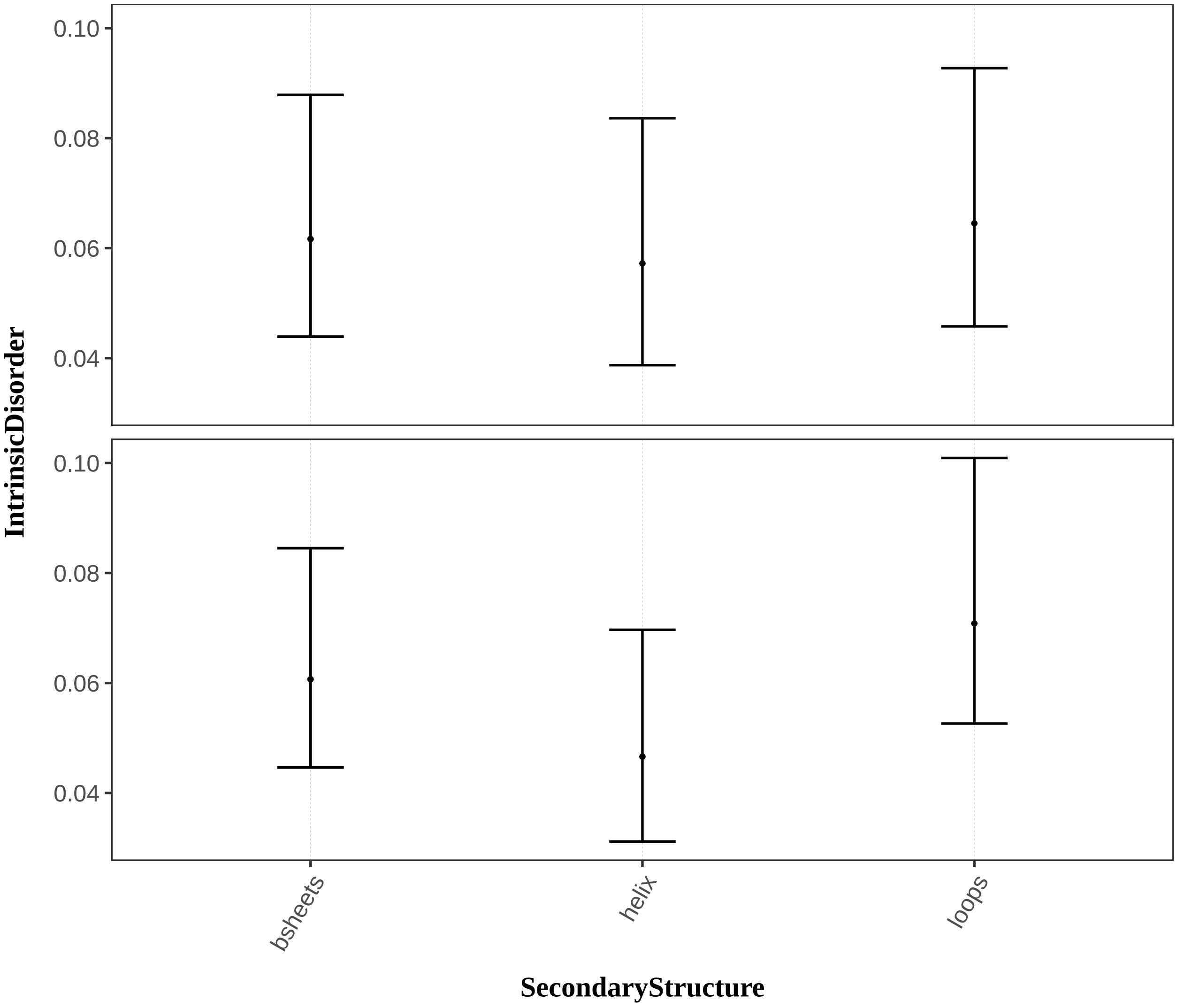

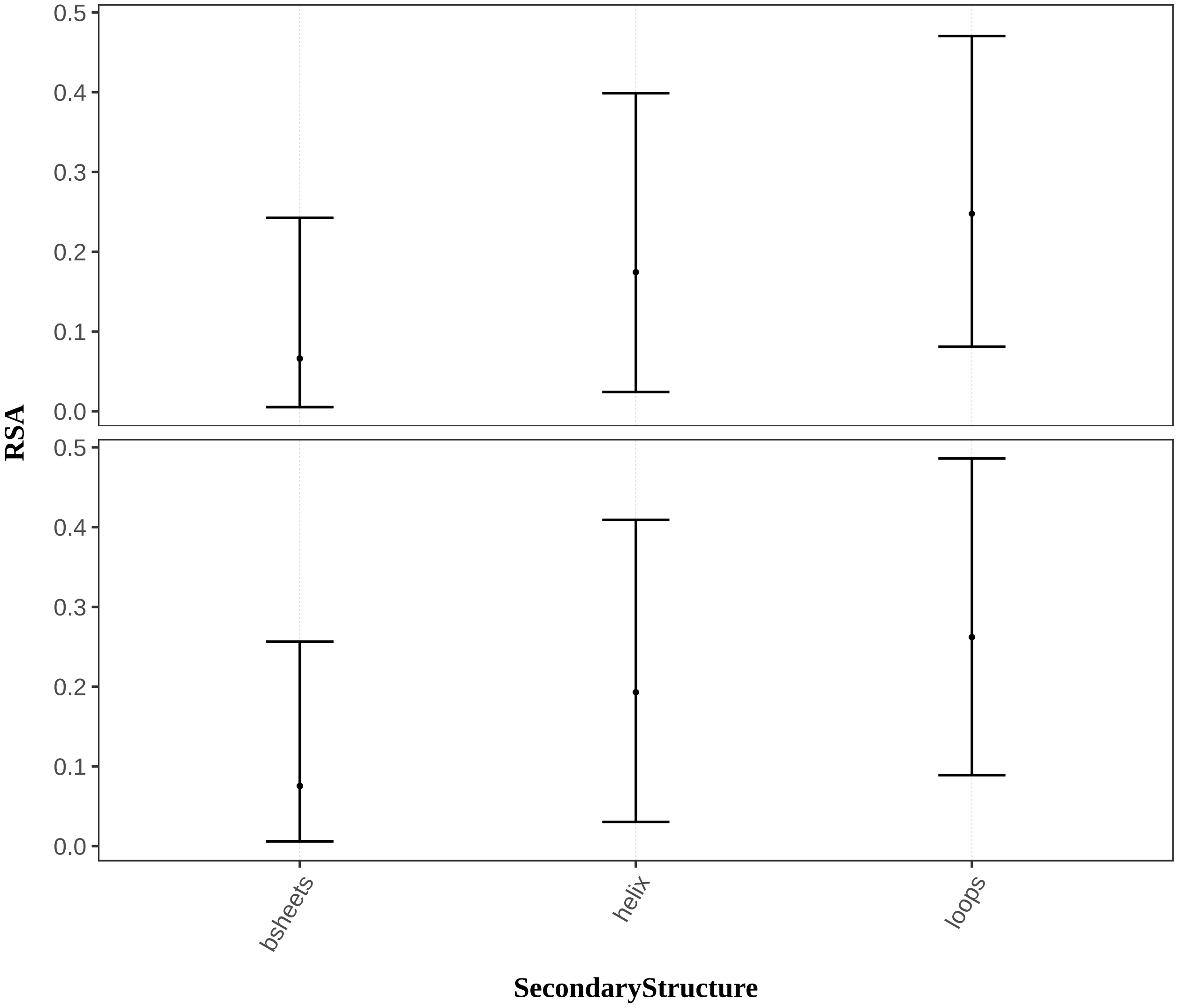


```r
# protein function/RSA
# Arabidopsis
tbl.arab <- subset(tbl.discrete[[1]], tbl.discrete[[1]]$species == "Arabidopsis")
plot.arab <- ggplot(tbl.arab, aes(x = cat.var1, y = med.var2)) +
  geom_point(size=0.5, col = "black") +
  geom_errorbar(aes(ymin=firstQ, ymax=thirdQ), width=.2) +
  ylab("Relative Solvent Accessibility") +
  xlab("Protein Functional Class") +
  scale_fill_grey() +
  scale_color_grey() +
  theme_bw() +
  theme.plot() +
  coord_flip()

plot.arab
```

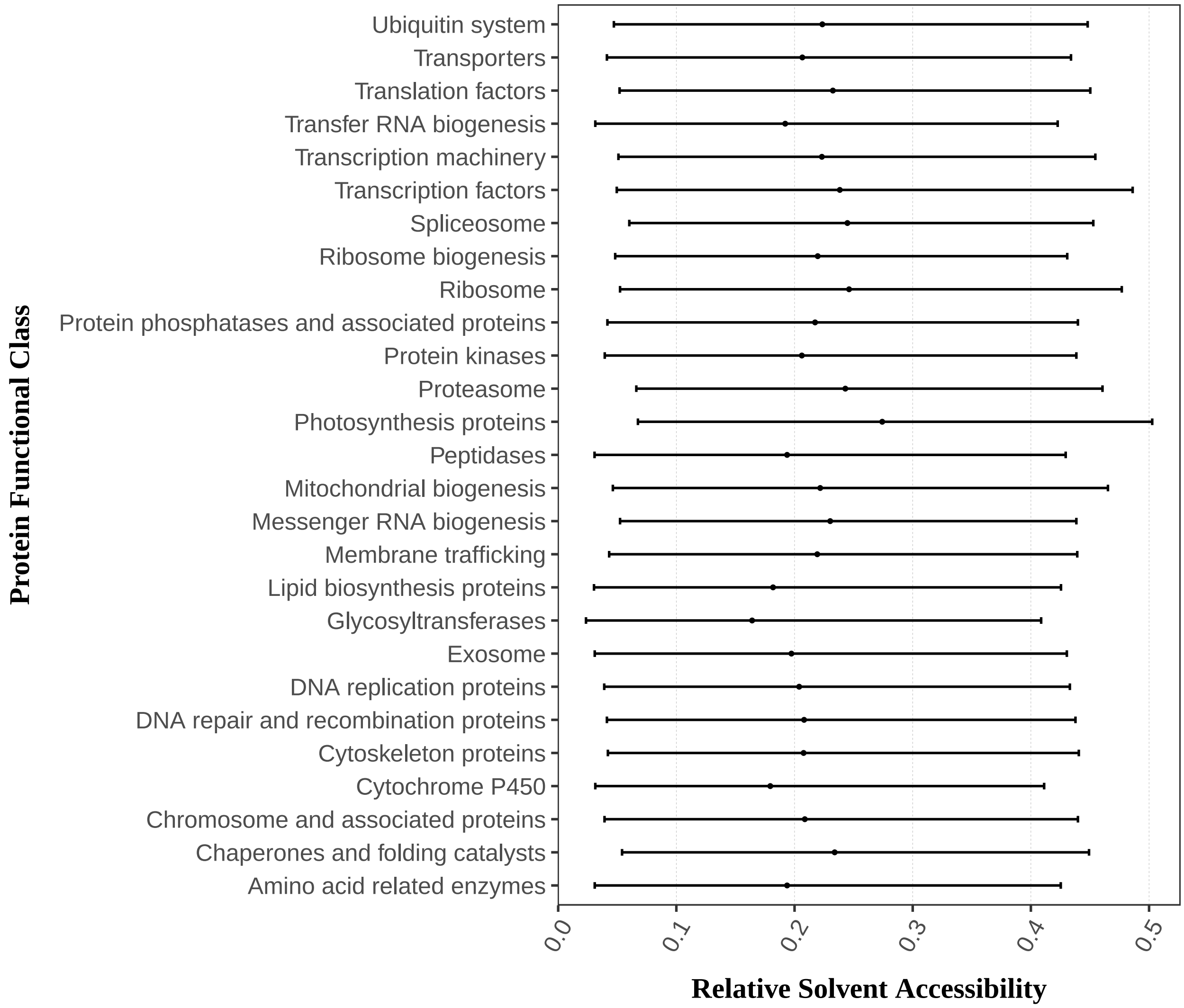


```r
# Drosophila
tbl.dmel <- subset(tbl.discrete[[1]], tbl.discrete[[1]]$species == "Drosophila")
plot.dmel <- ggplot(tbl.dmel, aes(x = cat.var1, y = med.var2)) +
  geom_point(size=0.5, col = "black") +
  geom_errorbar(aes(ymin=firstQ, ymax=thirdQ), width=.2) +
  ylab("Relative Solvent Accessibility") +
  xlab("Protein Functional Class") +
  scale_fill_grey() +
  scale_color_grey() +
  theme_bw() +
  theme.plot() +
  coord_flip()

plot.dmel
```

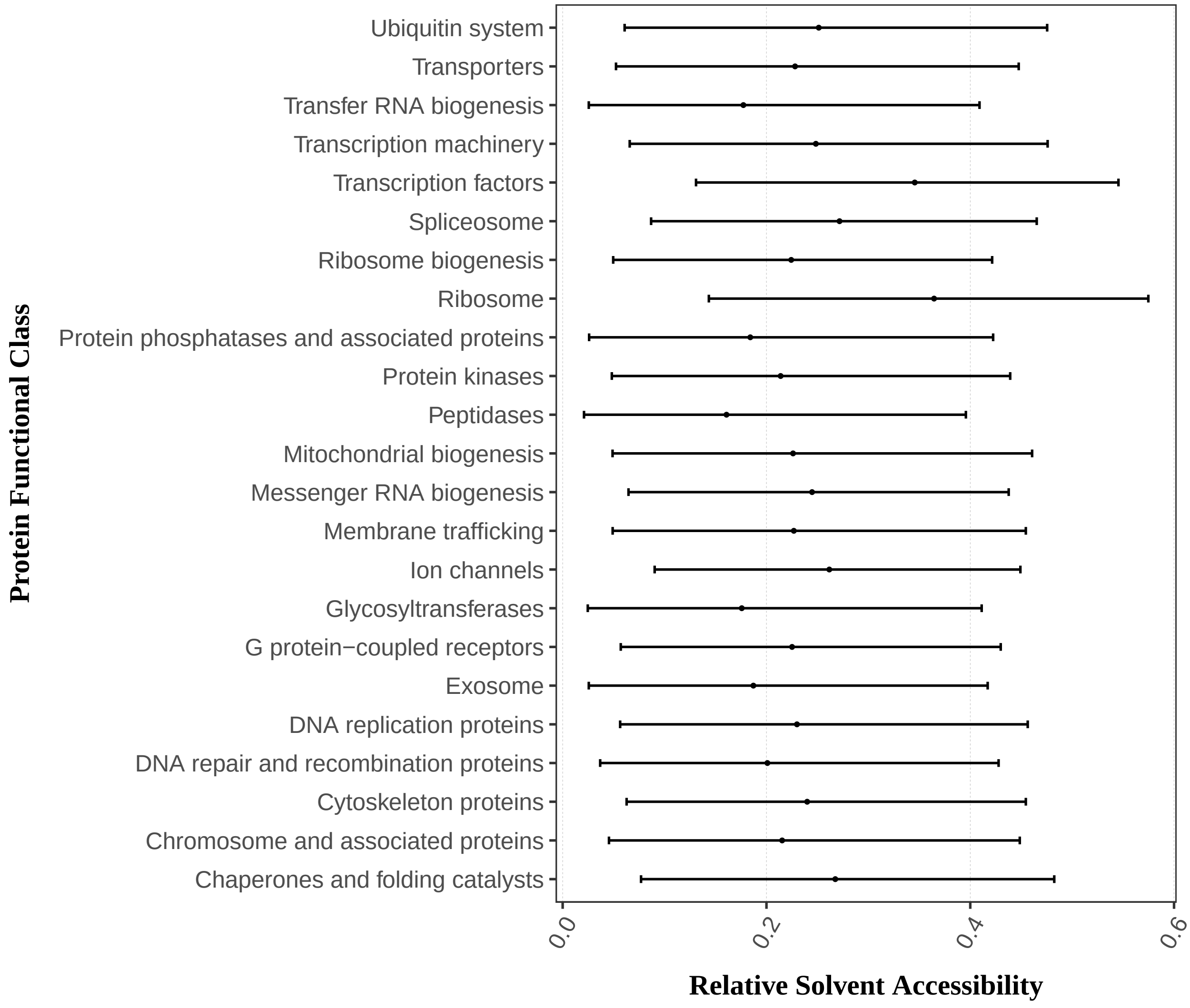
Ubiquitin system
Transporters
Transfer RNA biogenesis
Transcription machinery
Transcription factors
Spliceosome
Ribosome biogenesis
Ribosome
Protein phosphatases and associated proteins
Protein kinases
Peptidases
Mitochondrial biogenesis
Messenger RNA biogenesis
Membrane trafficking
Ion channels
Glycosyltransferases
G protein−coupled receptors
Exosome
DNA replication proteins
DNA repair and recombination proteins
Cytoskeleton proteins
Chromosome and associated proteins
Chaperones and folding catalysts
Protein Functional Class
0.0
0.2
0.4
0.6
Relative Solvent Accessibility
