## supplementary File S5 for "The impact of protein architecture on adaptive evolution"

### Correlation between variables

This is the script all plots and statistical analysis for the correlation between variables, refering to: Intrinsic Disorder/Secondary Structure, RSA/Active Site, RSA/Intrinsic Disorder, RSA/Protein Length and RSA/Secondary structure

For this purpose, two categories of RSA were considered: buried (RSA $<$ 0.05) and exposed (RSA $>=$ 0.05)

A function was created to call all files for each analysis performed. In this respect the order of the files is as follows:
files.correlations[1] <- Intrinsic Residue Disorder and Secondary Structure
files.correlations[2] <- RSA and Active Site
files.correlations[3] <- RSA and Residue Intrinsic Disorder
files.correlations[4] <- RSA and Protein Length
files.correlations[5] <- RSA and Secondary Structure

In these tables var 1 corresponds to the relative solvent accessibility and var 2 corresponds to the respective four different varaibles analysed (active site, disorder, protein length and secondary structure), with the exception of the correlation between intrinsic disorder and secondary structure, for which var1 corresponds to the structural motif and var2 the probability of residue intrinsic disorder.

The first part of the script removes bootstrap replicates for which the fitness effects parameters were not successfully fitted. For this purpose we discard 1% of the values above the maximum and below the minimum of each of the four parameters of fitness effects: Geman.neg, Gshape.neg, Gmean.neg and prop.pos.

```
setwd("/Users/moutinho/Dropbox/Data/Correlations/")

# Libraries
library(plyr)
library(dplyr)
library(data.table)
library(ggplot2)
library(reshape2)
library(doBy)
library(rlist)
library(stringr)
library(knitr)
library(kableExtra)
#

# calling all output tables
files.correlations <- list.files(".", ".csv")

# reading each table into a list
tbl.list <- lapply(files.correlations, read.table, header = TRUE)

## remove the outliers: 1% of the replicates below the min and
## 1% above the maximum
sub.tbl <- lapply(tbl.list, function(x) {
  ddply(x, c("species", "var1", "var2"), function(x) {
    sum.gmeanNeg <- summary(x$Gmean.neg)
    gmeanNeg.min1 <- as.numeric(sum.gmeanNeg[1]) + 0.01*as.numeric(sum.gmeanNeg[1])
    gmeanNeg.max1 <- as.numeric(sum.gmeanNeg[6]) - 0.01*as.numeric(sum.gmeanNeg[6])
    sum.gshapeNeg <- summary(x$Gshape.neg)
    gshapeNeg.min1 <- as.numeric(sum.gshapeNeg[1]) + 0.01*as.numeric(sum.gshapeNeg[1])
```

```
    gshapeNeg.max1 <- as.numeric(sum.gshapeNeg[6]) - 0.01*as.numeric(sum.gshapeNeg[6])
    sum.gmeanPos <- summary(x$Gmean.pos)
    gmeanPos.min1 <- as.numeric(sum.gmeanPos[1]) + 0.01*as.numeric(sum.gmeanPos[1])
    gmeanPos.max1 <- as.numeric(sum.gmeanPos[6]) - 0.01*as.numeric(sum.gmeanPos[6])
    sum.propPos <- summary(x$prop.pos)
    propPos.min1 <- as.numeric(sum.propPos[1]) + 0.01*as.numeric(sum.propPos[1])
    propPos.max1 <- as.numeric(sum.propPos[6]) - 0.01*as.numeric(sum.propPos[6])
    tbl <- x[!(x$Gmean.neg < gmeanNeg.min1 | x$Gmean.neg > gmeanNeg.max1 &
                 x$Gshape.neg < gshapeNeg.min1 | x$Gshape.neg > gshapeNeg.max1 &
                 x$Gmean.pos < gmeanPos.min1 | x$Gmean.pos > gmeanPos.max1 &
                 x$prop.pos < propPos.min1 | x$prop.pos > propPos.max1),]
  })
})
```

In the next chunk will take only the estimates concerning the rate of adaptive and non-adaptive substitutions, particularly: dnds, omegaNA and omegaA.

```
# In order to keep only the variables that we want to plot:
tbl.rates <- lapply(sub.tbl, function(x) {
  ddply(x, c("var", "species"), function(x) {
    melt(x, id.vars = c("var1", "var2"), measure.vars = c("dnds", "omegaNA", "omegaA"))
  })
})

# function to estimate the mean and satandard deviation
fun <- function(x){
  c(mean=mean(x), sd=sd(x))
}

# applying the above function to each output table for each value of each estimate
# (dnds, omegaA, omegaNA) for each value of the variable being analyzed for each species
tbl.stats <- lapply(tbl.rates, function(x) {
  summaryBy(value ~ variable + var1 + var2 + species + var, data=x, FUN = fun)
})

# to change the estimate name to the respective symbol
tbl.stats[[1]]$variable <- factor(tbl.stats[[1]]$variable,
                                  levels = c("dnds", "omegaNA", "omegaA"))
levels(tbl.stats[[1]]$variable) <- c(expression(omega), expression(omega[na]),
                                   expression(omega[a]))
tbl.stats[[2]]$variable <- factor(tbl.stats[[2]]$variable,
                                  levels = c("dnds", "omegaNA", "omegaA"))
levels(tbl.stats[[2]]$variable) <- c(expression(omega), expression(omega[na]),
                                   expression(omega[a]))
tbl.stats[[3]]$variable <- factor(tbl.stats[[3]]$variable,
                                  levels = c("dnds", "omegaNA", "omegaA"))
levels(tbl.stats[[3]]$variable) <- c(expression(omega), expression(omega[na]),
                                   expression(omega[a]))
tbl.stats[[4]]$variable <- factor(tbl.stats[[4]]$variable,
                                  levels = c("dnds", "omegaNA", "omegaA"))
levels(tbl.stats[[4]]$variable) <- c(expression(omega), expression(omega[na]),
                                   expression(omega[a]))
tbl.stats[[5]]$variable <- factor(tbl.stats[[5]]$variable,
                                  levels = c("dnds", "omegaNA", "omegaA"))
```

```
levels(tbl.stats[[5]]$variable) <- c(expression(omega), expression(omega[na]),
                                     expression(omega[a]))
```

The next chunk describes the script to plot the results of RSA/Active Site and RSA/Secondary Structure

```
# theme of the plot
theme.plot <- function(x) {
  theme(axis.title = element_text(face = "bold", color = "black", size=12,
                                  family = "Times"),
        text = element_text(size=12),
        axis.title.x = element_text(margin = margin(t = 9, r = 10, b = 0, l = 0)),
        axis.title.y = element_text(margin = margin(t = 9, r = 10, b = 0, l = 0)),
        panel.grid.minor=element_blank(),
        panel.grid.major = element_line(colour = "grey", linetype = "dashed", size = 0.1),
        panel.grid.major.y=element_blank(),
        strip.text.y = element_blank(),
        axis.text.x = element_text(angle = 60, hjust = 1),
        strip.background.x = element_rect(colour = "grey", fill = "gray92"),
        legend.text = element_text(size = 7),
        legend.title = element_text(size = 9, face = "bold"))
}

# plotting each of the output tables
plot.correlations.dis <- lapply(tbl.stats[c(2,5)], function(x) {
  ggplot(x, aes(x = var2, y = value.mean, fill = var1)) +
    geom_point(size=0.5, aes(fill = var1, col = var1)) +
    geom_errorbar(aes(ymin=value.mean + 1.96*value.sd,
                      ymax=value.mean - 1.96*value.sd, col = var1), width = .2) +
    facet_grid(species~variable, scales = "free_x", labeller = label_parsed) +
    ylab("") +
    xlab(as.character(x$var)) +
    scale_fill_grey() +
    scale_color_grey() +
    theme_bw() +
    theme.plot()
})

plot.correlations.dis
```

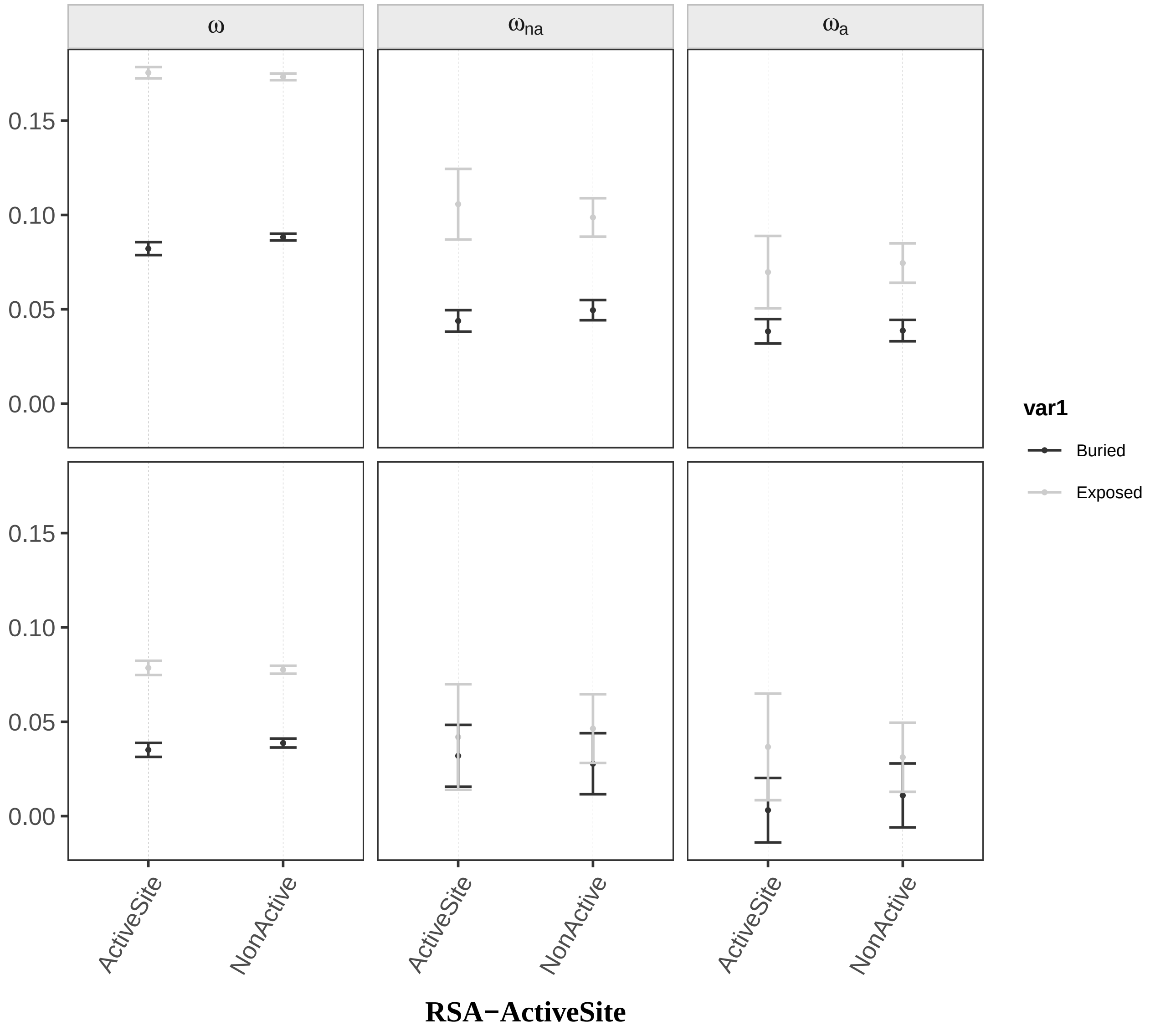
ω
$\omega_{na}$
$\omega_a$
0.15
0.10
0.05
0.00
0.15
0.10
0.05
0.00
ActiveSite
NonActive
ActiveSite
NonActive
ActiveSite
NonActive
RSA−ActiveSite
var1
Buried
Exposed

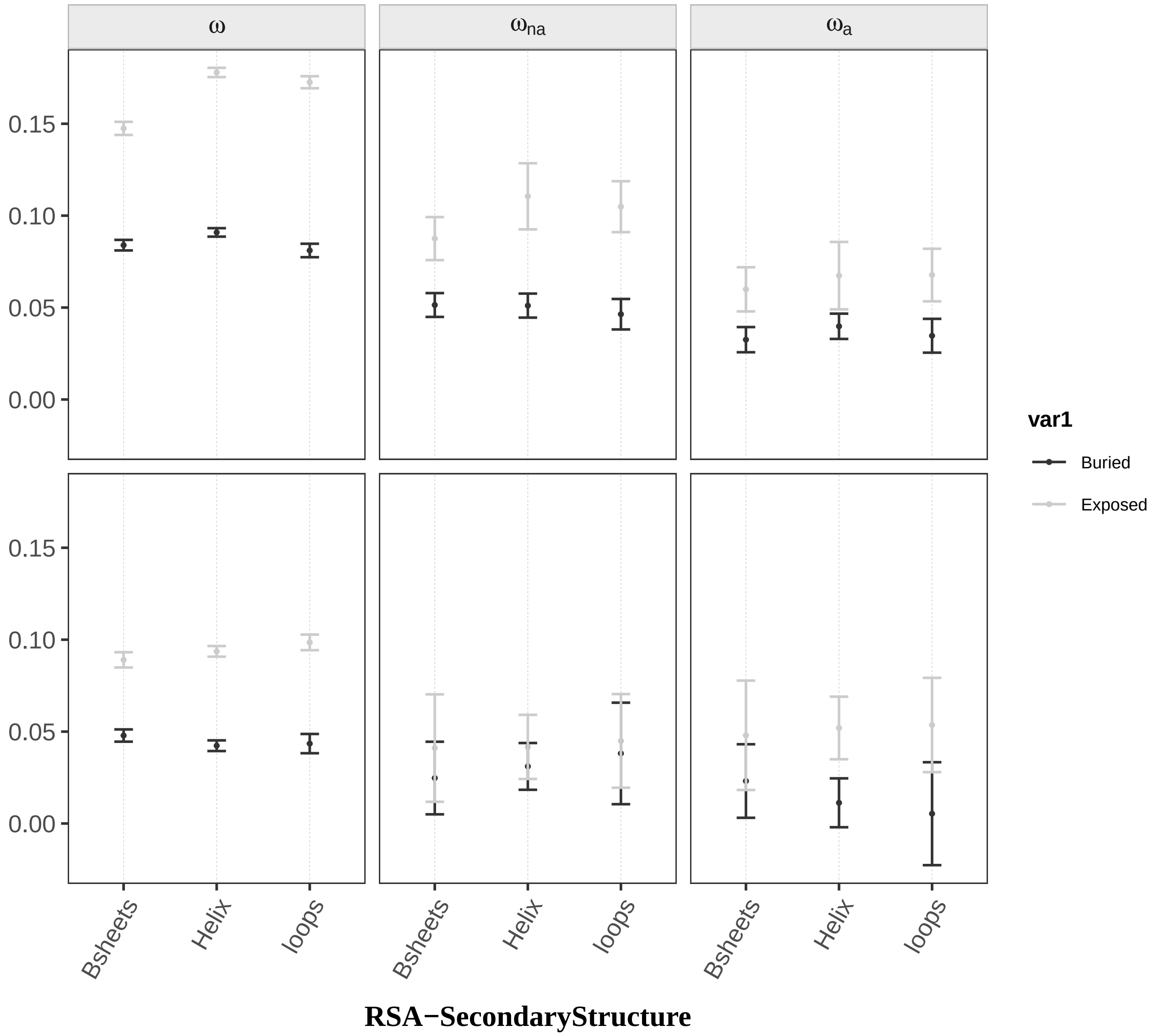


The next chunk describes the statistical analysis performed for the correlation of RSA/Active Site and RSA/Secondary Structure

```
####
#### RSA/Active Site
####

# checking the number of replicates per var.value, variable
# and species after removing 1% outliers
ps.nrep <- ddply(tbl.rates[[2]], c("species", "variable", "var1", "var2"),
                 function(x) {
    nrep <- nrow(x)
    data.frame(nrep)
})

# take the minimum number of replicates between categories
min.nrep <- ddply(ps.nrep, c("species", "var1", "var2"),
                  function(x) {
```

```r
  min.n <- min(x$nrep)
  data.frame(min.n)
})
min.nrep
```

| species | var1 | var2 | min.n |
|---|---|---|---|
| Arabidopsis | Buried | ActiveSite | 97 |
| Arabidopsis | Buried | NonActive | 97 |
| Arabidopsis | Exposed | ActiveSite | 96 |
| Arabidopsis | Exposed | NonActive | 98 |
| Drosophila | Buried | ActiveSite | 98 |
| Drosophila | Buried | NonActive | 97 |
| Drosophila | Exposed | ActiveSite | 97 |
| Drosophila | Exposed | NonActive | 98 |

```r
# 96 replicates Arabidopsis
nrep.arab <- subset(tbl.rates[[2]], tbl.rates[[2]]$species == "Arabidopsis")
tbl.arab <- ddply(nrep.arab, c("species", "var1", "var2", "variable"),
                  function(x) {
  x[sample(nrow(x), 96), ]
})

# 97 replicates for Drosophila
nrep.dmel <- subset(tbl.rates[[2]], tbl.rates[[2]]$species == "Drosophila")
tbl.dmel <- ddply(nrep.dmel, c("species", "var1", "var2", "variable"),
                  function(x) {
  x[sample(nrow(x), 97), ]
})

dat.ps <- rbind(tbl.arab, tbl.dmel)

# function to split the tables by the name of each variable
ps.split <- by(dat.ps, dat.ps[,c("var2")], function(y) y)

# to change the column name "value" to the respective id of the category
ps.value <- lapply(ps.split, function(x) {
  tbl <- data.frame(x)
  colnames(tbl)[6] <- as.character(unique(tbl$var2))
  tbl <- tbl[,-4]
  return(tbl)
})

# binding all values as columns
names.vars <- ps.value[[1]][,1:4]

tbl.estimate <- lapply(ps.value, function(x) {
  val <- x[,5]
})
tbl.ps <- list.cbind(tbl.estimate)
tbl.ps <- as.data.frame(tbl.ps)
tbl.ps <- cbind(names.vars, tbl.ps)
colnames(tbl.ps)[4] <- "estimate"

# now lets estimate the differences between the two variables
```

Table 1: Statistics for RSA/Active Site

| estimate | species | var1 | p.value |
|---|---|---|---|
| dnds | Arabidopsis | Buried | 0.0309278 |
| dnds | Arabidopsis | Exposed | 0.2164948 |
| omegaNA | Arabidopsis | Buried | 0.1134021 |
| omegaNA | Arabidopsis | Exposed | 0.5876289 |
| omegaA | Arabidopsis | Buried | 0.9175258 |
| omegaA | Arabidopsis | Exposed | 0.7319588 |

```
ps.dif <- ddply(tbl.ps, c("estimate", "species", "var1"), function(x){
  dif <- x$ActiveSite - x$NonActive
  tbl <- data.frame(dif)
})


## getting the p-value for each difference
## Arabidopsis
arab.dif <- subset(ps.dif, ps.dif$species == "Arabidopsis")
nboots <- 100
arab.p <- list()
for (i in 1:nboots) {
  arab.p[[i]] <- ddply(arab.dif, c("estimate", "species", "var1"),
                      function(x, N=96){
    c <- as.numeric(nrow(x[x$dif < 0,]))
    c2 <- as.numeric(nrow(x[x$dif > 0,]))
    m <- min(c, c2)
    p <- (2*m+1)/(N+1)
    tbl <- data.frame(m, p)
  })
}

arab.p.tbl <- rbindlist(arab.p)
arab.pvalue <- ddply(arab.p.tbl, c("estimate", "species", "var1"),
                    function(x) {
  p.value <- min(x$p)
  data.frame(p.value)
})

# showing the table
kable(arab.pvalue, caption = "Statistics for RSA/Active Site")
```

```
## Drosophila
dmel.dif <- subset(ps.dif, ps.dif$species == "Drosophila")
nboots <- 100
dmel.p <- list()
for (i in 1:nboots) {
  dmel.p[[i]] <- ddply(dmel.dif, c("estimate", "species", "var1"),
                      function(x, N=97){
    c <- as.numeric(nrow(x[x$dif < 0,]))
    c2 <- as.numeric(nrow(x[x$dif > 0,]))
    m <- min(c, c2)
```

Table 2: Statistics for RSA/Active Site

| estimate | species | var1 | p.value |
|---|---|---|---|
| dnds | Drosophila | Buried | 0.1122449 |
| dnds | Drosophila | Exposed | 0.6224490 |
| omegaNA | Drosophila | Buried | 0.6428571 |
| omegaNA | Drosophila | Exposed | 0.8673469 |
| omegaA | Drosophila | Buried | 0.4591837 |
| omegaA | Drosophila | Exposed | 0.8469388 |

```
    p <- (2*m+1)/(N+1)
    tbl <- data.frame(m, p)
  })
}
dmel.p.tbl <- rbindlist(dmel.p)
dmel.pvalue <- ddply(dmel.p.tbl, c("estimate", "species", "var1"),
                     function(x) {
  p.value <- min(x$p)
  data.frame(p.value)
})

# showing the table
kable(dmel.pvalue, caption = "Statistics for RSA/Active Site")
```

```
####
#### RSA/Secondary Structure
####

# checking the number of replicates per var.value, variable
# and species after removing 1% outliers
SS.nrep <- ddply(tbl.rates[[5]], c("species", "variable", "var1", "var2"),
                 function(x) {
    nrep <- nrow(x)
    data.frame(nrep)
})

# take the minimum number of replicates between categories
min.nrep <- ddply(SS.nrep, c("species", "var1", "var2"), function(x) {
  min.n <- min(x$nrep)
  data.frame(min.n)
})
min.nrep
```

| species | var1 | var2 | min.n |
|---|---|---|---|
| Arabidopsis | Buried | Bsheets | 98 |
| Arabidopsis | Buried | Helix | 95 |
| Arabidopsis | Buried | loops | 97 |
| Arabidopsis | Exposed | Bsheets | 97 |
| Arabidopsis | Exposed | Helix | 98 |
| Arabidopsis | Exposed | loops | 97 |
| Drosophila | Buried | Bsheets | 96 |
| Drosophila | Buried | Helix | 97 |
| Drosophila | Buried | loops | 97 |
| Drosophila | Exposed | Bsheets | 99 |
| Drosophila | Exposed | Helix | 98 |
| Drosophila | Exposed | loops | 99 |

```
# 95 replicates Arabidopsis
nrep.arab <- subset(tbl.rates[[5]], tbl.rates[[5]]$species == "Arabidopsis")
tbl.arab <- ddply(nrep.arab, c("species", "var1", "var2", "variable"),
                  function(x) {
  x[sample(nrow(x), 95), ]
})

# 96 replicates for Drosophila
nrep.dmel <- subset(tbl.rates[[5]], tbl.rates[[5]]$species == "Drosophila")
tbl.dmel <- ddply(nrep.dmel, c("species", "var1", "var2", "variable"),
                  function(x) {
  x[sample(nrow(x), 96), ]
})

dat.SS <- rbind(tbl.arab, tbl.dmel)

# function to split the tables by the name of each variable
SS.split <- by(dat.SS, dat.SS[,c("var2")], function(y) y)

# to change the column name "value" to the respective id of the category
SS.value <- lapply(SS.split, function(x) {
  tbl <- data.frame(x)
  colnames(tbl)[6] <- as.character(unique(tbl$var2))
  tbl <- tbl[,-4]
  return(tbl)
})

# binding all values as columns
names.vars <- SS.value[[1]][,1:4]

tbl.estimate <- lapply(SS.value, function(x) {
  val <- x[,5]
})
tbl.SS <- list.cbind(tbl.estimate)
tbl.SS <- as.data.frame(tbl.SS)
tbl.SS <- cbind(names.vars, tbl.SS)
colnames(tbl.SS)[4] <- "estimate"

# estimate the differences between columns
# to do so will duplicate the data.frame in order to subtract each column
```

```r
tbl2.SS <- tbl.SS[,5:7]

SS.dif <- cbind(tbl.SS[, c(1:4), drop=F],
        do.call(cbind, lapply(tbl2.SS[,1:2],
                              function(x) tbl.SS[,6:7]-x)))

# putting all variables in one column
SS.hist <- melt(SS.dif, id.vars = c("estimate", "species", "var1"),
                       measure.vars = c(names(SS.dif[5:6]),
                                        names(SS.dif[8])))

# puting NA values for when the difference is 0
# (i.e. when comparing the same variables)
split <- str_split_fixed(SS.hist$variable, "\\.", 2)
SS.hist$cat1 <- split[,2]
SS.hist$cat2 <- split[,1]

## getting the p-value for each difference
## N is not 100 because we took out the values that were outliers

## Arabidopsis
arab.hist <- subset(SS.hist, SS.hist$species == "Arabidopsis")
nboots <- 100
arab.p <- list()
for (i in 1:nboots) {
  arab.p[[i]] <- ddply(arab.hist,
                       c("species","estimate", "var1", "cat1", "cat2"),
                       function(x, N=95){
  c <- as.numeric(nrow(x[x$value < 0,]))
  c2 <- as.numeric(nrow(x[x$value > 0,]))
  m <- min(c, c2)
  p <- (2*m+1)/(N+1)
  tbl <- data.frame(c, c2, m, p)
  })
}

# correcting the p-value for multiple testing
arab.p.adj <- lapply(arab.p, function(x) {
  ddply(x,  c("species","estimate", "var1", "cat1", "cat2"),
        function(x) {
    p.value <- p.adjust(x$p)
    data.frame(p.value)
  })
})

# taking the minimum p-value
tbl.arab.p.adj <- rbindlist(arab.p.adj)
arab.pvalue <- ddply(tbl.arab.p.adj,
                     c("species","estimate", "var1", "cat1", "cat2"),
                     function(x) {
  p.value <- min(x$p.value)
  data.frame(p.value)
})
```

Table 3: Statistics for RSA/Secondary Structure

| species | estimate | var1 | cat1 | cat2 | p.value |
|---|---|---|---|---|---|
| Arabidopsis | dnds | Buried | Helix | Bsheets | 0.0104167 |
| Arabidopsis | dnds | Buried | loops | Bsheets | 0.3020833 |
| Arabidopsis | dnds | Buried | loops | Helix | 0.0104167 |
| Arabidopsis | dnds | Exposed | Helix | Bsheets | 0.0104167 |
| Arabidopsis | dnds | Exposed | loops | Bsheets | 0.0104167 |
| Arabidopsis | dnds | Exposed | loops | Helix | 0.0104167 |
| Arabidopsis | omegaNA | Buried | Helix | Bsheets | 0.9270833 |
| Arabidopsis | omegaNA | Buried | loops | Bsheets | 0.3229167 |
| Arabidopsis | omegaNA | Buried | loops | Helix | 0.3854167 |
| Arabidopsis | omegaNA | Exposed | Helix | Bsheets | 0.0312500 |
| Arabidopsis | omegaNA | Exposed | loops | Bsheets | 0.0520833 |
| Arabidopsis | omegaNA | Exposed | loops | Helix | 0.6979167 |
| Arabidopsis | omegaA | Buried | Helix | Bsheets | 0.1562500 |
| Arabidopsis | omegaA | Buried | loops | Bsheets | 0.7187500 |
| Arabidopsis | omegaA | Buried | loops | Helix | 0.4062500 |
| Arabidopsis | omegaA | Exposed | Helix | Bsheets | 0.6354167 |
| Arabidopsis | omegaA | Exposed | loops | Bsheets | 0.4479167 |
| Arabidopsis | omegaA | Exposed | loops | Helix | 0.8229167 |

```
# showing the table
kable(arab.pvalue, caption = "Statistics for RSA/Secondary Structure")
```

```
## Drosophila
dmel.hist <- subset(SS.hist, SS.hist$species == "Drosophila")
nboots <- 100
dmel.p <- list()
for (i in 1:nboots) {
  dmel.p[[i]] <- ddply(dmel.hist,
                       c("species","estimate", "var1", "cat1", "cat2"),
                       function(x, N=96){
  c <- as.numeric(nrow(x[x$value < 0,]))
  c2 <- as.numeric(nrow(x[x$value > 0,]))
  m <- min(c, c2)
  p <- (2*m+1)/(N+1)
  tbl <- data.frame(c, c2, m, p)
  })
}

# correcting the p-value for multiple testing
dmel.p.adj <- lapply(dmel.p, function(x) {
  ddply(x,  c("species","estimate", "var1", "cat1", "cat2"),
        function(x) {
    p.value <- p.adjust(x$p)
    data.frame(p.value)
  })
})

# taking the minimum p-value
```

Table 4: Statistics for RSA/Secondary Structure

| species | estimate | var1 | cat1 | cat2 | p.value |
|---|---|---|---|---|---|
| Drosophila | dnds | Buried | Helix | Bsheets | 0.0103093 |
| Drosophila | dnds | Buried | loops | Bsheets | 0.2577320 |
| Drosophila | dnds | Buried | loops | Helix | 0.6494845 |
| Drosophila | dnds | Exposed | Helix | Bsheets | 0.0927835 |
| Drosophila | dnds | Exposed | loops | Bsheets | 0.0103093 |
| Drosophila | dnds | Exposed | loops | Helix | 0.0515464 |
| Drosophila | omegaNA | Buried | Helix | Bsheets | 0.5463918 |
| Drosophila | omegaNA | Buried | loops | Bsheets | 0.5463918 |
| Drosophila | omegaNA | Buried | loops | Helix | 0.6907216 |
| Drosophila | omegaNA | Exposed | Helix | Bsheets | 0.9793814 |
| Drosophila | omegaNA | Exposed | loops | Bsheets | 0.7731959 |
| Drosophila | omegaNA | Exposed | loops | Helix | 0.8144330 |
| Drosophila | omegaA | Buried | Helix | Bsheets | 0.3195876 |
| Drosophila | omegaA | Buried | loops | Bsheets | 0.2371134 |
| Drosophila | omegaA | Buried | loops | Helix | 0.6494845 |
| Drosophila | omegaA | Exposed | Helix | Bsheets | 0.8762887 |
| Drosophila | omegaA | Exposed | loops | Bsheets | 0.8556701 |
| Drosophila | omegaA | Exposed | loops | Helix | 0.9175258 |

```r
tbl.dmel.p.adj <- rbindlist(dmel.p.adj)
dmel.pvalue <- ddply(tbl.dmel.p.adj,
                     c("species","estimate", "var1", "cat1", "cat2"),
                     function(x) {
  p.value <- min(x$p.value)
  data.frame(p.value)
})

# showing the table
kable(dmel.pvalue, caption = "Statistics for RSA/Secondary Structure")
```

The next chunk describes the script to plot the results of RSA/Disorder and RSA/Protein length.

```r
# theme of the plot
theme.plot <- function(x) {
  theme(axis.title = element_text(face = "bold", color = "black", size=12),
        text = element_text(size=12),
        axis.title.x = element_text(margin = margin(t = 18, r = 10, b = 0, l = 0)),
        axis.title.y = element_text(margin = margin(t = 18, r = 10, b = 0, l = 0)),
        panel.grid.minor=element_blank(),
        panel.grid.major = element_line(colour = "grey", linetype = "dashed", size = 0.2),
        panel.grid.major.y=element_blank(),
        axis.text.x = element_text(angle = 60, hjust = 1),
        legend.text = element_text(size = 7),
        legend.title = element_text(size = 10, face = "bold"))
}

# plotting the continuous variables
plot.correlations.conti <- lapply(tbl.stats[c(1,3:4)], function(x) {
  ggplot(x, aes(x = var2, y = value.mean, fill = var1)) +
```

```
  geom_line(size = 0.3, col = "black")+
  geom_ribbon(aes(ymin=value.mean + 1.96*value.sd,
                  ymax=value.mean - 1.96*value.sd, fill = var1), alpha = 0.8) +
  facet_grid(species~variable, scales = "free_x", labeller = label_parsed) +
  ylab("") +
  xlab(as.character(x$var)) +
  scale_fill_grey() +
  scale_color_grey() +
  theme_bw() +
  theme.plot()
})

plot.correlations.conti
```

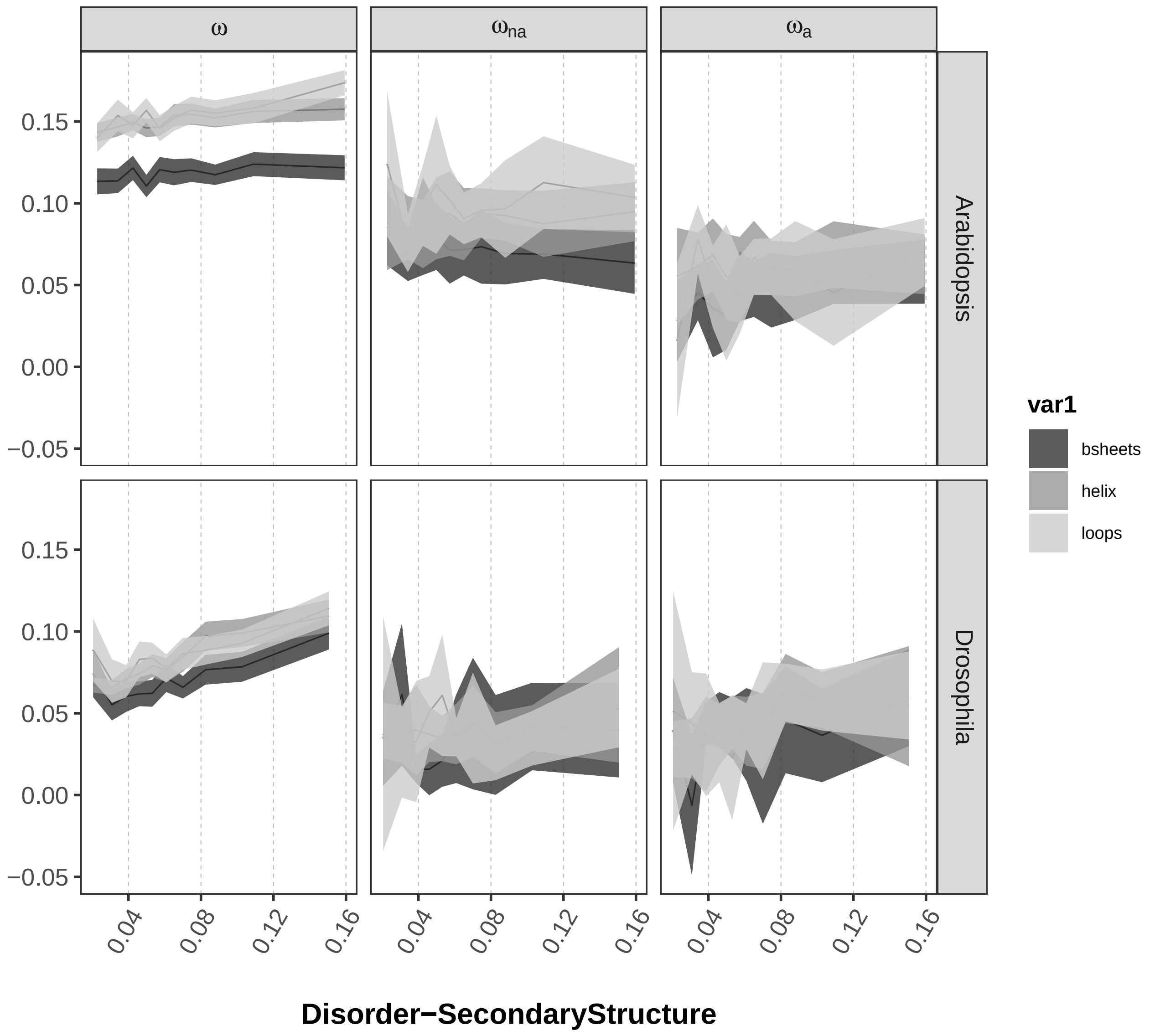

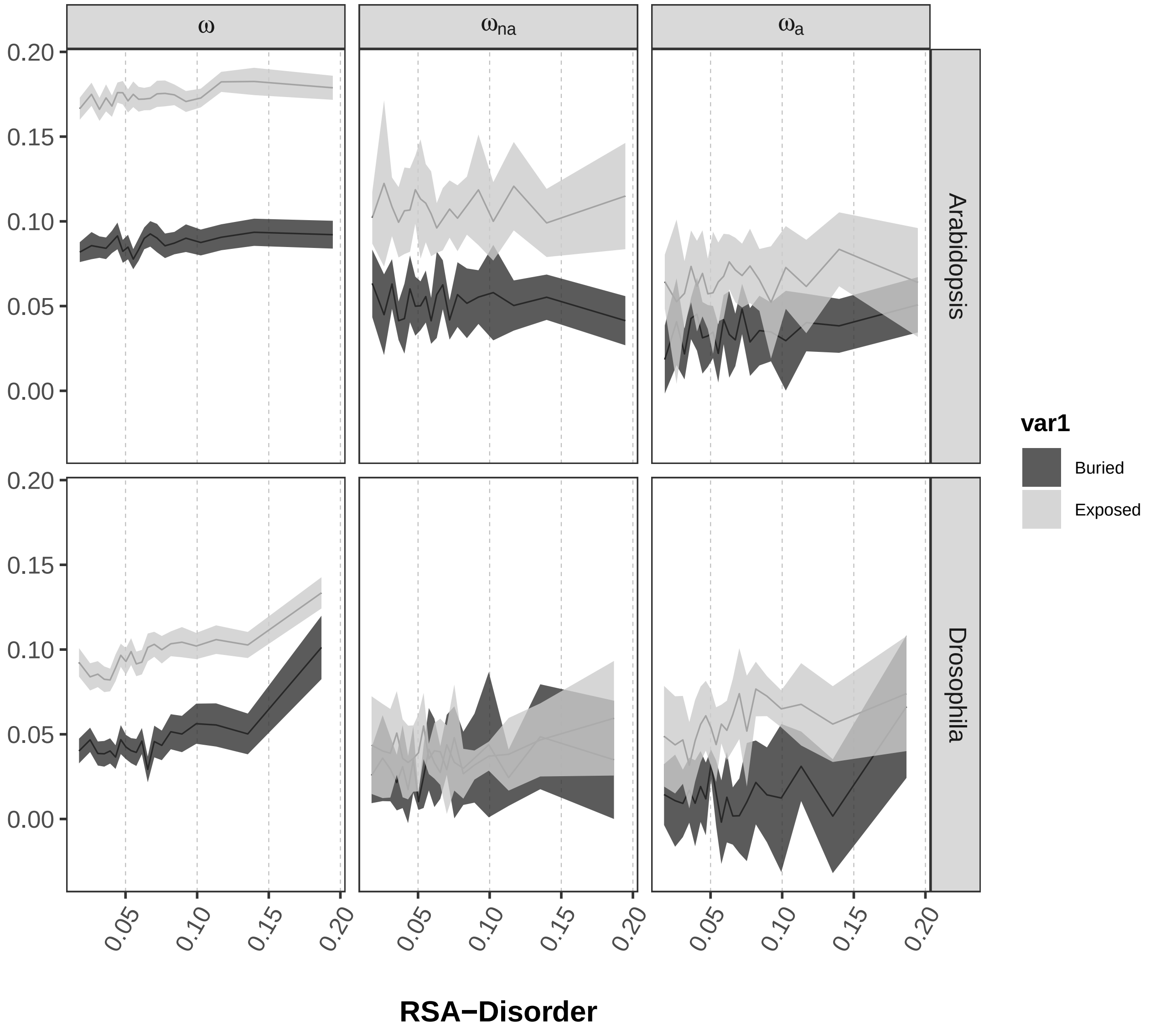

ω
ωna
ωa
0.20
0.15
0.10
0.05
0.00
Arabidopsis
Drosophila
0.05
0.10
0.15
0.20
RSA−Disorder
var1
Buried
Exposed

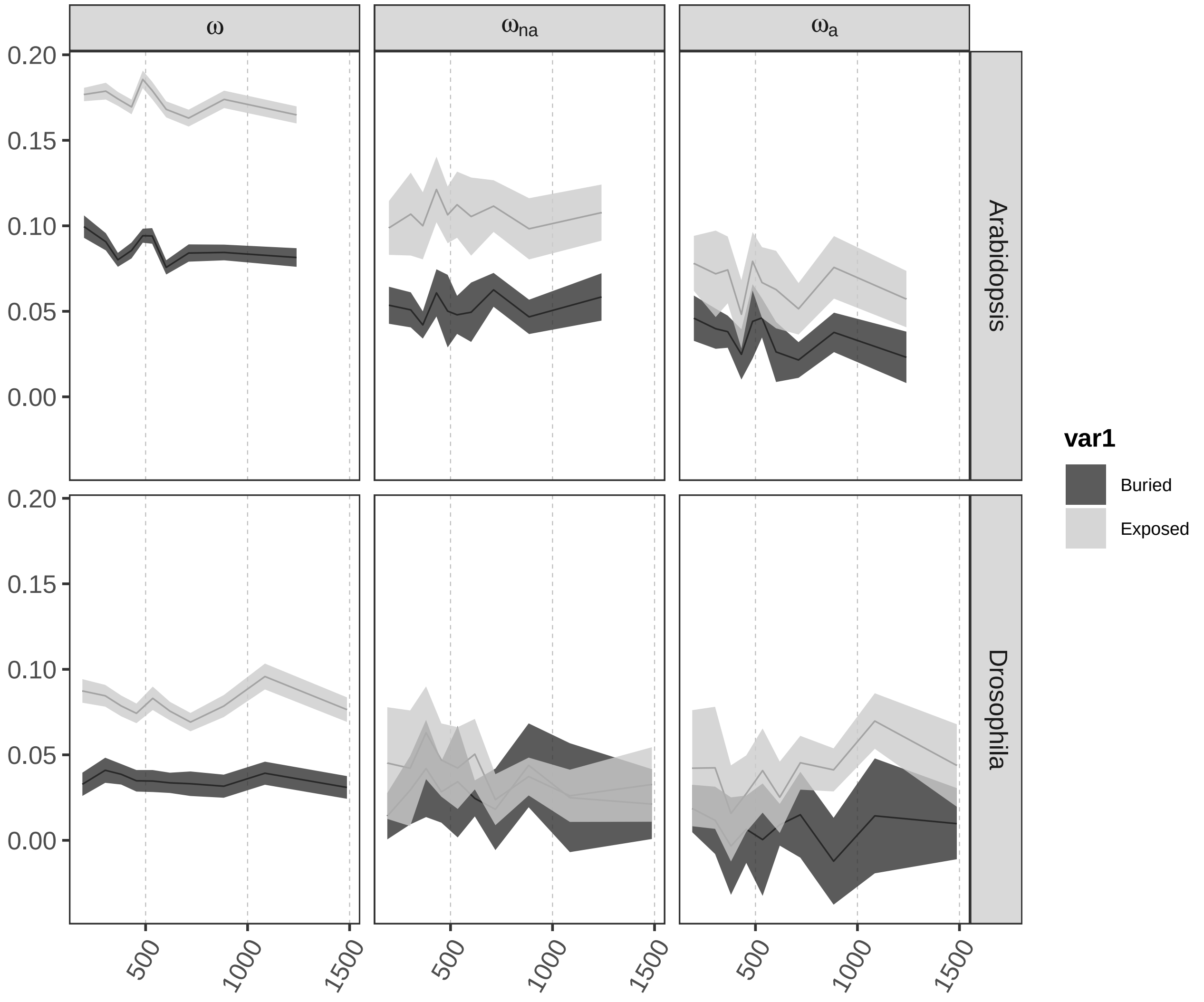


**RSA−ProteinLength**

The last section shows how the statistical analyses for RSA/Disorder and RSA/Protein length were performed.

```
stat <- lapply(tbl.stats[c(1,3:4)], function(x) {
  ddply(x, c("var", "species", "variable", "var1"), function(x) {
    var <- as.numeric(factor(x$var2))
    variable.value <- as.numeric(factor(x$value.mean))
    corr = cor.test(var, variable.value, method = "kendall", exact = FALSE)
    Kendall.tau = corr$estimate
    p.value = corr$p.value
    dat = data.frame(Kendall.tau, p.value)
  })
})

# showing the tables
for(i in stat) {
  print(kable(x = i, caption = paste0("Statistics for ", unique(i$var))))
}
```

Table 5: Statistics for Disorder-SecondaryStructure

| var | species | variable | var1 | Kendall.tau | p.value |
|---|---|---|---|---|---|
| Disorder-SecondaryStructure | Arabidopsis | omega | bsheets | 0.4222222 | 0.0892416 |
| Disorder-SecondaryStructure | Arabidopsis | omega | helix | 0.7333333 | 0.0031612 |
| Disorder-SecondaryStructure | Arabidopsis | omega | loops | 0.6444444 | 0.0094911 |
| Disorder-SecondaryStructure | Arabidopsis | omega[na] | bsheets | -0.5111111 | 0.0396687 |
| Disorder-SecondaryStructure | Arabidopsis | omega[na] | helix | 0.4222222 | 0.0892416 |
| Disorder-SecondaryStructure | Arabidopsis | omega[na] | loops | 0.0666667 | 0.7884467 |
| Disorder-SecondaryStructure | Arabidopsis | omega[a] | bsheets | 0.6888889 | 0.0055589 |
| Disorder-SecondaryStructure | Arabidopsis | omega[a] | helix | 0.2000000 | 0.4208286 |
| Disorder-SecondaryStructure | Arabidopsis | omega[a] | loops | 0.2000000 | 0.4208286 |
| Disorder-SecondaryStructure | Drosophila | omega | bsheets | 0.6888889 | 0.0055589 |
| Disorder-SecondaryStructure | Drosophila | omega | helix | 0.9111111 | 0.0002453 |
| Disorder-SecondaryStructure | Drosophila | omega | loops | 0.5555556 | 0.0253473 |
| Disorder-SecondaryStructure | Drosophila | omega[na] | bsheets | 0.1555556 | 0.5312500 |
| Disorder-SecondaryStructure | Drosophila | omega[na] | helix | 0.2444444 | 0.3251795 |
| Disorder-SecondaryStructure | Drosophila | omega[na] | loops | 0.1111111 | 0.6547208 |
| Disorder-SecondaryStructure | Drosophila | omega[a] | bsheets | 0.1555556 | 0.5312500 |
| Disorder-SecondaryStructure | Drosophila | omega[a] | helix | 0.7777778 | 0.0017451 |
| Disorder-SecondaryStructure | Drosophila | omega[a] | loops | 0.3333333 | 0.1797125 |

Table 6: Statistics for RSA-Disorder

| var | species | variable | var1 | Kendall.tau | p.value |
|---|---|---|---|---|---|
| RSA-Disorder | Arabidopsis | omega | Buried | 0.4526316 | 0.0052674 |
| RSA-Disorder | Arabidopsis | omega | Exposed | 0.3684211 | 0.0231409 |
| RSA-Disorder | Arabidopsis | omega[na] | Buried | -0.0631579 | 0.6970310 |
| RSA-Disorder | Arabidopsis | omega[na] | Exposed | -0.0210526 | 0.8967428 |
| RSA-Disorder | Arabidopsis | omega[a] | Buried | 0.2105263 | 0.1943659 |
| RSA-Disorder | Arabidopsis | omega[a] | Exposed | 0.2105263 | 0.1943659 |
| RSA-Disorder | Drosophila | omega | Buried | 0.5052632 | 0.0018416 |
| RSA-Disorder | Drosophila | omega | Exposed | 0.7157895 | 0.0000102 |
| RSA-Disorder | Drosophila | omega[na] | Buried | 0.2947368 | 0.0692355 |
| RSA-Disorder | Drosophila | omega[na] | Exposed | -0.0315789 | 0.8456547 |
| RSA-Disorder | Drosophila | omega[a] | Buried | 0.0842105 | 0.6036850 |
| RSA-Disorder | Drosophila | omega[a] | Exposed | 0.5368421 | 0.0009352 |

Table 7: Statistics for RSA-ProteinLength

| var | species | variable | var1 | Kendall.tau | p.value |
|---|---|---|---|---|---|
| RSA-ProteinLength | Arabidopsis | omega | Buried | -0.3777778 | 0.1283788 |
| RSA-ProteinLength | Arabidopsis | omega | Exposed | -0.4222222 | 0.0892416 |
| RSA-ProteinLength | Arabidopsis | omega[na] | Buried | -0.0222222 | 0.9287301 |
| RSA-ProteinLength | Arabidopsis | omega[na] | Exposed | 0.0666667 | 0.7884467 |
| RSA-ProteinLength | Arabidopsis | omega[a] | Buried | -0.4222222 | 0.0892416 |
| RSA-ProteinLength | Arabidopsis | omega[a] | Exposed | -0.2888889 | 0.2449288 |
| RSA-ProteinLength | Drosophila | omega | Buried | -0.4222222 | 0.0892416 |
| RSA-ProteinLength | Drosophila | omega | Exposed | -0.2444444 | 0.3251795 |
| RSA-ProteinLength | Drosophila | omega[na] | Buried | -0.0666667 | 0.7884467 |
| RSA-ProteinLength | Drosophila | omega[na] | Exposed | -0.4222222 | 0.0892416 |
| RSA-ProteinLength | Drosophila | omega[a] | Buried | -0.0666667 | 0.7884467 |
| RSA-ProteinLength | Drosophila | omega[a] | Exposed | 0.3333333 | 0.1797125 |
