## supplementary File S3 for "The impact of protein architecture on adaptive evolution"

### Discrete variables

This is the script describing all plots and statistical analysis regarding the analysis of discrete variables, refering to: secondary structure motif, binding affinity to dnak and protein location. A function was created to call all files for each analysis performed. In this respect the order of the files is as follows:
files.discrete[1] <- Molecular Chaperone dnaK output table
files.discrete[2] <- Protein Location output table
files.discrete[3] <- Secondary Structure motif output table

The first part of the script removes bootstrap replicates for which the fitness effects parameters were not successfully fitted. For this purpose we discard 1% of the values above the maximum and below the minimum of each of the four parameters of fitness effects: Geman.neg, Gshape.neg, Gmean.neg and prop.pos.

```
setwd("/Users/moutinho/Dropbox/Data/Discrete/")

# Libraries
library(plyr)
library(dplyr)
library(data.table)
library(ggplot2)
library(reshape2)
library(doBy)
library(knitr)
library(kableExtra)
#

# calling all output tables
files.discrete <- list.files(".", ".csv")

# reading each table into a list
tbl.list <- lapply(files.discrete, read.table, header = TRUE)

## remove the outliers: 1% of the replicates below the min and
## 1% above the maximum
sub.tbl <- lapply(tbl.list, function(x) {
  ddply(x, c("species", "var.value"), function(x) {
    sum.gmeanNeg <- summary(x$Gmean.neg)
    gmeanNeg.min1 <- as.numeric(sum.gmeanNeg[1]) + 0.01*as.numeric(sum.gmeanNeg[1])
    gmeanNeg.max1 <- as.numeric(sum.gmeanNeg[6]) - 0.01*as.numeric(sum.gmeanNeg[6])
    sum.gshapeNeg <- summary(x$Gshape.neg)
    gshapeNeg.min1 <- as.numeric(sum.gshapeNeg[1]) + 0.01*as.numeric(sum.gshapeNeg[1])
    gshapeNeg.max1 <- as.numeric(sum.gshapeNeg[6]) - 0.01*as.numeric(sum.gshapeNeg[6])
    sum.gmeanPos <- summary(x$Gmean.pos)
    gmeanPos.min1 <- as.numeric(sum.gmeanPos[1]) + 0.01*as.numeric(sum.gmeanPos[1])
    gmeanPos.max1 <- as.numeric(sum.gmeanPos[6]) - 0.01*as.numeric(sum.gmeanPos[6])
    sum.propPos <- summary(x$prop.pos)
    propPos.min1 <- as.numeric(sum.propPos[1]) + 0.01*as.numeric(sum.propPos[1])
    propPos.max1 <- as.numeric(sum.propPos[6]) - 0.01*as.numeric(sum.propPos[6])
    tbl <- x[!(x$Gmean.neg < gmeanNeg.min1 | x$Gmean.neg > gmeanNeg.max1 &
                 x$Gshape.neg < gshapeNeg.min1 | x$Gshape.neg > gshapeNeg.max1 &
                 x$Gmean.pos < gmeanPos.min1 | x$Gmean.pos > gmeanPos.max1 &
                 x$prop.pos < propPos.min1 | x$prop.pos > propPos.max1),]
  })
```

```
})
```

In the next chunk will take only the estimates concerning the rate of adaptive and non-adaptive substitutions, particularly: dnds, omegaNA and omegaA.

```
# In order to keep only the variables that we want to plot:
tbl.rates <- lapply(sub.tbl, function(x) {
  ddply(x, c("var", "species"), function(x) {
    melt(x, id.vars = c("var.value"), measure.vars = c("dnds", "omegaNA", "omegaA"))
  })
})

# function to estimate the mean and satandard deviation to plot the results with the
# mean of the bootstrap replicates and the 95% confidence interval

fun <- function(x){
  c(mean=mean(x), sd=sd(x))
}

# applying the above function to each output table for each value of each estimate
# (dnds, omegaA, omegaNA) for each value of the variable being analyzed for each species

tbl.stats <- lapply(tbl.rates, function(x) {
  summaryBy(value ~ variable + var.value + species + var, data=x, FUN = fun)
})

# to change the estimate name to the respective symbol
tbl.stats[[1]]$variable <- factor(tbl.stats[[1]]$variable,
                                  levels = c("dnds", "omegaNA", "omegaA"))
levels(tbl.stats[[1]]$variable) <- c(expression(omega), expression(omega[na]),
                                   expression(omega[a]))
tbl.stats[[2]]$variable <- factor(tbl.stats[[2]]$variable,
                                  levels = c("dnds", "omegaNA", "omegaA"))
levels(tbl.stats[[2]]$variable) <- c(expression(omega), expression(omega[na]),
                                   expression(omega[a]))
tbl.stats[[3]]$variable <- factor(tbl.stats[[3]]$variable,
                                  levels = c("dnds", "omegaNA", "omegaA"))
levels(tbl.stats[[3]]$variable) <- c(expression(omega), expression(omega[na]),
                                   expression(omega[a]))
```

The next chunck of the script shows the code used for plotting the results.

```
# theme of the plot
theme.plot <- function(x) {
  theme(axis.title = element_text(face = "bold", color = "black", size=14),
        text = element_text(size=14),
        axis.title.x = element_text(margin = margin(t = 18, r = 10, b = 0, l = 0)),
        axis.title.y = element_text(margin = margin(t = 18, r = 10, b = 0, l = 0)),
        panel.grid.minor=element_blank(),
        panel.grid.major = element_line(colour = "grey", linetype = "dashed", size = 0.2),
        panel.grid.major.y=element_blank(),
        #strip.text.y = element_blank(),
        axis.text.x = element_text(angle = 60, hjust = 1))
}
```

```r
# plotting each of the output tables
plot.discrete <- lapply(tbl.stats, function(x) {
  ggplot(x, aes(x = var.value, y = value.mean)) +
  geom_point(size=0.5, col = "black") +
  geom_errorbar(aes(ymin=value.mean + 1.96*value.sd,
                    ymax=value.mean - 1.96*value.sd), width = .2) +
  facet_grid(species~variable, scales = "free_x", labeller = label_parsed) +
  ylab("") +
  xlab(as.character(x$var)) +
  theme_bw() +
  theme.plot()
})

plot.discrete
```

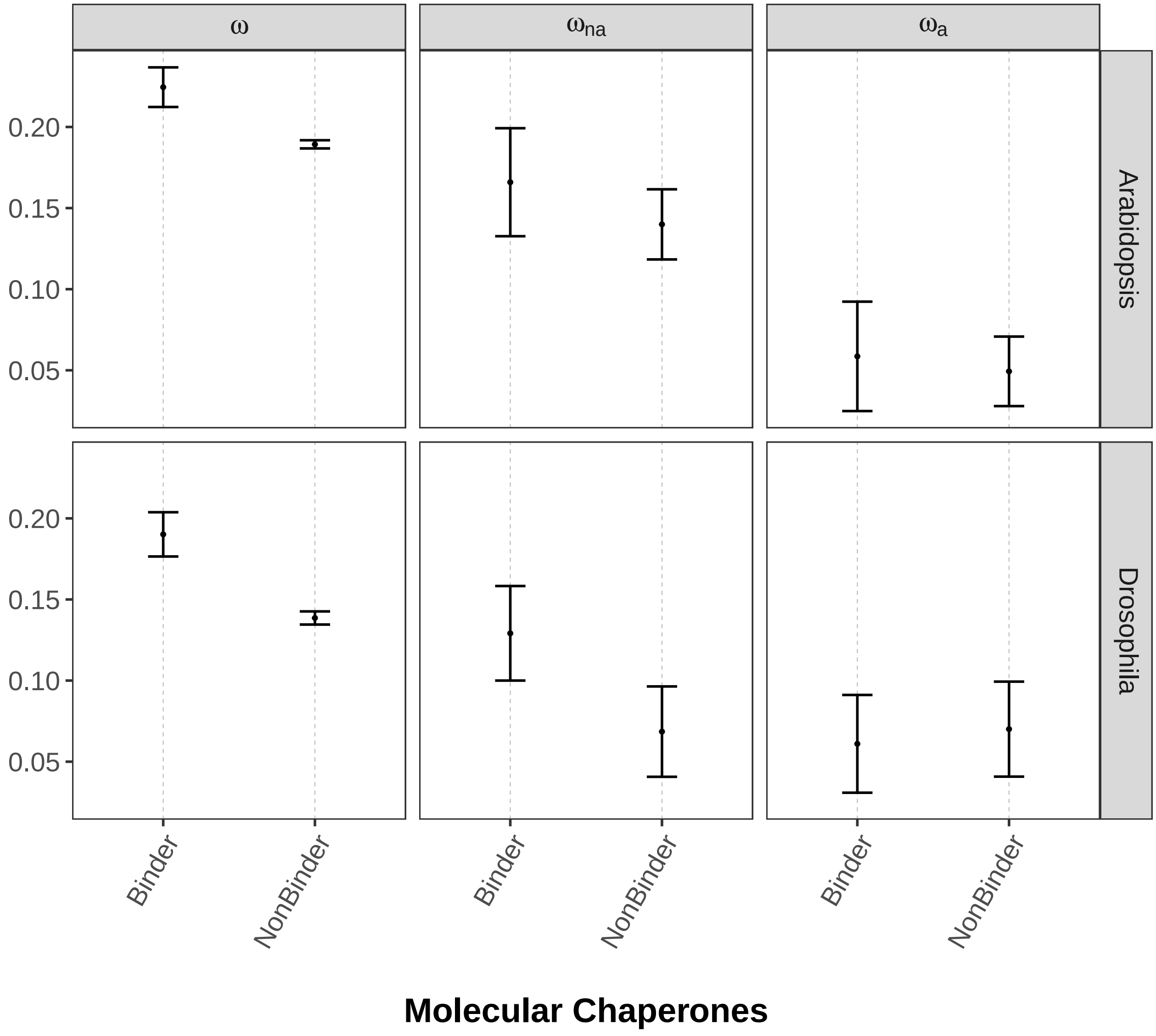

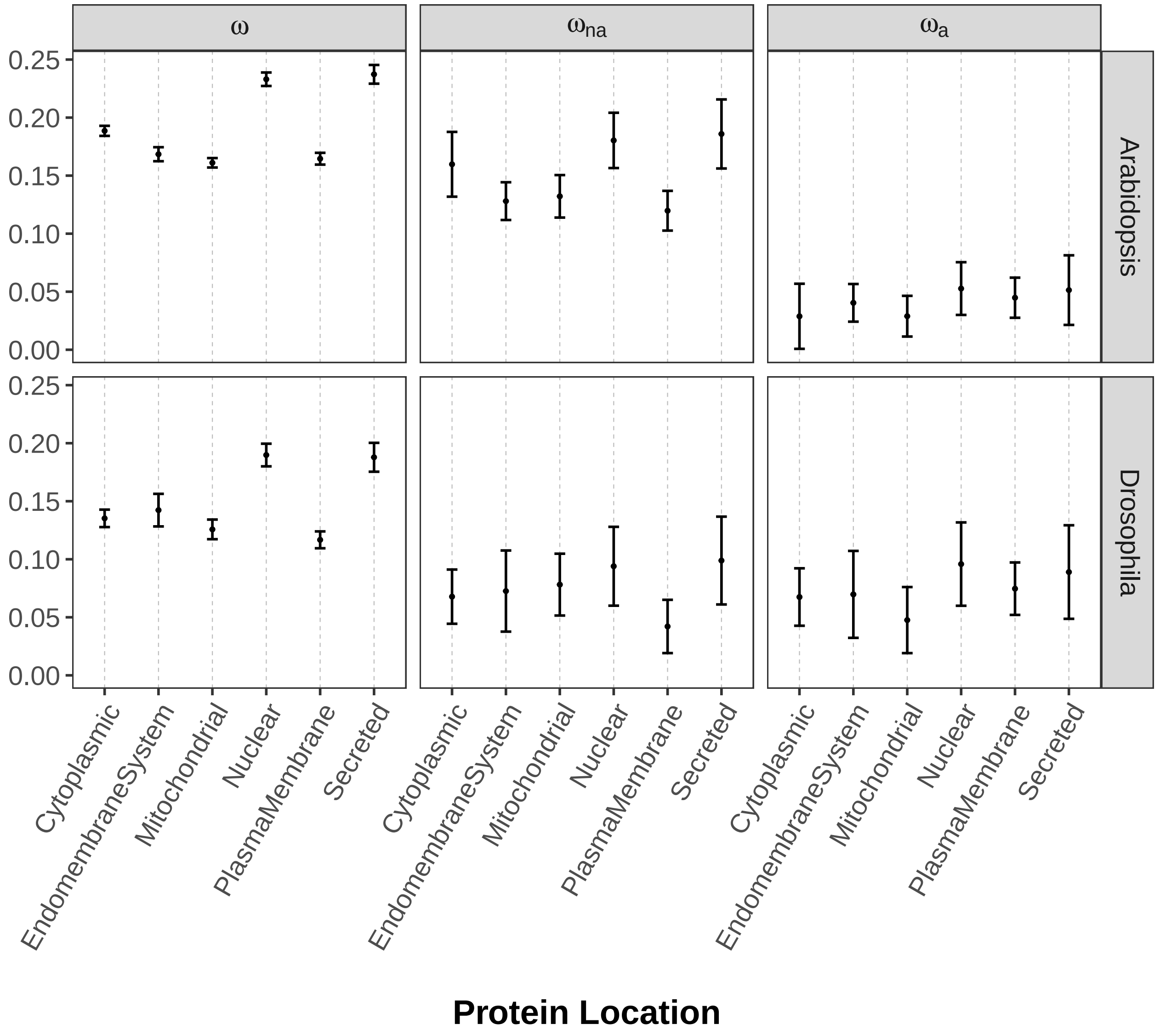

ω
$\omega_{na}$
$\omega_a$
Arabidopsis
Drosophila
0.25
0.20
0.15
0.10
0.05
0.00
Cytoplasmic
EndomembraneSystem
Mitochondrial
Nuclear
PlasmaMembrane
Secreted
Protein Location

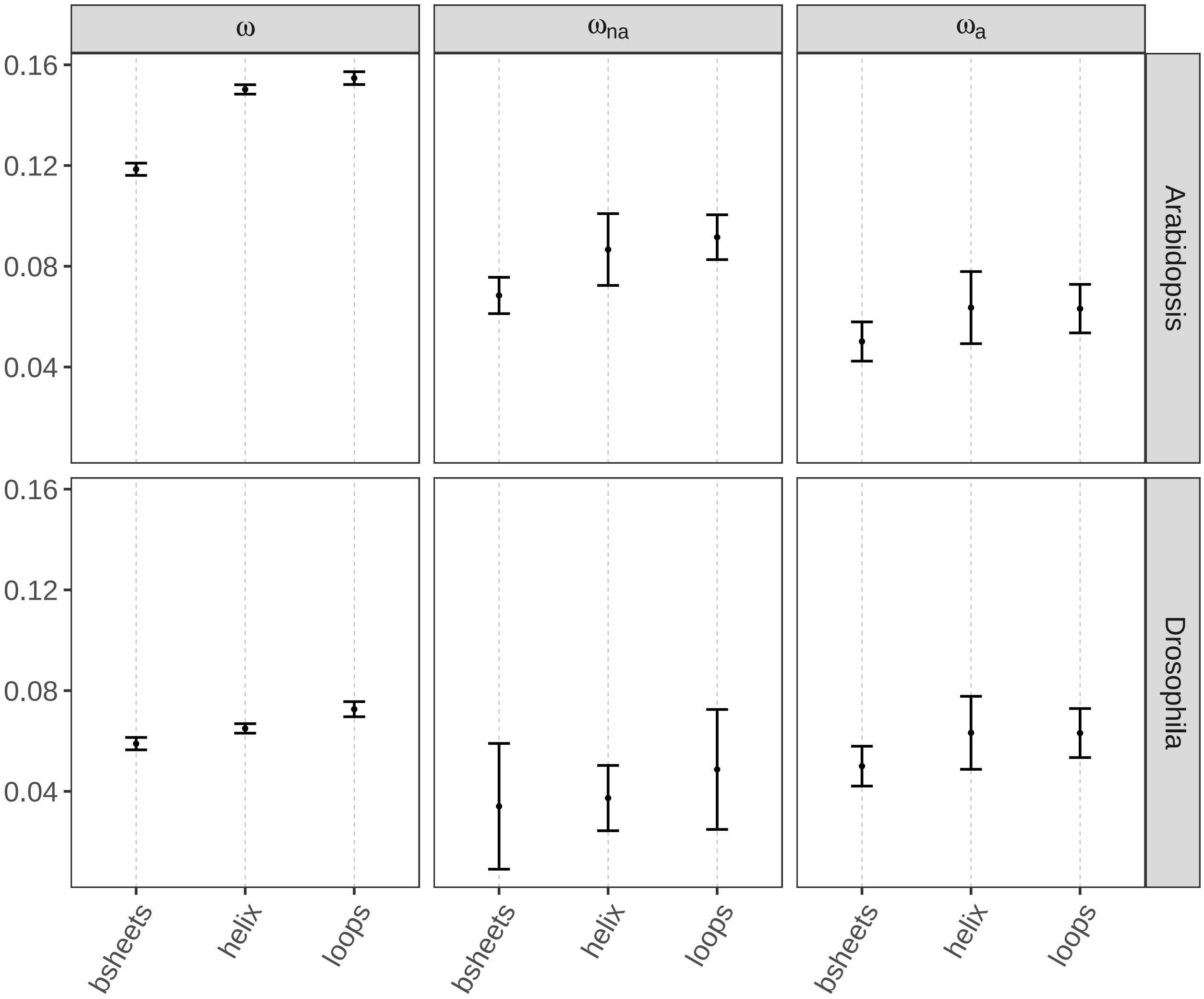


**Secondary Structure**

The next chunck of the script describes the statistical tests applied to the variables comparing two categories, specifically:
Binding affinity to dnaK

```
# libraries
library(rlist)
library(stringr)
#

######
###### Binding affinity to dnaK
######

# checking the number of replicates per var.value, variable
# and species after removing 1% outliers
dnak.nrep <- ddply(tbl.rates[[1]], c("species", "variable", "var.value"),
                   function(x) {
    nrep <- nrow(x)
    data.frame(nrep)
```

```r
})

# take the minimum number of replicates between categories
# take the minimum number of replicates between categories
min.nrep <- ddply(dnak.nrep, c("species", "variable"), function(x) {
  min.n <- min(x$nrep)
  data.frame(min.n)
})
kable(min.nrep)
```

| species | variable | min.n |
|---|---|---|
| Arabidopsis | dnds | 91 |
| Arabidopsis | omegaNA | 91 |
| Arabidopsis | omegaA | 91 |
| Drosophila | dnds | 57 |
| Drosophila | omegaNA | 57 |
| Drosophila | omegaA | 57 |

```r
# 91 replicates Arabidopsis
nrep.arab <- subset(tbl.rates[[1]], tbl.rates[[1]]$species == "Arabidopsis")
tbl.arab <- ddply(nrep.arab, c("species", "var.value", "variable"),
                  function(x) {
  x[sample(nrow(x), 91, replace = FALSE), ]
})

# 57 replicates for Drosophila
nrep.dmel <- subset(tbl.rates[[1]], tbl.rates[[1]]$species == "Drosophila")
tbl.dmel <- ddply(nrep.dmel, c("species", "var.value", "variable"),
                  function(x) {
  x[sample(nrow(x), 57, replace = FALSE), ]
})

dat.dnak <- rbind(tbl.arab, tbl.dmel)

# function to split the tables by the name of each variable
dnak.split <- by(dat.dnak, dat.dnak[,c("var.value")], function(y) y)

# to change the column name "value" to the respective id of the category
dnak.value <- lapply(dnak.split, function(x) {
  tbl <- data.frame(x)
  colnames(tbl)[5] <- as.character(unique(tbl$var.value))
  tbl <- tbl[,-3]
  return(tbl)
})

# binding all values as columns
names.vars <- dnak.value[[1]][,1:3]

tbl.estimate <- lapply(dnak.value, function(x) {
  val <- x[,4]
})
tbl.dnak <- list.cbind(tbl.estimate)
tbl.dnak <- as.data.frame(tbl.dnak)
tbl.dnak <- cbind(names.vars, tbl.dnak)
```

Table 1: Statistics for the binding affinity to dnaK

| variable | species | p.value |
|---|---|---|
| dnds | Arabidopsis | 0.0108696 |
| omegaNA | Arabidopsis | 0.3152174 |
| omegaA | Arabidopsis | 0.6847826 |

```
# now lets estimate the differences between the two variables
dnak.dif <- ddply(tbl.dnak, c("variable", "species"), function(x){
  dif <- x$Binder - x$NonBinder
  tbl <- data.frame(dif)
})

# Getting the p-value for each difference. Since we are sampling across
# bootstrap replicates, the p-values will be estimated 100 times and
# then the minimum p-value will be used

# Arabidopsis
arab.dif <- subset(dnak.dif, dnak.dif$species == "Arabidopsis")
nboots <- 100
arab.p <- list()
for (i in 1:nboots) {
  arab.p[[i]] <- ddply(arab.dif, c("variable", "species"), function(x, N=91){
    c <- as.numeric(nrow(x[x$dif < 0,]))
    c2 <- as.numeric(nrow(x[x$dif > 0,]))
    m <- min(c, c2)
    p <- (2*m+1)/(N+1)
    tbl <- data.frame(m, p)
  })
}
arab.p.tbl <- rbindlist(arab.p)
arab.pvalue <- ddply(arab.p.tbl, c("variable", "species"), function(x) {
  p.value <- min(x$p)
  data.frame(p.value)
})

# showing the table
kable(arab.pvalue, caption = "Statistics for the binding affinity to dnaK")
```

```
# Drosophila
dmel.dif <- subset(dnak.dif, dnak.dif$species == "Drosophila")
nboots <- 100
dmel.p <- list()
for (i in 1:nboots) {
  dmel.p[[i]] <- ddply(dmel.dif, c("variable", "species"), function(x, N=57){
    c <- as.numeric(nrow(x[x$dif < 0,]))
    c2 <- as.numeric(nrow(x[x$dif > 0,]))
    m <- min(c, c2)
    p <- (2*m+1)/(N+1)
    tbl <- data.frame(m, p)
  })
}
```

Table 2: Statistics for the binding affinity to dnaK

| variable | species | p.value |
|---|---|---|
| dnds | Drosophila | 0.0172414 |
| omegaNA | Drosophila | 0.0172414 |
| omegaA | Drosophila | 0.6724138 |

```
dmel.p.tbl <- rbindlist(dmel.p)
dmel.pvalue <- ddply(dmel.p.tbl, c("variable", "species"), function(x) {
  p.value <- min(x$p)
  data.frame(p.value)
})

# showing the table
kable(dmel.pvalue, caption = "Statistics for the binding affinity to dnaK")
```

The next section displays the statistical tests used for variables comparing three or more categories, specifically:
Protein Location
Secondary structure

```
# libraries
library(rlist)
library(stringr)
#

######
###### Protein Location
######

# checking the number of replicates per var.value, variable
# and species after removing 1% outliers
loc.nrep <- ddply(tbl.rates[[2]], c("species", "variable", "var.value"),
                  function(x) {
    nrep <- nrow(x)
    data.frame(nrep)
})

# take the minimum number of replicates between categories
min.nrep <- ddply(loc.nrep, c("species", "variable"), function(x) {
  min.n <- min(x$nrep)
  data.frame(min.n)
})
kable(min.nrep)
```

| species | variable | min.n |
|---|---|---|
| Arabidopsis | dnds | 77 |
| Arabidopsis | omegaNA | 77 |
| Arabidopsis | omegaA | 77 |
| Drosophila | dnds | 95 |
| Drosophila | omegaNA | 95 |
| Drosophila | omegaA | 95 |

```r
# 77 replicates Arabidopsis
nrep.arab <- subset(tbl.rates[[2]], tbl.rates[[2]]$species == "Arabidopsis")
tbl.arab <- ddply(nrep.arab, c("species", "var.value", "variable"),
                  function(x) {
  x[sample(nrow(x), 77, replace = FALSE), ]
})

# 95 replicates for Drosophila
nrep.dmel <- subset(tbl.rates[[2]], tbl.rates[[2]]$species == "Drosophila")
tbl.dmel <- ddply(nrep.dmel, c("species", "var.value", "variable"),
                  function(x) {
  x[sample(nrow(x), 95, replace = FALSE), ]
})

dat.loc <- rbind(tbl.arab, tbl.dmel)

# function to split the tables by the name of each variable
loc.split <- by(dat.loc, dat.loc[,c("var.value")], function(y) y)

# to change the column name "value" to the respective id of the category
loc.value <- lapply(loc.split, function(x) {
  tbl <- data.frame(x)
  colnames(tbl)[5] <- as.character(unique(tbl$var.value))
  tbl <- tbl[,-3]
  return(tbl)
})

# binding all values as columns
names.vars <- loc.value[[1]][,1:3]

tbl.estimate <- lapply(loc.value, function(x) {
  val <- x[,4]
})
tbl.loc <- list.cbind(tbl.estimate)
tbl.loc <- as.data.frame(tbl.loc)
tbl.loc <- cbind(tbl.loc, names.vars)
colnames(tbl.loc)[9] <- "estimate"

# estimate the differences between columns
# to do so will duplicate the data.frame in order to subtract each column
tbl2.loc <- tbl.loc[,1:6]

# doing the differences in a way that we count only for one of the differences
loc.dif <- cbind(tbl.loc[, c(8,9), drop=F],
        do.call(cbind, lapply(tbl2.loc[,2:6],
                              function(x) tbl.loc[,1:2]-x)),
        do.call(cbind, lapply(tbl2.loc[,4:6],
                              function(x) tbl.loc[,3:4]-x)))


# putting all variables in one column
# not counting with the columns comparing the same value
loc.hist <- melt(loc.dif, id.vars = c("estimate", "species"),
```

```
                 measure.vars = c(names(loc.dif[3]), names(loc.dif[5:13]),
                                  names(loc.dif[15:18])))

# puting NA values for when the difference is 0
# (i.e. when comparing the same variables)
split <- str_split_fixed(loc.hist$variable, "\\.", 2)
loc.hist$var1 <- split[,1]
loc.hist$var2 <- split[,2]

## getting the p-value for each difference. Since we are sampling across
# bootstrap replicates, the p-values will be estimated 100 times and
# then the minimum p-value will be used

## Arabidopsis
arab.hist <- subset(loc.hist, loc.hist$species == "Arabidopsis")
nboots <- 100
arab.p <- list()
for (i in 1:nboots) {
  arab.p[[i]] <- ddply(arab.hist, c("species", "estimate", "var1", "var2"),
                       function(x, N=77){
    c <- as.numeric(nrow(x[x$value < 0,]))
    c2 <- as.numeric(nrow(x[x$value > 0,]))
    m <- min(c, c2)
    p <- (2*m+1)/(N+1)
    tbl <- data.frame(m, p)
  })
}
# correcting the p-value for multiple testing
arab.p.adj <- lapply(arab.p, function(x) {
  ddply(x,  c("species","estimate", "var1", "var2"), function(x) {
    p.value <- p.adjust(x$p)
    data.frame(p.value)
  })
})

# taking the minimum p-value
tbl.arab.p.adj <- rbindlist(arab.p.adj)
arab.pvalue <- ddply(tbl.arab.p.adj, c("species","estimate", "var1", "var2"),
                     function(x) {
  p.value <- min(x$p.value)
  data.frame(p.value)
})

# showing the table
kable(arab.pvalue, caption= "Statistics for protein location")
```

```
## Drosophila
dmel.hist <- subset(loc.hist, loc.hist$species == "Drosophila")
nboots <- 100
dmel.p <- list()
for (i in 1:100) {
  dmel.p[[i]] <- ddply(dmel.hist, c("species", "estimate", "var1", "var2"),
                       function(x, N=95){
    c <- as.numeric(nrow(x[x$value < 0,]))
```

Table 3: Statistics for protein location

| species | estimate | var1 | var2 | p.value |
|---|---|---|---|---|
| Arabidopsis | dnds | EndomembraneSystem | Cytoplasmic | 0.0128205 |
| Arabidopsis | dnds | Mitochondrial | Cytoplasmic | 0.0128205 |
| Arabidopsis | dnds | Mitochondrial | EndomembraneSystem | 0.0128205 |
| Arabidopsis | dnds | Nuclear | Cytoplasmic | 0.0128205 |
| Arabidopsis | dnds | Nuclear | EndomembraneSystem | 0.0128205 |
| Arabidopsis | dnds | Nuclear | Mitochondrial | 0.0128205 |
| Arabidopsis | dnds | PlasmaMembrane | Cytoplasmic | 0.0128205 |
| Arabidopsis | dnds | PlasmaMembrane | EndomembraneSystem | 0.2179487 |
| Arabidopsis | dnds | PlasmaMembrane | Mitochondrial | 0.2948718 |
| Arabidopsis | dnds | PlasmaMembrane | Nuclear | 0.0128205 |
| Arabidopsis | dnds | Secreted | Cytoplasmic | 0.0128205 |
| Arabidopsis | dnds | Secreted | EndomembraneSystem | 0.0128205 |
| Arabidopsis | dnds | Secreted | Mitochondrial | 0.0128205 |
| Arabidopsis | dnds | Secreted | Nuclear | 0.4487179 |
| Arabidopsis | omegaNA | EndomembraneSystem | Cytoplasmic | 0.0384615 |
| Arabidopsis | omegaNA | Mitochondrial | Cytoplasmic | 0.1410256 |
| Arabidopsis | omegaNA | Mitochondrial | EndomembraneSystem | 0.7820513 |
| Arabidopsis | omegaNA | Nuclear | Cytoplasmic | 0.2948718 |
| Arabidopsis | omegaNA | Nuclear | EndomembraneSystem | 0.0128205 |
| Arabidopsis | omegaNA | Nuclear | Mitochondrial | 0.0128205 |
| Arabidopsis | omegaNA | PlasmaMembrane | Cytoplasmic | 0.0128205 |
| Arabidopsis | omegaNA | PlasmaMembrane | EndomembraneSystem | 0.4230769 |
| Arabidopsis | omegaNA | PlasmaMembrane | Mitochondrial | 0.3461538 |
| Arabidopsis | omegaNA | PlasmaMembrane | Nuclear | 0.0128205 |
| Arabidopsis | omegaNA | Secreted | Cytoplasmic | 0.1923077 |
| Arabidopsis | omegaNA | Secreted | EndomembraneSystem | 0.0128205 |
| Arabidopsis | omegaNA | Secreted | Mitochondrial | 0.0128205 |
| Arabidopsis | omegaNA | Secreted | Nuclear | 0.6538462 |
| Arabidopsis | omegaA | EndomembraneSystem | Cytoplasmic | 0.6282051 |
| Arabidopsis | omegaA | Mitochondrial | Cytoplasmic | 0.9871795 |
| Arabidopsis | omegaA | Mitochondrial | EndomembraneSystem | 0.2692308 |
| Arabidopsis | omegaA | Nuclear | Cytoplasmic | 0.2435897 |
| Arabidopsis | omegaA | Nuclear | EndomembraneSystem | 0.4487179 |
| Arabidopsis | omegaA | Nuclear | Mitochondrial | 0.1410256 |
| Arabidopsis | omegaA | PlasmaMembrane | Cytoplasmic | 0.4230769 |
| Arabidopsis | omegaA | PlasmaMembrane | EndomembraneSystem | 0.8333333 |
| Arabidopsis | omegaA | PlasmaMembrane | Mitochondrial | 0.0897436 |
| Arabidopsis | omegaA | PlasmaMembrane | Nuclear | 0.5256410 |
| Arabidopsis | omegaA | Secreted | Cytoplasmic | 0.1923077 |
| Arabidopsis | omegaA | Secreted | EndomembraneSystem | 0.5256410 |
| Arabidopsis | omegaA | Secreted | Mitochondrial | 0.2179487 |
| Arabidopsis | omegaA | Secreted | Nuclear | 0.8076923 |

```
    c2 <- as.numeric(nrow(x[x$value > 0,]))
    m <- min(c, c2)
    p <- (2*m+1)/(N+1)
    tbl <- data.frame(m, p)
  })
}

## correcting the p-value for multiple testing
dmel.p.adj <- lapply(dmel.p, function(x) {
  ddply(x,  c("species","estimate", "var1", "var2"),
        function(x) {
    p.value <- p.adjust(x$p)
    data.frame(p.value)
  })
})

# taking the minimum p-value
tbl.dmel.p.adj <- rbindlist(dmel.p.adj)
dmel.pvalue <- ddply(tbl.dmel.p.adj, c("species","estimate", "var1", "var2"),
                     function(x) {
  p.value <- min(x$p.value)
  data.frame(p.value)
})

# showing the table
kable(dmel.pvalue, caption = "Statistics for protein location")
```

```
######
###### Secondary Structure
######

# checking the number of replicates per var.value,
# variable and species after removing 1% outliers
SS.nrep <- ddply(tbl.rates[[3]], c("species", "variable", "var.value"),
                 function(x) {
    nrep <- nrow(x)
    data.frame(nrep)
})

# take the minimum number of replicates between categories
min.nrep <- ddply(SS.nrep, c("species", "variable"),
                  function(x) {
  min.n <- min(x$nrep)
  data.frame(min.n)
})
kable(min.nrep)
```

| species | variable | min.n |
|---|---|---|
| Arabidopsis | dnds | 98 |
| Arabidopsis | omegaNA | 98 |
| Arabidopsis | omegaA | 98 |
| Drosophila | dnds | 95 |
| Drosophila | omegaNA | 95 |
| Drosophila | omegaA | 95 |

Table 4: Statistics for protein location

| species | estimate | var1 | var2 | p.value |
|---|---|---|---|---|
| Drosophila | dnds | EndomembraneSystem | Cytoplasmic | 0.4270833 |
| Drosophila | dnds | Mitochondrial | Cytoplasmic | 0.1145833 |
| Drosophila | dnds | Mitochondrial | EndomembraneSystem | 0.0312500 |
| Drosophila | dnds | Nuclear | Cytoplasmic | 0.0104167 |
| Drosophila | dnds | Nuclear | EndomembraneSystem | 0.0104167 |
| Drosophila | dnds | Nuclear | Mitochondrial | 0.0104167 |
| Drosophila | dnds | PlasmaMembrane | Cytoplasmic | 0.0104167 |
| Drosophila | dnds | PlasmaMembrane | EndomembraneSystem | 0.0312500 |
| Drosophila | dnds | PlasmaMembrane | Mitochondrial | 0.0937500 |
| Drosophila | dnds | PlasmaMembrane | Nuclear | 0.0104167 |
| Drosophila | dnds | Secreted | Cytoplasmic | 0.0104167 |
| Drosophila | dnds | Secreted | EndomembraneSystem | 0.0104167 |
| Drosophila | dnds | Secreted | Mitochondrial | 0.0104167 |
| Drosophila | dnds | Secreted | Nuclear | 0.7812500 |
| Drosophila | omegaNA | EndomembraneSystem | Cytoplasmic | 0.8437500 |
| Drosophila | omegaNA | Mitochondrial | Cytoplasmic | 0.3437500 |
| Drosophila | omegaNA | Mitochondrial | EndomembraneSystem | 0.9270833 |
| Drosophila | omegaNA | Nuclear | Cytoplasmic | 0.2812500 |
| Drosophila | omegaNA | Nuclear | EndomembraneSystem | 0.3020833 |
| Drosophila | omegaNA | Nuclear | Mitochondrial | 0.2812500 |
| Drosophila | omegaNA | PlasmaMembrane | Cytoplasmic | 0.3020833 |
| Drosophila | omegaNA | PlasmaMembrane | EndomembraneSystem | 0.1770833 |
| Drosophila | omegaNA | PlasmaMembrane | Mitochondrial | 0.1145833 |
| Drosophila | omegaNA | PlasmaMembrane | Nuclear | 0.0729167 |
| Drosophila | omegaNA | Secreted | Cytoplasmic | 0.0937500 |
| Drosophila | omegaNA | Secreted | EndomembraneSystem | 0.3437500 |
| Drosophila | omegaNA | Secreted | Mitochondrial | 0.3645833 |
| Drosophila | omegaNA | Secreted | Nuclear | 0.9687500 |
| Drosophila | omegaA | EndomembraneSystem | Cytoplasmic | 0.9062500 |
| Drosophila | omegaA | Mitochondrial | Cytoplasmic | 0.3437500 |
| Drosophila | omegaA | Mitochondrial | EndomembraneSystem | 0.2395833 |
| Drosophila | omegaA | Nuclear | Cytoplasmic | 0.1562500 |
| Drosophila | omegaA | Nuclear | EndomembraneSystem | 0.3854167 |
| Drosophila | omegaA | Nuclear | Mitochondrial | 0.0520833 |
| Drosophila | omegaA | PlasmaMembrane | Cytoplasmic | 0.5520833 |
| Drosophila | omegaA | PlasmaMembrane | EndomembraneSystem | 0.8437500 |
| Drosophila | omegaA | PlasmaMembrane | Mitochondrial | 0.1979167 |
| Drosophila | omegaA | PlasmaMembrane | Nuclear | 0.1354167 |
| Drosophila | omegaA | Secreted | Cytoplasmic | 0.3229167 |
| Drosophila | omegaA | Secreted | EndomembraneSystem | 0.4895833 |
| Drosophila | omegaA | Secreted | Mitochondrial | 0.1354167 |
| Drosophila | omegaA | Secreted | Nuclear | 0.9895833 |

```r
# 98 replicates Arabidopsis
nrep.arab <- subset(tbl.rates[[3]], tbl.rates[[3]]$species == "Arabidopsis")
tbl.arab <- ddply(nrep.arab, c("species", "var.value", "variable"),
                  function(x) {
  x[sample(nrow(x), 98), ]
})

# 95 replicates for Drosophila
nrep.dmel <- subset(tbl.rates[[3]], tbl.rates[[3]]$species == "Drosophila")
tbl.dmel <- ddply(nrep.dmel, c("species", "var.value", "variable"),
                  function(x) {
  x[sample(nrow(x), 95), ]
})

dat.SS <- rbind(tbl.arab, tbl.dmel)

# function to split the tables by the name of each variable
SS.split <- by(dat.SS, dat.SS[,c("var.value")], function(y) y)

# to change the column name "value" to the respective id of the category
SS.value <- lapply(SS.split, function(x) {
  tbl <- data.frame(x)
  colnames(tbl)[5] <- as.character(unique(tbl$var.value))
  tbl <- tbl[,-3]
  return(tbl)
})

# binding all values as columns
names.vars <- SS.value[[1]][,1:3]

tbl.estimate <- lapply(SS.value, function(x) {
  val <- x[,4]
})
tbl.SS <- list.cbind(tbl.estimate)
tbl.SS <- as.data.frame(tbl.SS)
tbl.SS <- cbind(tbl.SS, names.vars)
colnames(tbl.SS)[6] <- "estimate"

# estimate the differences between columns
# to do so will duplicate the data.frame in order to subtract each column
tbl2.SS <- tbl.SS[,1:3]

SS.dif <- cbind(tbl.SS[, c(5,6), drop=F],
        do.call(cbind, lapply(tbl2.SS[,1:2],
                              function(x) tbl.SS[,2:3]-x)))

# putting all variables in one column
SS.hist <- melt(SS.dif, id.vars = c("estimate", "species"),
                        measure.vars = c(names(SS.dif[3:4]),
                                         names(SS.dif[6])))

# puting NA values for when the difference is 0
# (i.e. when comparing the same variables)
```

```r
split <- str_split_fixed(SS.hist$variable, "\\.", 2)
SS.hist$var1 <- split[,2]
SS.hist$var2 <- split[,1]

## getting the p-value for each difference
## N is not 100 because we took out the values that were outliers
## Arabidopsis
arab.hist <- subset(SS.hist, SS.hist$species == "Arabidopsis")
nboots <- 100
arab.p <- list()
for (i in 1:100) {
  arab.p[[i]] <- ddply(arab.hist, c("species","estimate", "var1", "var2"),
                       function(x, N=98){
    c <- as.numeric(nrow(x[x$value < 0,]))
    c2 <- as.numeric(nrow(x[x$value > 0,]))
    m <- min(c, c2)
    p <- (2*m+1)/(N+1)
    tbl <- data.frame(c, c2, m, p)
  })
}

# correcting the p-value for multiple testing
arab.p.adj <- lapply(arab.p, function(x) {
  ddply(x,  c("species","estimate", "var1", "var2"),
        function(x) {
    p.value <- p.adjust(x$p)
    data.frame(p.value)
  })
})

# taking the minimum p-value
tbl.arab.p.adj <- rbindlist(arab.p.adj)
arab.pvalue <- ddply(tbl.arab.p.adj, c("species","estimate", "var1", "var2"),
                     function(x) {
  p.value <- min(x$p.value)
  data.frame(p.value)
})

# showing the table
kable(arab.pvalue, caption = "Statistics for secondary structure")
```

```r
## Drosophila
dmel.hist <- subset(SS.hist, SS.hist$species == "Drosophila")
nboots <- 100
dmel.p <- list()
for (i in 1:nboots) {
  dmel.p[[i]] <- ddply(dmel.hist, c("species","estimate", "var1", "var2"),
                       function(x, N=95){
    c <- as.numeric(nrow(x[x$value < 0,]))
    c2 <- as.numeric(nrow(x[x$value > 0,]))
    m <- min(c, c2)
    p <- (2*m+1)/(N+1)
    tbl <- data.frame(c, c2, m, p)
  })
```

Table 5: Statistics for secondary structure

| species | estimate | var1 | var2 | p.value |
|---|---|---|---|---|
| Arabidopsis | dnds | helix | bsheets | 0.0101010 |
| Arabidopsis | dnds | loops | bsheets | 0.0101010 |
| Arabidopsis | dnds | loops | helix | 0.0303030 |
| Arabidopsis | omegaNA | helix | bsheets | 0.0303030 |
| Arabidopsis | omegaNA | loops | bsheets | 0.0101010 |
| Arabidopsis | omegaNA | loops | helix | 0.5959596 |
| Arabidopsis | omegaA | helix | bsheets | 0.0707071 |
| Arabidopsis | omegaA | loops | bsheets | 0.0707071 |
| Arabidopsis | omegaA | loops | helix | 0.9191919 |

Table 6: Statistics for secondary structure

| species | estimate | var1 | var2 | p.value |
|---|---|---|---|---|
| Drosophila | dnds | helix | bsheets | 0.0104167 |
| Drosophila | dnds | loops | bsheets | 0.0104167 |
| Drosophila | dnds | loops | helix | 0.0104167 |
| Drosophila | omegaNA | helix | bsheets | 0.8229167 |
| Drosophila | omegaNA | loops | bsheets | 0.4062500 |
| Drosophila | omegaNA | loops | helix | 0.4270833 |
| Drosophila | omegaA | helix | bsheets | 0.0520833 |
| Drosophila | omegaA | loops | bsheets | 0.0729167 |
| Drosophila | omegaA | loops | helix | 0.9687500 |

```
}

# correcting the p-value for multiple testing
dmel.p.adj <- lapply(dmel.p, function(x) {
  ddply(x,  c("species","estimate", "var1", "var2"), function(x) {
    p.value <- p.adjust(x$p)
    tbl <- data.frame(p.value)
  })
})

# taking the minimum p-value
tbl.dmel.p.adj <- rbindlist(dmel.p.adj)
dmel.pvalue <- ddply(tbl.dmel.p.adj, c("species","estimate", "var1", "var2"),
                     function(x) {
  p.value <- min(x$p.value)
  data.frame(p.value)
})

# showing the table
kable(dmel.pvalue, caption = "Statistics for secondary structure")
```
