## supplementary File S2 for "The impact of protein architecture on adaptive evolution"

### Continuous variables

This is the script with all plots and statistical analysis regarding the analysis of continuous variables, refering to: Intrinsic Residue Disorder, Gene Expression, Number of Introns, Number of Protein-Protein Interactions, Proportion of Disordered Residues, Protein Length, Recombination Rate and Relative Solvent Accessibility.

A function was created to call all files for each analysis performed. In this respect the order of the files is as follows:
files.continuous[1] <- Intrinsic Residue Disorder
files.continuous[2] <- Gene Expression
files.continuous[3] <- Number of Introns
files.continuous[4] <- Number of Protein-Protein interactions
files.continuous[5] <- Proportion of Disordered Residues
files.continuous[6] <- Protein Length
files.continuous[7] <- Recombination Rate
files.continuous[8] <- Relative Solvent Accessibility (RSA)

```
tbl.stats[[6]]$variable <- factor(tbl.stats[[6]]$variable,
                                  levels = c("dnds", "omegaNA", "omegaA"))
levels(tbl.stats[[6]]$variable) <- c(expression(omega), expression(omega[na]),
                                   expression(omega[a]))
tbl.stats[[7]]$variable <- factor(tbl.stats[[7]]$variable,
                                  levels = c("dnds", "omegaNA", "omegaA"))
levels(tbl.stats[[7]]$variable) <- c(expression(omega), expression(omega[na]),
                                   expression(omega[a]))
tbl.stats[[8]]$variable <- factor(tbl.stats[[8]]$variable,
                                  levels = c("dnds", "omegaNA", "omegaA"))
levels(tbl.stats[[8]]$variable) <- c(expression(omega), expression(omega[na]),
                                   expression(omega[a]))
```

In the next chunk the script to plot the results is represented. The same order will follow.

```
# theme of the plot
theme.plot <- function(x) {
  theme(axis.title = element_text(face = "bold", color = "black", size=14),
        text = element_text(size=14),
        axis.title.x = element_text(margin = margin(t = 18, r = 10, b = 0, l = 0)),
        axis.title.y = element_text(margin = margin(t = 18, r = 10, b = 0, l = 0)),
        panel.grid.minor=element_blank(),
        panel.grid.major = element_line(colour = "grey", linetype = "dashed", size = 0.2),
        panel.grid.major.y=element_blank(),
        #strip.text.y = element_blank(),
        axis.text.x = element_text(angle = 60, hjust = 1))
}

# plotting each of the output tables
plot.continuous <- lapply(tbl.stats, function(x) {
  ggplot(x, aes(x = var.value, y = value.mean)) +
  geom_line(col = "black", size = 0.2)+
  geom_ribbon(aes(ymin=value.mean + 1.96*value.sd,
                  ymax=value.mean - 1.96*value.sd), alpha=0.2) +
  geom_point(size=.2)+
  facet_grid(species~variable, scales = "free_x", labeller = label_parsed) +
  ylab("") +
  xlab(as.character(x$var)) +
  scale_x_sqrt() +
  theme_bw() +
  theme.plot()
})

plot.continuous
```

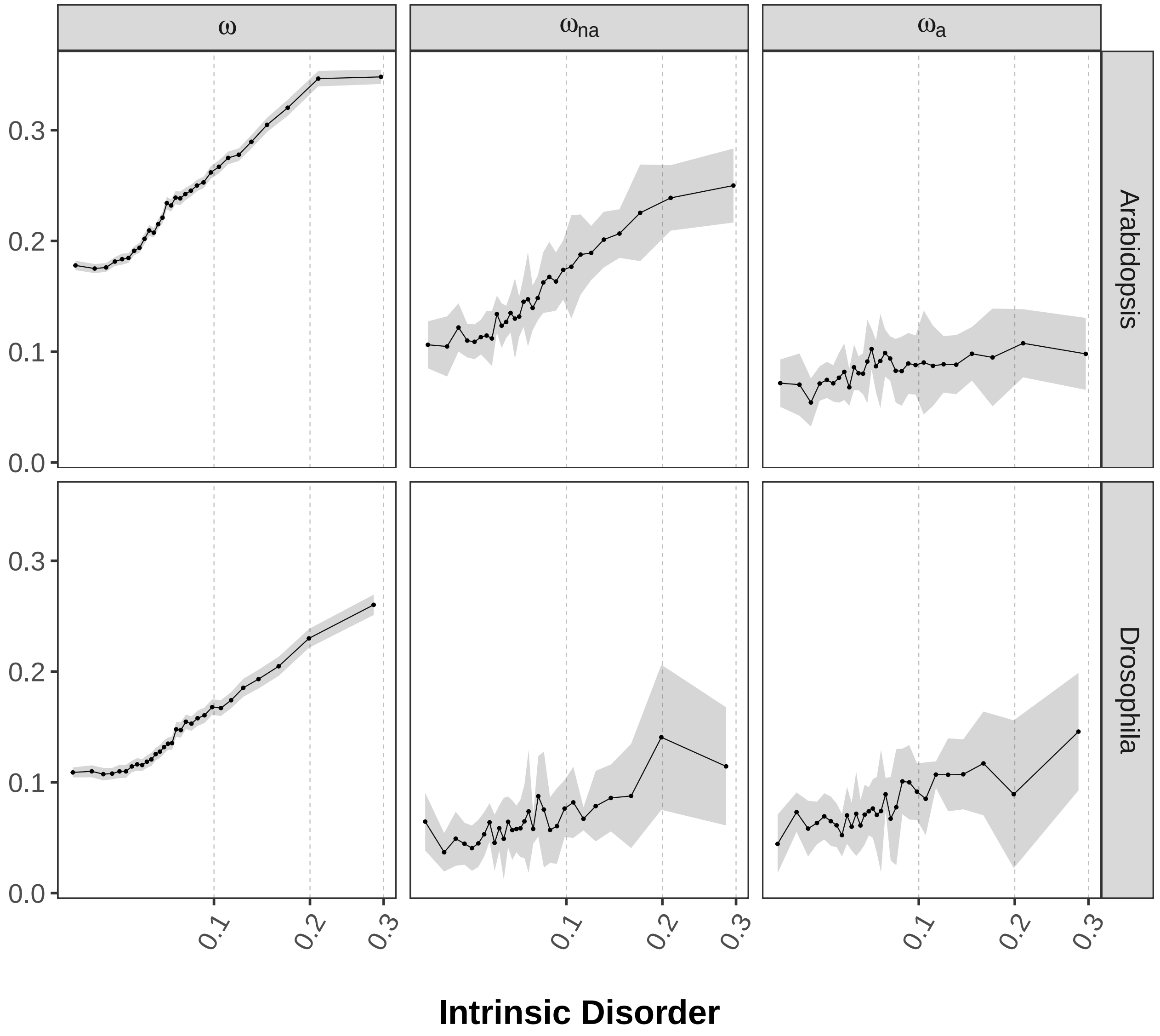
ω
$\omega_{na}$
$\omega_a$
Arabidopsis
Drosophila
0.0
0.1
0.2
0.3
0.1
0.2
0.3
Intrinsic Disorder

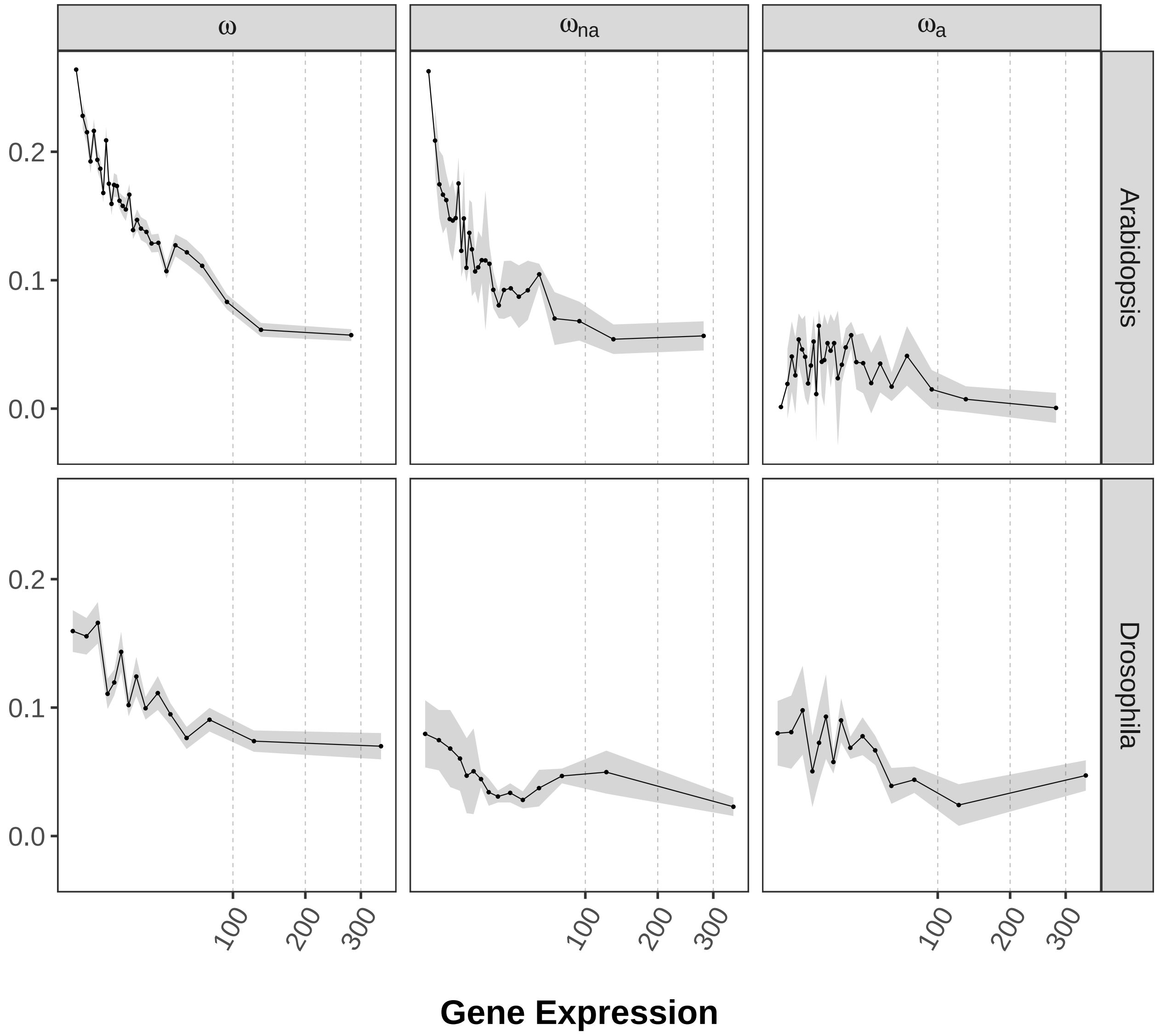
ω
ωna
ωa
Arabidopsis
Drosophila
0.0
0.1
0.2
100
200
300
Gene Expression

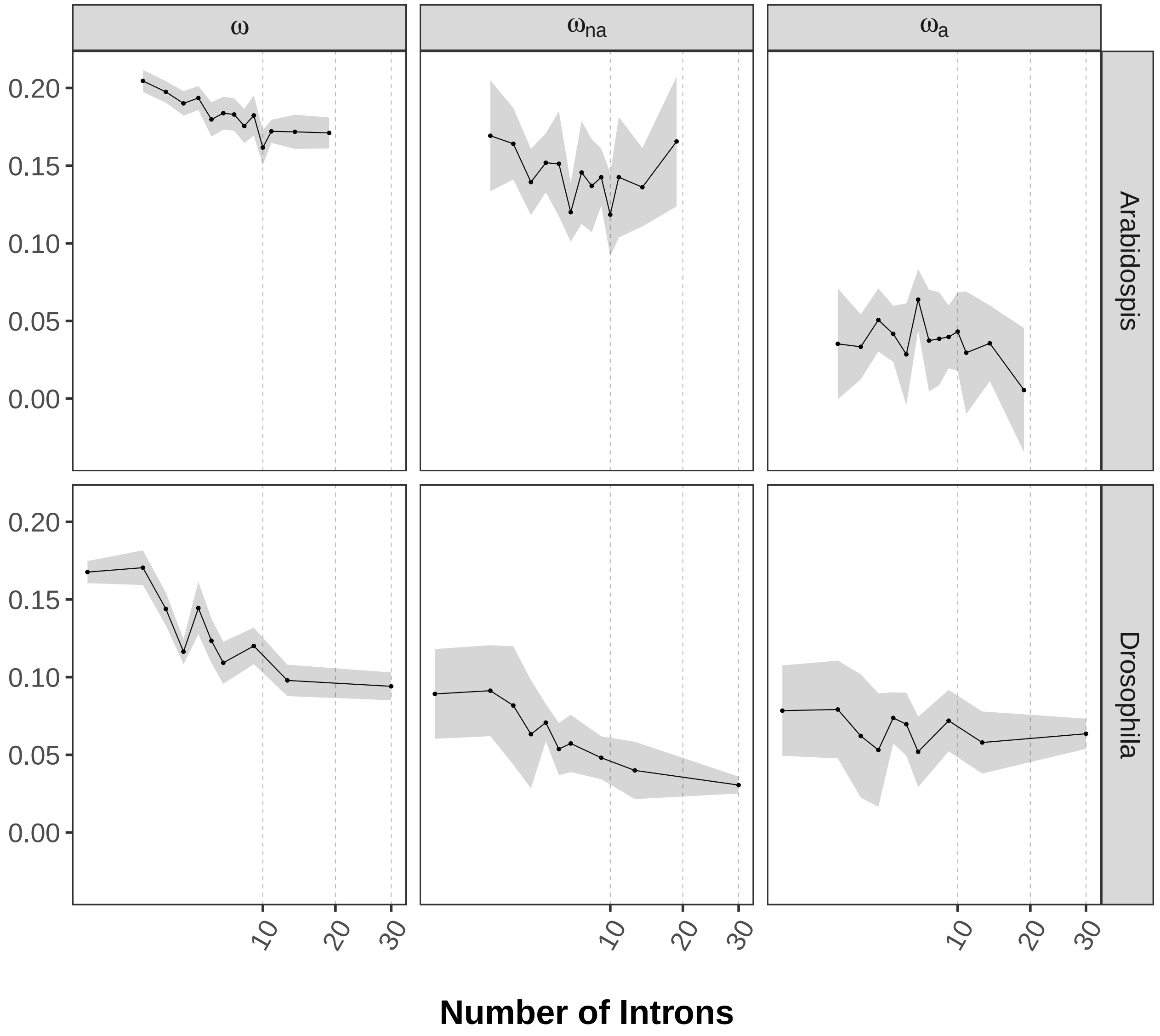
ω
ω_na
ω_a
Arabidospis
Drosophila
0.20
0.15
0.10
0.05
0.00
10
20
30
Number of Introns

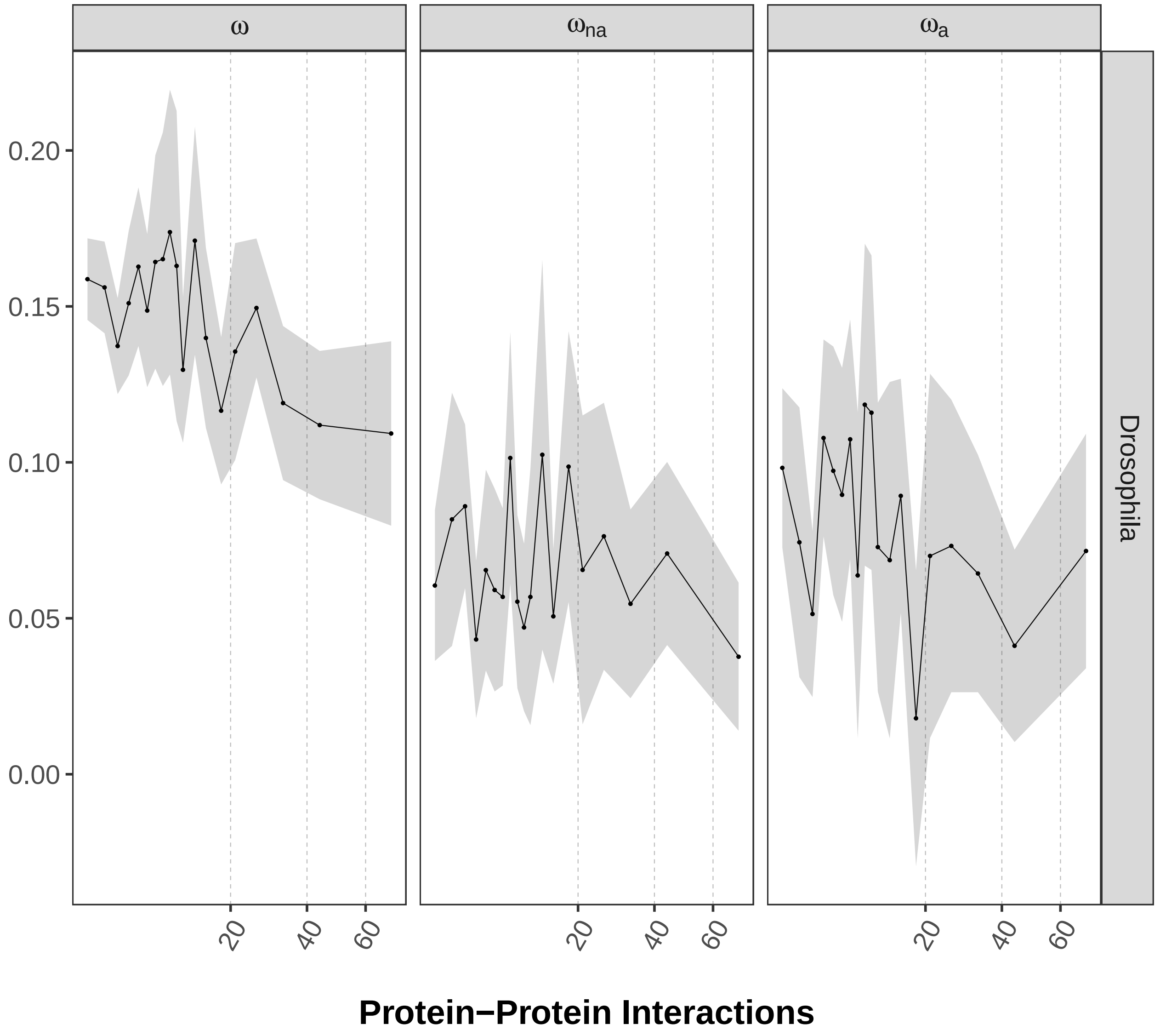

ω
ω_na
ω_a
Drosophila
0.20
0.15
0.10
0.05
0.00
20
40
60
Protein−Protein Interactions

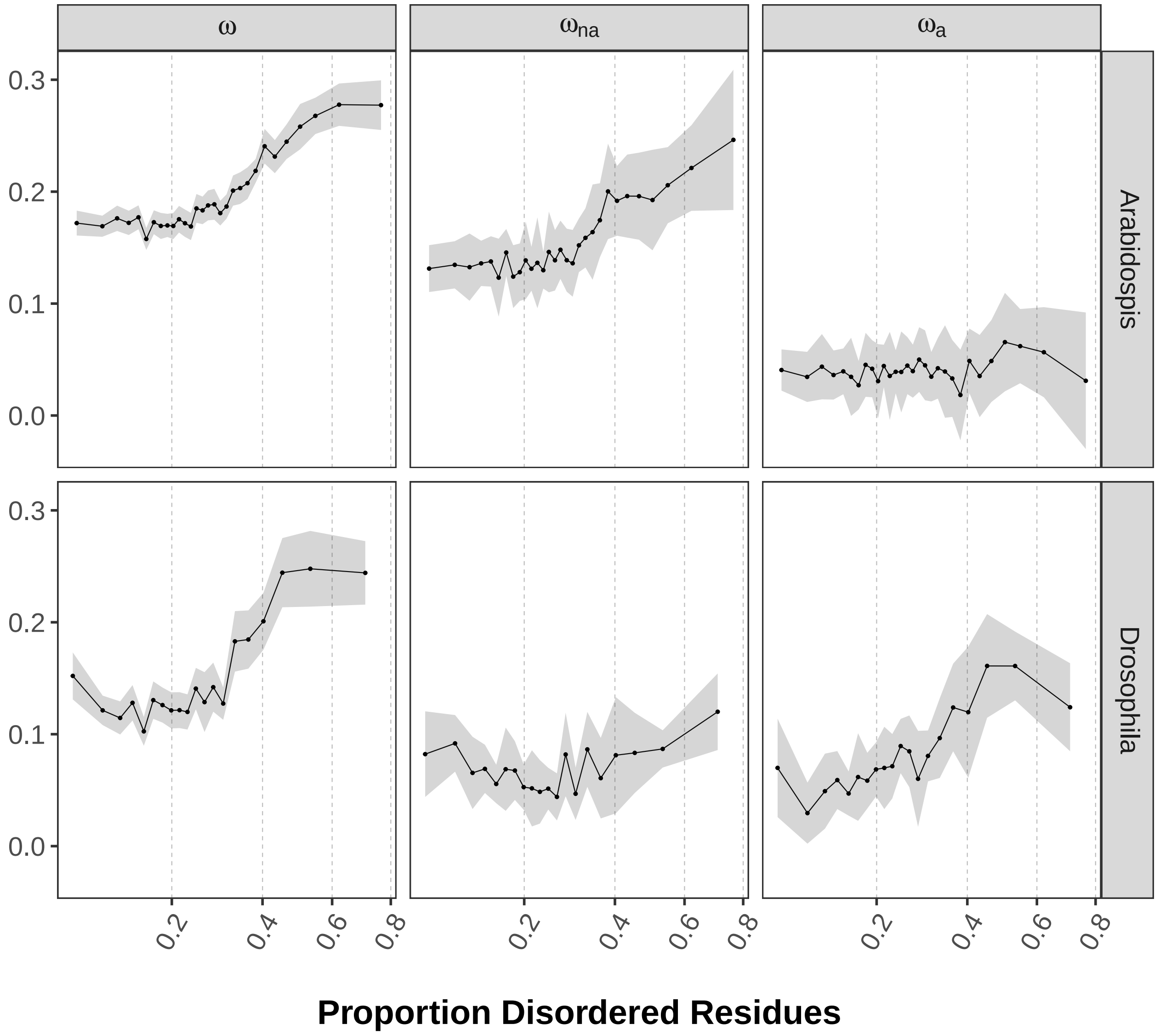
ω
$\omega_{na}$
$\omega_a$
Arabidospis
Drosophila
0.3
0.2
0.1
0.0
0.2
0.4
0.6
0.8
Proportion Disordered Residues

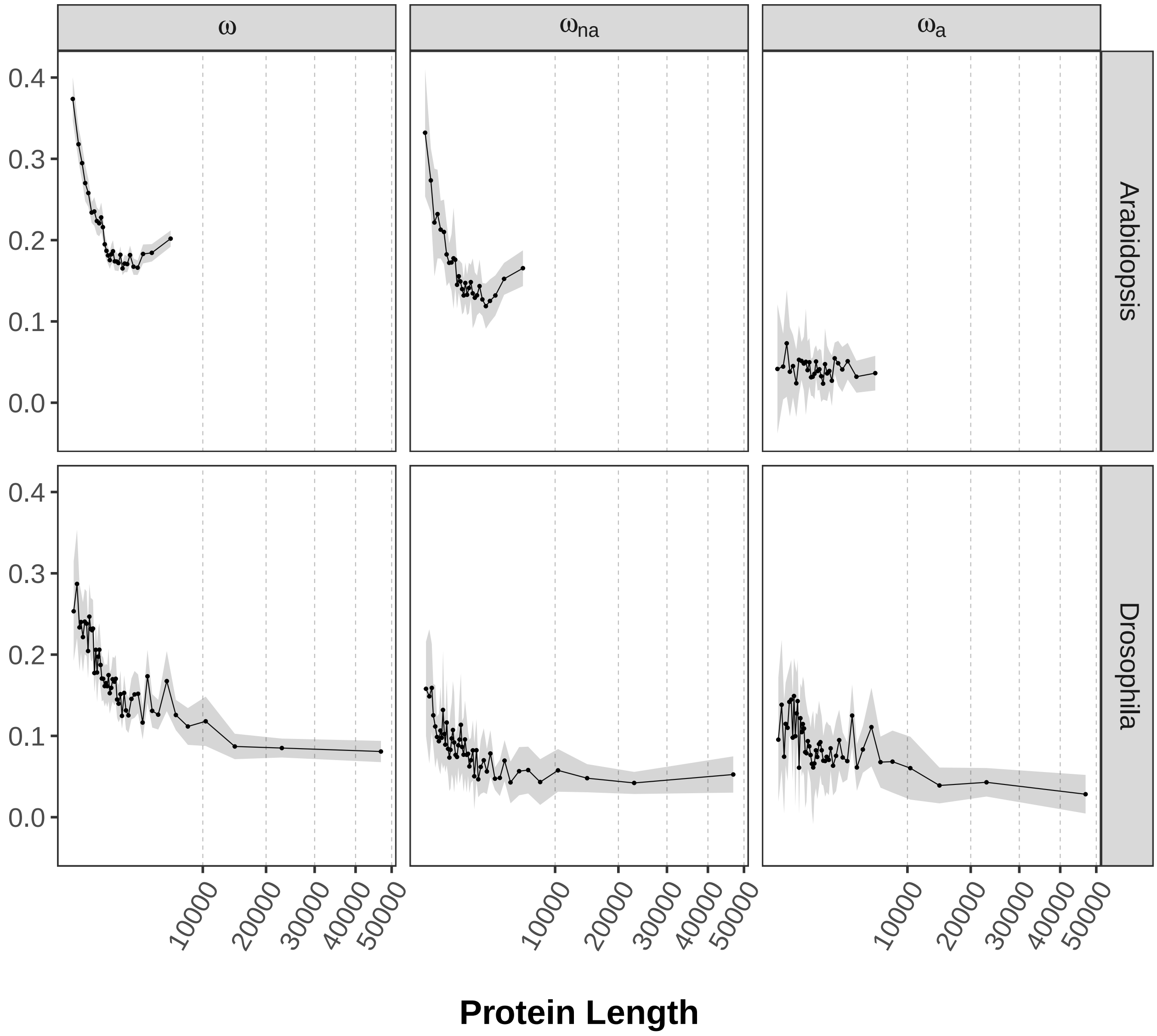

ω
$\omega_{na}$
$\omega_a$
Arabidopsis
Drosophila
0.4
0.3
0.2
0.1
0.0
10000
20000
30000
40000
50000
Protein Length

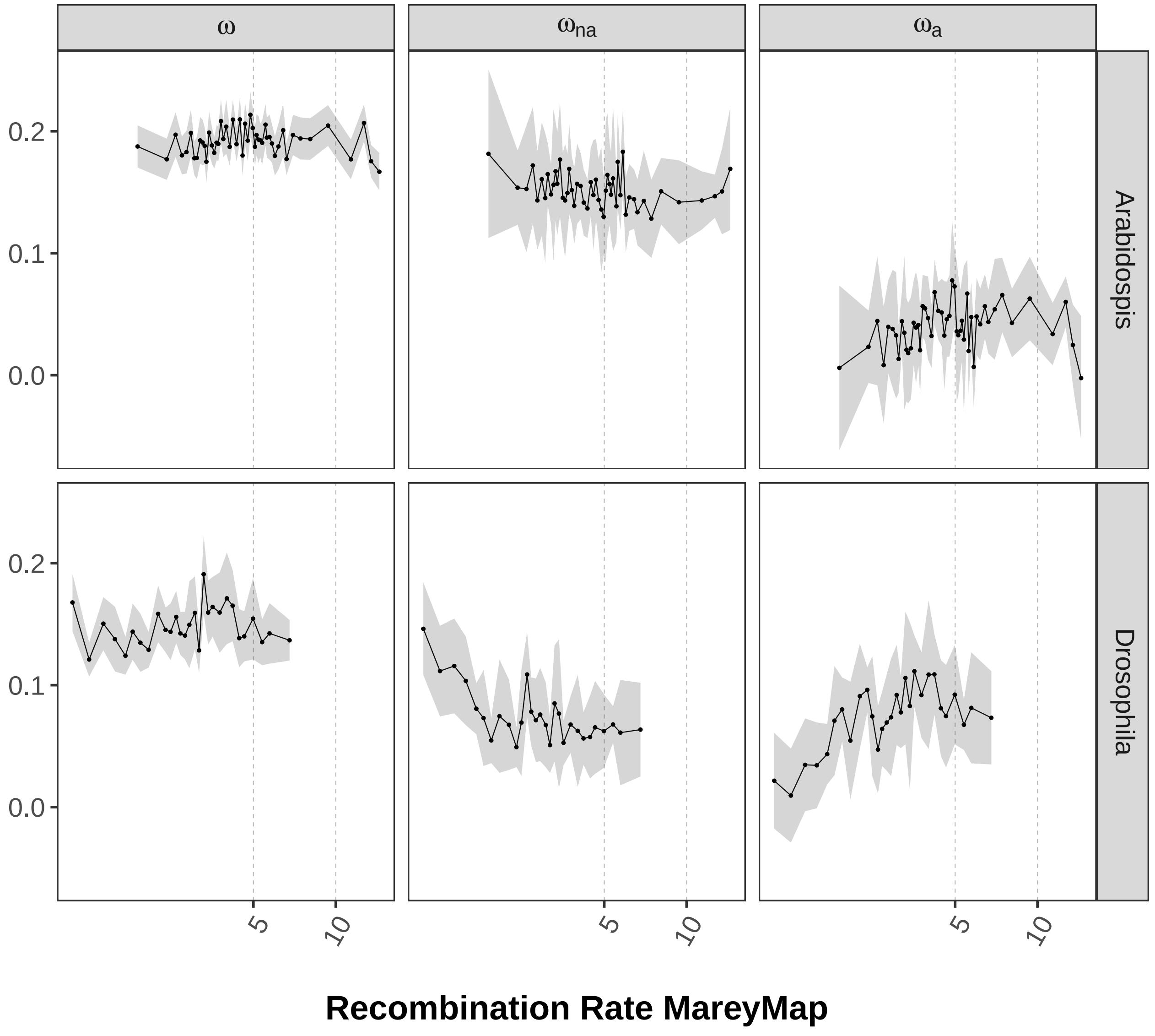
ω
ωna
ωa
Arabidospis
Drosophila
0.2
0.1
0.0
5
10
Recombination Rate MareyMap

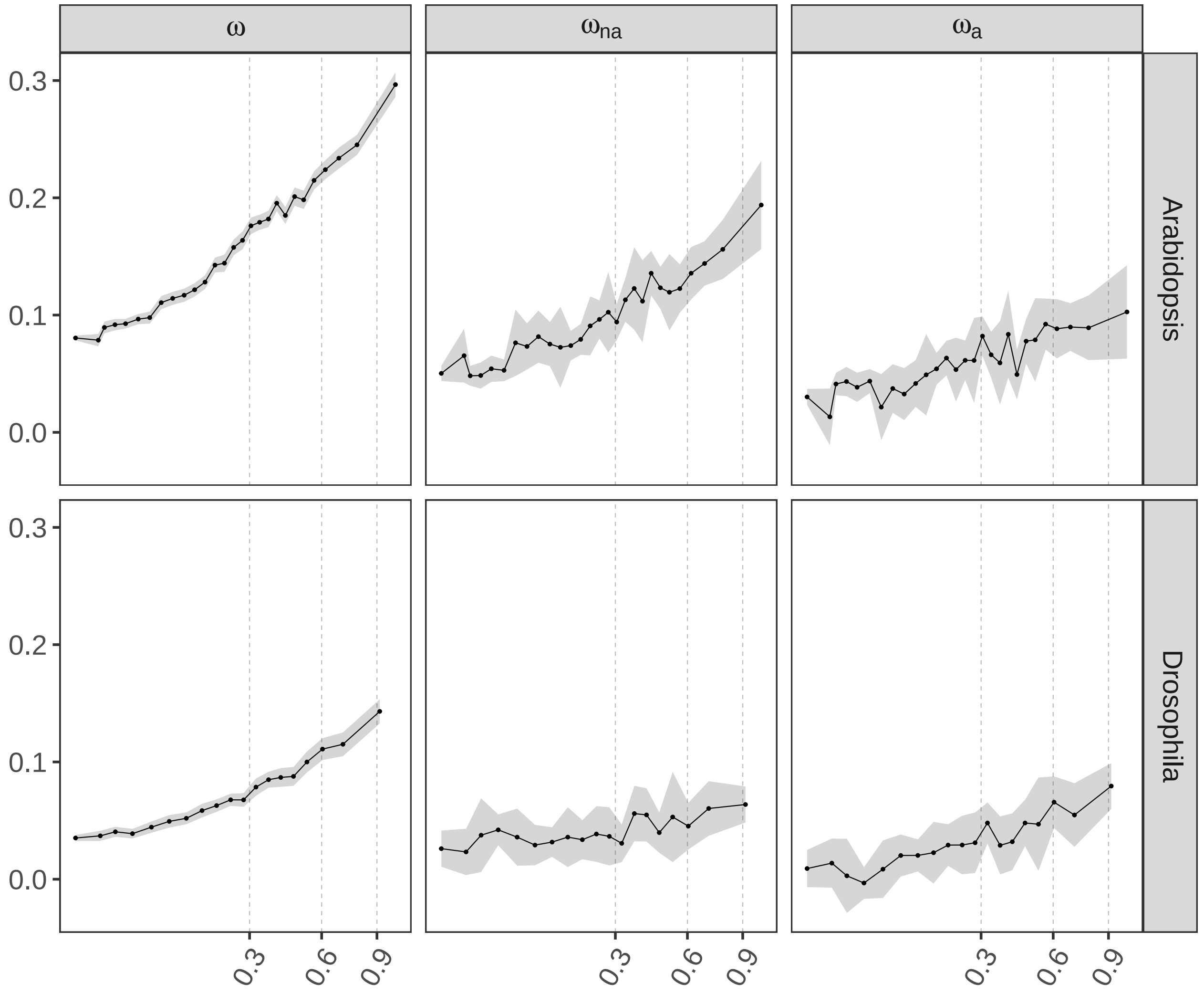


**Relative Solvent Accessibility**

The last section shows how the statistical analyses were performed.

```
stat <- lapply(tbl.stats, function(x) {
  ddply(x, c("var", "species", "variable"), function(x) {
    var <- as.numeric(factor(x$var.value))
    variable.value <- as.numeric(factor(x$value.mean))
    corr = cor.test(var, variable.value, method = "kendall", exact = FALSE)
    Kendall.tau = corr$estimate
    p.value = corr$p.value
    dat = data.frame(Kendall.tau, p.value)
  })
})

# showing the tables
for(i in stat) {
  print(kable(x = i, caption = paste0("Statistics for ", unique(i$var))))
}
```

Table 1: Statistics for Intrinsic Disorder

| var | species | variable | Kendall.tau | p.value |
|---|---|---|---|---|
| Intrinsic Disorder | Arabidopsis | omega | 0.9770115 | 0.0e+00 |
| Intrinsic Disorder | Arabidopsis | omega[na] | 0.9172414 | 0.0e+00 |
| Intrinsic Disorder | Arabidopsis | omega[a] | 0.6000000 | 3.2e-06 |
| Intrinsic Disorder | Drosophila | omega | 0.9540230 | 0.0e+00 |
| Intrinsic Disorder | Drosophila | omega[na] | 0.6689655 | 2.0e-07 |
| Intrinsic Disorder | Drosophila | omega[a] | 0.7057471 | 0.0e+00 |

Table 2: Statistics for Gene Expression

| var | species | variable | Kendall.tau | p.value |
|---|---|---|---|---|
| Gene Expression | Arabidopsis | omega | -0.8942529 | 0.0000000 |
| Gene Expression | Arabidopsis | omega[na] | -0.8160920 | 0.0000000 |
| Gene Expression | Arabidopsis | omega[a] | -0.1586207 | 0.2183112 |
| Gene Expression | Drosophila | omega | -0.7714286 | 0.0000611 |
| Gene Expression | Drosophila | omega[na] | -0.6190476 | 0.0012969 |
| Gene Expression | Drosophila | omega[a] | -0.5047619 | 0.0087205 |

Table 3: Statistics for Number of Introns

| var | species | variable | Kendall.tau | p.value |
|---|---|---|---|---|
| Number of Introns | Arabidospis | omega | -0.7948718 | 0.0001552 |
| Number of Introns | Arabidospis | omega[na] | -0.3589744 | 0.0875902 |
| Number of Introns | Arabidospis | omega[a] | -0.1538462 | 0.4641035 |
| Number of Introns | Drosophila | omega | -0.7333333 | 0.0031612 |
| Number of Introns | Drosophila | omega[na] | -0.8666667 | 0.0004862 |
| Number of Introns | Drosophila | omega[a] | -0.3333333 | 0.1797125 |

Table 4: Statistics for Protein-Protein Interactions

| var | species | variable | Kendall.tau | p.value |
|---|---|---|---|---|
| Protein-Protein Interactions | Drosophila | omega | -0.3684211 | 0.0275179 |
| Protein-Protein Interactions | Drosophila | omega[na] | -0.1111111 | 0.5062259 |
| Protein-Protein Interactions | Drosophila | omega[a] | -0.3099415 | 0.0637055 |

Table 5: Statistics for Proportion Disordered Residues

| var | species | variable | Kendall.tau | p.value |
|---|---|---|---|---|
| Proportion Disordered Residues | Arabidospis | omega | 0.7517241 | 0.0000000 |
| Proportion Disordered Residues | Arabidospis | omega[na] | 0.7333333 | 0.0000000 |
| Proportion Disordered Residues | Arabidospis | omega[a] | 0.1908046 | 0.1386584 |
| Proportion Disordered Residues | Drosophila | omega | 0.5684211 | 0.0004584 |
| Proportion Disordered Residues | Drosophila | omega[na] | 0.0631579 | 0.6970310 |
| Proportion Disordered Residues | Drosophila | omega[a] | 0.7263158 | 0.0000076 |

Table 6: Statistics for Protein Length

| var | species | variable | Kendall.tau | p.value |
|---|---|---|---|---|
| Protein Length | Arabidopsis | omega | -0.6781609 | 0.0000001 |
| Protein Length | Arabidopsis | omega[na] | -0.6735632 | 0.0000002 |
| Protein Length | Arabidopsis | omega[a] | -0.1310345 | 0.3091826 |
| Protein Length | Drosophila | omega | -0.7763265 | 0.0000000 |
| Protein Length | Drosophila | omega[na] | -0.6963265 | 0.0000000 |
| Protein Length | Drosophila | omega[a] | -0.4775510 | 0.0000010 |

Table 7: Statistics for Recombination Rate MareyMap

| var | species | variable | Kendall.tau | p.value |
|---|---|---|---|---|
| Recombination Rate MareyMap | Arabidospis | omega | 0.0857143 | 0.3797755 |
| Recombination Rate MareyMap | Arabidospis | omega[na] | -0.2212245 | 0.0233978 |
| Recombination Rate MareyMap | Arabidospis | omega[a] | 0.2065306 | 0.0343185 |
| Recombination Rate MareyMap | Drosophila | omega | 0.0758621 | 0.5560263 |
| Recombination Rate MareyMap | Drosophila | omega[na] | -0.4022989 | 0.0017952 |
| Recombination Rate MareyMap | Drosophila | omega[a] | 0.3839080 | 0.0028876 |

Table 8: Statistics for Relative Solvent Accessibility

| var | species | variable | Kendall.tau | p.value |
|---|---|---|---|---|
| Relative Solvent Accessibility | Arabidopsis | omega | 0.9841270 | 0.0000000 |
| Relative Solvent Accessibility | Arabidopsis | omega[na] | 0.8465608 | 0.0000000 |
| Relative Solvent Accessibility | Arabidopsis | omega[a] | 0.7513228 | 0.0000000 |
| Relative Solvent Accessibility | Drosophila | omega | 0.9766082 | 0.0000000 |
| Relative Solvent Accessibility | Drosophila | omega[na] | 0.5789474 | 0.0005331 |
| Relative Solvent Accessibility | Drosophila | omega[a] | 0.8128655 | 0.0000012 |
