## supplementary File S4 for "The impact of protein architecture on adaptive evolution"

### Protein Functional Class

This notebook describes all plots and statistical analysis performed with the categories of protein functional class.

The first part of the script removes bootstrap replicates for which the fitness effects parameters were not successfully fitted. For this purpose we discard 1% of the values above the maximum and below the minimum of each of the four parameters of fitness effects: Geman.neg, Gshape.neg, Gmean.neg and prop.pos.

```
setwd("/Users/moutinho/Dropbox/Data/Discrete/FunctionalClass/")

# Libraries
library(plyr)
library(dplyr)
library(data.table)
library(ggplot2)
library(reshape2)
library(doBy)
library(knitr)
library(kableExtra)
#

# calling all output tables
tbl.keggsmall <- read.table(file = "SmallProteinClass.csv", sep = "\t", header = TRUE)

## remove the outliers: 1% of the replicates below the min and
## 1% above the maximum

sub.keggSmall <- ddply(tbl.keggsmall, c("species", "var.value"), function(x) {
  sum.gmeanNeg <- summary(x$Gmean.neg)
  gmeanNeg.min1 <- as.numeric(sum.gmeanNeg[1]) + 0.01*as.numeric(sum.gmeanNeg[1])
  gmeanNeg.max1 <- as.numeric(sum.gmeanNeg[6]) - 0.01*as.numeric(sum.gmeanNeg[6])
  sum.gshapeNeg <- summary(x$Gshape.neg)
  gshapeNeg.min1 <- as.numeric(sum.gshapeNeg[1]) + 0.01*as.numeric(sum.gshapeNeg[1])
  gshapeNeg.max1 <- as.numeric(sum.gshapeNeg[6]) - 0.01*as.numeric(sum.gshapeNeg[6])
  sum.gmeanPos <- summary(x$Gmean.pos)
  gmeanPos.min1 <- as.numeric(sum.gmeanPos[1]) + 0.01*as.numeric(sum.gmeanPos[1])
  gmeanPos.max1 <- as.numeric(sum.gmeanPos[6]) - 0.01*as.numeric(sum.gmeanPos[6])
  sum.propPos <- summary(x$prop.pos)
  propPos.min1 <- as.numeric(sum.propPos[1]) + 0.01*as.numeric(sum.propPos[1])
  propPos.max1 <- as.numeric(sum.propPos[6]) - 0.01*as.numeric(sum.propPos[6])
  tbl <- x[!(x$Gmean.neg < gmeanNeg.min1 | x$Gmean.neg > gmeanNeg.max1 &
             x$Gshape.neg < gshapeNeg.min1 | x$Gshape.neg > gshapeNeg.max1 &
             x$Gmean.pos < gmeanPos.min1 | x$Gmean.pos > gmeanPos.max1 &
             x$prop.pos < propPos.min1 | x$prop.pos > propPos.max1),]
})
```

In the next chunk will take only the estimates concerning the rate of adaptive and non-adaptive substitutions, particularly: dnds, omegaNA and omegaA.

```
# In order to keep only the variables that we want to plot:
tbl.rates <- ddply(sub.keggSmall, c("species"), function(x) {
    melt(x, id.vars = c("var.value"), measure.vars = c("dnds", "omegaNA", "omegaA"))
})
```

```r
# function to estimate the mean and satandard deviation to plot the results with the
# mean of the bootstrap replicates and the 95% confidence interval

fun <- function(x){
  c(mean=mean(x), sd=sd(x))
}

# applying the above function to each output table for each value of each estimate
# (dnds, omegaA, omegaNA) for each value of the variable being analyzed for each species

tbl.stats <- summaryBy(value ~ variable + var.value + species, data=tbl.rates, FUN = fun)

# to change the estimate name to the respective symbol
tbl.stats$variable <- factor(tbl.stats$variable, levels = c("dnds", "omegaNA", "omegaA"))
levels(tbl.stats$variable) <- c(expression(omega), expression(omega[na]),
                                  expression(omega[a]))
```

The next chunck of the script shows the code used for plotting the results. The plots were done separately for the two species because the categories analysed did not overlap completely.

```r
# theme of the plot
theme.plot <- function(x) {
  theme(axis.title = element_text(face = "bold", color = "black", size=12,
                                  family = "Times"),
        text = element_text(size=12),
        axis.title.x = element_text(margin = margin(t = 9, r = 18, b = 5, l = 5)),
        axis.title.y = element_text(margin = margin(t = 9, r = 18, b = 5, l = 5)),
        panel.grid.minor=element_blank(),
        panel.grid.major = element_line(colour = "grey", linetype = "dashed", size = 0.1),
        panel.grid.major.y=element_blank(),
        strip.text.y = element_blank(),
        axis.text.x = element_text(angle = 60, hjust = 1),
        strip.background.x = element_rect(colour = "grey", fill = "gray92"),
        panel.spacing = unit(0.75, "lines"))
}

###
### Arabidopsis
###

stat.arab <- subset(tbl.stats, tbl.stats$species == "Arabidopsis")

# to order the plot according to values of omegaA
omegaA.arab <- subset(stat.arab, stat.arab$variable == "omega[a]")
omegaA.arab <- omegaA.arab[order(omegaA.arab[,4]),]

plot.arab <- ggplot(stat.arab, aes(x = var.value, y = value.mean)) +
  geom_point(size=1, col = "black") +
  geom_errorbar(aes(ymin=value.mean + 1.96*value.sd,
                    ymax=value.mean - 1.96*value.sd), width = .2) +
  scale_x_discrete(limits = as.character(omegaA.arab$var.value)) +
  ylab("") +
  xlab("Protein Functional Classification") +
  facet_grid(~variable, labeller = label_parsed) +
```

```r
  theme_bw() +
  theme.plot() +
  coord_flip()

plot.arab
```

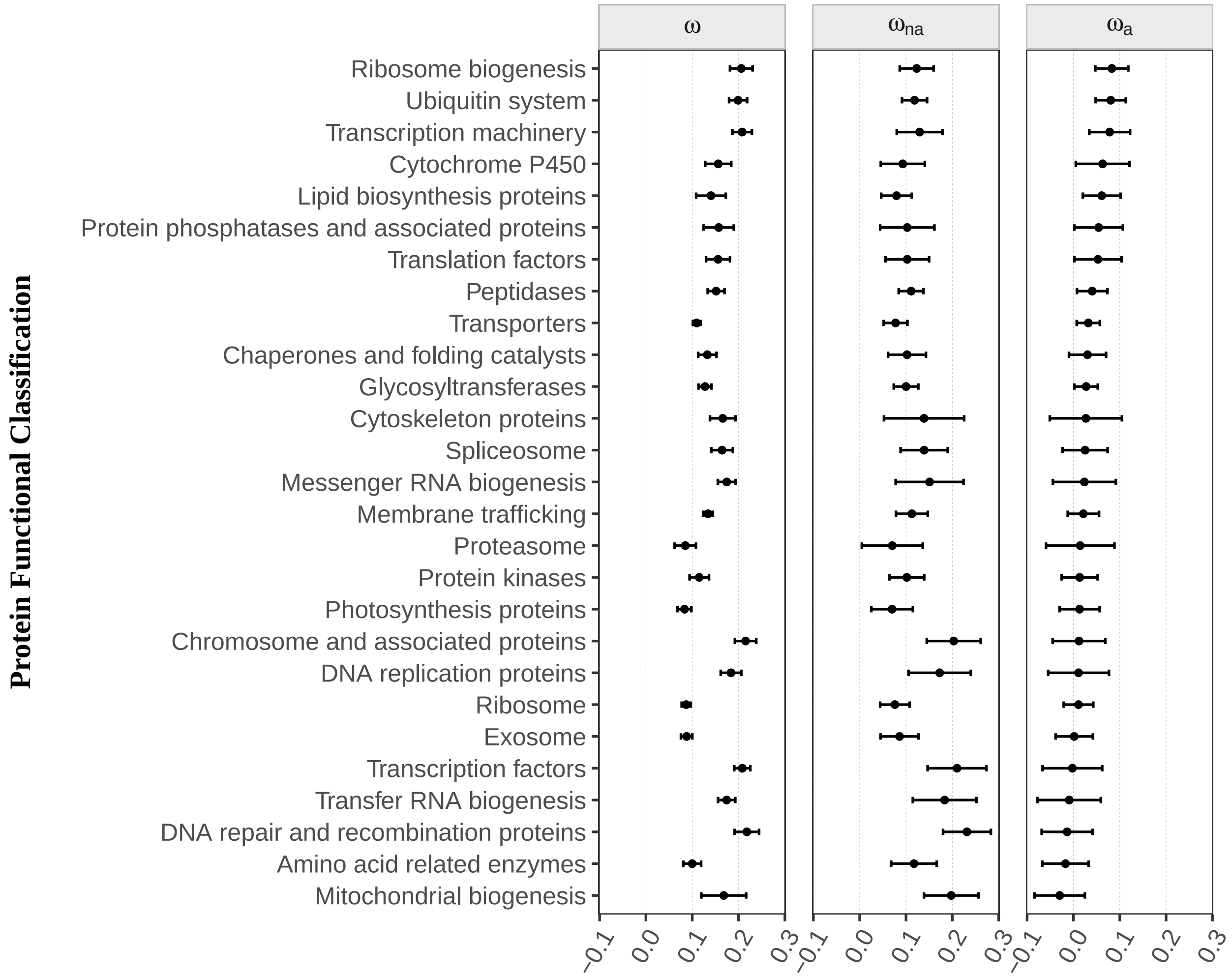


```r
###
### Drosophila
###

stat.dmel <- subset(tbl.stats, tbl.stats$species == "Drosophila")

# to order the plot according to values of omegaA
omegaA.dmel <- subset(stat.dmel, stat.dmel$variable == "omega[a]")
omegaA.dmel <- omegaA.dmel[order(omegaA.dmel[,4]),]

plot.dmel <- ggplot(stat.dmel, aes(x = var.value, y = value.mean)) +
  geom_point(size=1, col = "black") +
  geom_errorbar(aes(ymin=value.mean + 1.96*value.sd,
                    ymax=value.mean - 1.96*value.sd), width = .2) +
  scale_x_discrete(limits = as.character(omegaA.dmel$var.value)) +
  ylab("") +
```

```r
  xlab("Protein Functional Classification") +
  scale_y_continuous(limits = c(-0.1, 0.30)) +
  facet_grid(~variable, labeller = label_parsed) +
  theme_bw() +
  theme.plot() +
  coord_flip()

plot.dmel
```

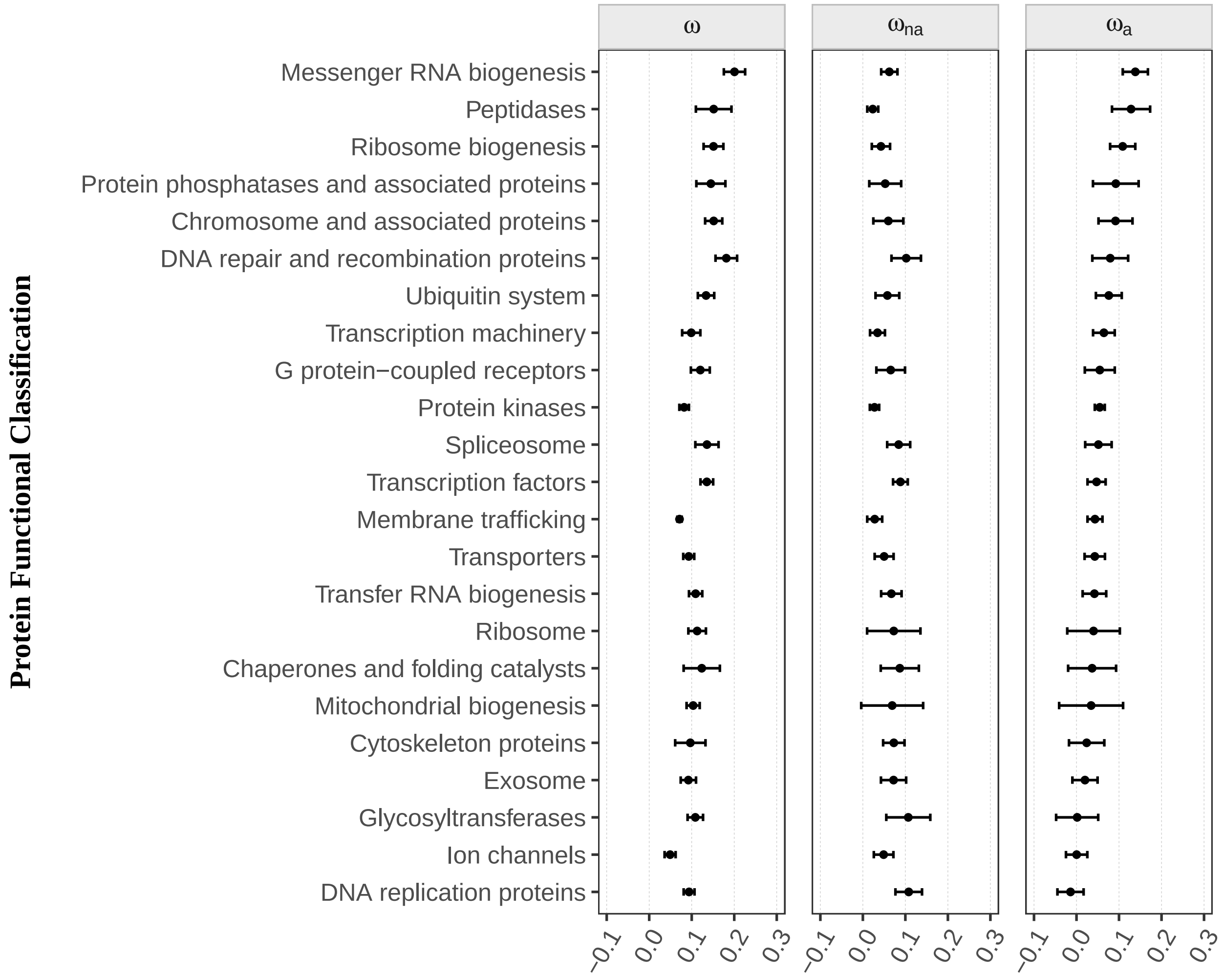


The next chunck of the script describes the statistical analysis.

```r
# libraries
library(rlist)
library(stringr)
library(rowr)
#

# checking the number of replicates per var.value, variable and
# species after removing 1% outliers
kegg.nrep <- ddply(tbl.rates, c("species", "variable", "var.value"),
                   function(x) {
    nrep <- nrow(x)
```

```r
    data.frame(nrep)
})

# take the minimum number of replicates between categories
min.nrep <- ddply(kegg.nrep, c("species", "variable"), function(x) {
  min.n <- min(x$nrep)
  data.frame(min.n)
})
min.nrep
```

| species | variable | min.n |
|---|---|---|
| Arabidopsis | dnds | 28 |
| Arabidopsis | omegaNA | 28 |
| Arabidopsis | omegaA | 28 |
| Drosophila | dnds | 53 |
| Drosophila | omegaNA | 53 |
| Drosophila | omegaA | 53 |

```r
# 28 replicates for Arabidopsis
nrep.arab <- subset(tbl.rates, tbl.rates$species == "Arabidopsis")
tbl.arab <- ddply(nrep.arab, c("species", "var.value", "variable"),
                  function(x) {
  x[sample(nrow(x), 28), ]
})

# 53 replicates for Drosophila
nrep.dmel <- subset(tbl.rates, tbl.rates$species == "Drosophila")
tbl.dmel <- ddply(nrep.dmel, c("species", "var.value", "variable"),
                  function(x) {
  x[sample(nrow(x), 53), ]
})

dat.kegg <- rbind(tbl.arab, tbl.dmel)

# function to split the tables by the name of each variable
kegg.split <- by(dat.kegg, dat.kegg[,c("var.value")], function(y) y)

# to change the column name "value" to the respective id of the category
kegg.value <- lapply(kegg.split, function(x) {
  tbl <- data.frame(x)
  colnames(tbl)[4] <- as.character(unique(tbl$var.value))
  tbl <- tbl[,-2]
  return(tbl)
})

# checking which category has the maximum rows to combine all variables
row.n <- lapply(kegg.value, function(x) {
  nrow(x)
})

species <- kegg.value[[2]][,1]
estimate <- kegg.value[[2]][,2]

tbl.estimate <- lapply(kegg.value, function(x) {
```

```r
  val <- x[,3]
})

# binding all values as columns
tbl.kegg <- list.cbind(tbl.estimate)
tbl.kegg <- as.data.frame(tbl.kegg)
tbl.kegg <- data.frame(tbl.kegg, species, estimate, fix.empty.names = TRUE)
names(tbl.kegg) <- gsub(x = names(tbl.kegg), pattern = "\\.", replacement = "_")

# estimate the differences between columns
# to do so will duplicate the data.frame in order to subtract each column
tbl2.kegg <- tbl.kegg[,1:29]

# doing the differences in a way that we count only for one of the differences
kegg.dif <- cbind(tbl.kegg[, c(30,31), drop=F],
        do.call(cbind, lapply(tbl2.kegg[,2:29],
                              function(x) tbl.kegg[,1:2]-x)),
        do.call(cbind, lapply(tbl2.kegg[,4:29],
                              function(x) tbl.kegg[,3:4]-x)),
        do.call(cbind, lapply(tbl2.kegg[,6:29],
                              function(x) tbl.kegg[,5:6]-x)),
        do.call(cbind, lapply(tbl2.kegg[,8:29],
                              function(x) tbl.kegg[,7:8]-x)),
        do.call(cbind, lapply(tbl2.kegg[,10:29],
                              function(x) tbl.kegg[,9:10]-x)),
        do.call(cbind, lapply(tbl2.kegg[,12:29],
                              function(x) tbl.kegg[,11:12]-x)),
        do.call(cbind, lapply(tbl2.kegg[,14:29],
                              function(x) tbl.kegg[,13:14]-x)),
        do.call(cbind, lapply(tbl2.kegg[,16:29],
                              function(x) tbl.kegg[,15:16]-x)),
        do.call(cbind, lapply(tbl2.kegg[,18:29],
                              function(x) tbl.kegg[,17:18]-x)),
        do.call(cbind, lapply(tbl2.kegg[,20:29],
                              function(x) tbl.kegg[,19:20]-x)),
        do.call(cbind, lapply(tbl2.kegg[,22:29],
                              function(x) tbl.kegg[,21:22]-x)),
        do.call(cbind, lapply(tbl2.kegg[,24:29],
                              function(x) tbl.kegg[,23:24]-x)),
        do.call(cbind, lapply(tbl2.kegg[,26:29],
                              function(x) tbl.kegg[,25:26]-x)),
        do.call(cbind, lapply(tbl2.kegg[,28:29],
                              function(x) tbl.kegg[,27:28]-x)))


# putting all variables in one column
# not counting with the columns comparing the same value
kegg.hist <- melt(kegg.dif, id.vars = c("estimate", "species"),
                  measure.vars = c(names(kegg.dif[3:ncol(kegg.dif)])))

# puting NA values for when the difference is 0 (i.e. when
# comparing the same variables)
split <- str_split_fixed(kegg.hist$variable, "\\.", 2)
```

```r
kegg.hist$var1 <- split[,1]
kegg.hist$var2 <- split[,2]

# removing the variables being compared to itself
kegg.hist$value[kegg.hist$var1 == kegg.hist$var2] <- NA
kegg.hist <- na.omit(kegg.hist[1:6])

## getting the p-value for each difference
## Arabidopsis
arab.hist <- subset(kegg.hist, kegg.hist$species == "Arabidopsis")
nboots <- 100
arab.smallkegg.p <- list()
for (i in 1:nboots) {
  arab.smallkegg.p[[i]] <- ddply(arab.hist, c("species", "estimate",
                                              "var1", "var2"),
                                 function(x, N=28){
    c <- as.numeric(nrow(x[x$value < 0,]))
    c2 <- as.numeric(nrow(x[x$value > 0,]))
    m <- min(c, c2)
    p <- (2*m+1)/(N+1)
    tbl <- data.frame(m, p)
  })
}

# correcting the p-value for multiple testing
arab.kegg.p.adj <- lapply(arab.smallkegg.p, function(x) {
  ddply(x,  c("species","estimate", "var1", "var2"), function(x) {
    p.value <- p.adjust(x$p)
    data.frame(p.value)
  })
})

# taking the minimum p-value of the replicates performed
tbl.arab.p.adj <- rbindlist(arab.kegg.p.adj)
arab.pvalue <- ddply(tbl.arab.p.adj, c("species","estimate", "var1", "var2"),
                     function(x) {
  p.value <- min(x$p.value)
  data.frame(p.value)
})
# showing the table
kable(arab.pvalue, format = "latex", booktabs = TRUE, longtable = TRUE) %>%
  kable_styling(latex_options = c("hold_position", "repeat_header"),
              font_size = 7)
```

| species | estimate | var1 | var2 | p.value |
|---|---|---|---|---|
| Arabidopsis | dnds | Chaperones_and_folding_catalysts | Amino_acid_related_enzymes | 0.1034483 |
| Arabidopsis | dnds | Chromosome_and_associated_proteins | Amino_acid_related_enzymes | 0.0344828 |
| Arabidopsis | dnds | Chromosome_and_associated_proteins | Chaperones_and_folding_catalysts | 0.0344828 |
| Arabidopsis | dnds | Cytochrome_P450 | Amino_acid_related_enzymes | 0.0344828 |
| Arabidopsis | dnds | Cytochrome_P450 | Chaperones_and_folding_catalysts | 0.2413793 |
| Arabidopsis | dnds | Cytochrome_P450 | Chromosome_and_associated_proteins | 0.0344828 |
| Arabidopsis | dnds | Cytoskeleton_proteins | Amino_acid_related_enzymes | 0.0344828 |
| Arabidopsis | dnds | Cytoskeleton_proteins | Chaperones_and_folding_catalysts | 0.0344828 |
| Arabidopsis | dnds | Cytoskeleton_proteins | Chromosome_and_associated_proteins | 0.0344828 |
| Arabidopsis | dnds | Cytoskeleton_proteins | Cytochrome_P450 | 0.5172414 |

*(continued)*

| species | estimate | var1 | var2 | p.value |
|---|---|---|---|---|
| Arabidopsis | dnds | DNA__repair__and__recombination__proteins | Amino__acid__related__enzymes | 0.0344828 |
| Arabidopsis | dnds | DNA__repair__and__recombination__proteins | Chaperones__and__folding__catalysts | 0.0344828 |
| Arabidopsis | dnds | DNA__repair__and__recombination__proteins | Chromosome__and__associated__proteins | 0.9310345 |
| Arabidopsis | dnds | DNA__repair__and__recombination__proteins | Cytochrome__P450 | 0.0344828 |
| Arabidopsis | dnds | DNA__repair__and__recombination__proteins | Cytoskeleton__proteins | 0.0344828 |
| Arabidopsis | dnds | DNA__replication__proteins | Amino__acid__related__enzymes | 0.0344828 |
| Arabidopsis | dnds | DNA__replication__proteins | Chaperones__and__folding__catalysts | 0.0344828 |
| Arabidopsis | dnds | DNA__replication__proteins | Chromosome__and__associated__proteins | 0.0344828 |
| Arabidopsis | dnds | DNA__replication__proteins | Cytochrome__P450 | 0.3103448 |
| Arabidopsis | dnds | DNA__replication__proteins | Cytoskeleton__proteins | 0.3103448 |
| Arabidopsis | dnds | DNA__replication__proteins | DNA__repair__and__recombination__proteins | 0.1034483 |
| Arabidopsis | dnds | Exosome | Amino__acid__related__enzymes | 0.1724138 |
| Arabidopsis | dnds | Exosome | Chaperones__and__folding__catalysts | 0.0344828 |
| Arabidopsis | dnds | Exosome | Chromosome__and__associated__proteins | 0.0344828 |
| Arabidopsis | dnds | Exosome | Cytochrome__P450 | 0.0344828 |
| Arabidopsis | dnds | Exosome | Cytoskeleton__proteins | 0.0344828 |
| Arabidopsis | dnds | Exosome | DNA__repair__and__recombination__proteins | 0.0344828 |
| Arabidopsis | dnds | Exosome | DNA__replication__proteins | 0.0344828 |
| Arabidopsis | dnds | G__protein__coupled__receptors | Amino__acid__related__enzymes | 0.3103448 |
| Arabidopsis | dnds | G__protein__coupled__receptors | Chaperones__and__folding__catalysts | 0.3793103 |
| Arabidopsis | dnds | G__protein__coupled__receptors | Chromosome__and__associated__proteins | 0.0344828 |
| Arabidopsis | dnds | G__protein__coupled__receptors | Cytochrome__P450 | 0.1034483 |
| Arabidopsis | dnds | G__protein__coupled__receptors | Cytoskeleton__proteins | 0.0344828 |
| Arabidopsis | dnds | G__protein__coupled__receptors | DNA__repair__and__recombination__proteins | 0.0344828 |
| Arabidopsis | dnds | G__protein__coupled__receptors | DNA__replication__proteins | 0.0344828 |
| Arabidopsis | dnds | G__protein__coupled__receptors | Exosome | 0.0344828 |
| Arabidopsis | dnds | Glycosyltransferases | Amino__acid__related__enzymes | 0.1034483 |
| Arabidopsis | dnds | Glycosyltransferases | Chaperones__and__folding__catalysts | 0.7931034 |
| Arabidopsis | dnds | Glycosyltransferases | Chromosome__and__associated__proteins | 0.0344828 |
| Arabidopsis | dnds | Glycosyltransferases | Cytochrome__P450 | 0.0344828 |
| Arabidopsis | dnds | Glycosyltransferases | Cytoskeleton__proteins | 0.0344828 |
| Arabidopsis | dnds | Glycosyltransferases | DNA__repair__and__recombination__proteins | 0.0344828 |
| Arabidopsis | dnds | Glycosyltransferases | DNA__replication__proteins | 0.0344828 |
| Arabidopsis | dnds | Glycosyltransferases | Exosome | 0.0344828 |
| Arabidopsis | dnds | Glycosyltransferases | G__protein__coupled__receptors | 0.3793103 |
| Arabidopsis | dnds | Ion__channels | Amino__acid__related__enzymes | 0.0344828 |
| Arabidopsis | dnds | Ion__channels | Chaperones__and__folding__catalysts | 0.0344828 |
| Arabidopsis | dnds | Ion__channels | Chromosome__and__associated__proteins | 0.0344828 |
| Arabidopsis | dnds | Ion__channels | Cytochrome__P450 | 0.0344828 |
| Arabidopsis | dnds | Ion__channels | Cytoskeleton__proteins | 0.0344828 |
| Arabidopsis | dnds | Ion__channels | DNA__repair__and__recombination__proteins | 0.0344828 |
| Arabidopsis | dnds | Ion__channels | DNA__replication__proteins | 0.0344828 |
| Arabidopsis | dnds | Ion__channels | Exosome | 0.0344828 |
| Arabidopsis | dnds | Ion__channels | G__protein__coupled__receptors | 0.0344828 |
| Arabidopsis | dnds | Ion__channels | Glycosyltransferases | 0.0344828 |
| Arabidopsis | dnds | Lipid__biosynthesis__proteins | Amino__acid__related__enzymes | 0.0344828 |
| Arabidopsis | dnds | Lipid__biosynthesis__proteins | Chaperones__and__folding__catalysts | 0.8620690 |
| Arabidopsis | dnds | Lipid__biosynthesis__proteins | Chromosome__and__associated__proteins | 0.0344828 |
| Arabidopsis | dnds | Lipid__biosynthesis__proteins | Cytochrome__P450 | 0.4482759 |
| Arabidopsis | dnds | Lipid__biosynthesis__proteins | Cytoskeleton__proteins | 0.3793103 |
| Arabidopsis | dnds | Lipid__biosynthesis__proteins | DNA__repair__and__recombination__proteins | 0.0344828 |
| Arabidopsis | dnds | Lipid__biosynthesis__proteins | DNA__replication__proteins | 0.1034483 |
| Arabidopsis | dnds | Lipid__biosynthesis__proteins | Exosome | 0.0344828 |
| Arabidopsis | dnds | Lipid__biosynthesis__proteins | G__protein__coupled__receptors | 0.3793103 |
| Arabidopsis | dnds | Lipid__biosynthesis__proteins | Glycosyltransferases | 0.6551724 |
| Arabidopsis | dnds | Lipid__biosynthesis__proteins | Ion__channels | 0.0344828 |
| Arabidopsis | dnds | Membrane__trafficking | Amino__acid__related__enzymes | 0.1034483 |
| Arabidopsis | dnds | Membrane__trafficking | Chaperones__and__folding__catalysts | 1.0000000 |
| Arabidopsis | dnds | Membrane__trafficking | Chromosome__and__associated__proteins | 0.0344828 |
| Arabidopsis | dnds | Membrane__trafficking | Cytochrome__P450 | 0.1034483 |
| Arabidopsis | dnds | Membrane__trafficking | Cytoskeleton__proteins | 0.0344828 |

*(continued)*

| species | estimate | var1 | var2 | p.value |
|---|---|---|---|---|
| Arabidopsis | dnds | Membrane__trafficking | DNA__repair__and__recombination__proteins | 0.0344828 |
| Arabidopsis | dnds | Membrane__trafficking | DNA__replication__proteins | 0.0344828 |
| Arabidopsis | dnds | Membrane__trafficking | Exosome | 0.0344828 |
| Arabidopsis | dnds | Membrane__trafficking | G__protein__coupled__receptors | 0.3103448 |
| Arabidopsis | dnds | Membrane__trafficking | Glycosyltransferases | 0.5172414 |
| Arabidopsis | dnds | Membrane__trafficking | Ion__channels | 0.0344828 |
| Arabidopsis | dnds | Membrane__trafficking | Lipid__biosynthesis__proteins | 1.0000000 |
| Arabidopsis | dnds | Messenger__RNA__biogenesis | Amino__acid__related__enzymes | 0.0344828 |
| Arabidopsis | dnds | Messenger__RNA__biogenesis | Chaperones__and__folding__catalysts | 0.0344828 |
| Arabidopsis | dnds | Messenger__RNA__biogenesis | Chromosome__and__associated__proteins | 0.0344828 |
| Arabidopsis | dnds | Messenger__RNA__biogenesis | Cytochrome__P450 | 0.3793103 |
| Arabidopsis | dnds | Messenger__RNA__biogenesis | Cytoskeleton__proteins | 0.6551724 |
| Arabidopsis | dnds | Messenger__RNA__biogenesis | DNA__repair__and__recombination__proteins | 0.0344828 |
| Arabidopsis | dnds | Messenger__RNA__biogenesis | DNA__replication__proteins | 0.4482759 |
| Arabidopsis | dnds | Messenger__RNA__biogenesis | Exosome | 0.0344828 |
| Arabidopsis | dnds | Messenger__RNA__biogenesis | G__protein__coupled__receptors | 0.0344828 |
| Arabidopsis | dnds | Messenger__RNA__biogenesis | Glycosyltransferases | 0.0344828 |
| Arabidopsis | dnds | Messenger__RNA__biogenesis | Ion__channels | 0.0344828 |
| Arabidopsis | dnds | Messenger__RNA__biogenesis | Lipid__biosynthesis__proteins | 0.1724138 |
| Arabidopsis | dnds | Messenger__RNA__biogenesis | Membrane__trafficking | 0.0344828 |
| Arabidopsis | dnds | Mitochondrial__biogenesis | Amino__acid__related__enzymes | 0.0344828 |
| Arabidopsis | dnds | Mitochondrial__biogenesis | Chaperones__and__folding__catalysts | 0.3793103 |
| Arabidopsis | dnds | Mitochondrial__biogenesis | Chromosome__and__associated__proteins | 0.1034483 |
| Arabidopsis | dnds | Mitochondrial__biogenesis | Cytochrome__P450 | 0.8620690 |
| Arabidopsis | dnds | Mitochondrial__biogenesis | Cytoskeleton__proteins | 0.7931034 |
| Arabidopsis | dnds | Mitochondrial__biogenesis | DNA__repair__and__recombination__proteins | 0.1034483 |
| Arabidopsis | dnds | Mitochondrial__biogenesis | DNA__replication__proteins | 0.5862069 |
| Arabidopsis | dnds | Mitochondrial__biogenesis | Exosome | 0.0344828 |
| Arabidopsis | dnds | Mitochondrial__biogenesis | G__protein__coupled__receptors | 0.1034483 |
| Arabidopsis | dnds | Mitochondrial__biogenesis | Glycosyltransferases | 0.2413793 |
| Arabidopsis | dnds | Mitochondrial__biogenesis | Ion__channels | 0.0344828 |
| Arabidopsis | dnds | Mitochondrial__biogenesis | Lipid__biosynthesis__proteins | 0.3793103 |
| Arabidopsis | dnds | Mitochondrial__biogenesis | Membrane__trafficking | 0.2413793 |
| Arabidopsis | dnds | Mitochondrial__biogenesis | Messenger__RNA__biogenesis | 0.7241379 |
| Arabidopsis | dnds | Peptidases | Amino__acid__related__enzymes | 0.0344828 |
| Arabidopsis | dnds | Peptidases | Chaperones__and__folding__catalysts | 0.1724138 |
| Arabidopsis | dnds | Peptidases | Chromosome__and__associated__proteins | 0.0344828 |
| Arabidopsis | dnds | Peptidases | Cytochrome__P450 | 0.6551724 |
| Arabidopsis | dnds | Peptidases | Cytoskeleton__proteins | 0.4482759 |
| Arabidopsis | dnds | Peptidases | DNA__repair__and__recombination__proteins | 0.0344828 |
| Arabidopsis | dnds | Peptidases | DNA__replication__proteins | 0.0344828 |
| Arabidopsis | dnds | Peptidases | Exosome | 0.0344828 |
| Arabidopsis | dnds | Peptidases | G__protein__coupled__receptors | 0.1034483 |
| Arabidopsis | dnds | Peptidases | Glycosyltransferases | 0.1034483 |
| Arabidopsis | dnds | Peptidases | Ion__channels | 0.0344828 |
| Arabidopsis | dnds | Peptidases | Lipid__biosynthesis__proteins | 0.5172414 |
| Arabidopsis | dnds | Peptidases | Membrane__trafficking | 0.1724138 |
| Arabidopsis | dnds | Peptidases | Messenger__RNA__biogenesis | 0.1724138 |
| Arabidopsis | dnds | Peptidases | Mitochondrial__biogenesis | 0.8620690 |
| Arabidopsis | dnds | Photosynthesis__proteins | Amino__acid__related__enzymes | 0.2413793 |
| Arabidopsis | dnds | Photosynthesis__proteins | Chaperones__and__folding__catalysts | 0.0344828 |
| Arabidopsis | dnds | Photosynthesis__proteins | Chromosome__and__associated__proteins | 0.0344828 |
| Arabidopsis | dnds | Photosynthesis__proteins | Cytochrome__P450 | 0.0344828 |
| Arabidopsis | dnds | Photosynthesis__proteins | Cytoskeleton__proteins | 0.0344828 |
| Arabidopsis | dnds | Photosynthesis__proteins | DNA__repair__and__recombination__proteins | 0.0344828 |
| Arabidopsis | dnds | Photosynthesis__proteins | DNA__replication__proteins | 0.0344828 |
| Arabidopsis | dnds | Photosynthesis__proteins | Exosome | 0.9310345 |
| Arabidopsis | dnds | Photosynthesis__proteins | G__protein__coupled__receptors | 0.0344828 |
| Arabidopsis | dnds | Photosynthesis__proteins | Glycosyltransferases | 0.0344828 |
| Arabidopsis | dnds | Photosynthesis__proteins | Ion__channels | 0.0344828 |
| Arabidopsis | dnds | Photosynthesis__proteins | Lipid__biosynthesis__proteins | 0.0344828 |
| Arabidopsis | dnds | Photosynthesis__proteins | Membrane__trafficking | 0.0344828 |

*(continued)*

| species | estimate | var1 | var2 | p.value |
|---|---|---|---|---|
| Arabidopsis | dnds | Photosynthesis__proteins | Messenger__RNA__biogenesis | 0.0344828 |
| Arabidopsis | dnds | Photosynthesis__proteins | Mitochondrial__biogenesis | 0.0344828 |
| Arabidopsis | dnds | Photosynthesis__proteins | Peptidases | 0.0344828 |
| Arabidopsis | dnds | Proteasome | Amino__acid__related__enzymes | 0.3793103 |
| Arabidopsis | dnds | Proteasome | Chaperones__and__folding__catalysts | 0.0344828 |
| Arabidopsis | dnds | Proteasome | Chromosome__and__associated__proteins | 0.0344828 |
| Arabidopsis | dnds | Proteasome | Cytochrome__P450 | 0.0344828 |
| Arabidopsis | dnds | Proteasome | Cytoskeleton__proteins | 0.0344828 |
| Arabidopsis | dnds | Proteasome | DNA__repair__and__recombination__proteins | 0.0344828 |
| Arabidopsis | dnds | Proteasome | DNA__replication__proteins | 0.0344828 |
| Arabidopsis | dnds | Proteasome | Exosome | 0.7931034 |
| Arabidopsis | dnds | Proteasome | G__protein__coupled__receptors | 0.0344828 |
| Arabidopsis | dnds | Proteasome | Glycosyltransferases | 0.0344828 |
| Arabidopsis | dnds | Proteasome | Ion__channels | 0.0344828 |
| Arabidopsis | dnds | Proteasome | Lipid__biosynthesis__proteins | 0.0344828 |
| Arabidopsis | dnds | Proteasome | Membrane__trafficking | 0.0344828 |
| Arabidopsis | dnds | Proteasome | Messenger__RNA__biogenesis | 0.0344828 |
| Arabidopsis | dnds | Proteasome | Mitochondrial__biogenesis | 0.0344828 |
| Arabidopsis | dnds | Proteasome | Peptidases | 0.0344828 |
| Arabidopsis | dnds | Proteasome | Photosynthesis__proteins | 0.7931034 |
| Arabidopsis | dnds | Protein__kinases | Amino__acid__related__enzymes | 0.3793103 |
| Arabidopsis | dnds | Protein__kinases | Chaperones__and__folding__catalysts | 0.3103448 |
| Arabidopsis | dnds | Protein__kinases | Chromosome__and__associated__proteins | 0.0344828 |
| Arabidopsis | dnds | Protein__kinases | Cytochrome__P450 | 0.0344828 |
| Arabidopsis | dnds | Protein__kinases | Cytoskeleton__proteins | 0.0344828 |
| Arabidopsis | dnds | Protein__kinases | DNA__repair__and__recombination__proteins | 0.0344828 |
| Arabidopsis | dnds | Protein__kinases | DNA__replication__proteins | 0.0344828 |
| Arabidopsis | dnds | Protein__kinases | Exosome | 0.0344828 |
| Arabidopsis | dnds | Protein__kinases | G__protein__coupled__receptors | 0.9310345 |
| Arabidopsis | dnds | Protein__kinases | Glycosyltransferases | 0.3103448 |
| Arabidopsis | dnds | Protein__kinases | Ion__channels | 0.0344828 |
| Arabidopsis | dnds | Protein__kinases | Lipid__biosynthesis__proteins | 0.2413793 |
| Arabidopsis | dnds | Protein__kinases | Membrane__trafficking | 0.2413793 |
| Arabidopsis | dnds | Protein__kinases | Messenger__RNA__biogenesis | 0.0344828 |
| Arabidopsis | dnds | Protein__kinases | Mitochondrial__biogenesis | 0.0344828 |
| Arabidopsis | dnds | Protein__kinases | Peptidases | 0.0344828 |
| Arabidopsis | dnds | Protein__kinases | Photosynthesis__proteins | 0.1034483 |
| Arabidopsis | dnds | Protein__kinases | Proteasome | 0.1034483 |
| Arabidopsis | dnds | Protein__phosphatases__and__associated__proteins | Amino__acid__related__enzymes | 0.0344828 |
| Arabidopsis | dnds | Protein__phosphatases__and__associated__proteins | Chaperones__and__folding__catalysts | 0.2413793 |
| Arabidopsis | dnds | Protein__phosphatases__and__associated__proteins | Chromosome__and__associated__proteins | 0.0344828 |
| Arabidopsis | dnds | Protein__phosphatases__and__associated__proteins | Cytochrome__P450 | 0.8620690 |
| Arabidopsis | dnds | Protein__phosphatases__and__associated__proteins | Cytoskeleton__proteins | 0.6551724 |
| Arabidopsis | dnds | Protein__phosphatases__and__associated__proteins | DNA__repair__and__recombination__proteins | 0.0344828 |
| Arabidopsis | dnds | Protein__phosphatases__and__associated__proteins | DNA__replication__proteins | 0.3103448 |
| Arabidopsis | dnds | Protein__phosphatases__and__associated__proteins | Exosome | 0.0344828 |
| Arabidopsis | dnds | Protein__phosphatases__and__associated__proteins | G__protein__coupled__receptors | 0.0344828 |
| Arabidopsis | dnds | Protein__phosphatases__and__associated__proteins | Glycosyltransferases | 0.2413793 |
| Arabidopsis | dnds | Protein__phosphatases__and__associated__proteins | Ion__channels | 0.0344828 |
| Arabidopsis | dnds | Protein__phosphatases__and__associated__proteins | Lipid__biosynthesis__proteins | 0.4482759 |
| Arabidopsis | dnds | Protein__phosphatases__and__associated__proteins | Membrane__trafficking | 0.1724138 |
| Arabidopsis | dnds | Protein__phosphatases__and__associated__proteins | Messenger__RNA__biogenesis | 0.5862069 |
| Arabidopsis | dnds | Protein__phosphatases__and__associated__proteins | Mitochondrial__biogenesis | 0.7931034 |
| Arabidopsis | dnds | Protein__phosphatases__and__associated__proteins | Peptidases | 0.8620690 |
| Arabidopsis | dnds | Protein__phosphatases__and__associated__proteins | Photosynthesis__proteins | 0.0344828 |
| Arabidopsis | dnds | Protein__phosphatases__and__associated__proteins | Proteasome | 0.0344828 |
| Arabidopsis | dnds | Protein__phosphatases__and__associated__proteins | Protein__kinases | 0.0344828 |
| Arabidopsis | dnds | Ribosome | Amino__acid__related__enzymes | 0.3103448 |
| Arabidopsis | dnds | Ribosome | Chaperones__and__folding__catalysts | 0.0344828 |
| Arabidopsis | dnds | Ribosome | Chromosome__and__associated__proteins | 0.0344828 |
| Arabidopsis | dnds | Ribosome | Cytochrome__P450 | 0.0344828 |
| Arabidopsis | dnds | Ribosome | Cytoskeleton__proteins | 0.0344828 |

*(continued)*

| species | estimate | var1 | var2 | p.value |
|---|---|---|---|---|
| Arabidopsis | dnds | Ribosome | DNA__repair__and__recombination__proteins | 0.0344828 |
| Arabidopsis | dnds | Ribosome | DNA__replication__proteins | 0.0344828 |
| Arabidopsis | dnds | Ribosome | Exosome | 0.9310345 |
| Arabidopsis | dnds | Ribosome | G__protein__coupled__receptors | 0.0344828 |
| Arabidopsis | dnds | Ribosome | Glycosyltransferases | 0.0344828 |
| Arabidopsis | dnds | Ribosome | Ion__channels | 0.0344828 |
| Arabidopsis | dnds | Ribosome | Lipid__biosynthesis__proteins | 0.0344828 |
| Arabidopsis | dnds | Ribosome | Membrane__trafficking | 0.0344828 |
| Arabidopsis | dnds | Ribosome | Messenger__RNA__biogenesis | 0.0344828 |
| Arabidopsis | dnds | Ribosome | Mitochondrial__biogenesis | 0.0344828 |
| Arabidopsis | dnds | Ribosome | Peptidases | 0.0344828 |
| Arabidopsis | dnds | Ribosome | Photosynthesis__proteins | 0.6551724 |
| Arabidopsis | dnds | Ribosome | Proteasome | 0.9310345 |
| Arabidopsis | dnds | Ribosome | Protein__kinases | 0.0344828 |
| Arabidopsis | dnds | Ribosome | Protein__phosphatases__and__associated__proteins | 0.0344828 |
| Arabidopsis | dnds | Ribosome__biogenesis | Amino__acid__related__enzymes | 0.0344828 |
| Arabidopsis | dnds | Ribosome__biogenesis | Chaperones__and__folding__catalysts | 0.0344828 |
| Arabidopsis | dnds | Ribosome__biogenesis | Chromosome__and__associated__proteins | 0.6551724 |
| Arabidopsis | dnds | Ribosome__biogenesis | Cytochrome__P450 | 0.0344828 |
| Arabidopsis | dnds | Ribosome__biogenesis | Cytoskeleton__proteins | 0.0344828 |
| Arabidopsis | dnds | Ribosome__biogenesis | DNA__repair__and__recombination__proteins | 0.6551724 |
| Arabidopsis | dnds | Ribosome__biogenesis | DNA__replication__proteins | 0.1034483 |
| Arabidopsis | dnds | Ribosome__biogenesis | Exosome | 0.0344828 |
| Arabidopsis | dnds | Ribosome__biogenesis | G__protein__coupled__receptors | 0.0344828 |
| Arabidopsis | dnds | Ribosome__biogenesis | Glycosyltransferases | 0.0344828 |
| Arabidopsis | dnds | Ribosome__biogenesis | Ion__channels | 0.0344828 |
| Arabidopsis | dnds | Ribosome__biogenesis | Lipid__biosynthesis__proteins | 0.0344828 |
| Arabidopsis | dnds | Ribosome__biogenesis | Membrane__trafficking | 0.0344828 |
| Arabidopsis | dnds | Ribosome__biogenesis | Messenger__RNA__biogenesis | 0.0344828 |
| Arabidopsis | dnds | Ribosome__biogenesis | Mitochondrial__biogenesis | 0.2413793 |
| Arabidopsis | dnds | Ribosome__biogenesis | Peptidases | 0.0344828 |
| Arabidopsis | dnds | Ribosome__biogenesis | Photosynthesis__proteins | 0.0344828 |
| Arabidopsis | dnds | Ribosome__biogenesis | Proteasome | 0.0344828 |
| Arabidopsis | dnds | Ribosome__biogenesis | Protein__kinases | 0.0344828 |
| Arabidopsis | dnds | Ribosome__biogenesis | Protein__phosphatases__and__associated__proteins | 0.0344828 |
| Arabidopsis | dnds | Ribosome__biogenesis | Ribosome | 0.0344828 |
| Arabidopsis | dnds | Spliceosome | Amino__acid__related__enzymes | 0.0344828 |
| Arabidopsis | dnds | Spliceosome | Chaperones__and__folding__catalysts | 0.1034483 |
| Arabidopsis | dnds | Spliceosome | Chromosome__and__associated__proteins | 0.0344828 |
| Arabidopsis | dnds | Spliceosome | Cytochrome__P450 | 0.7241379 |
| Arabidopsis | dnds | Spliceosome | Cytoskeleton__proteins | 0.9310345 |
| Arabidopsis | dnds | Spliceosome | DNA__repair__and__recombination__proteins | 0.0344828 |
| Arabidopsis | dnds | Spliceosome | DNA__replication__proteins | 0.2413793 |
| Arabidopsis | dnds | Spliceosome | Exosome | 0.0344828 |
| Arabidopsis | dnds | Spliceosome | G__protein__coupled__receptors | 0.0344828 |
| Arabidopsis | dnds | Spliceosome | Glycosyltransferases | 0.0344828 |
| Arabidopsis | dnds | Spliceosome | Ion__channels | 0.0344828 |
| Arabidopsis | dnds | Spliceosome | Lipid__biosynthesis__proteins | 0.2413793 |
| Arabidopsis | dnds | Spliceosome | Membrane__trafficking | 0.0344828 |
| Arabidopsis | dnds | Spliceosome | Messenger__RNA__biogenesis | 0.5862069 |
| Arabidopsis | dnds | Spliceosome | Mitochondrial__biogenesis | 0.9310345 |
| Arabidopsis | dnds | Spliceosome | Peptidases | 0.3793103 |
| Arabidopsis | dnds | Spliceosome | Photosynthesis__proteins | 0.0344828 |
| Arabidopsis | dnds | Spliceosome | Proteasome | 0.0344828 |
| Arabidopsis | dnds | Spliceosome | Protein__kinases | 0.0344828 |
| Arabidopsis | dnds | Spliceosome | Protein__phosphatases__and__associated__proteins | 0.8620690 |
| Arabidopsis | dnds | Spliceosome | Ribosome | 0.0344828 |
| Arabidopsis | dnds | Spliceosome | Ribosome__biogenesis | 0.0344828 |
| Arabidopsis | dnds | Transcription__factors | Amino__acid__related__enzymes | 0.0344828 |
| Arabidopsis | dnds | Transcription__factors | Chaperones__and__folding__catalysts | 0.0344828 |
| Arabidopsis | dnds | Transcription__factors | Chromosome__and__associated__proteins | 0.7241379 |

*(continued)*

| species | estimate | var1 | var2 | p.value |
|---|---|---|---|---|
| Arabidopsis | dnds | Transcription__factors | Cytochrome__P450 | 0.0344828 |
| Arabidopsis | dnds | Transcription__factors | Cytoskeleton__proteins | 0.0344828 |
| Arabidopsis | dnds | Transcription__factors | DNA__repair__and__recombination__proteins | 0.3793103 |
| Arabidopsis | dnds | Transcription__factors | DNA__replication__proteins | 0.1034483 |
| Arabidopsis | dnds | Transcription__factors | Exosome | 0.0344828 |
| Arabidopsis | dnds | Transcription__factors | G__protein__coupled__receptors | 0.0344828 |
| Arabidopsis | dnds | Transcription__factors | Glycosyltransferases | 0.0344828 |
| Arabidopsis | dnds | Transcription__factors | Ion__channels | 0.0344828 |
| Arabidopsis | dnds | Transcription__factors | Lipid__biosynthesis__proteins | 0.0344828 |
| Arabidopsis | dnds | Transcription__factors | Membrane__trafficking | 0.0344828 |
| Arabidopsis | dnds | Transcription__factors | Messenger__RNA__biogenesis | 0.0344828 |
| Arabidopsis | dnds | Transcription__factors | Mitochondrial__biogenesis | 0.1034483 |
| Arabidopsis | dnds | Transcription__factors | Peptidases | 0.0344828 |
| Arabidopsis | dnds | Transcription__factors | Photosynthesis__proteins | 0.0344828 |
| Arabidopsis | dnds | Transcription__factors | Proteasome | 0.0344828 |
| Arabidopsis | dnds | Transcription__factors | Protein__kinases | 0.0344828 |
| Arabidopsis | dnds | Transcription__factors | Protein__phosphatases__and__associated__proteins | 0.0344828 |
| Arabidopsis | dnds | Transcription__factors | Ribosome | 0.0344828 |
| Arabidopsis | dnds | Transcription__factors | Ribosome__biogenesis | 0.8620690 |
| Arabidopsis | dnds | Transcription__factors | Spliceosome | 0.0344828 |
| Arabidopsis | dnds | Transcription__machinery | Amino__acid__related__enzymes | 0.0344828 |
| Arabidopsis | dnds | Transcription__machinery | Chaperones__and__folding__catalysts | 0.0344828 |
| Arabidopsis | dnds | Transcription__machinery | Chromosome__and__associated__proteins | 0.3103448 |
| Arabidopsis | dnds | Transcription__machinery | Cytochrome__P450 | 0.0344828 |
| Arabidopsis | dnds | Transcription__machinery | Cytoskeleton__proteins | 0.0344828 |
| Arabidopsis | dnds | Transcription__machinery | DNA__repair__and__recombination__proteins | 0.5862069 |
| Arabidopsis | dnds | Transcription__machinery | DNA__replication__proteins | 0.0344828 |
| Arabidopsis | dnds | Transcription__machinery | Exosome | 0.0344828 |
| Arabidopsis | dnds | Transcription__machinery | G__protein__coupled__receptors | 0.0344828 |
| Arabidopsis | dnds | Transcription__machinery | Glycosyltransferases | 0.0344828 |
| Arabidopsis | dnds | Transcription__machinery | Ion__channels | 0.0344828 |
| Arabidopsis | dnds | Transcription__machinery | Lipid__biosynthesis__proteins | 0.0344828 |
| Arabidopsis | dnds | Transcription__machinery | Membrane__trafficking | 0.0344828 |
| Arabidopsis | dnds | Transcription__machinery | Messenger__RNA__biogenesis | 0.0344828 |
| Arabidopsis | dnds | Transcription__machinery | Mitochondrial__biogenesis | 0.1034483 |
| Arabidopsis | dnds | Transcription__machinery | Peptidases | 0.0344828 |
| Arabidopsis | dnds | Transcription__machinery | Photosynthesis__proteins | 0.0344828 |
| Arabidopsis | dnds | Transcription__machinery | Proteasome | 0.0344828 |
| Arabidopsis | dnds | Transcription__machinery | Protein__kinases | 0.0344828 |
| Arabidopsis | dnds | Transcription__machinery | Protein__phosphatases__and__associated__proteins | 0.0344828 |
| Arabidopsis | dnds | Transcription__machinery | Ribosome | 0.0344828 |
| Arabidopsis | dnds | Transcription__machinery | Ribosome__biogenesis | 0.9310345 |
| Arabidopsis | dnds | Transcription__machinery | Spliceosome | 0.0344828 |
| Arabidopsis | dnds | Transcription__machinery | Transcription__factors | 0.8620690 |
| Arabidopsis | dnds | Transfer__RNA__biogenesis | Amino__acid__related__enzymes | 0.0344828 |
| Arabidopsis | dnds | Transfer__RNA__biogenesis | Chaperones__and__folding__catalysts | 0.0344828 |
| Arabidopsis | dnds | Transfer__RNA__biogenesis | Chromosome__and__associated__proteins | 0.0344828 |
| Arabidopsis | dnds | Transfer__RNA__biogenesis | Cytochrome__P450 | 0.3103448 |
| Arabidopsis | dnds | Transfer__RNA__biogenesis | Cytoskeleton__proteins | 0.5172414 |
| Arabidopsis | dnds | Transfer__RNA__biogenesis | DNA__repair__and__recombination__proteins | 0.0344828 |
| Arabidopsis | dnds | Transfer__RNA__biogenesis | DNA__replication__proteins | 0.7931034 |
| Arabidopsis | dnds | Transfer__RNA__biogenesis | Exosome | 0.0344828 |
| Arabidopsis | dnds | Transfer__RNA__biogenesis | G__protein__coupled__receptors | 0.0344828 |
| Arabidopsis | dnds | Transfer__RNA__biogenesis | Glycosyltransferases | 0.0344828 |
| Arabidopsis | dnds | Transfer__RNA__biogenesis | Ion__channels | 0.0344828 |
| Arabidopsis | dnds | Transfer__RNA__biogenesis | Lipid__biosynthesis__proteins | 0.0344828 |
| Arabidopsis | dnds | Transfer__RNA__biogenesis | Membrane__trafficking | 0.0344828 |
| Arabidopsis | dnds | Transfer__RNA__biogenesis | Messenger__RNA__biogenesis | 0.6551724 |
| Arabidopsis | dnds | Transfer__RNA__biogenesis | Mitochondrial__biogenesis | 0.5862069 |
| Arabidopsis | dnds | Transfer__RNA__biogenesis | Peptidases | 0.1034483 |
| Arabidopsis | dnds | Transfer__RNA__biogenesis | Photosynthesis__proteins | 0.0344828 |
| Arabidopsis | dnds | Transfer__RNA__biogenesis | Proteasome | 0.0344828 |

*(continued)*

| species | estimate | var1 | var2 | p.value |
|---|---|---|---|---|
| Arabidopsis | dnds | Transfer__RNA__biogenesis | Protein__kinases | 0.0344828 |
| Arabidopsis | dnds | Transfer__RNA__biogenesis | Protein__phosphatases__and__associated__proteins | 0.3103448 |
| Arabidopsis | dnds | Transfer__RNA__biogenesis | Ribosome | 0.0344828 |
| Arabidopsis | dnds | Transfer__RNA__biogenesis | Ribosome__biogenesis | 0.0344828 |
| Arabidopsis | dnds | Transfer__RNA__biogenesis | Spliceosome | 0.3103448 |
| Arabidopsis | dnds | Transfer__RNA__biogenesis | Transcription__factors | 0.0344828 |
| Arabidopsis | dnds | Transfer__RNA__biogenesis | Transcription__machinery | 0.0344828 |
| Arabidopsis | dnds | Translation__factors | Amino__acid__related__enzymes | 0.0344828 |
| Arabidopsis | dnds | Translation__factors | Chaperones__and__folding__catalysts | 0.3103448 |
| Arabidopsis | dnds | Translation__factors | Chromosome__and__associated__proteins | 0.0344828 |
| Arabidopsis | dnds | Translation__factors | Cytochrome__P450 | 0.7241379 |
| Arabidopsis | dnds | Translation__factors | Cytoskeleton__proteins | 0.7241379 |
| Arabidopsis | dnds | Translation__factors | DNA__repair__and__recombination__proteins | 0.0344828 |
| Arabidopsis | dnds | Translation__factors | DNA__replication__proteins | 0.3103448 |
| Arabidopsis | dnds | Translation__factors | Exosome | 0.0344828 |
| Arabidopsis | dnds | Translation__factors | G__protein__coupled__receptors | 0.1034483 |
| Arabidopsis | dnds | Translation__factors | Glycosyltransferases | 0.0344828 |
| Arabidopsis | dnds | Translation__factors | Ion__channels | 0.0344828 |
| Arabidopsis | dnds | Translation__factors | Lipid__biosynthesis__proteins | 0.3793103 |
| Arabidopsis | dnds | Translation__factors | Membrane__trafficking | 0.1724138 |
| Arabidopsis | dnds | Translation__factors | Messenger__RNA__biogenesis | 0.3103448 |
| Arabidopsis | dnds | Translation__factors | Mitochondrial__biogenesis | 0.7931034 |
| Arabidopsis | dnds | Translation__factors | Peptidases | 0.7931034 |
| Arabidopsis | dnds | Translation__factors | Photosynthesis__proteins | 0.0344828 |
| Arabidopsis | dnds | Translation__factors | Proteasome | 0.0344828 |
| Arabidopsis | dnds | Translation__factors | Protein__kinases | 0.0344828 |
| Arabidopsis | dnds | Translation__factors | Protein__phosphatases__and__associated__proteins | 0.9310345 |
| Arabidopsis | dnds | Translation__factors | Ribosome | 0.0344828 |
| Arabidopsis | dnds | Translation__factors | Ribosome__biogenesis | 0.0344828 |
| Arabidopsis | dnds | Translation__factors | Spliceosome | 0.7241379 |
| Arabidopsis | dnds | Translation__factors | Transcription__factors | 0.0344828 |
| Arabidopsis | dnds | Translation__factors | Transcription__machinery | 0.0344828 |
| Arabidopsis | dnds | Translation__factors | Transfer__RNA__biogenesis | 0.3103448 |
| Arabidopsis | dnds | Transporters | Amino__acid__related__enzymes | 0.2413793 |
| Arabidopsis | dnds | Transporters | Chaperones__and__folding__catalysts | 0.1034483 |
| Arabidopsis | dnds | Transporters | Chromosome__and__associated__proteins | 0.0344828 |
| Arabidopsis | dnds | Transporters | Cytochrome__P450 | 0.0344828 |
| Arabidopsis | dnds | Transporters | Cytoskeleton__proteins | 0.0344828 |
| Arabidopsis | dnds | Transporters | DNA__repair__and__recombination__proteins | 0.0344828 |
| Arabidopsis | dnds | Transporters | DNA__replication__proteins | 0.0344828 |
| Arabidopsis | dnds | Transporters | Exosome | 0.0344828 |
| Arabidopsis | dnds | Transporters | G__protein__coupled__receptors | 0.6551724 |
| Arabidopsis | dnds | Transporters | Glycosyltransferases | 0.0344828 |
| Arabidopsis | dnds | Transporters | Ion__channels | 0.0344828 |
| Arabidopsis | dnds | Transporters | Lipid__biosynthesis__proteins | 0.1724138 |
| Arabidopsis | dnds | Transporters | Membrane__trafficking | 0.0344828 |
| Arabidopsis | dnds | Transporters | Messenger__RNA__biogenesis | 0.0344828 |
| Arabidopsis | dnds | Transporters | Mitochondrial__biogenesis | 0.0344828 |
| Arabidopsis | dnds | Transporters | Peptidases | 0.0344828 |
| Arabidopsis | dnds | Transporters | Photosynthesis__proteins | 0.0344828 |
| Arabidopsis | dnds | Transporters | Proteasome | 0.1034483 |
| Arabidopsis | dnds | Transporters | Protein__kinases | 0.7931034 |
| Arabidopsis | dnds | Transporters | Protein__phosphatases__and__associated__proteins | 0.0344828 |
| Arabidopsis | dnds | Transporters | Ribosome | 0.0344828 |
| Arabidopsis | dnds | Transporters | Ribosome__biogenesis | 0.0344828 |
| Arabidopsis | dnds | Transporters | Spliceosome | 0.0344828 |
| Arabidopsis | dnds | Transporters | Transcription__factors | 0.0344828 |
| Arabidopsis | dnds | Transporters | Transcription__machinery | 0.0344828 |
| Arabidopsis | dnds | Transporters | Transfer__RNA__biogenesis | 0.0344828 |
| Arabidopsis | dnds | Transporters | Translation__factors | 0.0344828 |
| Arabidopsis | dnds | Ubiquitin__system | Amino__acid__related__enzymes | 0.0344828 |
| Arabidopsis | dnds | Ubiquitin__system | Chaperones__and__folding__catalysts | 0.0344828 |

*(continued)*

| species | estimate | var1 | var2 | p.value |
|---|---|---|---|---|
| Arabidopsis | dnds | Ubiquitin__system | Chromosome__and__associated__proteins | 0.2413793 |
| Arabidopsis | dnds | Ubiquitin__system | Cytochrome__P450 | 0.0344828 |
| Arabidopsis | dnds | Ubiquitin__system | Cytoskeleton__proteins | 0.0344828 |
| Arabidopsis | dnds | Ubiquitin__system | DNA__repair__and__recombination__proteins | 0.3793103 |
| Arabidopsis | dnds | Ubiquitin__system | DNA__replication__proteins | 0.2413793 |
| Arabidopsis | dnds | Ubiquitin__system | Exosome | 0.0344828 |
| Arabidopsis | dnds | Ubiquitin__system | G__protein__coupled__receptors | 0.0344828 |
| Arabidopsis | dnds | Ubiquitin__system | Glycosyltransferases | 0.0344828 |
| Arabidopsis | dnds | Ubiquitin__system | Ion__channels | 0.0344828 |
| Arabidopsis | dnds | Ubiquitin__system | Lipid__biosynthesis__proteins | 0.0344828 |
| Arabidopsis | dnds | Ubiquitin__system | Membrane__trafficking | 0.0344828 |
| Arabidopsis | dnds | Ubiquitin__system | Messenger__RNA__biogenesis | 0.1034483 |
| Arabidopsis | dnds | Ubiquitin__system | Mitochondrial__biogenesis | 0.3103448 |
| Arabidopsis | dnds | Ubiquitin__system | Peptidases | 0.0344828 |
| Arabidopsis | dnds | Ubiquitin__system | Photosynthesis__proteins | 0.0344828 |
| Arabidopsis | dnds | Ubiquitin__system | Proteasome | 0.0344828 |
| Arabidopsis | dnds | Ubiquitin__system | Protein__kinases | 0.0344828 |
| Arabidopsis | dnds | Ubiquitin__system | Protein__phosphatases__and__associated__proteins | 0.0344828 |
| Arabidopsis | dnds | Ubiquitin__system | Ribosome | 0.0344828 |
| Arabidopsis | dnds | Ubiquitin__system | Ribosome__biogenesis | 0.7931034 |
| Arabidopsis | dnds | Ubiquitin__system | Spliceosome | 0.1034483 |
| Arabidopsis | dnds | Ubiquitin__system | Transcription__factors | 0.5172414 |
| Arabidopsis | dnds | Ubiquitin__system | Transcription__machinery | 0.5862069 |
| Arabidopsis | dnds | Ubiquitin__system | Transfer__RNA__biogenesis | 0.1034483 |
| Arabidopsis | dnds | Ubiquitin__system | Translation__factors | 0.1034483 |
| Arabidopsis | dnds | Ubiquitin__system | Transporters | 0.0344828 |
| Arabidopsis | omegaNA | Chaperones__and__folding__catalysts | Amino__acid__related__enzymes | 0.5172414 |
| Arabidopsis | omegaNA | Chromosome__and__associated__proteins | Amino__acid__related__enzymes | 0.1034483 |
| Arabidopsis | omegaNA | Chromosome__and__associated__proteins | Chaperones__and__folding__catalysts | 0.0344828 |
| Arabidopsis | omegaNA | Cytochrome__P450 | Amino__acid__related__enzymes | 0.3793103 |
| Arabidopsis | omegaNA | Cytochrome__P450 | Chaperones__and__folding__catalysts | 0.7931034 |
| Arabidopsis | omegaNA | Cytochrome__P450 | Chromosome__and__associated__proteins | 0.0344828 |
| Arabidopsis | omegaNA | Cytoskeleton__proteins | Amino__acid__related__enzymes | 0.7241379 |
| Arabidopsis | omegaNA | Cytoskeleton__proteins | Chaperones__and__folding__catalysts | 0.5172414 |
| Arabidopsis | omegaNA | Cytoskeleton__proteins | Chromosome__and__associated__proteins | 0.2413793 |
| Arabidopsis | omegaNA | Cytoskeleton__proteins | Cytochrome__P450 | 0.5172414 |
| Arabidopsis | omegaNA | DNA__repair__and__recombination__proteins | Amino__acid__related__enzymes | 0.0344828 |
| Arabidopsis | omegaNA | DNA__repair__and__recombination__proteins | Chaperones__and__folding__catalysts | 0.0344828 |
| Arabidopsis | omegaNA | DNA__repair__and__recombination__proteins | Chromosome__and__associated__proteins | 0.4482759 |
| Arabidopsis | omegaNA | DNA__repair__and__recombination__proteins | Cytochrome__P450 | 0.0344828 |
| Arabidopsis | omegaNA | DNA__repair__and__recombination__proteins | Cytoskeleton__proteins | 0.1724138 |
| Arabidopsis | omegaNA | DNA__replication__proteins | Amino__acid__related__enzymes | 0.1034483 |
| Arabidopsis | omegaNA | DNA__replication__proteins | Chaperones__and__folding__catalysts | 0.0344828 |
| Arabidopsis | omegaNA | DNA__replication__proteins | Chromosome__and__associated__proteins | 0.4482759 |
| Arabidopsis | omegaNA | DNA__replication__proteins | Cytochrome__P450 | 0.1034483 |
| Arabidopsis | omegaNA | DNA__replication__proteins | Cytoskeleton__proteins | 0.3793103 |
| Arabidopsis | omegaNA | DNA__replication__proteins | DNA__repair__and__recombination__proteins | 0.1724138 |
| Arabidopsis | omegaNA | Exosome | Amino__acid__related__enzymes | 0.2413793 |
| Arabidopsis | omegaNA | Exosome | Chaperones__and__folding__catalysts | 0.6551724 |
| Arabidopsis | omegaNA | Exosome | Chromosome__and__associated__proteins | 0.0344828 |
| Arabidopsis | omegaNA | Exosome | Cytochrome__P450 | 0.7241379 |
| Arabidopsis | omegaNA | Exosome | Cytoskeleton__proteins | 0.2413793 |
| Arabidopsis | omegaNA | Exosome | DNA__repair__and__recombination__proteins | 0.0344828 |
| Arabidopsis | omegaNA | Exosome | DNA__replication__proteins | 0.1034483 |
| Arabidopsis | omegaNA | G__protein__coupled__receptors | Amino__acid__related__enzymes | 0.9310345 |
| Arabidopsis | omegaNA | G__protein__coupled__receptors | Chaperones__and__folding__catalysts | 0.6551724 |
| Arabidopsis | omegaNA | G__protein__coupled__receptors | Chromosome__and__associated__proteins | 0.0344828 |
| Arabidopsis | omegaNA | G__protein__coupled__receptors | Cytochrome__P450 | 0.3793103 |
| Arabidopsis | omegaNA | G__protein__coupled__receptors | Cytoskeleton__proteins | 1.0000000 |
| Arabidopsis | omegaNA | G__protein__coupled__receptors | DNA__repair__and__recombination__proteins | 0.0344828 |
| Arabidopsis | omegaNA | G__protein__coupled__receptors | DNA__replication__proteins | 0.0344828 |

*(continued)*

| species | estimate | var1 | var2 | p.value |
|---|---|---|---|---|
| Arabidopsis | omegaNA | G__protein__coupled__receptors | Exosome | 0.2413793 |
| Arabidopsis | omegaNA | Glycosyltransferases | Amino__acid__related__enzymes | 0.5172414 |
| Arabidopsis | omegaNA | Glycosyltransferases | Chaperones__and__folding__catalysts | 0.9310345 |
| Arabidopsis | omegaNA | Glycosyltransferases | Chromosome__and__associated__proteins | 0.0344828 |
| Arabidopsis | omegaNA | Glycosyltransferases | Cytochrome__P450 | 0.4482759 |
| Arabidopsis | omegaNA | Glycosyltransferases | Cytoskeleton__proteins | 0.6551724 |
| Arabidopsis | omegaNA | Glycosyltransferases | DNA__repair__and__recombination__proteins | 0.0344828 |
| Arabidopsis | omegaNA | Glycosyltransferases | DNA__replication__proteins | 0.0344828 |
| Arabidopsis | omegaNA | Glycosyltransferases | Exosome | 0.5172414 |
| Arabidopsis | omegaNA | Glycosyltransferases | G__protein__coupled__receptors | 0.3793103 |
| Arabidopsis | omegaNA | Ion__channels | Amino__acid__related__enzymes | 0.0344828 |
| Arabidopsis | omegaNA | Ion__channels | Chaperones__and__folding__catalysts | 0.0344828 |
| Arabidopsis | omegaNA | Ion__channels | Chromosome__and__associated__proteins | 0.0344828 |
| Arabidopsis | omegaNA | Ion__channels | Cytochrome__P450 | 0.0344828 |
| Arabidopsis | omegaNA | Ion__channels | Cytoskeleton__proteins | 0.0344828 |
| Arabidopsis | omegaNA | Ion__channels | DNA__repair__and__recombination__proteins | 0.0344828 |
| Arabidopsis | omegaNA | Ion__channels | DNA__replication__proteins | 0.0344828 |
| Arabidopsis | omegaNA | Ion__channels | Exosome | 0.1034483 |
| Arabidopsis | omegaNA | Ion__channels | G__protein__coupled__receptors | 0.1724138 |
| Arabidopsis | omegaNA | Ion__channels | Glycosyltransferases | 0.0344828 |
| Arabidopsis | omegaNA | Lipid__biosynthesis__proteins | Amino__acid__related__enzymes | 0.3103448 |
| Arabidopsis | omegaNA | Lipid__biosynthesis__proteins | Chaperones__and__folding__catalysts | 0.7241379 |
| Arabidopsis | omegaNA | Lipid__biosynthesis__proteins | Chromosome__and__associated__proteins | 0.0344828 |
| Arabidopsis | omegaNA | Lipid__biosynthesis__proteins | Cytochrome__P450 | 0.7931034 |
| Arabidopsis | omegaNA | Lipid__biosynthesis__proteins | Cytoskeleton__proteins | 0.2413793 |
| Arabidopsis | omegaNA | Lipid__biosynthesis__proteins | DNA__repair__and__recombination__proteins | 0.0344828 |
| Arabidopsis | omegaNA | Lipid__biosynthesis__proteins | DNA__replication__proteins | 0.0344828 |
| Arabidopsis | omegaNA | Lipid__biosynthesis__proteins | Exosome | 0.8620690 |
| Arabidopsis | omegaNA | Lipid__biosynthesis__proteins | G__protein__coupled__receptors | 0.3103448 |
| Arabidopsis | omegaNA | Lipid__biosynthesis__proteins | Glycosyltransferases | 0.4482759 |
| Arabidopsis | omegaNA | Lipid__biosynthesis__proteins | Ion__channels | 0.1724138 |
| Arabidopsis | omegaNA | Membrane__trafficking | Amino__acid__related__enzymes | 0.8620690 |
| Arabidopsis | omegaNA | Membrane__trafficking | Chaperones__and__folding__catalysts | 0.5172414 |
| Arabidopsis | omegaNA | Membrane__trafficking | Chromosome__and__associated__proteins | 0.0344828 |
| Arabidopsis | omegaNA | Membrane__trafficking | Cytochrome__P450 | 0.3103448 |
| Arabidopsis | omegaNA | Membrane__trafficking | Cytoskeleton__proteins | 0.6551724 |
| Arabidopsis | omegaNA | Membrane__trafficking | DNA__repair__and__recombination__proteins | 0.0344828 |
| Arabidopsis | omegaNA | Membrane__trafficking | DNA__replication__proteins | 0.0344828 |
| Arabidopsis | omegaNA | Membrane__trafficking | Exosome | 0.3103448 |
| Arabidopsis | omegaNA | Membrane__trafficking | G__protein__coupled__receptors | 0.7931034 |
| Arabidopsis | omegaNA | Membrane__trafficking | Glycosyltransferases | 0.5862069 |
| Arabidopsis | omegaNA | Membrane__trafficking | Ion__channels | 0.0344828 |
| Arabidopsis | omegaNA | Membrane__trafficking | Lipid__biosynthesis__proteins | 0.2413793 |
| Arabidopsis | omegaNA | Messenger__RNA__biogenesis | Amino__acid__related__enzymes | 0.7241379 |
| Arabidopsis | omegaNA | Messenger__RNA__biogenesis | Chaperones__and__folding__catalysts | 0.2413793 |
| Arabidopsis | omegaNA | Messenger__RNA__biogenesis | Chromosome__and__associated__proteins | 0.3793103 |
| Arabidopsis | omegaNA | Messenger__RNA__biogenesis | Cytochrome__P450 | 0.1724138 |
| Arabidopsis | omegaNA | Messenger__RNA__biogenesis | Cytoskeleton__proteins | 0.6551724 |
| Arabidopsis | omegaNA | Messenger__RNA__biogenesis | DNA__repair__and__recombination__proteins | 0.0344828 |
| Arabidopsis | omegaNA | Messenger__RNA__biogenesis | DNA__replication__proteins | 0.5862069 |
| Arabidopsis | omegaNA | Messenger__RNA__biogenesis | Exosome | 0.1724138 |
| Arabidopsis | omegaNA | Messenger__RNA__biogenesis | G__protein__coupled__receptors | 0.4482759 |
| Arabidopsis | omegaNA | Messenger__RNA__biogenesis | Glycosyltransferases | 0.2413793 |
| Arabidopsis | omegaNA | Messenger__RNA__biogenesis | Ion__channels | 0.0344828 |
| Arabidopsis | omegaNA | Messenger__RNA__biogenesis | Lipid__biosynthesis__proteins | 0.1724138 |
| Arabidopsis | omegaNA | Messenger__RNA__biogenesis | Membrane__trafficking | 0.3793103 |
| Arabidopsis | omegaNA | Mitochondrial__biogenesis | Amino__acid__related__enzymes | 0.1034483 |
| Arabidopsis | omegaNA | Mitochondrial__biogenesis | Chaperones__and__folding__catalysts | 0.0344828 |
| Arabidopsis | omegaNA | Mitochondrial__biogenesis | Chromosome__and__associated__proteins | 0.7931034 |
| Arabidopsis | omegaNA | Mitochondrial__biogenesis | Cytochrome__P450 | 0.0344828 |
| Arabidopsis | omegaNA | Mitochondrial__biogenesis | Cytoskeleton__proteins | 0.1724138 |
| Arabidopsis | omegaNA | Mitochondrial__biogenesis | DNA__repair__and__recombination__proteins | 0.3103448 |

*(continued)*

| species | estimate | var1 | var2 | p.value |
|---|---|---|---|---|
| Arabidopsis | omegaNA | Mitochondrial__biogenesis | DNA__replication__proteins | 0.5172414 |
| Arabidopsis | omegaNA | Mitochondrial__biogenesis | Exosome | 0.0344828 |
| Arabidopsis | omegaNA | Mitochondrial__biogenesis | G__protein__coupled__receptors | 0.0344828 |
| Arabidopsis | omegaNA | Mitochondrial__biogenesis | Glycosyltransferases | 0.0344828 |
| Arabidopsis | omegaNA | Mitochondrial__biogenesis | Ion__channels | 0.0344828 |
| Arabidopsis | omegaNA | Mitochondrial__biogenesis | Lipid__biosynthesis__proteins | 0.0344828 |
| Arabidopsis | omegaNA | Mitochondrial__biogenesis | Membrane__trafficking | 0.0344828 |
| Arabidopsis | omegaNA | Mitochondrial__biogenesis | Messenger__RNA__biogenesis | 0.2413793 |
| Arabidopsis | omegaNA | Peptidases | Amino__acid__related__enzymes | 0.7241379 |
| Arabidopsis | omegaNA | Peptidases | Chaperones__and__folding__catalysts | 0.6551724 |
| Arabidopsis | omegaNA | Peptidases | Chromosome__and__associated__proteins | 0.0344828 |
| Arabidopsis | omegaNA | Peptidases | Cytochrome__P450 | 0.2413793 |
| Arabidopsis | omegaNA | Peptidases | Cytoskeleton__proteins | 0.7931034 |
| Arabidopsis | omegaNA | Peptidases | DNA__repair__and__recombination__proteins | 0.0344828 |
| Arabidopsis | omegaNA | Peptidases | DNA__replication__proteins | 0.0344828 |
| Arabidopsis | omegaNA | Peptidases | Exosome | 0.3103448 |
| Arabidopsis | omegaNA | Peptidases | G__protein__coupled__receptors | 0.7931034 |
| Arabidopsis | omegaNA | Peptidases | Glycosyltransferases | 0.5172414 |
| Arabidopsis | omegaNA | Peptidases | Ion__channels | 0.0344828 |
| Arabidopsis | omegaNA | Peptidases | Lipid__biosynthesis__proteins | 0.1724138 |
| Arabidopsis | omegaNA | Peptidases | Membrane__trafficking | 0.8620690 |
| Arabidopsis | omegaNA | Peptidases | Messenger__RNA__biogenesis | 0.3793103 |
| Arabidopsis | omegaNA | Peptidases | Mitochondrial__biogenesis | 0.0344828 |
| Arabidopsis | omegaNA | Photosynthesis__proteins | Amino__acid__related__enzymes | 0.3103448 |
| Arabidopsis | omegaNA | Photosynthesis__proteins | Chaperones__and__folding__catalysts | 0.5172414 |
| Arabidopsis | omegaNA | Photosynthesis__proteins | Chromosome__and__associated__proteins | 0.0344828 |
| Arabidopsis | omegaNA | Photosynthesis__proteins | Cytochrome__P450 | 0.5862069 |
| Arabidopsis | omegaNA | Photosynthesis__proteins | Cytoskeleton__proteins | 0.3103448 |
| Arabidopsis | omegaNA | Photosynthesis__proteins | DNA__repair__and__recombination__proteins | 0.0344828 |
| Arabidopsis | omegaNA | Photosynthesis__proteins | DNA__replication__proteins | 0.1034483 |
| Arabidopsis | omegaNA | Photosynthesis__proteins | Exosome | 0.7241379 |
| Arabidopsis | omegaNA | Photosynthesis__proteins | G__protein__coupled__receptors | 0.2413793 |
| Arabidopsis | omegaNA | Photosynthesis__proteins | Glycosyltransferases | 0.3103448 |
| Arabidopsis | omegaNA | Photosynthesis__proteins | Ion__channels | 0.3793103 |
| Arabidopsis | omegaNA | Photosynthesis__proteins | Lipid__biosynthesis__proteins | 0.8620690 |
| Arabidopsis | omegaNA | Photosynthesis__proteins | Membrane__trafficking | 0.1034483 |
| Arabidopsis | omegaNA | Photosynthesis__proteins | Messenger__RNA__biogenesis | 0.1724138 |
| Arabidopsis | omegaNA | Photosynthesis__proteins | Mitochondrial__biogenesis | 0.0344828 |
| Arabidopsis | omegaNA | Photosynthesis__proteins | Peptidases | 0.3103448 |
| Arabidopsis | omegaNA | Proteasome | Amino__acid__related__enzymes | 0.2413793 |
| Arabidopsis | omegaNA | Proteasome | Chaperones__and__folding__catalysts | 0.5862069 |
| Arabidopsis | omegaNA | Proteasome | Chromosome__and__associated__proteins | 0.0344828 |
| Arabidopsis | omegaNA | Proteasome | Cytochrome__P450 | 0.7931034 |
| Arabidopsis | omegaNA | Proteasome | Cytoskeleton__proteins | 0.2413793 |
| Arabidopsis | omegaNA | Proteasome | DNA__repair__and__recombination__proteins | 0.0344828 |
| Arabidopsis | omegaNA | Proteasome | DNA__replication__proteins | 0.0344828 |
| Arabidopsis | omegaNA | Proteasome | Exosome | 0.7241379 |
| Arabidopsis | omegaNA | Proteasome | G__protein__coupled__receptors | 0.1724138 |
| Arabidopsis | omegaNA | Proteasome | Glycosyltransferases | 0.2413793 |
| Arabidopsis | omegaNA | Proteasome | Ion__channels | 0.5172414 |
| Arabidopsis | omegaNA | Proteasome | Lipid__biosynthesis__proteins | 0.7241379 |
| Arabidopsis | omegaNA | Proteasome | Membrane__trafficking | 0.1724138 |
| Arabidopsis | omegaNA | Proteasome | Messenger__RNA__biogenesis | 0.0344828 |
| Arabidopsis | omegaNA | Proteasome | Mitochondrial__biogenesis | 0.0344828 |
| Arabidopsis | omegaNA | Proteasome | Peptidases | 0.3793103 |
| Arabidopsis | omegaNA | Proteasome | Photosynthesis__proteins | 0.8620690 |
| Arabidopsis | omegaNA | Protein__kinases | Amino__acid__related__enzymes | 0.5862069 |
| Arabidopsis | omegaNA | Protein__kinases | Chaperones__and__folding__catalysts | 0.9310345 |
| Arabidopsis | omegaNA | Protein__kinases | Chromosome__and__associated__proteins | 0.0344828 |
| Arabidopsis | omegaNA | Protein__kinases | Cytochrome__P450 | 0.7241379 |
| Arabidopsis | omegaNA | Protein__kinases | Cytoskeleton__proteins | 0.6551724 |
| Arabidopsis | omegaNA | Protein__kinases | DNA__repair__and__recombination__proteins | 0.0344828 |

*(continued)*

| species | estimate | var1 | var2 | p.value |
|---|---|---|---|---|
| Arabidopsis | omegaNA | Protein__kinases | DNA__replication__proteins | 0.0344828 |
| Arabidopsis | omegaNA | Protein__kinases | Exosome | 0.5862069 |
| Arabidopsis | omegaNA | Protein__kinases | G__protein__coupled__receptors | 0.7241379 |
| Arabidopsis | omegaNA | Protein__kinases | Glycosyltransferases | 0.9310345 |
| Arabidopsis | omegaNA | Protein__kinases | Ion__channels | 0.0344828 |
| Arabidopsis | omegaNA | Protein__kinases | Lipid__biosynthesis__proteins | 0.5862069 |
| Arabidopsis | omegaNA | Protein__kinases | Membrane__trafficking | 0.5172414 |
| Arabidopsis | omegaNA | Protein__kinases | Messenger__RNA__biogenesis | 0.1724138 |
| Arabidopsis | omegaNA | Protein__kinases | Mitochondrial__biogenesis | 0.0344828 |
| Arabidopsis | omegaNA | Protein__kinases | Peptidases | 0.7241379 |
| Arabidopsis | omegaNA | Protein__kinases | Photosynthesis__proteins | 0.5172414 |
| Arabidopsis | omegaNA | Protein__kinases | Proteasome | 0.3793103 |
| Arabidopsis | omegaNA | Protein__phosphatases__and__associated__proteins | Amino__acid__related__enzymes | 0.7241379 |
| Arabidopsis | omegaNA | Protein__phosphatases__and__associated__proteins | Chaperones__and__folding__catalysts | 1.0000000 |
| Arabidopsis | omegaNA | Protein__phosphatases__and__associated__proteins | Chromosome__and__associated__proteins | 0.1034483 |
| Arabidopsis | omegaNA | Protein__phosphatases__and__associated__proteins | Cytochrome__P450 | 0.5862069 |
| Arabidopsis | omegaNA | Protein__phosphatases__and__associated__proteins | Cytoskeleton__proteins | 0.6551724 |
| Arabidopsis | omegaNA | Protein__phosphatases__and__associated__proteins | DNA__repair__and__recombination__proteins | 0.0344828 |
| Arabidopsis | omegaNA | Protein__phosphatases__and__associated__proteins | DNA__replication__proteins | 0.1034483 |
| Arabidopsis | omegaNA | Protein__phosphatases__and__associated__proteins | Exosome | 0.6551724 |
| Arabidopsis | omegaNA | Protein__phosphatases__and__associated__proteins | G__protein__coupled__receptors | 0.7241379 |
| Arabidopsis | omegaNA | Protein__phosphatases__and__associated__proteins | Glycosyltransferases | 1.0000000 |
| Arabidopsis | omegaNA | Protein__phosphatases__and__associated__proteins | Ion__channels | 0.0344828 |
| Arabidopsis | omegaNA | Protein__phosphatases__and__associated__proteins | Lipid__biosynthesis__proteins | 0.7241379 |
| Arabidopsis | omegaNA | Protein__phosphatases__and__associated__proteins | Membrane__trafficking | 0.7931034 |
| Arabidopsis | omegaNA | Protein__phosphatases__and__associated__proteins | Messenger__RNA__biogenesis | 0.6551724 |
| Arabidopsis | omegaNA | Protein__phosphatases__and__associated__proteins | Mitochondrial__biogenesis | 0.1034483 |
| Arabidopsis | omegaNA | Protein__phosphatases__and__associated__proteins | Peptidases | 0.7931034 |
| Arabidopsis | omegaNA | Protein__phosphatases__and__associated__proteins | Photosynthesis__proteins | 0.4482759 |
| Arabidopsis | omegaNA | Protein__phosphatases__and__associated__proteins | Proteasome | 0.3793103 |
| Arabidopsis | omegaNA | Protein__phosphatases__and__associated__proteins | Protein__kinases | 0.9310345 |
| Arabidopsis | omegaNA | Ribosome | Amino__acid__related__enzymes | 0.1034483 |
| Arabidopsis | omegaNA | Ribosome | Chaperones__and__folding__catalysts | 0.4482759 |
| Arabidopsis | omegaNA | Ribosome | Chromosome__and__associated__proteins | 0.0344828 |
| Arabidopsis | omegaNA | Ribosome | Cytochrome__P450 | 0.5172414 |
| Arabidopsis | omegaNA | Ribosome | Cytoskeleton__proteins | 0.1724138 |
| Arabidopsis | omegaNA | Ribosome | DNA__repair__and__recombination__proteins | 0.0344828 |
| Arabidopsis | omegaNA | Ribosome | DNA__replication__proteins | 0.0344828 |
| Arabidopsis | omegaNA | Ribosome | Exosome | 1.0000000 |
| Arabidopsis | omegaNA | Ribosome | G__protein__coupled__receptors | 0.2413793 |
| Arabidopsis | omegaNA | Ribosome | Glycosyltransferases | 0.3793103 |
| Arabidopsis | omegaNA | Ribosome | Ion__channels | 0.3793103 |
| Arabidopsis | omegaNA | Ribosome | Lipid__biosynthesis__proteins | 0.9310345 |
| Arabidopsis | omegaNA | Ribosome | Membrane__trafficking | 0.0344828 |
| Arabidopsis | omegaNA | Ribosome | Messenger__RNA__biogenesis | 0.1034483 |
| Arabidopsis | omegaNA | Ribosome | Mitochondrial__biogenesis | 0.0344828 |
| Arabidopsis | omegaNA | Ribosome | Peptidases | 0.1034483 |
| Arabidopsis | omegaNA | Ribosome | Photosynthesis__proteins | 0.9310345 |
| Arabidopsis | omegaNA | Ribosome | Proteasome | 0.9310345 |
| Arabidopsis | omegaNA | Ribosome | Protein__kinases | 0.3103448 |
| Arabidopsis | omegaNA | Ribosome | Protein__phosphatases__and__associated__proteins | 0.4482759 |
| Arabidopsis | omegaNA | Ribosome__biogenesis | Amino__acid__related__enzymes | 0.9310345 |
| Arabidopsis | omegaNA | Ribosome__biogenesis | Chaperones__and__folding__catalysts | 0.3793103 |
| Arabidopsis | omegaNA | Ribosome__biogenesis | Chromosome__and__associated__proteins | 0.0344828 |
| Arabidopsis | omegaNA | Ribosome__biogenesis | Cytochrome__P450 | 0.2413793 |
| Arabidopsis | omegaNA | Ribosome__biogenesis | Cytoskeleton__proteins | 0.9310345 |
| Arabidopsis | omegaNA | Ribosome__biogenesis | DNA__repair__and__recombination__proteins | 0.0344828 |
| Arabidopsis | omegaNA | Ribosome__biogenesis | DNA__replication__proteins | 0.1724138 |
| Arabidopsis | omegaNA | Ribosome__biogenesis | Exosome | 0.2413793 |
| Arabidopsis | omegaNA | Ribosome__biogenesis | G__protein__coupled__receptors | 0.8620690 |
| Arabidopsis | omegaNA | Ribosome__biogenesis | Glycosyltransferases | 0.4482759 |

*(continued)*

| species | estimate | var1 | var2 | p.value |
|---|---|---|---|---|
| Arabidopsis | omegaNA | Ribosome__biogenesis | Ion__channels | 0.0344828 |
| Arabidopsis | omegaNA | Ribosome__biogenesis | Lipid__biosynthesis__proteins | 0.1724138 |
| Arabidopsis | omegaNA | Ribosome__biogenesis | Membrane__trafficking | 0.7931034 |
| Arabidopsis | omegaNA | Ribosome__biogenesis | Messenger__RNA__biogenesis | 0.7241379 |
| Arabidopsis | omegaNA | Ribosome__biogenesis | Mitochondrial__biogenesis | 0.1034483 |
| Arabidopsis | omegaNA | Ribosome__biogenesis | Peptidases | 0.5862069 |
| Arabidopsis | omegaNA | Ribosome__biogenesis | Photosynthesis__proteins | 0.2413793 |
| Arabidopsis | omegaNA | Ribosome__biogenesis | Proteasome | 0.1034483 |
| Arabidopsis | omegaNA | Ribosome__biogenesis | Protein__kinases | 0.3103448 |
| Arabidopsis | omegaNA | Ribosome__biogenesis | Protein__phosphatases__and__associated__proteins | 0.5172414 |
| Arabidopsis | omegaNA | Ribosome__biogenesis | Ribosome | 0.1034483 |
| Arabidopsis | omegaNA | Spliceosome | Amino__acid__related__enzymes | 0.4482759 |
| Arabidopsis | omegaNA | Spliceosome | Chaperones__and__folding__catalysts | 0.3103448 |
| Arabidopsis | omegaNA | Spliceosome | Chromosome__and__associated__proteins | 0.1724138 |
| Arabidopsis | omegaNA | Spliceosome | Cytochrome__P450 | 0.1034483 |
| Arabidopsis | omegaNA | Spliceosome | Cytoskeleton__proteins | 0.6551724 |
| Arabidopsis | omegaNA | Spliceosome | DNA__repair__and__recombination__proteins | 0.0344828 |
| Arabidopsis | omegaNA | Spliceosome | DNA__replication__proteins | 0.3103448 |
| Arabidopsis | omegaNA | Spliceosome | Exosome | 0.0344828 |
| Arabidopsis | omegaNA | Spliceosome | G__protein__coupled__receptors | 0.1724138 |
| Arabidopsis | omegaNA | Spliceosome | Glycosyltransferases | 0.2413793 |
| Arabidopsis | omegaNA | Spliceosome | Ion__channels | 0.0344828 |
| Arabidopsis | omegaNA | Spliceosome | Lipid__biosynthesis__proteins | 0.1034483 |
| Arabidopsis | omegaNA | Spliceosome | Membrane__trafficking | 0.3103448 |
| Arabidopsis | omegaNA | Spliceosome | Messenger__RNA__biogenesis | 0.8620690 |
| Arabidopsis | omegaNA | Spliceosome | Mitochondrial__biogenesis | 0.1034483 |
| Arabidopsis | omegaNA | Spliceosome | Peptidases | 0.3103448 |
| Arabidopsis | omegaNA | Spliceosome | Photosynthesis__proteins | 0.1034483 |
| Arabidopsis | omegaNA | Spliceosome | Proteasome | 0.1724138 |
| Arabidopsis | omegaNA | Spliceosome | Protein__kinases | 0.2413793 |
| Arabidopsis | omegaNA | Spliceosome | Protein__phosphatases__and__associated__proteins | 0.4482759 |
| Arabidopsis | omegaNA | Spliceosome | Ribosome | 0.1034483 |
| Arabidopsis | omegaNA | Spliceosome | Ribosome__biogenesis | 0.6551724 |
| Arabidopsis | omegaNA | Transcription__factors | Amino__acid__related__enzymes | 0.1034483 |
| Arabidopsis | omegaNA | Transcription__factors | Chaperones__and__folding__catalysts | 0.1034483 |
| Arabidopsis | omegaNA | Transcription__factors | Chromosome__and__associated__proteins | 0.7931034 |
| Arabidopsis | omegaNA | Transcription__factors | Cytochrome__P450 | 0.0344828 |
| Arabidopsis | omegaNA | Transcription__factors | Cytoskeleton__proteins | 0.2413793 |
| Arabidopsis | omegaNA | Transcription__factors | DNA__repair__and__recombination__proteins | 0.5172414 |
| Arabidopsis | omegaNA | Transcription__factors | DNA__replication__proteins | 0.3103448 |
| Arabidopsis | omegaNA | Transcription__factors | Exosome | 0.0344828 |
| Arabidopsis | omegaNA | Transcription__factors | G__protein__coupled__receptors | 0.0344828 |
| Arabidopsis | omegaNA | Transcription__factors | Glycosyltransferases | 0.0344828 |
| Arabidopsis | omegaNA | Transcription__factors | Ion__channels | 0.0344828 |
| Arabidopsis | omegaNA | Transcription__factors | Lipid__biosynthesis__proteins | 0.0344828 |
| Arabidopsis | omegaNA | Transcription__factors | Membrane__trafficking | 0.0344828 |
| Arabidopsis | omegaNA | Transcription__factors | Messenger__RNA__biogenesis | 0.1724138 |
| Arabidopsis | omegaNA | Transcription__factors | Mitochondrial__biogenesis | 0.7931034 |
| Arabidopsis | omegaNA | Transcription__factors | Peptidases | 0.0344828 |
| Arabidopsis | omegaNA | Transcription__factors | Photosynthesis__proteins | 0.0344828 |
| Arabidopsis | omegaNA | Transcription__factors | Proteasome | 0.0344828 |
| Arabidopsis | omegaNA | Transcription__factors | Protein__kinases | 0.0344828 |
| Arabidopsis | omegaNA | Transcription__factors | Protein__phosphatases__and__associated__proteins | 0.1034483 |
| Arabidopsis | omegaNA | Transcription__factors | Ribosome | 0.0344828 |
| Arabidopsis | omegaNA | Transcription__factors | Ribosome__biogenesis | 0.0344828 |
| Arabidopsis | omegaNA | Transcription__factors | Spliceosome | 0.1724138 |
| Arabidopsis | omegaNA | Transcription__machinery | Amino__acid__related__enzymes | 0.5862069 |
| Arabidopsis | omegaNA | Transcription__machinery | Chaperones__and__folding__catalysts | 0.3793103 |
| Arabidopsis | omegaNA | Transcription__machinery | Chromosome__and__associated__proteins | 0.1034483 |
| Arabidopsis | omegaNA | Transcription__machinery | Cytochrome__P450 | 0.2413793 |
| Arabidopsis | omegaNA | Transcription__machinery | Cytoskeleton__proteins | 1.0000000 |
| Arabidopsis | omegaNA | Transcription__machinery | DNA__repair__and__recombination__proteins | 0.0344828 |

*(continued)*

| species | estimate | var1 | var2 | p.value |
|---|---|---|---|---|
| Arabidopsis | omegaNA | Transcription__machinery | DNA__replication__proteins | 0.2413793 |
| Arabidopsis | omegaNA | Transcription__machinery | Exosome | 0.3103448 |
| Arabidopsis | omegaNA | Transcription__machinery | G__protein__coupled__receptors | 0.5172414 |
| Arabidopsis | omegaNA | Transcription__machinery | Glycosyltransferases | 0.3103448 |
| Arabidopsis | omegaNA | Transcription__machinery | Ion__channels | 0.0344828 |
| Arabidopsis | omegaNA | Transcription__machinery | Lipid__biosynthesis__proteins | 0.2413793 |
| Arabidopsis | omegaNA | Transcription__machinery | Membrane__trafficking | 0.6551724 |
| Arabidopsis | omegaNA | Transcription__machinery | Messenger__RNA__biogenesis | 0.6551724 |
| Arabidopsis | omegaNA | Transcription__machinery | Mitochondrial__biogenesis | 0.1034483 |
| Arabidopsis | omegaNA | Transcription__machinery | Peptidases | 0.6551724 |
| Arabidopsis | omegaNA | Transcription__machinery | Photosynthesis__proteins | 0.1034483 |
| Arabidopsis | omegaNA | Transcription__machinery | Proteasome | 0.0344828 |
| Arabidopsis | omegaNA | Transcription__machinery | Protein__kinases | 0.3793103 |
| Arabidopsis | omegaNA | Transcription__machinery | Protein__phosphatases__and__associated__proteins | 0.7241379 |
| Arabidopsis | omegaNA | Transcription__machinery | Ribosome | 0.1034483 |
| Arabidopsis | omegaNA | Transcription__machinery | Ribosome__biogenesis | 0.7241379 |
| Arabidopsis | omegaNA | Transcription__machinery | Spliceosome | 0.4482759 |
| Arabidopsis | omegaNA | Transcription__machinery | Transcription__factors | 0.1034483 |
| Arabidopsis | omegaNA | Transfer__RNA__biogenesis | Amino__acid__related__enzymes | 0.1724138 |
| Arabidopsis | omegaNA | Transfer__RNA__biogenesis | Chaperones__and__folding__catalysts | 0.1034483 |
| Arabidopsis | omegaNA | Transfer__RNA__biogenesis | Chromosome__and__associated__proteins | 0.6551724 |
| Arabidopsis | omegaNA | Transfer__RNA__biogenesis | Cytochrome__P450 | 0.1034483 |
| Arabidopsis | omegaNA | Transfer__RNA__biogenesis | Cytoskeleton__proteins | 0.3793103 |
| Arabidopsis | omegaNA | Transfer__RNA__biogenesis | DNA__repair__and__recombination__proteins | 0.2413793 |
| Arabidopsis | omegaNA | Transfer__RNA__biogenesis | DNA__replication__proteins | 0.7241379 |
| Arabidopsis | omegaNA | Transfer__RNA__biogenesis | Exosome | 0.0344828 |
| Arabidopsis | omegaNA | Transfer__RNA__biogenesis | G__protein__coupled__receptors | 0.1034483 |
| Arabidopsis | omegaNA | Transfer__RNA__biogenesis | Glycosyltransferases | 0.0344828 |
| Arabidopsis | omegaNA | Transfer__RNA__biogenesis | Ion__channels | 0.0344828 |
| Arabidopsis | omegaNA | Transfer__RNA__biogenesis | Lipid__biosynthesis__proteins | 0.0344828 |
| Arabidopsis | omegaNA | Transfer__RNA__biogenesis | Membrane__trafficking | 0.1034483 |
| Arabidopsis | omegaNA | Transfer__RNA__biogenesis | Messenger__RNA__biogenesis | 0.5172414 |
| Arabidopsis | omegaNA | Transfer__RNA__biogenesis | Mitochondrial__biogenesis | 0.6551724 |
| Arabidopsis | omegaNA | Transfer__RNA__biogenesis | Peptidases | 0.0344828 |
| Arabidopsis | omegaNA | Transfer__RNA__biogenesis | Photosynthesis__proteins | 0.0344828 |
| Arabidopsis | omegaNA | Transfer__RNA__biogenesis | Proteasome | 0.0344828 |
| Arabidopsis | omegaNA | Transfer__RNA__biogenesis | Protein__kinases | 0.0344828 |
| Arabidopsis | omegaNA | Transfer__RNA__biogenesis | Protein__phosphatases__and__associated__proteins | 0.2413793 |
| Arabidopsis | omegaNA | Transfer__RNA__biogenesis | Ribosome | 0.0344828 |
| Arabidopsis | omegaNA | Transfer__RNA__biogenesis | Ribosome__biogenesis | 0.1724138 |
| Arabidopsis | omegaNA | Transfer__RNA__biogenesis | Spliceosome | 0.3793103 |
| Arabidopsis | omegaNA | Transfer__RNA__biogenesis | Transcription__factors | 0.5172414 |
| Arabidopsis | omegaNA | Transfer__RNA__biogenesis | Transcription__machinery | 0.4482759 |
| Arabidopsis | omegaNA | Translation__factors | Amino__acid__related__enzymes | 0.9310345 |
| Arabidopsis | omegaNA | Translation__factors | Chaperones__and__folding__catalysts | 0.9310345 |
| Arabidopsis | omegaNA | Translation__factors | Chromosome__and__associated__proteins | 0.0344828 |
| Arabidopsis | omegaNA | Translation__factors | Cytochrome__P450 | 0.4482759 |
| Arabidopsis | omegaNA | Translation__factors | Cytoskeleton__proteins | 1.0000000 |
| Arabidopsis | omegaNA | Translation__factors | DNA__repair__and__recombination__proteins | 0.0344828 |
| Arabidopsis | omegaNA | Translation__factors | DNA__replication__proteins | 0.1724138 |
| Arabidopsis | omegaNA | Translation__factors | Exosome | 0.5862069 |
| Arabidopsis | omegaNA | Translation__factors | G__protein__coupled__receptors | 0.7931034 |
| Arabidopsis | omegaNA | Translation__factors | Glycosyltransferases | 0.6551724 |
| Arabidopsis | omegaNA | Translation__factors | Ion__channels | 0.1034483 |
| Arabidopsis | omegaNA | Translation__factors | Lipid__biosynthesis__proteins | 0.3103448 |
| Arabidopsis | omegaNA | Translation__factors | Membrane__trafficking | 0.9310345 |
| Arabidopsis | omegaNA | Translation__factors | Messenger__RNA__biogenesis | 0.3793103 |
| Arabidopsis | omegaNA | Translation__factors | Mitochondrial__biogenesis | 0.0344828 |
| Arabidopsis | omegaNA | Translation__factors | Peptidases | 0.7241379 |
| Arabidopsis | omegaNA | Translation__factors | Photosynthesis__proteins | 0.3793103 |
| Arabidopsis | omegaNA | Translation__factors | Proteasome | 0.3103448 |
| Arabidopsis | omegaNA | Translation__factors | Protein__kinases | 0.7931034 |

*(continued)*

| species | estimate | var1 | var2 | p.value |
|---|---|---|---|---|
| Arabidopsis | omegaNA | Translation__factors | Protein__phosphatases__and__associated__proteins | 0.9310345 |
| Arabidopsis | omegaNA | Translation__factors | Ribosome | 0.5172414 |
| Arabidopsis | omegaNA | Translation__factors | Ribosome__biogenesis | 0.5862069 |
| Arabidopsis | omegaNA | Translation__factors | Spliceosome | 0.3793103 |
| Arabidopsis | omegaNA | Translation__factors | Transcription__factors | 0.0344828 |
| Arabidopsis | omegaNA | Translation__factors | Transcription__machinery | 0.3793103 |
| Arabidopsis | omegaNA | Translation__factors | Transfer__RNA__biogenesis | 0.0344828 |
| Arabidopsis | omegaNA | Transporters | Amino__acid__related__enzymes | 0.2413793 |
| Arabidopsis | omegaNA | Transporters | Chaperones__and__folding__catalysts | 0.3793103 |
| Arabidopsis | omegaNA | Transporters | Chromosome__and__associated__proteins | 0.0344828 |
| Arabidopsis | omegaNA | Transporters | Cytochrome__P450 | 0.6551724 |
| Arabidopsis | omegaNA | Transporters | Cytoskeleton__proteins | 0.2413793 |
| Arabidopsis | omegaNA | Transporters | DNA__repair__and__recombination__proteins | 0.0344828 |
| Arabidopsis | omegaNA | Transporters | DNA__replication__proteins | 0.0344828 |
| Arabidopsis | omegaNA | Transporters | Exosome | 1.0000000 |
| Arabidopsis | omegaNA | Transporters | G__protein__coupled__receptors | 0.2413793 |
| Arabidopsis | omegaNA | Transporters | Glycosyltransferases | 0.1724138 |
| Arabidopsis | omegaNA | Transporters | Ion__channels | 0.1034483 |
| Arabidopsis | omegaNA | Transporters | Lipid__biosynthesis__proteins | 0.7931034 |
| Arabidopsis | omegaNA | Transporters | Membrane__trafficking | 0.1034483 |
| Arabidopsis | omegaNA | Transporters | Messenger__RNA__biogenesis | 0.0344828 |
| Arabidopsis | omegaNA | Transporters | Mitochondrial__biogenesis | 0.0344828 |
| Arabidopsis | omegaNA | Transporters | Peptidases | 0.1724138 |
| Arabidopsis | omegaNA | Transporters | Photosynthesis__proteins | 1.0000000 |
| Arabidopsis | omegaNA | Transporters | Proteasome | 0.8620690 |
| Arabidopsis | omegaNA | Transporters | Protein__kinases | 0.3793103 |
| Arabidopsis | omegaNA | Transporters | Protein__phosphatases__and__associated__proteins | 0.3793103 |
| Arabidopsis | omegaNA | Transporters | Ribosome | 1.0000000 |
| Arabidopsis | omegaNA | Transporters | Ribosome__biogenesis | 0.0344828 |
| Arabidopsis | omegaNA | Transporters | Spliceosome | 0.0344828 |
| Arabidopsis | omegaNA | Transporters | Transcription__factors | 0.0344828 |
| Arabidopsis | omegaNA | Transporters | Transcription__machinery | 0.0344828 |
| Arabidopsis | omegaNA | Transporters | Transfer__RNA__biogenesis | 0.0344828 |
| Arabidopsis | omegaNA | Transporters | Translation__factors | 0.3793103 |
| Arabidopsis | omegaNA | Ubiquitin__system | Amino__acid__related__enzymes | 0.8620690 |
| Arabidopsis | omegaNA | Ubiquitin__system | Chaperones__and__folding__catalysts | 0.4482759 |
| Arabidopsis | omegaNA | Ubiquitin__system | Chromosome__and__associated__proteins | 0.1034483 |
| Arabidopsis | omegaNA | Ubiquitin__system | Cytochrome__P450 | 0.2413793 |
| Arabidopsis | omegaNA | Ubiquitin__system | Cytoskeleton__proteins | 1.0000000 |
| Arabidopsis | omegaNA | Ubiquitin__system | DNA__repair__and__recombination__proteins | 0.0344828 |
| Arabidopsis | omegaNA | Ubiquitin__system | DNA__replication__proteins | 0.0344828 |
| Arabidopsis | omegaNA | Ubiquitin__system | Exosome | 0.2413793 |
| Arabidopsis | omegaNA | Ubiquitin__system | G__protein__coupled__receptors | 0.7931034 |
| Arabidopsis | omegaNA | Ubiquitin__system | Glycosyltransferases | 0.4482759 |
| Arabidopsis | omegaNA | Ubiquitin__system | Ion__channels | 0.0344828 |
| Arabidopsis | omegaNA | Ubiquitin__system | Lipid__biosynthesis__proteins | 0.1034483 |
| Arabidopsis | omegaNA | Ubiquitin__system | Membrane__trafficking | 0.6551724 |
| Arabidopsis | omegaNA | Ubiquitin__system | Messenger__RNA__biogenesis | 0.6551724 |
| Arabidopsis | omegaNA | Ubiquitin__system | Mitochondrial__biogenesis | 0.0344828 |
| Arabidopsis | omegaNA | Ubiquitin__system | Peptidases | 0.5862069 |
| Arabidopsis | omegaNA | Ubiquitin__system | Photosynthesis__proteins | 0.1724138 |
| Arabidopsis | omegaNA | Ubiquitin__system | Proteasome | 0.0344828 |
| Arabidopsis | omegaNA | Ubiquitin__system | Protein__kinases | 0.5172414 |
| Arabidopsis | omegaNA | Ubiquitin__system | Protein__phosphatases__and__associated__proteins | 0.6551724 |
| Arabidopsis | omegaNA | Ubiquitin__system | Ribosome | 0.0344828 |
| Arabidopsis | omegaNA | Ubiquitin__system | Ribosome__biogenesis | 1.0000000 |
| Arabidopsis | omegaNA | Ubiquitin__system | Spliceosome | 0.3793103 |
| Arabidopsis | omegaNA | Ubiquitin__system | Transcription__factors | 0.0344828 |
| Arabidopsis | omegaNA | Ubiquitin__system | Transcription__machinery | 0.7241379 |
| Arabidopsis | omegaNA | Ubiquitin__system | Transfer__RNA__biogenesis | 0.1724138 |
| Arabidopsis | omegaNA | Ubiquitin__system | Translation__factors | 0.6551724 |

*(continued)*

| species | estimate | var1 | var2 | p.value |
|---|---|---|---|---|
| Arabidopsis | omegaNA | Ubiquitin__system | Transporters | 0.0344828 |
| Arabidopsis | omegaA | Chaperones__and__folding__catalysts | Amino__acid__related__enzymes | 0.1034483 |
| Arabidopsis | omegaA | Chromosome__and__associated__proteins | Amino__acid__related__enzymes | 0.3793103 |
| Arabidopsis | omegaA | Chromosome__and__associated__proteins | Chaperones__and__folding__catalysts | 0.4482759 |
| Arabidopsis | omegaA | Cytochrome__P450 | Amino__acid__related__enzymes | 0.1034483 |
| Arabidopsis | omegaA | Cytochrome__P450 | Chaperones__and__folding__catalysts | 0.3103448 |
| Arabidopsis | omegaA | Cytochrome__P450 | Chromosome__and__associated__proteins | 0.1724138 |
| Arabidopsis | omegaA | Cytoskeleton__proteins | Amino__acid__related__enzymes | 0.3793103 |
| Arabidopsis | omegaA | Cytoskeleton__proteins | Chaperones__and__folding__catalysts | 0.8620690 |
| Arabidopsis | omegaA | Cytoskeleton__proteins | Chromosome__and__associated__proteins | 0.7931034 |
| Arabidopsis | omegaA | Cytoskeleton__proteins | Cytochrome__P450 | 0.4482759 |
| Arabidopsis | omegaA | DNA__repair__and__recombination__proteins | Amino__acid__related__enzymes | 0.9310345 |
| Arabidopsis | omegaA | DNA__repair__and__recombination__proteins | Chaperones__and__folding__catalysts | 0.2413793 |
| Arabidopsis | omegaA | DNA__repair__and__recombination__proteins | Chromosome__and__associated__proteins | 0.6551724 |
| Arabidopsis | omegaA | DNA__repair__and__recombination__proteins | Cytochrome__P450 | 0.1034483 |
| Arabidopsis | omegaA | DNA__repair__and__recombination__proteins | Cytoskeleton__proteins | 0.5862069 |
| Arabidopsis | omegaA | DNA__replication__proteins | Amino__acid__related__enzymes | 0.5862069 |
| Arabidopsis | omegaA | DNA__replication__proteins | Chaperones__and__folding__catalysts | 0.5172414 |
| Arabidopsis | omegaA | DNA__replication__proteins | Chromosome__and__associated__proteins | 0.9310345 |
| Arabidopsis | omegaA | DNA__replication__proteins | Cytochrome__P450 | 0.2413793 |
| Arabidopsis | omegaA | DNA__replication__proteins | Cytoskeleton__proteins | 0.8620690 |
| Arabidopsis | omegaA | DNA__replication__proteins | DNA__repair__and__recombination__proteins | 0.5862069 |
| Arabidopsis | omegaA | Exosome | Amino__acid__related__enzymes | 0.6551724 |
| Arabidopsis | omegaA | Exosome | Chaperones__and__folding__catalysts | 0.3793103 |
| Arabidopsis | omegaA | Exosome | Chromosome__and__associated__proteins | 0.9310345 |
| Arabidopsis | omegaA | Exosome | Cytochrome__P450 | 0.1034483 |
| Arabidopsis | omegaA | Exosome | Cytoskeleton__proteins | 0.5862069 |
| Arabidopsis | omegaA | Exosome | DNA__repair__and__recombination__proteins | 0.5862069 |
| Arabidopsis | omegaA | Exosome | DNA__replication__proteins | 0.7241379 |
| Arabidopsis | omegaA | G__protein__coupled__receptors | Amino__acid__related__enzymes | 0.0344828 |
| Arabidopsis | omegaA | G__protein__coupled__receptors | Chaperones__and__folding__catalysts | 0.3793103 |
| Arabidopsis | omegaA | G__protein__coupled__receptors | Chromosome__and__associated__proteins | 0.0344828 |
| Arabidopsis | omegaA | G__protein__coupled__receptors | Cytochrome__P450 | 0.9310345 |
| Arabidopsis | omegaA | G__protein__coupled__receptors | Cytoskeleton__proteins | 0.5862069 |
| Arabidopsis | omegaA | G__protein__coupled__receptors | DNA__repair__and__recombination__proteins | 0.0344828 |
| Arabidopsis | omegaA | G__protein__coupled__receptors | DNA__replication__proteins | 0.1724138 |
| Arabidopsis | omegaA | G__protein__coupled__receptors | Exosome | 0.0344828 |
| Arabidopsis | omegaA | Glycosyltransferases | Amino__acid__related__enzymes | 0.1034483 |
| Arabidopsis | omegaA | Glycosyltransferases | Chaperones__and__folding__catalysts | 0.8620690 |
| Arabidopsis | omegaA | Glycosyltransferases | Chromosome__and__associated__proteins | 0.7931034 |
| Arabidopsis | omegaA | Glycosyltransferases | Cytochrome__P450 | 0.3103448 |
| Arabidopsis | omegaA | Glycosyltransferases | Cytoskeleton__proteins | 0.9310345 |
| Arabidopsis | omegaA | Glycosyltransferases | DNA__repair__and__recombination__proteins | 0.3103448 |
| Arabidopsis | omegaA | Glycosyltransferases | DNA__replication__proteins | 0.7241379 |
| Arabidopsis | omegaA | Glycosyltransferases | Exosome | 0.3103448 |
| Arabidopsis | omegaA | Glycosyltransferases | G__protein__coupled__receptors | 0.0344828 |
| Arabidopsis | omegaA | Ion__channels | Amino__acid__related__enzymes | 0.0344828 |
| Arabidopsis | omegaA | Ion__channels | Chaperones__and__folding__catalysts | 0.5862069 |
| Arabidopsis | omegaA | Ion__channels | Chromosome__and__associated__proteins | 0.1034483 |
| Arabidopsis | omegaA | Ion__channels | Cytochrome__P450 | 0.7241379 |
| Arabidopsis | omegaA | Ion__channels | Cytoskeleton__proteins | 0.7931034 |
| Arabidopsis | omegaA | Ion__channels | DNA__repair__and__recombination__proteins | 0.1724138 |
| Arabidopsis | omegaA | Ion__channels | DNA__replication__proteins | 0.3103448 |
| Arabidopsis | omegaA | Ion__channels | Exosome | 0.0344828 |
| Arabidopsis | omegaA | Ion__channels | G__protein__coupled__receptors | 0.6551724 |
| Arabidopsis | omegaA | Ion__channels | Glycosyltransferases | 0.1034483 |
| Arabidopsis | omegaA | Lipid__biosynthesis__proteins | Amino__acid__related__enzymes | 0.0344828 |
| Arabidopsis | omegaA | Lipid__biosynthesis__proteins | Chaperones__and__folding__catalysts | 0.3103448 |
| Arabidopsis | omegaA | Lipid__biosynthesis__proteins | Chromosome__and__associated__proteins | 0.1034483 |
| Arabidopsis | omegaA | Lipid__biosynthesis__proteins | Cytochrome__P450 | 0.7931034 |
| Arabidopsis | omegaA | Lipid__biosynthesis__proteins | Cytoskeleton__proteins | 0.4482759 |
| Arabidopsis | omegaA | Lipid__biosynthesis__proteins | DNA__repair__and__recombination__proteins | 0.1034483 |

*(continued)*

| species | estimate | var1 | var2 | p.value |
|---|---|---|---|---|
| Arabidopsis | omegaA | Lipid__biosynthesis__proteins | DNA__replication__proteins | 0.0344828 |
| Arabidopsis | omegaA | Lipid__biosynthesis__proteins | Exosome | 0.0344828 |
| Arabidopsis | omegaA | Lipid__biosynthesis__proteins | G__protein__coupled__receptors | 0.9310345 |
| Arabidopsis | omegaA | Lipid__biosynthesis__proteins | Glycosyltransferases | 0.2413793 |
| Arabidopsis | omegaA | Lipid__biosynthesis__proteins | Ion__channels | 0.6551724 |
| Arabidopsis | omegaA | Membrane__trafficking | Amino__acid__related__enzymes | 0.1034483 |
| Arabidopsis | omegaA | Membrane__trafficking | Chaperones__and__folding__catalysts | 0.7241379 |
| Arabidopsis | omegaA | Membrane__trafficking | Chromosome__and__associated__proteins | 0.7931034 |
| Arabidopsis | omegaA | Membrane__trafficking | Cytochrome__P450 | 0.1724138 |
| Arabidopsis | omegaA | Membrane__trafficking | Cytoskeleton__proteins | 1.0000000 |
| Arabidopsis | omegaA | Membrane__trafficking | DNA__repair__and__recombination__proteins | 0.2413793 |
| Arabidopsis | omegaA | Membrane__trafficking | DNA__replication__proteins | 0.6551724 |
| Arabidopsis | omegaA | Membrane__trafficking | Exosome | 0.3793103 |
| Arabidopsis | omegaA | Membrane__trafficking | G__protein__coupled__receptors | 0.1724138 |
| Arabidopsis | omegaA | Membrane__trafficking | Glycosyltransferases | 0.8620690 |
| Arabidopsis | omegaA | Membrane__trafficking | Ion__channels | 0.1724138 |
| Arabidopsis | omegaA | Membrane__trafficking | Lipid__biosynthesis__proteins | 0.1724138 |
| Arabidopsis | omegaA | Messenger__RNA__biogenesis | Amino__acid__related__enzymes | 0.5172414 |
| Arabidopsis | omegaA | Messenger__RNA__biogenesis | Chaperones__and__folding__catalysts | 0.7931034 |
| Arabidopsis | omegaA | Messenger__RNA__biogenesis | Chromosome__and__associated__proteins | 0.8620690 |
| Arabidopsis | omegaA | Messenger__RNA__biogenesis | Cytochrome__P450 | 0.3793103 |
| Arabidopsis | omegaA | Messenger__RNA__biogenesis | Cytoskeleton__proteins | 0.8620690 |
| Arabidopsis | omegaA | Messenger__RNA__biogenesis | DNA__repair__and__recombination__proteins | 0.5862069 |
| Arabidopsis | omegaA | Messenger__RNA__biogenesis | DNA__replication__proteins | 0.9310345 |
| Arabidopsis | omegaA | Messenger__RNA__biogenesis | Exosome | 0.7241379 |
| Arabidopsis | omegaA | Messenger__RNA__biogenesis | G__protein__coupled__receptors | 0.3103448 |
| Arabidopsis | omegaA | Messenger__RNA__biogenesis | Glycosyltransferases | 0.8620690 |
| Arabidopsis | omegaA | Messenger__RNA__biogenesis | Ion__channels | 0.4482759 |
| Arabidopsis | omegaA | Messenger__RNA__biogenesis | Lipid__biosynthesis__proteins | 0.1724138 |
| Arabidopsis | omegaA | Messenger__RNA__biogenesis | Membrane__trafficking | 0.9310345 |
| Arabidopsis | omegaA | Mitochondrial__biogenesis | Amino__acid__related__enzymes | 0.8620690 |
| Arabidopsis | omegaA | Mitochondrial__biogenesis | Chaperones__and__folding__catalysts | 0.1034483 |
| Arabidopsis | omegaA | Mitochondrial__biogenesis | Chromosome__and__associated__proteins | 0.3793103 |
| Arabidopsis | omegaA | Mitochondrial__biogenesis | Cytochrome__P450 | 0.1034483 |
| Arabidopsis | omegaA | Mitochondrial__biogenesis | Cytoskeleton__proteins | 0.3793103 |
| Arabidopsis | omegaA | Mitochondrial__biogenesis | DNA__repair__and__recombination__proteins | 0.6551724 |
| Arabidopsis | omegaA | Mitochondrial__biogenesis | DNA__replication__proteins | 0.4482759 |
| Arabidopsis | omegaA | Mitochondrial__biogenesis | Exosome | 0.5172414 |
| Arabidopsis | omegaA | Mitochondrial__biogenesis | G__protein__coupled__receptors | 0.0344828 |
| Arabidopsis | omegaA | Mitochondrial__biogenesis | Glycosyltransferases | 0.0344828 |
| Arabidopsis | omegaA | Mitochondrial__biogenesis | Ion__channels | 0.0344828 |
| Arabidopsis | omegaA | Mitochondrial__biogenesis | Lipid__biosynthesis__proteins | 0.0344828 |
| Arabidopsis | omegaA | Mitochondrial__biogenesis | Membrane__trafficking | 0.2413793 |
| Arabidopsis | omegaA | Mitochondrial__biogenesis | Messenger__RNA__biogenesis | 0.3793103 |
| Arabidopsis | omegaA | Peptidases | Amino__acid__related__enzymes | 0.0344828 |
| Arabidopsis | omegaA | Peptidases | Chaperones__and__folding__catalysts | 0.7931034 |
| Arabidopsis | omegaA | Peptidases | Chromosome__and__associated__proteins | 0.5862069 |
| Arabidopsis | omegaA | Peptidases | Cytochrome__P450 | 0.4482759 |
| Arabidopsis | omegaA | Peptidases | Cytoskeleton__proteins | 0.8620690 |
| Arabidopsis | omegaA | Peptidases | DNA__repair__and__recombination__proteins | 0.1724138 |
| Arabidopsis | omegaA | Peptidases | DNA__replication__proteins | 0.3793103 |
| Arabidopsis | omegaA | Peptidases | Exosome | 0.0344828 |
| Arabidopsis | omegaA | Peptidases | G__protein__coupled__receptors | 0.3793103 |
| Arabidopsis | omegaA | Peptidases | Glycosyltransferases | 0.3793103 |
| Arabidopsis | omegaA | Peptidases | Ion__channels | 0.5172414 |
| Arabidopsis | omegaA | Peptidases | Lipid__biosynthesis__proteins | 0.3793103 |
| Arabidopsis | omegaA | Peptidases | Membrane__trafficking | 0.6551724 |
| Arabidopsis | omegaA | Peptidases | Messenger__RNA__biogenesis | 0.5172414 |
| Arabidopsis | omegaA | Peptidases | Mitochondrial__biogenesis | 0.0344828 |
| Arabidopsis | omegaA | Photosynthesis__proteins | Amino__acid__related__enzymes | 0.2413793 |
| Arabidopsis | omegaA | Photosynthesis__proteins | Chaperones__and__folding__catalysts | 0.5862069 |
| Arabidopsis | omegaA | Photosynthesis__proteins | Chromosome__and__associated__proteins | 0.9310345 |

*(continued)*

| species | estimate | var1 | var2 | p.value |
|---|---|---|---|---|
| Arabidopsis | omegaA | Photosynthesis__proteins | Cytochrome__P450 | 0.1724138 |
| Arabidopsis | omegaA | Photosynthesis__proteins | Cytoskeleton__proteins | 0.8620690 |
| Arabidopsis | omegaA | Photosynthesis__proteins | DNA__repair__and__recombination__proteins | 0.5862069 |
| Arabidopsis | omegaA | Photosynthesis__proteins | DNA__replication__proteins | 0.8620690 |
| Arabidopsis | omegaA | Photosynthesis__proteins | Exosome | 0.6551724 |
| Arabidopsis | omegaA | Photosynthesis__proteins | G__protein__coupled__receptors | 0.0344828 |
| Arabidopsis | omegaA | Photosynthesis__proteins | Glycosyltransferases | 0.5862069 |
| Arabidopsis | omegaA | Photosynthesis__proteins | Ion__channels | 0.0344828 |
| Arabidopsis | omegaA | Photosynthesis__proteins | Lipid__biosynthesis__proteins | 0.1034483 |
| Arabidopsis | omegaA | Photosynthesis__proteins | Membrane__trafficking | 0.7931034 |
| Arabidopsis | omegaA | Photosynthesis__proteins | Messenger__RNA__biogenesis | 0.7241379 |
| Arabidopsis | omegaA | Photosynthesis__proteins | Mitochondrial__biogenesis | 0.2413793 |
| Arabidopsis | omegaA | Photosynthesis__proteins | Peptidases | 0.2413793 |
| Arabidopsis | omegaA | Proteasome | Amino__acid__related__enzymes | 0.5172414 |
| Arabidopsis | omegaA | Proteasome | Chaperones__and__folding__catalysts | 0.6551724 |
| Arabidopsis | omegaA | Proteasome | Chromosome__and__associated__proteins | 0.9310345 |
| Arabidopsis | omegaA | Proteasome | Cytochrome__P450 | 0.3793103 |
| Arabidopsis | omegaA | Proteasome | Cytoskeleton__proteins | 0.8620690 |
| Arabidopsis | omegaA | Proteasome | DNA__repair__and__recombination__proteins | 0.5172414 |
| Arabidopsis | omegaA | Proteasome | DNA__replication__proteins | 1.0000000 |
| Arabidopsis | omegaA | Proteasome | Exosome | 0.8620690 |
| Arabidopsis | omegaA | Proteasome | G__protein__coupled__receptors | 0.1724138 |
| Arabidopsis | omegaA | Proteasome | Glycosyltransferases | 0.7241379 |
| Arabidopsis | omegaA | Proteasome | Ion__channels | 0.3793103 |
| Arabidopsis | omegaA | Proteasome | Lipid__biosynthesis__proteins | 0.2413793 |
| Arabidopsis | omegaA | Proteasome | Membrane__trafficking | 0.7931034 |
| Arabidopsis | omegaA | Proteasome | Messenger__RNA__biogenesis | 0.9310345 |
| Arabidopsis | omegaA | Proteasome | Mitochondrial__biogenesis | 0.5172414 |
| Arabidopsis | omegaA | Proteasome | Peptidases | 0.5172414 |
| Arabidopsis | omegaA | Proteasome | Photosynthesis__proteins | 0.9310345 |
| Arabidopsis | omegaA | Protein__kinases | Amino__acid__related__enzymes | 0.3793103 |
| Arabidopsis | omegaA | Protein__kinases | Chaperones__and__folding__catalysts | 0.3793103 |
| Arabidopsis | omegaA | Protein__kinases | Chromosome__and__associated__proteins | 0.9310345 |
| Arabidopsis | omegaA | Protein__kinases | Cytochrome__P450 | 0.1034483 |
| Arabidopsis | omegaA | Protein__kinases | Cytoskeleton__proteins | 0.8620690 |
| Arabidopsis | omegaA | Protein__kinases | DNA__repair__and__recombination__proteins | 0.5172414 |
| Arabidopsis | omegaA | Protein__kinases | DNA__replication__proteins | 0.9310345 |
| Arabidopsis | omegaA | Protein__kinases | Exosome | 0.6551724 |
| Arabidopsis | omegaA | Protein__kinases | G__protein__coupled__receptors | 0.0344828 |
| Arabidopsis | omegaA | Protein__kinases | Glycosyltransferases | 0.7241379 |
| Arabidopsis | omegaA | Protein__kinases | Ion__channels | 0.1034483 |
| Arabidopsis | omegaA | Protein__kinases | Lipid__biosynthesis__proteins | 0.0344828 |
| Arabidopsis | omegaA | Protein__kinases | Membrane__trafficking | 0.7931034 |
| Arabidopsis | omegaA | Protein__kinases | Messenger__RNA__biogenesis | 0.7931034 |
| Arabidopsis | omegaA | Protein__kinases | Mitochondrial__biogenesis | 0.3103448 |
| Arabidopsis | omegaA | Protein__kinases | Peptidases | 0.1034483 |
| Arabidopsis | omegaA | Protein__kinases | Photosynthesis__proteins | 1.0000000 |
| Arabidopsis | omegaA | Protein__kinases | Proteasome | 1.0000000 |
| Arabidopsis | omegaA | Protein__phosphatases__and__associated__proteins | Amino__acid__related__enzymes | 0.0344828 |
| Arabidopsis | omegaA | Protein__phosphatases__and__associated__proteins | Chaperones__and__folding__catalysts | 0.5862069 |
| Arabidopsis | omegaA | Protein__phosphatases__and__associated__proteins | Chromosome__and__associated__proteins | 0.1724138 |
| Arabidopsis | omegaA | Protein__phosphatases__and__associated__proteins | Cytochrome__P450 | 0.8620690 |
| Arabidopsis | omegaA | Protein__phosphatases__and__associated__proteins | Cytoskeleton__proteins | 0.5862069 |
| Arabidopsis | omegaA | Protein__phosphatases__and__associated__proteins | DNA__repair__and__recombination__proteins | 0.1724138 |
| Arabidopsis | omegaA | Protein__phosphatases__and__associated__proteins | DNA__replication__proteins | 0.2413793 |
| Arabidopsis | omegaA | Protein__phosphatases__and__associated__proteins | Exosome | 0.1724138 |
| Arabidopsis | omegaA | Protein__phosphatases__and__associated__proteins | G__protein__coupled__receptors | 0.8620690 |
| Arabidopsis | omegaA | Protein__phosphatases__and__associated__proteins | Glycosyltransferases | 0.3103448 |
| Arabidopsis | omegaA | Protein__phosphatases__and__associated__proteins | Ion__channels | 0.7931034 |
| Arabidopsis | omegaA | Protein__phosphatases__and__associated__proteins | Lipid__biosynthesis__proteins | 0.7931034 |
| Arabidopsis | omegaA | Protein__phosphatases__and__associated__proteins | Membrane__trafficking | 0.3103448 |

*(continued)*

| species | estimate | var1 | var2 | p.value |
|---|---|---|---|---|
| Arabidopsis | omegaA | Protein__phosphatases__and__associated__proteins | Messenger__RNA__biogenesis | 0.4482759 |
| Arabidopsis | omegaA | Protein__phosphatases__and__associated__proteins | Mitochondrial__biogenesis | 0.0344828 |
| Arabidopsis | omegaA | Protein__phosphatases__and__associated__proteins | Peptidases | 0.4482759 |
| Arabidopsis | omegaA | Protein__phosphatases__and__associated__proteins | Photosynthesis__proteins | 0.1724138 |
| Arabidopsis | omegaA | Protein__phosphatases__and__associated__proteins | Proteasome | 0.2413793 |
| Arabidopsis | omegaA | Protein__phosphatases__and__associated__proteins | Protein__kinases | 0.1724138 |
| Arabidopsis | omegaA | Ribosome | Amino__acid__related__enzymes | 0.3103448 |
| Arabidopsis | omegaA | Ribosome | Chaperones__and__folding__catalysts | 0.5862069 |
| Arabidopsis | omegaA | Ribosome | Chromosome__and__associated__proteins | 0.9310345 |
| Arabidopsis | omegaA | Ribosome | Cytochrome__P450 | 0.1724138 |
| Arabidopsis | omegaA | Ribosome | Cytoskeleton__proteins | 0.9310345 |
| Arabidopsis | omegaA | Ribosome | DNA__repair__and__recombination__proteins | 0.3793103 |
| Arabidopsis | omegaA | Ribosome | DNA__replication__proteins | 1.0000000 |
| Arabidopsis | omegaA | Ribosome | Exosome | 0.7931034 |
| Arabidopsis | omegaA | Ribosome | G__protein__coupled__receptors | 0.0344828 |
| Arabidopsis | omegaA | Ribosome | Glycosyltransferases | 0.5172414 |
| Arabidopsis | omegaA | Ribosome | Ion__channels | 0.1724138 |
| Arabidopsis | omegaA | Ribosome | Lipid__biosynthesis__proteins | 0.1034483 |
| Arabidopsis | omegaA | Ribosome | Membrane__trafficking | 0.7241379 |
| Arabidopsis | omegaA | Ribosome | Messenger__RNA__biogenesis | 0.9310345 |
| Arabidopsis | omegaA | Ribosome | Mitochondrial__biogenesis | 0.0344828 |
| Arabidopsis | omegaA | Ribosome | Peptidases | 0.2413793 |
| Arabidopsis | omegaA | Ribosome | Photosynthesis__proteins | 0.7931034 |
| Arabidopsis | omegaA | Ribosome | Proteasome | 0.8620690 |
| Arabidopsis | omegaA | Ribosome | Protein__kinases | 0.7931034 |
| Arabidopsis | omegaA | Ribosome | Protein__phosphatases__and__associated__proteins | 0.2413793 |
| Arabidopsis | omegaA | Ribosome__biogenesis | Amino__acid__related__enzymes | 0.0344828 |
| Arabidopsis | omegaA | Ribosome__biogenesis | Chaperones__and__folding__catalysts | 0.1034483 |
| Arabidopsis | omegaA | Ribosome__biogenesis | Chromosome__and__associated__proteins | 0.0344828 |
| Arabidopsis | omegaA | Ribosome__biogenesis | Cytochrome__P450 | 0.5862069 |
| Arabidopsis | omegaA | Ribosome__biogenesis | Cytoskeleton__proteins | 0.1724138 |
| Arabidopsis | omegaA | Ribosome__biogenesis | DNA__repair__and__recombination__proteins | 0.0344828 |
| Arabidopsis | omegaA | Ribosome__biogenesis | DNA__replication__proteins | 0.1034483 |
| Arabidopsis | omegaA | Ribosome__biogenesis | Exosome | 0.0344828 |
| Arabidopsis | omegaA | Ribosome__biogenesis | G__protein__coupled__receptors | 0.5172414 |
| Arabidopsis | omegaA | Ribosome__biogenesis | Glycosyltransferases | 0.1034483 |
| Arabidopsis | omegaA | Ribosome__biogenesis | Ion__channels | 0.1724138 |
| Arabidopsis | omegaA | Ribosome__biogenesis | Lipid__biosynthesis__proteins | 0.5172414 |
| Arabidopsis | omegaA | Ribosome__biogenesis | Membrane__trafficking | 0.1034483 |
| Arabidopsis | omegaA | Ribosome__biogenesis | Messenger__RNA__biogenesis | 0.1034483 |
| Arabidopsis | omegaA | Ribosome__biogenesis | Mitochondrial__biogenesis | 0.0344828 |
| Arabidopsis | omegaA | Ribosome__biogenesis | Peptidases | 0.1034483 |
| Arabidopsis | omegaA | Ribosome__biogenesis | Photosynthesis__proteins | 0.0344828 |
| Arabidopsis | omegaA | Ribosome__biogenesis | Proteasome | 0.0344828 |
| Arabidopsis | omegaA | Ribosome__biogenesis | Protein__kinases | 0.0344828 |
| Arabidopsis | omegaA | Ribosome__biogenesis | Protein__phosphatases__and__associated__proteins | 0.5862069 |
| Arabidopsis | omegaA | Ribosome__biogenesis | Ribosome | 0.0344828 |
| Arabidopsis | omegaA | Spliceosome | Amino__acid__related__enzymes | 0.1724138 |
| Arabidopsis | omegaA | Spliceosome | Chaperones__and__folding__catalysts | 0.8620690 |
| Arabidopsis | omegaA | Spliceosome | Chromosome__and__associated__proteins | 0.9310345 |
| Arabidopsis | omegaA | Spliceosome | Cytochrome__P450 | 0.3103448 |
| Arabidopsis | omegaA | Spliceosome | Cytoskeleton__proteins | 0.9310345 |
| Arabidopsis | omegaA | Spliceosome | DNA__repair__and__recombination__proteins | 0.3793103 |
| Arabidopsis | omegaA | Spliceosome | DNA__replication__proteins | 0.7241379 |
| Arabidopsis | omegaA | Spliceosome | Exosome | 0.3793103 |
| Arabidopsis | omegaA | Spliceosome | G__protein__coupled__receptors | 0.1034483 |
| Arabidopsis | omegaA | Spliceosome | Glycosyltransferases | 0.9310345 |
| Arabidopsis | omegaA | Spliceosome | Ion__channels | 0.2413793 |
| Arabidopsis | omegaA | Spliceosome | Lipid__biosynthesis__proteins | 0.1034483 |
| Arabidopsis | omegaA | Spliceosome | Membrane__trafficking | 0.9310345 |
| Arabidopsis | omegaA | Spliceosome | Messenger__RNA__biogenesis | 0.8620690 |
| Arabidopsis | omegaA | Spliceosome | Mitochondrial__biogenesis | 0.2413793 |

*(continued)*

| species | estimate | var1 | var2 | p.value |
|---|---|---|---|---|
| Arabidopsis | omegaA | Spliceosome | Peptidases | 0.5862069 |
| Arabidopsis | omegaA | Spliceosome | Photosynthesis__proteins | 0.5172414 |
| Arabidopsis | omegaA | Spliceosome | Proteasome | 0.7241379 |
| Arabidopsis | omegaA | Spliceosome | Protein__kinases | 0.5862069 |
| Arabidopsis | omegaA | Spliceosome | Protein__phosphatases__and__associated__proteins | 0.1724138 |
| Arabidopsis | omegaA | Spliceosome | Ribosome | 0.4482759 |
| Arabidopsis | omegaA | Spliceosome | Ribosome__biogenesis | 0.1034483 |
| Arabidopsis | omegaA | Transcription__factors | Amino__acid__related__enzymes | 0.5862069 |
| Arabidopsis | omegaA | Transcription__factors | Chaperones__and__folding__catalysts | 0.5172414 |
| Arabidopsis | omegaA | Transcription__factors | Chromosome__and__associated__proteins | 0.7931034 |
| Arabidopsis | omegaA | Transcription__factors | Cytochrome__P450 | 0.1724138 |
| Arabidopsis | omegaA | Transcription__factors | Cytoskeleton__proteins | 0.7241379 |
| Arabidopsis | omegaA | Transcription__factors | DNA__repair__and__recombination__proteins | 0.7931034 |
| Arabidopsis | omegaA | Transcription__factors | DNA__replication__proteins | 0.7241379 |
| Arabidopsis | omegaA | Transcription__factors | Exosome | 0.9310345 |
| Arabidopsis | omegaA | Transcription__factors | G__protein__coupled__receptors | 0.0344828 |
| Arabidopsis | omegaA | Transcription__factors | Glycosyltransferases | 0.6551724 |
| Arabidopsis | omegaA | Transcription__factors | Ion__channels | 0.1034483 |
| Arabidopsis | omegaA | Transcription__factors | Lipid__biosynthesis__proteins | 0.0344828 |
| Arabidopsis | omegaA | Transcription__factors | Membrane__trafficking | 0.5172414 |
| Arabidopsis | omegaA | Transcription__factors | Messenger__RNA__biogenesis | 0.5862069 |
| Arabidopsis | omegaA | Transcription__factors | Mitochondrial__biogenesis | 0.5862069 |
| Arabidopsis | omegaA | Transcription__factors | Peptidases | 0.3103448 |
| Arabidopsis | omegaA | Transcription__factors | Photosynthesis__proteins | 0.7931034 |
| Arabidopsis | omegaA | Transcription__factors | Proteasome | 0.7241379 |
| Arabidopsis | omegaA | Transcription__factors | Protein__kinases | 0.6551724 |
| Arabidopsis | omegaA | Transcription__factors | Protein__phosphatases__and__associated__proteins | 0.1724138 |
| Arabidopsis | omegaA | Transcription__factors | Ribosome | 0.7931034 |
| Arabidopsis | omegaA | Transcription__factors | Ribosome__biogenesis | 0.1034483 |
| Arabidopsis | omegaA | Transcription__factors | Spliceosome | 0.5172414 |
| Arabidopsis | omegaA | Transcription__machinery | Amino__acid__related__enzymes | 0.0344828 |
| Arabidopsis | omegaA | Transcription__machinery | Chaperones__and__folding__catalysts | 0.1034483 |
| Arabidopsis | omegaA | Transcription__machinery | Chromosome__and__associated__proteins | 0.1034483 |
| Arabidopsis | omegaA | Transcription__machinery | Cytochrome__P450 | 0.5862069 |
| Arabidopsis | omegaA | Transcription__machinery | Cytoskeleton__proteins | 0.2413793 |
| Arabidopsis | omegaA | Transcription__machinery | DNA__repair__and__recombination__proteins | 0.0344828 |
| Arabidopsis | omegaA | Transcription__machinery | DNA__replication__proteins | 0.0344828 |
| Arabidopsis | omegaA | Transcription__machinery | Exosome | 0.1034483 |
| Arabidopsis | omegaA | Transcription__machinery | G__protein__coupled__receptors | 0.5862069 |
| Arabidopsis | omegaA | Transcription__machinery | Glycosyltransferases | 0.1724138 |
| Arabidopsis | omegaA | Transcription__machinery | Ion__channels | 0.3103448 |
| Arabidopsis | omegaA | Transcription__machinery | Lipid__biosynthesis__proteins | 0.4482759 |
| Arabidopsis | omegaA | Transcription__machinery | Membrane__trafficking | 0.1034483 |
| Arabidopsis | omegaA | Transcription__machinery | Messenger__RNA__biogenesis | 0.1034483 |
| Arabidopsis | omegaA | Transcription__machinery | Mitochondrial__biogenesis | 0.0344828 |
| Arabidopsis | omegaA | Transcription__machinery | Peptidases | 0.2413793 |
| Arabidopsis | omegaA | Transcription__machinery | Photosynthesis__proteins | 0.1034483 |
| Arabidopsis | omegaA | Transcription__machinery | Proteasome | 0.1724138 |
| Arabidopsis | omegaA | Transcription__machinery | Protein__kinases | 0.1034483 |
| Arabidopsis | omegaA | Transcription__machinery | Protein__phosphatases__and__associated__proteins | 0.7241379 |
| Arabidopsis | omegaA | Transcription__machinery | Ribosome | 0.1034483 |
| Arabidopsis | omegaA | Transcription__machinery | Ribosome__biogenesis | 0.9310345 |
| Arabidopsis | omegaA | Transcription__machinery | Spliceosome | 0.1034483 |
| Arabidopsis | omegaA | Transcription__machinery | Transcription__factors | 0.0344828 |
| Arabidopsis | omegaA | Transfer__RNA__biogenesis | Amino__acid__related__enzymes | 0.6551724 |
| Arabidopsis | omegaA | Transfer__RNA__biogenesis | Chaperones__and__folding__catalysts | 0.4482759 |
| Arabidopsis | omegaA | Transfer__RNA__biogenesis | Chromosome__and__associated__proteins | 0.7931034 |
| Arabidopsis | omegaA | Transfer__RNA__biogenesis | Cytochrome__P450 | 0.1724138 |
| Arabidopsis | omegaA | Transfer__RNA__biogenesis | Cytoskeleton__proteins | 0.7931034 |
| Arabidopsis | omegaA | Transfer__RNA__biogenesis | DNA__repair__and__recombination__proteins | 0.7931034 |
| Arabidopsis | omegaA | Transfer__RNA__biogenesis | DNA__replication__proteins | 0.6551724 |
| Arabidopsis | omegaA | Transfer__RNA__biogenesis | Exosome | 0.7931034 |

*(continued)*

| species | estimate | var1 | var2 | p.value |
|---|---|---|---|---|
| Arabidopsis | omegaA | Transfer__RNA__biogenesis | G__protein__coupled__receptors | 0.0344828 |
| Arabidopsis | omegaA | Transfer__RNA__biogenesis | Glycosyltransferases | 0.2413793 |
| Arabidopsis | omegaA | Transfer__RNA__biogenesis | Ion__channels | 0.0344828 |
| Arabidopsis | omegaA | Transfer__RNA__biogenesis | Lipid__biosynthesis__proteins | 0.1034483 |
| Arabidopsis | omegaA | Transfer__RNA__biogenesis | Membrane__trafficking | 0.5172414 |
| Arabidopsis | omegaA | Transfer__RNA__biogenesis | Messenger__RNA__biogenesis | 0.5172414 |
| Arabidopsis | omegaA | Transfer__RNA__biogenesis | Mitochondrial__biogenesis | 0.7241379 |
| Arabidopsis | omegaA | Transfer__RNA__biogenesis | Peptidases | 0.1724138 |
| Arabidopsis | omegaA | Transfer__RNA__biogenesis | Photosynthesis__proteins | 0.6551724 |
| Arabidopsis | omegaA | Transfer__RNA__biogenesis | Proteasome | 0.8620690 |
| Arabidopsis | omegaA | Transfer__RNA__biogenesis | Protein__kinases | 0.8620690 |
| Arabidopsis | omegaA | Transfer__RNA__biogenesis | Protein__phosphatases__and__associated__proteins | 0.1724138 |
| Arabidopsis | omegaA | Transfer__RNA__biogenesis | Ribosome | 0.7931034 |
| Arabidopsis | omegaA | Transfer__RNA__biogenesis | Ribosome__biogenesis | 0.0344828 |
| Arabidopsis | omegaA | Transfer__RNA__biogenesis | Spliceosome | 0.5862069 |
| Arabidopsis | omegaA | Transfer__RNA__biogenesis | Transcription__factors | 0.7931034 |
| Arabidopsis | omegaA | Transfer__RNA__biogenesis | Transcription__machinery | 0.1034483 |
| Arabidopsis | omegaA | Translation__factors | Amino__acid__related__enzymes | 0.1034483 |
| Arabidopsis | omegaA | Translation__factors | Chaperones__and__folding__catalysts | 0.3793103 |
| Arabidopsis | omegaA | Translation__factors | Chromosome__and__associated__proteins | 0.1724138 |
| Arabidopsis | omegaA | Translation__factors | Cytochrome__P450 | 0.7931034 |
| Arabidopsis | omegaA | Translation__factors | Cytoskeleton__proteins | 0.6551724 |
| Arabidopsis | omegaA | Translation__factors | DNA__repair__and__recombination__proteins | 0.1034483 |
| Arabidopsis | omegaA | Translation__factors | DNA__replication__proteins | 0.2413793 |
| Arabidopsis | omegaA | Translation__factors | Exosome | 0.1034483 |
| Arabidopsis | omegaA | Translation__factors | G__protein__coupled__receptors | 0.7931034 |
| Arabidopsis | omegaA | Translation__factors | Glycosyltransferases | 0.2413793 |
| Arabidopsis | omegaA | Translation__factors | Ion__channels | 0.8620690 |
| Arabidopsis | omegaA | Translation__factors | Lipid__biosynthesis__proteins | 0.8620690 |
| Arabidopsis | omegaA | Translation__factors | Membrane__trafficking | 0.4482759 |
| Arabidopsis | omegaA | Translation__factors | Messenger__RNA__biogenesis | 0.5862069 |
| Arabidopsis | omegaA | Translation__factors | Mitochondrial__biogenesis | 0.0344828 |
| Arabidopsis | omegaA | Translation__factors | Peptidases | 0.5862069 |
| Arabidopsis | omegaA | Translation__factors | Photosynthesis__proteins | 0.2413793 |
| Arabidopsis | omegaA | Translation__factors | Proteasome | 0.3793103 |
| Arabidopsis | omegaA | Translation__factors | Protein__kinases | 0.1724138 |
| Arabidopsis | omegaA | Translation__factors | Protein__phosphatases__and__associated__proteins | 0.7931034 |
| Arabidopsis | omegaA | Translation__factors | Ribosome | 0.1034483 |
| Arabidopsis | omegaA | Translation__factors | Ribosome__biogenesis | 0.5172414 |
| Arabidopsis | omegaA | Translation__factors | Spliceosome | 0.5172414 |
| Arabidopsis | omegaA | Translation__factors | Transcription__factors | 0.1724138 |
| Arabidopsis | omegaA | Translation__factors | Transcription__machinery | 0.3793103 |
| Arabidopsis | omegaA | Translation__factors | Transfer__RNA__biogenesis | 0.1034483 |
| Arabidopsis | omegaA | Transporters | Amino__acid__related__enzymes | 0.1034483 |
| Arabidopsis | omegaA | Transporters | Chaperones__and__folding__catalysts | 0.8620690 |
| Arabidopsis | omegaA | Transporters | Chromosome__and__associated__proteins | 0.4482759 |
| Arabidopsis | omegaA | Transporters | Cytochrome__P450 | 0.3103448 |
| Arabidopsis | omegaA | Transporters | Cytoskeleton__proteins | 0.8620690 |
| Arabidopsis | omegaA | Transporters | DNA__repair__and__recombination__proteins | 0.1034483 |
| Arabidopsis | omegaA | Transporters | DNA__replication__proteins | 0.5172414 |
| Arabidopsis | omegaA | Transporters | Exosome | 0.1724138 |
| Arabidopsis | omegaA | Transporters | G__protein__coupled__receptors | 0.1724138 |
| Arabidopsis | omegaA | Transporters | Glycosyltransferases | 0.7241379 |
| Arabidopsis | omegaA | Transporters | Ion__channels | 0.2413793 |
| Arabidopsis | omegaA | Transporters | Lipid__biosynthesis__proteins | 0.2413793 |
| Arabidopsis | omegaA | Transporters | Membrane__trafficking | 0.4482759 |
| Arabidopsis | omegaA | Transporters | Messenger__RNA__biogenesis | 0.7931034 |
| Arabidopsis | omegaA | Transporters | Mitochondrial__biogenesis | 0.0344828 |
| Arabidopsis | omegaA | Transporters | Peptidases | 0.8620690 |
| Arabidopsis | omegaA | Transporters | Photosynthesis__proteins | 0.3793103 |
| Arabidopsis | omegaA | Transporters | Proteasome | 0.8620690 |

*(continued)*

| species | estimate | var1 | var2 | p.value |
|---|---|---|---|---|
| Arabidopsis | omegaA | Transporters | Protein_kinases | 0.3103448 |
| Arabidopsis | omegaA | Transporters | Protein_phosphatases_and_associated_proteins | 0.3103448 |
| Arabidopsis | omegaA | Transporters | Ribosome | 0.4482759 |
| Arabidopsis | omegaA | Transporters | Ribosome_biogenesis | 0.1034483 |
| Arabidopsis | omegaA | Transporters | Spliceosome | 0.7931034 |
| Arabidopsis | omegaA | Transporters | Transcription_factors | 0.3793103 |
| Arabidopsis | omegaA | Transporters | Transcription_machinery | 0.1034483 |
| Arabidopsis | omegaA | Transporters | Transfer_RNA_biogenesis | 0.1724138 |
| Arabidopsis | omegaA | Transporters | Translation_factors | 0.4482759 |
| Arabidopsis | omegaA | Ubiquitin_system | Amino_acid_related_enzymes | 0.0344828 |
| Arabidopsis | omegaA | Ubiquitin_system | Chaperones_and_folding_catalysts | 0.1724138 |
| Arabidopsis | omegaA | Ubiquitin_system | Chromosome_and_associated_proteins | 0.0344828 |
| Arabidopsis | omegaA | Ubiquitin_system | Cytochrome_P450 | 0.7931034 |
| Arabidopsis | omegaA | Ubiquitin_system | Cytoskeleton_proteins | 0.3103448 |
| Arabidopsis | omegaA | Ubiquitin_system | DNA_repair_and_recombination_proteins | 0.0344828 |
| Arabidopsis | omegaA | Ubiquitin_system | DNA_replication_proteins | 0.1034483 |
| Arabidopsis | omegaA | Ubiquitin_system | Exosome | 0.0344828 |
| Arabidopsis | omegaA | Ubiquitin_system | G_protein_coupled_receptors | 0.6551724 |
| Arabidopsis | omegaA | Ubiquitin_system | Glycosyltransferases | 0.0344828 |
| Arabidopsis | omegaA | Ubiquitin_system | Ion_channels | 0.3103448 |
| Arabidopsis | omegaA | Ubiquitin_system | Lipid_biosynthesis_proteins | 0.5172414 |
| Arabidopsis | omegaA | Ubiquitin_system | Membrane_trafficking | 0.1034483 |
| Arabidopsis | omegaA | Ubiquitin_system | Messenger_RNA_biogenesis | 0.1034483 |
| Arabidopsis | omegaA | Ubiquitin_system | Mitochondrial_biogenesis | 0.0344828 |
| Arabidopsis | omegaA | Ubiquitin_system | Peptidases | 0.1724138 |
| Arabidopsis | omegaA | Ubiquitin_system | Photosynthesis_proteins | 0.0344828 |
| Arabidopsis | omegaA | Ubiquitin_system | Proteasome | 0.1034483 |
| Arabidopsis | omegaA | Ubiquitin_system | Protein_kinases | 0.0344828 |
| Arabidopsis | omegaA | Ubiquitin_system | Protein_phosphatases_and_associated_proteins | 0.5862069 |
| Arabidopsis | omegaA | Ubiquitin_system | Ribosome | 0.0344828 |
| Arabidopsis | omegaA | Ubiquitin_system | Ribosome_biogenesis | 1.0000000 |
| Arabidopsis | omegaA | Ubiquitin_system | Spliceosome | 0.1034483 |
| Arabidopsis | omegaA | Ubiquitin_system | Transcription_factors | 0.0344828 |
| Arabidopsis | omegaA | Ubiquitin_system | Transcription_machinery | 0.7241379 |
| Arabidopsis | omegaA | Ubiquitin_system | Transfer_RNA_biogenesis | 0.1034483 |
| Arabidopsis | omegaA | Ubiquitin_system | Translation_factors | 0.3103448 |
| Arabidopsis | omegaA | Ubiquitin_system | Transporters | 0.1034483 |

```r
## Drosophila
dmel.hist <- subset(kegg.hist, kegg.hist$species == "Drosophila")
nboots <- 100
dmel.smallkegg.p <- list()
for (i in 1:nboots) {
  dmel.smallkegg.p[[i]] <- ddply(dmel.hist, c("species", "estimate",
                                              "var1", "var2"),
                                 function(x, N=53){
    c <- as.numeric(nrow(x[x$value < 0,]))
    c2 <- as.numeric(nrow(x[x$value > 0,]))
    m <- min(c, c2)
    p <- (2*m+1)/(N+1)
    tbl <- data.frame(m, p)
  }
)}

# correcting the p-value for multiple testing
dmel.kegg.p.adj <- lapply(dmel.smallkegg.p, function(x) {
  ddply(x,  c("species","estimate", "var1", "var2"), function(x) {
```

```
    p.value <- p.adjust(x$p)
    data.frame(p.value)
  })
})

# taking the minimum p-value of the replicates performed
tbl.dmel.p.adj <- rbindlist(dmel.kegg.p.adj)
dmel.pvalue <- ddply(tbl.dmel.p.adj, c("species","estimate", "var1", "var2"),
                     function(x) {
  p.value <- min(x$p.value)
  data.frame(p.value)
})

# showing the table
kable(dmel.pvalue, format = "latex", booktabs = TRUE, longtable = TRUE) %>%
  kable_styling(latex_options = c("hold_position", "repeat_header"),
                font_size = 7)
```

| species | estimate | var1 | var2 | p.value |
|---|---|---|---|---|
| Drosophila | dnds | Chaperones_and_folding_catalysts | Amino_acid_related_enzymes | 0.4629630 |
| Drosophila | dnds | Chromosome_and_associated_proteins | Amino_acid_related_enzymes | 0.1296296 |
| Drosophila | dnds | Chromosome_and_associated_proteins | Chaperones_and_folding_catalysts | 0.1666667 |
| Drosophila | dnds | Cytochrome_P450 | Amino_acid_related_enzymes | 0.7592593 |
| Drosophila | dnds | Cytochrome_P450 | Chaperones_and_folding_catalysts | 0.9444444 |
| Drosophila | dnds | Cytochrome_P450 | Chromosome_and_associated_proteins | 0.6111111 |
| Drosophila | dnds | Cytoskeleton_proteins | Amino_acid_related_enzymes | 0.7592593 |
| Drosophila | dnds | Cytoskeleton_proteins | Chaperones_and_folding_catalysts | 0.3518519 |
| Drosophila | dnds | Cytoskeleton_proteins | Chromosome_and_associated_proteins | 0.0185185 |
| Drosophila | dnds | Cytoskeleton_proteins | Cytochrome_P450 | 0.6111111 |
| Drosophila | dnds | DNA_repair_and_recombination_proteins | Amino_acid_related_enzymes | 0.0555556 |
| Drosophila | dnds | DNA_repair_and_recombination_proteins | Chaperones_and_folding_catalysts | 0.0185185 |
| Drosophila | dnds | DNA_repair_and_recombination_proteins | Chromosome_and_associated_proteins | 0.0555556 |
| Drosophila | dnds | DNA_repair_and_recombination_proteins | Cytochrome_P450 | 0.1666667 |
| Drosophila | dnds | DNA_repair_and_recombination_proteins | Cytoskeleton_proteins | 0.0185185 |
| Drosophila | dnds | DNA_replication_proteins | Amino_acid_related_enzymes | 0.4259259 |
| Drosophila | dnds | DNA_replication_proteins | Chaperones_and_folding_catalysts | 0.2037037 |
| Drosophila | dnds | DNA_replication_proteins | Chromosome_and_associated_proteins | 0.0185185 |
| Drosophila | dnds | DNA_replication_proteins | Cytochrome_P450 | 0.6111111 |
| Drosophila | dnds | DNA_replication_proteins | Cytoskeleton_proteins | 0.9074074 |
| Drosophila | dnds | DNA_replication_proteins | DNA_repair_and_recombination_proteins | 0.0185185 |
| Drosophila | dnds | Exosome | Amino_acid_related_enzymes | 0.5000000 |
| Drosophila | dnds | Exosome | Chaperones_and_folding_catalysts | 0.2037037 |
| Drosophila | dnds | Exosome | Chromosome_and_associated_proteins | 0.0185185 |
| Drosophila | dnds | Exosome | Cytochrome_P450 | 0.5370370 |
| Drosophila | dnds | Exosome | Cytoskeleton_proteins | 0.9074074 |
| Drosophila | dnds | Exosome | DNA_repair_and_recombination_proteins | 0.0185185 |
| Drosophila | dnds | Exosome | DNA_replication_proteins | 0.8703704 |
| Drosophila | dnds | G_protein_coupled_receptors | Amino_acid_related_enzymes | 0.0185185 |
| Drosophila | dnds | G_protein_coupled_receptors | Chaperones_and_folding_catalysts | 0.0185185 |
| Drosophila | dnds | G_protein_coupled_receptors | Chromosome_and_associated_proteins | 0.0185185 |
| Drosophila | dnds | G_protein_coupled_receptors | Cytochrome_P450 | 0.0925926 |
| Drosophila | dnds | G_protein_coupled_receptors | Cytoskeleton_proteins | 0.1666667 |
| Drosophila | dnds | G_protein_coupled_receptors | DNA_repair_and_recombination_proteins | 0.0185185 |
| Drosophila | dnds | G_protein_coupled_receptors | DNA_replication_proteins | 0.0555556 |
| Drosophila | dnds | G_protein_coupled_receptors | Exosome | 0.0925926 |
| Drosophila | dnds | Glycosyltransferases | Amino_acid_related_enzymes | 0.7962963 |
| Drosophila | dnds | Glycosyltransferases | Chaperones_and_folding_catalysts | 0.4629630 |
| Drosophila | dnds | Glycosyltransferases | Chromosome_and_associated_proteins | 0.0185185 |
| Drosophila | dnds | Glycosyltransferases | Cytochrome_P450 | 0.7592593 |
| Drosophila | dnds | Glycosyltransferases | Cytoskeleton_proteins | 0.6111111 |

*(continued)*

| species | estimate | var1 | var2 | p.value |
|---|---|---|---|---|
| Drosophila | dnds | Glycosyltransferases | DNA__repair__and__recombination__proteins | 0.0185185 |
| Drosophila | dnds | Glycosyltransferases | DNA__replication__proteins | 0.2037037 |
| Drosophila | dnds | Glycosyltransferases | Exosome | 0.2037037 |
| Drosophila | dnds | Glycosyltransferases | G__protein__coupled__receptors | 0.0555556 |
| Drosophila | dnds | Ion__channels | Amino__acid__related__enzymes | 0.0185185 |
| Drosophila | dnds | Ion__channels | Chaperones__and__folding__catalysts | 0.0185185 |
| Drosophila | dnds | Ion__channels | Chromosome__and__associated__proteins | 0.0185185 |
| Drosophila | dnds | Ion__channels | Cytochrome__P450 | 0.0185185 |
| Drosophila | dnds | Ion__channels | Cytoskeleton__proteins | 0.0185185 |
| Drosophila | dnds | Ion__channels | DNA__repair__and__recombination__proteins | 0.0185185 |
| Drosophila | dnds | Ion__channels | DNA__replication__proteins | 0.0185185 |
| Drosophila | dnds | Ion__channels | Exosome | 0.0185185 |
| Drosophila | dnds | Ion__channels | G__protein__coupled__receptors | 0.1296296 |
| Drosophila | dnds | Ion__channels | Glycosyltransferases | 0.0185185 |
| Drosophila | dnds | Lipid__biosynthesis__proteins | Amino__acid__related__enzymes | 0.7962963 |
| Drosophila | dnds | Lipid__biosynthesis__proteins | Chaperones__and__folding__catalysts | 0.7222222 |
| Drosophila | dnds | Lipid__biosynthesis__proteins | Chromosome__and__associated__proteins | 0.2037037 |
| Drosophila | dnds | Lipid__biosynthesis__proteins | Cytochrome__P450 | 0.6111111 |
| Drosophila | dnds | Lipid__biosynthesis__proteins | Cytoskeleton__proteins | 0.7592593 |
| Drosophila | dnds | Lipid__biosynthesis__proteins | DNA__repair__and__recombination__proteins | 0.0185185 |
| Drosophila | dnds | Lipid__biosynthesis__proteins | DNA__replication__proteins | 0.7222222 |
| Drosophila | dnds | Lipid__biosynthesis__proteins | Exosome | 0.6111111 |
| Drosophila | dnds | Lipid__biosynthesis__proteins | G__protein__coupled__receptors | 0.1296296 |
| Drosophila | dnds | Lipid__biosynthesis__proteins | Glycosyltransferases | 0.9074074 |
| Drosophila | dnds | Lipid__biosynthesis__proteins | Ion__channels | 0.0185185 |
| Drosophila | dnds | Membrane__trafficking | Amino__acid__related__enzymes | 0.0185185 |
| Drosophila | dnds | Membrane__trafficking | Chaperones__and__folding__catalysts | 0.0185185 |
| Drosophila | dnds | Membrane__trafficking | Chromosome__and__associated__proteins | 0.0185185 |
| Drosophila | dnds | Membrane__trafficking | Cytochrome__P450 | 0.2407407 |
| Drosophila | dnds | Membrane__trafficking | Cytoskeleton__proteins | 0.1296296 |
| Drosophila | dnds | Membrane__trafficking | DNA__repair__and__recombination__proteins | 0.0185185 |
| Drosophila | dnds | Membrane__trafficking | DNA__replication__proteins | 0.0185185 |
| Drosophila | dnds | Membrane__trafficking | Exosome | 0.0185185 |
| Drosophila | dnds | Membrane__trafficking | G__protein__coupled__receptors | 0.2037037 |
| Drosophila | dnds | Membrane__trafficking | Glycosyltransferases | 0.0185185 |
| Drosophila | dnds | Membrane__trafficking | Ion__channels | 0.0185185 |
| Drosophila | dnds | Membrane__trafficking | Lipid__biosynthesis__proteins | 0.2777778 |
| Drosophila | dnds | Messenger__RNA__biogenesis | Amino__acid__related__enzymes | 0.0555556 |
| Drosophila | dnds | Messenger__RNA__biogenesis | Chaperones__and__folding__catalysts | 0.0185185 |
| Drosophila | dnds | Messenger__RNA__biogenesis | Chromosome__and__associated__proteins | 0.0185185 |
| Drosophila | dnds | Messenger__RNA__biogenesis | Cytochrome__P450 | 0.0185185 |
| Drosophila | dnds | Messenger__RNA__biogenesis | Cytoskeleton__proteins | 0.0185185 |
| Drosophila | dnds | Messenger__RNA__biogenesis | DNA__repair__and__recombination__proteins | 0.3888889 |
| Drosophila | dnds | Messenger__RNA__biogenesis | DNA__replication__proteins | 0.0185185 |
| Drosophila | dnds | Messenger__RNA__biogenesis | Exosome | 0.0185185 |
| Drosophila | dnds | Messenger__RNA__biogenesis | G__protein__coupled__receptors | 0.0185185 |
| Drosophila | dnds | Messenger__RNA__biogenesis | Glycosyltransferases | 0.0185185 |
| Drosophila | dnds | Messenger__RNA__biogenesis | Ion__channels | 0.0185185 |
| Drosophila | dnds | Messenger__RNA__biogenesis | Lipid__biosynthesis__proteins | 0.0185185 |
| Drosophila | dnds | Messenger__RNA__biogenesis | Membrane__trafficking | 0.0185185 |
| Drosophila | dnds | Mitochondrial__biogenesis | Amino__acid__related__enzymes | 0.9814815 |
| Drosophila | dnds | Mitochondrial__biogenesis | Chaperones__and__folding__catalysts | 0.2777778 |
| Drosophila | dnds | Mitochondrial__biogenesis | Chromosome__and__associated__proteins | 0.0185185 |
| Drosophila | dnds | Mitochondrial__biogenesis | Cytochrome__P450 | 0.7222222 |
| Drosophila | dnds | Mitochondrial__biogenesis | Cytoskeleton__proteins | 0.7222222 |
| Drosophila | dnds | Mitochondrial__biogenesis | DNA__repair__and__recombination__proteins | 0.0185185 |
| Drosophila | dnds | Mitochondrial__biogenesis | DNA__replication__proteins | 0.3518519 |
| Drosophila | dnds | Mitochondrial__biogenesis | Exosome | 0.3148148 |
| Drosophila | dnds | Mitochondrial__biogenesis | G__protein__coupled__receptors | 0.0185185 |
| Drosophila | dnds | Mitochondrial__biogenesis | Glycosyltransferases | 0.5740741 |
| Drosophila | dnds | Mitochondrial__biogenesis | Ion__channels | 0.0185185 |
| Drosophila | dnds | Mitochondrial__biogenesis | Lipid__biosynthesis__proteins | 0.8333333 |

(continued)

| species | estimate | var1 | var2 | p.value |
|---|---|---|---|---|
| Drosophila | dnds | Mitochondrial__biogenesis | Membrane__trafficking | 0.0185185 |
| Drosophila | dnds | Mitochondrial__biogenesis | Messenger__RNA__biogenesis | 0.0185185 |
| Drosophila | dnds | Peptidases | Amino__acid__related__enzymes | 0.1296296 |
| Drosophila | dnds | Peptidases | Chaperones__and__folding__catalysts | 0.2777778 |
| Drosophila | dnds | Peptidases | Chromosome__and__associated__proteins | 0.9814815 |
| Drosophila | dnds | Peptidases | Cytochrome__P450 | 0.6851852 |
| Drosophila | dnds | Peptidases | Cytoskeleton__proteins | 0.0555556 |
| Drosophila | dnds | Peptidases | DNA__repair__and__recombination__proteins | 0.2037037 |
| Drosophila | dnds | Peptidases | DNA__replication__proteins | 0.0185185 |
| Drosophila | dnds | Peptidases | Exosome | 0.0185185 |
| Drosophila | dnds | Peptidases | G__protein__coupled__receptors | 0.0185185 |
| Drosophila | dnds | Peptidases | Glycosyltransferases | 0.0185185 |
| Drosophila | dnds | Peptidases | Ion__channels | 0.0185185 |
| Drosophila | dnds | Peptidases | Lipid__biosynthesis__proteins | 0.3148148 |
| Drosophila | dnds | Peptidases | Membrane__trafficking | 0.0185185 |
| Drosophila | dnds | Peptidases | Messenger__RNA__biogenesis | 0.0555556 |
| Drosophila | dnds | Peptidases | Mitochondrial__biogenesis | 0.0185185 |
| Drosophila | dnds | Photosynthesis__proteins | Amino__acid__related__enzymes | 0.2777778 |
| Drosophila | dnds | Photosynthesis__proteins | Chaperones__and__folding__catalysts | 0.1296296 |
| Drosophila | dnds | Photosynthesis__proteins | Chromosome__and__associated__proteins | 0.0555556 |
| Drosophila | dnds | Photosynthesis__proteins | Cytochrome__P450 | 0.2777778 |
| Drosophila | dnds | Photosynthesis__proteins | Cytoskeleton__proteins | 0.5740741 |
| Drosophila | dnds | Photosynthesis__proteins | DNA__repair__and__recombination__proteins | 0.0185185 |
| Drosophila | dnds | Photosynthesis__proteins | DNA__replication__proteins | 0.3888889 |
| Drosophila | dnds | Photosynthesis__proteins | Exosome | 0.5740741 |
| Drosophila | dnds | Photosynthesis__proteins | G__protein__coupled__receptors | 0.2777778 |
| Drosophila | dnds | Photosynthesis__proteins | Glycosyltransferases | 0.1296296 |
| Drosophila | dnds | Photosynthesis__proteins | Ion__channels | 0.0185185 |
| Drosophila | dnds | Photosynthesis__proteins | Lipid__biosynthesis__proteins | 0.4259259 |
| Drosophila | dnds | Photosynthesis__proteins | Membrane__trafficking | 0.4259259 |
| Drosophila | dnds | Photosynthesis__proteins | Messenger__RNA__biogenesis | 0.0185185 |
| Drosophila | dnds | Photosynthesis__proteins | Mitochondrial__biogenesis | 0.1666667 |
| Drosophila | dnds | Photosynthesis__proteins | Peptidases | 0.0555556 |
| Drosophila | dnds | Proteasome | Amino__acid__related__enzymes | 0.3148148 |
| Drosophila | dnds | Proteasome | Chaperones__and__folding__catalysts | 0.1296296 |
| Drosophila | dnds | Proteasome | Chromosome__and__associated__proteins | 0.0185185 |
| Drosophila | dnds | Proteasome | Cytochrome__P450 | 0.3888889 |
| Drosophila | dnds | Proteasome | Cytoskeleton__proteins | 0.7222222 |
| Drosophila | dnds | Proteasome | DNA__repair__and__recombination__proteins | 0.0185185 |
| Drosophila | dnds | Proteasome | DNA__replication__proteins | 0.6111111 |
| Drosophila | dnds | Proteasome | Exosome | 0.6851852 |
| Drosophila | dnds | Proteasome | G__protein__coupled__receptors | 0.3888889 |
| Drosophila | dnds | Proteasome | Glycosyltransferases | 0.2407407 |
| Drosophila | dnds | Proteasome | Ion__channels | 0.0185185 |
| Drosophila | dnds | Proteasome | Lipid__biosynthesis__proteins | 0.3518519 |
| Drosophila | dnds | Proteasome | Membrane__trafficking | 0.6851852 |
| Drosophila | dnds | Proteasome | Messenger__RNA__biogenesis | 0.0185185 |
| Drosophila | dnds | Proteasome | Mitochondrial__biogenesis | 0.2777778 |
| Drosophila | dnds | Proteasome | Peptidases | 0.0555556 |
| Drosophila | dnds | Proteasome | Photosynthesis__proteins | 0.9444444 |
| Drosophila | dnds | Protein__kinases | Amino__acid__related__enzymes | 0.1666667 |
| Drosophila | dnds | Protein__kinases | Chaperones__and__folding__catalysts | 0.0925926 |
| Drosophila | dnds | Protein__kinases | Chromosome__and__associated__proteins | 0.0185185 |
| Drosophila | dnds | Protein__kinases | Cytochrome__P450 | 0.3888889 |
| Drosophila | dnds | Protein__kinases | Cytoskeleton__proteins | 0.5740741 |
| Drosophila | dnds | Protein__kinases | DNA__repair__and__recombination__proteins | 0.0185185 |
| Drosophila | dnds | Protein__kinases | DNA__replication__proteins | 0.2037037 |
| Drosophila | dnds | Protein__kinases | Exosome | 0.3148148 |
| Drosophila | dnds | Protein__kinases | G__protein__coupled__receptors | 0.0555556 |
| Drosophila | dnds | Protein__kinases | Glycosyltransferases | 0.0185185 |
| Drosophila | dnds | Protein__kinases | Ion__channels | 0.0185185 |
| Drosophila | dnds | Protein__kinases | Lipid__biosynthesis__proteins | 0.3888889 |

*(continued)*

| species | estimate | var1 | var2 | p.value |
|---|---|---|---|---|
| Drosophila | dnds | Protein__kinases | Membrane__trafficking | 0.0555556 |
| Drosophila | dnds | Protein__kinases | Messenger__RNA__biogenesis | 0.0185185 |
| Drosophila | dnds | Protein__kinases | Mitochondrial__biogenesis | 0.0555556 |
| Drosophila | dnds | Protein__kinases | Peptidases | 0.0185185 |
| Drosophila | dnds | Protein__kinases | Photosynthesis__proteins | 0.9074074 |
| Drosophila | dnds | Protein__kinases | Proteasome | 0.9074074 |
| Drosophila | dnds | Protein__phosphatases__and__associated__proteins | Amino__acid__related__enzymes | 0.2407407 |
| Drosophila | dnds | Protein__phosphatases__and__associated__proteins | Chaperones__and__folding__catalysts | 0.4629630 |
| Drosophila | dnds | Protein__phosphatases__and__associated__proteins | Chromosome__and__associated__proteins | 0.4259259 |
| Drosophila | dnds | Protein__phosphatases__and__associated__proteins | Cytochrome__P450 | 0.9444444 |
| Drosophila | dnds | Protein__phosphatases__and__associated__proteins | Cytoskeleton__proteins | 0.0555556 |
| Drosophila | dnds | Protein__phosphatases__and__associated__proteins | DNA__repair__and__recombination__proteins | 0.1296296 |
| Drosophila | dnds | Protein__phosphatases__and__associated__proteins | DNA__replication__proteins | 0.0185185 |
| Drosophila | dnds | Protein__phosphatases__and__associated__proteins | Exosome | 0.0185185 |
| Drosophila | dnds | Protein__phosphatases__and__associated__proteins | G__protein__coupled__receptors | 0.0185185 |
| Drosophila | dnds | Protein__phosphatases__and__associated__proteins | Glycosyltransferases | 0.0185185 |
| Drosophila | dnds | Protein__phosphatases__and__associated__proteins | Ion__channels | 0.0185185 |
| Drosophila | dnds | Protein__phosphatases__and__associated__proteins | Lipid__biosynthesis__proteins | 0.4259259 |
| Drosophila | dnds | Protein__phosphatases__and__associated__proteins | Membrane__trafficking | 0.0185185 |
| Drosophila | dnds | Protein__phosphatases__and__associated__proteins | Messenger__RNA__biogenesis | 0.0185185 |
| Drosophila | dnds | Protein__phosphatases__and__associated__proteins | Mitochondrial__biogenesis | 0.0185185 |
| Drosophila | dnds | Protein__phosphatases__and__associated__proteins | Peptidases | 0.5000000 |
| Drosophila | dnds | Protein__phosphatases__and__associated__proteins | Photosynthesis__proteins | 0.0555556 |
| Drosophila | dnds | Protein__phosphatases__and__associated__proteins | Proteasome | 0.0555556 |
| Drosophila | dnds | Protein__phosphatases__and__associated__proteins | Protein__kinases | 0.0185185 |
| Drosophila | dnds | Ribosome | Amino__acid__related__enzymes | 0.6481481 |
| Drosophila | dnds | Ribosome | Chaperones__and__folding__catalysts | 0.6851852 |
| Drosophila | dnds | Ribosome | Chromosome__and__associated__proteins | 0.0185185 |
| Drosophila | dnds | Ribosome | Cytochrome__P450 | 0.8333333 |
| Drosophila | dnds | Ribosome | Cytoskeleton__proteins | 0.3888889 |
| Drosophila | dnds | Ribosome | DNA__repair__and__recombination__proteins | 0.0185185 |
| Drosophila | dnds | Ribosome | DNA__replication__proteins | 0.0555556 |
| Drosophila | dnds | Ribosome | Exosome | 0.0555556 |
| Drosophila | dnds | Ribosome | G__protein__coupled__receptors | 0.0555556 |
| Drosophila | dnds | Ribosome | Glycosyltransferases | 0.5370370 |
| Drosophila | dnds | Ribosome | Ion__channels | 0.0185185 |
| Drosophila | dnds | Ribosome | Lipid__biosynthesis__proteins | 0.9444444 |
| Drosophila | dnds | Ribosome | Membrane__trafficking | 0.0185185 |
| Drosophila | dnds | Ribosome | Messenger__RNA__biogenesis | 0.0185185 |
| Drosophila | dnds | Ribosome | Mitochondrial__biogenesis | 0.2407407 |
| Drosophila | dnds | Ribosome | Peptidases | 0.0925926 |
| Drosophila | dnds | Ribosome | Photosynthesis__proteins | 0.1296296 |
| Drosophila | dnds | Ribosome | Proteasome | 0.1296296 |
| Drosophila | dnds | Ribosome | Protein__kinases | 0.0185185 |
| Drosophila | dnds | Ribosome | Protein__phosphatases__and__associated__proteins | 0.0925926 |
| Drosophila | dnds | Ribosome__biogenesis | Amino__acid__related__enzymes | 0.1296296 |
| Drosophila | dnds | Ribosome__biogenesis | Chaperones__and__folding__catalysts | 0.1666667 |
| Drosophila | dnds | Ribosome__biogenesis | Chromosome__and__associated__proteins | 0.6851852 |
| Drosophila | dnds | Ribosome__biogenesis | Cytochrome__P450 | 0.7222222 |
| Drosophila | dnds | Ribosome__biogenesis | Cytoskeleton__proteins | 0.0185185 |
| Drosophila | dnds | Ribosome__biogenesis | DNA__repair__and__recombination__proteins | 0.0925926 |
| Drosophila | dnds | Ribosome__biogenesis | DNA__replication__proteins | 0.0185185 |
| Drosophila | dnds | Ribosome__biogenesis | Exosome | 0.0185185 |
| Drosophila | dnds | Ribosome__biogenesis | G__protein__coupled__receptors | 0.0185185 |
| Drosophila | dnds | Ribosome__biogenesis | Glycosyltransferases | 0.0185185 |
| Drosophila | dnds | Ribosome__biogenesis | Ion__channels | 0.0185185 |
| Drosophila | dnds | Ribosome__biogenesis | Lipid__biosynthesis__proteins | 0.3518519 |
| Drosophila | dnds | Ribosome__biogenesis | Membrane__trafficking | 0.0185185 |
| Drosophila | dnds | Ribosome__biogenesis | Messenger__RNA__biogenesis | 0.0185185 |
| Drosophila | dnds | Ribosome__biogenesis | Mitochondrial__biogenesis | 0.0185185 |
| Drosophila | dnds | Ribosome__biogenesis | Peptidases | 0.9074074 |

(continued)

| species | estimate | var1 | var2 | p.value |
|---|---|---|---|---|
| Drosophila | dnds | Ribosome__biogenesis | Photosynthesis__proteins | 0.0185185 |
| Drosophila | dnds | Ribosome__biogenesis | Proteasome | 0.0555556 |
| Drosophila | dnds | Ribosome__biogenesis | Protein__kinases | 0.0185185 |
| Drosophila | dnds | Ribosome__biogenesis | Protein__phosphatases__and__associated__proteins | 0.7592593 |
| Drosophila | dnds | Ribosome__biogenesis | Ribosome | 0.0925926 |
| Drosophila | dnds | Spliceosome | Amino__acid__related__enzymes | 0.2407407 |
| Drosophila | dnds | Spliceosome | Chaperones__and__folding__catalysts | 0.5740741 |
| Drosophila | dnds | Spliceosome | Chromosome__and__associated__proteins | 0.4629630 |
| Drosophila | dnds | Spliceosome | Cytochrome__P450 | 0.9814815 |
| Drosophila | dnds | Spliceosome | Cytoskeleton__proteins | 0.2037037 |
| Drosophila | dnds | Spliceosome | DNA__repair__and__recombination__proteins | 0.0555556 |
| Drosophila | dnds | Spliceosome | DNA__replication__proteins | 0.0185185 |
| Drosophila | dnds | Spliceosome | Exosome | 0.0185185 |
| Drosophila | dnds | Spliceosome | G__protein__coupled__receptors | 0.0185185 |
| Drosophila | dnds | Spliceosome | Glycosyltransferases | 0.1296296 |
| Drosophila | dnds | Spliceosome | Ion__channels | 0.0185185 |
| Drosophila | dnds | Spliceosome | Lipid__biosynthesis__proteins | 0.6111111 |
| Drosophila | dnds | Spliceosome | Membrane__trafficking | 0.0185185 |
| Drosophila | dnds | Spliceosome | Messenger__RNA__biogenesis | 0.0555556 |
| Drosophila | dnds | Spliceosome | Mitochondrial__biogenesis | 0.0185185 |
| Drosophila | dnds | Spliceosome | Peptidases | 0.5000000 |
| Drosophila | dnds | Spliceosome | Photosynthesis__proteins | 0.0555556 |
| Drosophila | dnds | Spliceosome | Proteasome | 0.0555556 |
| Drosophila | dnds | Spliceosome | Protein__kinases | 0.0185185 |
| Drosophila | dnds | Spliceosome | Protein__phosphatases__and__associated__proteins | 0.7962963 |
| Drosophila | dnds | Spliceosome | Ribosome | 0.2407407 |
| Drosophila | dnds | Spliceosome | Ribosome__biogenesis | 0.4629630 |
| Drosophila | dnds | Transcription__factors | Amino__acid__related__enzymes | 0.1666667 |
| Drosophila | dnds | Transcription__factors | Chaperones__and__folding__catalysts | 0.5000000 |
| Drosophila | dnds | Transcription__factors | Chromosome__and__associated__proteins | 0.1296296 |
| Drosophila | dnds | Transcription__factors | Cytochrome__P450 | 0.9814815 |
| Drosophila | dnds | Transcription__factors | Cytoskeleton__proteins | 0.0555556 |
| Drosophila | dnds | Transcription__factors | DNA__repair__and__recombination__proteins | 0.0185185 |
| Drosophila | dnds | Transcription__factors | DNA__replication__proteins | 0.0185185 |
| Drosophila | dnds | Transcription__factors | Exosome | 0.0185185 |
| Drosophila | dnds | Transcription__factors | G__protein__coupled__receptors | 0.0185185 |
| Drosophila | dnds | Transcription__factors | Glycosyltransferases | 0.0185185 |
| Drosophila | dnds | Transcription__factors | Ion__channels | 0.0185185 |
| Drosophila | dnds | Transcription__factors | Lipid__biosynthesis__proteins | 0.5370370 |
| Drosophila | dnds | Transcription__factors | Membrane__trafficking | 0.0185185 |
| Drosophila | dnds | Transcription__factors | Messenger__RNA__biogenesis | 0.0185185 |
| Drosophila | dnds | Transcription__factors | Mitochondrial__biogenesis | 0.0185185 |
| Drosophila | dnds | Transcription__factors | Peptidases | 0.3148148 |
| Drosophila | dnds | Transcription__factors | Photosynthesis__proteins | 0.0555556 |
| Drosophila | dnds | Transcription__factors | Proteasome | 0.0555556 |
| Drosophila | dnds | Transcription__factors | Protein__kinases | 0.0185185 |
| Drosophila | dnds | Transcription__factors | Protein__phosphatases__and__associated__proteins | 0.8333333 |
| Drosophila | dnds | Transcription__factors | Ribosome | 0.1296296 |
| Drosophila | dnds | Transcription__factors | Ribosome__biogenesis | 0.3888889 |
| Drosophila | dnds | Transcription__factors | Spliceosome | 0.9444444 |
| Drosophila | dnds | Transcription__machinery | Amino__acid__related__enzymes | 0.7222222 |
| Drosophila | dnds | Transcription__machinery | Chaperones__and__folding__catalysts | 0.3518519 |
| Drosophila | dnds | Transcription__machinery | Chromosome__and__associated__proteins | 0.0185185 |
| Drosophila | dnds | Transcription__machinery | Cytochrome__P450 | 0.6111111 |
| Drosophila | dnds | Transcription__machinery | Cytoskeleton__proteins | 0.9074074 |
| Drosophila | dnds | Transcription__machinery | DNA__repair__and__recombination__proteins | 0.0185185 |
| Drosophila | dnds | Transcription__machinery | DNA__replication__proteins | 0.7222222 |
| Drosophila | dnds | Transcription__machinery | Exosome | 0.7962963 |
| Drosophila | dnds | Transcription__machinery | G__protein__coupled__receptors | 0.0555556 |
| Drosophila | dnds | Transcription__machinery | Glycosyltransferases | 0.5740741 |
| Drosophila | dnds | Transcription__machinery | Ion__channels | 0.0185185 |
| Drosophila | dnds | Transcription__machinery | Lipid__biosynthesis__proteins | 0.7222222 |

*(continued)*

| species | estimate | var1 | var2 | p.value |
|---|---|---|---|---|
| Drosophila | dnds | Transcription__machinery | Membrane__trafficking | 0.0185185 |
| Drosophila | dnds | Transcription__machinery | Messenger__RNA__biogenesis | 0.0185185 |
| Drosophila | dnds | Transcription__machinery | Mitochondrial__biogenesis | 0.6851852 |
| Drosophila | dnds | Transcription__machinery | Peptidases | 0.0555556 |
| Drosophila | dnds | Transcription__machinery | Photosynthesis__proteins | 0.3518519 |
| Drosophila | dnds | Transcription__machinery | Proteasome | 0.3518519 |
| Drosophila | dnds | Transcription__machinery | Protein__kinases | 0.1296296 |
| Drosophila | dnds | Transcription__machinery | Protein__phosphatases__and__associated__proteins | 0.0555556 |
| Drosophila | dnds | Transcription__machinery | Ribosome | 0.3888889 |
| Drosophila | dnds | Transcription__machinery | Ribosome__biogenesis | 0.0185185 |
| Drosophila | dnds | Transcription__machinery | Spliceosome | 0.0555556 |
| Drosophila | dnds | Transcription__machinery | Transcription__factors | 0.0185185 |
| Drosophila | dnds | Transfer__RNA__biogenesis | Amino__acid__related__enzymes | 0.8703704 |
| Drosophila | dnds | Transfer__RNA__biogenesis | Chaperones__and__folding__catalysts | 0.5000000 |
| Drosophila | dnds | Transfer__RNA__biogenesis | Chromosome__and__associated__proteins | 0.0185185 |
| Drosophila | dnds | Transfer__RNA__biogenesis | Cytochrome__P450 | 0.7962963 |
| Drosophila | dnds | Transfer__RNA__biogenesis | Cytoskeleton__proteins | 0.5370370 |
| Drosophila | dnds | Transfer__RNA__biogenesis | DNA__repair__and__recombination__proteins | 0.0185185 |
| Drosophila | dnds | Transfer__RNA__biogenesis | DNA__replication__proteins | 0.0925926 |
| Drosophila | dnds | Transfer__RNA__biogenesis | Exosome | 0.2037037 |
| Drosophila | dnds | Transfer__RNA__biogenesis | G__protein__coupled__receptors | 0.0185185 |
| Drosophila | dnds | Transfer__RNA__biogenesis | Glycosyltransferases | 0.9814815 |
| Drosophila | dnds | Transfer__RNA__biogenesis | Ion__channels | 0.0185185 |
| Drosophila | dnds | Transfer__RNA__biogenesis | Lipid__biosynthesis__proteins | 0.9444444 |
| Drosophila | dnds | Transfer__RNA__biogenesis | Membrane__trafficking | 0.0185185 |
| Drosophila | dnds | Transfer__RNA__biogenesis | Messenger__RNA__biogenesis | 0.0185185 |
| Drosophila | dnds | Transfer__RNA__biogenesis | Mitochondrial__biogenesis | 0.4629630 |
| Drosophila | dnds | Transfer__RNA__biogenesis | Peptidases | 0.0185185 |
| Drosophila | dnds | Transfer__RNA__biogenesis | Photosynthesis__proteins | 0.1296296 |
| Drosophila | dnds | Transfer__RNA__biogenesis | Proteasome | 0.1296296 |
| Drosophila | dnds | Transfer__RNA__biogenesis | Protein__kinases | 0.0185185 |
| Drosophila | dnds | Transfer__RNA__biogenesis | Protein__phosphatases__and__associated__proteins | 0.0555556 |
| Drosophila | dnds | Transfer__RNA__biogenesis | Ribosome | 0.9814815 |
| Drosophila | dnds | Transfer__RNA__biogenesis | Ribosome__biogenesis | 0.0185185 |
| Drosophila | dnds | Transfer__RNA__biogenesis | Spliceosome | 0.1666667 |
| Drosophila | dnds | Transfer__RNA__biogenesis | Transcription__factors | 0.0185185 |
| Drosophila | dnds | Transfer__RNA__biogenesis | Transcription__machinery | 0.6851852 |
| Drosophila | dnds | Translation__factors | Amino__acid__related__enzymes | 0.4629630 |
| Drosophila | dnds | Translation__factors | Chaperones__and__folding__catalysts | 0.7962963 |
| Drosophila | dnds | Translation__factors | Chromosome__and__associated__proteins | 0.6481481 |
| Drosophila | dnds | Translation__factors | Cytochrome__P450 | 0.8333333 |
| Drosophila | dnds | Translation__factors | Cytoskeleton__proteins | 0.3888889 |
| Drosophila | dnds | Translation__factors | DNA__repair__and__recombination__proteins | 0.0925926 |
| Drosophila | dnds | Translation__factors | DNA__replication__proteins | 0.3148148 |
| Drosophila | dnds | Translation__factors | Exosome | 0.3148148 |
| Drosophila | dnds | Translation__factors | G__protein__coupled__receptors | 0.0185185 |
| Drosophila | dnds | Translation__factors | Glycosyltransferases | 0.5740741 |
| Drosophila | dnds | Translation__factors | Ion__channels | 0.0185185 |
| Drosophila | dnds | Translation__factors | Lipid__biosynthesis__proteins | 0.3148148 |
| Drosophila | dnds | Translation__factors | Membrane__trafficking | 0.0555556 |
| Drosophila | dnds | Translation__factors | Messenger__RNA__biogenesis | 0.0555556 |
| Drosophila | dnds | Translation__factors | Mitochondrial__biogenesis | 0.3888889 |
| Drosophila | dnds | Translation__factors | Peptidases | 0.6851852 |
| Drosophila | dnds | Translation__factors | Photosynthesis__proteins | 0.1666667 |
| Drosophila | dnds | Translation__factors | Proteasome | 0.1296296 |
| Drosophila | dnds | Translation__factors | Protein__kinases | 0.0925926 |
| Drosophila | dnds | Translation__factors | Protein__phosphatases__and__associated__proteins | 0.8703704 |
| Drosophila | dnds | Translation__factors | Ribosome | 0.5370370 |
| Drosophila | dnds | Translation__factors | Ribosome__biogenesis | 0.7592593 |
| Drosophila | dnds | Translation__factors | Spliceosome | 0.9814815 |
| Drosophila | dnds | Translation__factors | Transcription__factors | 0.9814815 |
| Drosophila | dnds | Translation__factors | Transcription__machinery | 0.3518519 |

*(continued)*

| species | estimate | var1 | var2 | p.value |
|---|---|---|---|---|
| Drosophila | dnds | Translation__factors | Transfer__RNA__biogenesis | 0.5370370 |
| Drosophila | dnds | Transporters | Amino__acid__related__enzymes | 0.3888889 |
| Drosophila | dnds | Transporters | Chaperones__and__folding__catalysts | 0.2037037 |
| Drosophila | dnds | Transporters | Chromosome__and__associated__proteins | 0.0185185 |
| Drosophila | dnds | Transporters | Cytochrome__P450 | 0.6851852 |
| Drosophila | dnds | Transporters | Cytoskeleton__proteins | 0.7592593 |
| Drosophila | dnds | Transporters | DNA__repair__and__recombination__proteins | 0.0185185 |
| Drosophila | dnds | Transporters | DNA__replication__proteins | 0.8703704 |
| Drosophila | dnds | Transporters | Exosome | 0.9814815 |
| Drosophila | dnds | Transporters | G__protein__coupled__receptors | 0.0925926 |
| Drosophila | dnds | Transporters | Glycosyltransferases | 0.1666667 |
| Drosophila | dnds | Transporters | Ion__channels | 0.0185185 |
| Drosophila | dnds | Transporters | Lipid__biosynthesis__proteins | 0.7222222 |
| Drosophila | dnds | Transporters | Membrane__trafficking | 0.0185185 |
| Drosophila | dnds | Transporters | Messenger__RNA__biogenesis | 0.0185185 |
| Drosophila | dnds | Transporters | Mitochondrial__biogenesis | 0.3518519 |
| Drosophila | dnds | Transporters | Peptidases | 0.0185185 |
| Drosophila | dnds | Transporters | Photosynthesis__proteins | 0.5000000 |
| Drosophila | dnds | Transporters | Proteasome | 0.7962963 |
| Drosophila | dnds | Transporters | Protein__kinases | 0.2407407 |
| Drosophila | dnds | Transporters | Protein__phosphatases__and__associated__proteins | 0.0185185 |
| Drosophila | dnds | Transporters | Ribosome | 0.0555556 |
| Drosophila | dnds | Transporters | Ribosome__biogenesis | 0.0185185 |
| Drosophila | dnds | Transporters | Spliceosome | 0.0185185 |
| Drosophila | dnds | Transporters | Transcription__factors | 0.0185185 |
| Drosophila | dnds | Transporters | Transcription__machinery | 0.6851852 |
| Drosophila | dnds | Transporters | Transfer__RNA__biogenesis | 0.0925926 |
| Drosophila | dnds | Transporters | Translation__factors | 0.2777778 |
| Drosophila | dnds | Ubiquitin__system | Amino__acid__related__enzymes | 0.2407407 |
| Drosophila | dnds | Ubiquitin__system | Chaperones__and__folding__catalysts | 0.6481481 |
| Drosophila | dnds | Ubiquitin__system | Chromosome__and__associated__proteins | 0.1296296 |
| Drosophila | dnds | Ubiquitin__system | Cytochrome__P450 | 0.8703704 |
| Drosophila | dnds | Ubiquitin__system | Cytoskeleton__proteins | 0.0185185 |
| Drosophila | dnds | Ubiquitin__system | DNA__repair__and__recombination__proteins | 0.0185185 |
| Drosophila | dnds | Ubiquitin__system | DNA__replication__proteins | 0.0185185 |
| Drosophila | dnds | Ubiquitin__system | Exosome | 0.0185185 |
| Drosophila | dnds | Ubiquitin__system | G__protein__coupled__receptors | 0.0185185 |
| Drosophila | dnds | Ubiquitin__system | Glycosyltransferases | 0.0185185 |
| Drosophila | dnds | Ubiquitin__system | Ion__channels | 0.0185185 |
| Drosophila | dnds | Ubiquitin__system | Lipid__biosynthesis__proteins | 0.5740741 |
| Drosophila | dnds | Ubiquitin__system | Membrane__trafficking | 0.0185185 |
| Drosophila | dnds | Ubiquitin__system | Messenger__RNA__biogenesis | 0.0185185 |
| Drosophila | dnds | Ubiquitin__system | Mitochondrial__biogenesis | 0.0555556 |
| Drosophila | dnds | Ubiquitin__system | Peptidases | 0.3888889 |
| Drosophila | dnds | Ubiquitin__system | Photosynthesis__proteins | 0.0555556 |
| Drosophila | dnds | Ubiquitin__system | Proteasome | 0.0555556 |
| Drosophila | dnds | Ubiquitin__system | Protein__kinases | 0.0185185 |
| Drosophila | dnds | Ubiquitin__system | Protein__phosphatases__and__associated__proteins | 0.5000000 |
| Drosophila | dnds | Ubiquitin__system | Ribosome | 0.1666667 |
| Drosophila | dnds | Ubiquitin__system | Ribosome__biogenesis | 0.2037037 |
| Drosophila | dnds | Ubiquitin__system | Spliceosome | 0.7592593 |
| Drosophila | dnds | Ubiquitin__system | Transcription__factors | 0.9444444 |
| Drosophila | dnds | Ubiquitin__system | Transcription__machinery | 0.0185185 |
| Drosophila | dnds | Ubiquitin__system | Transfer__RNA__biogenesis | 0.0555556 |
| Drosophila | dnds | Ubiquitin__system | Translation__factors | 0.8703704 |
| Drosophila | dnds | Ubiquitin__system | Transporters | 0.0185185 |
| Drosophila | omegaNA | Chaperones__and__folding__catalysts | Amino__acid__related__enzymes | 0.7222222 |
| Drosophila | omegaNA | Chromosome__and__associated__proteins | Amino__acid__related__enzymes | 0.9444444 |
| Drosophila | omegaNA | Chromosome__and__associated__proteins | Chaperones__and__folding__catalysts | 0.4259259 |
| Drosophila | omegaNA | Cytochrome__P450 | Amino__acid__related__enzymes | 0.1666667 |
| Drosophila | omegaNA | Cytochrome__P450 | Chaperones__and__folding__catalysts | 0.9814815 |

*(continued)*

| species | estimate | var1 | var2 | p.value |
|---|---|---|---|---|
| Drosophila | omegaNA | Cytochrome__P450 | Chromosome__and__associated__proteins | 0.4629630 |
| Drosophila | omegaNA | Cytoskeleton__proteins | Amino__acid__related__enzymes | 0.9444444 |
| Drosophila | omegaNA | Cytoskeleton__proteins | Chaperones__and__folding__catalysts | 0.4629630 |
| Drosophila | omegaNA | Cytoskeleton__proteins | Chromosome__and__associated__proteins | 0.6111111 |
| Drosophila | omegaNA | Cytoskeleton__proteins | Cytochrome__P450 | 0.7222222 |
| Drosophila | omegaNA | DNA__repair__and__recombination__proteins | Amino__acid__related__enzymes | 0.5740741 |
| Drosophila | omegaNA | DNA__repair__and__recombination__proteins | Chaperones__and__folding__catalysts | 0.5740741 |
| Drosophila | omegaNA | DNA__repair__and__recombination__proteins | Chromosome__and__associated__proteins | 0.1296296 |
| Drosophila | omegaNA | DNA__repair__and__recombination__proteins | Cytochrome__P450 | 0.8333333 |
| Drosophila | omegaNA | DNA__repair__and__recombination__proteins | Cytoskeleton__proteins | 0.2037037 |
| Drosophila | omegaNA | DNA__replication__proteins | Amino__acid__related__enzymes | 0.3518519 |
| Drosophila | omegaNA | DNA__replication__proteins | Chaperones__and__folding__catalysts | 0.4629630 |
| Drosophila | omegaNA | DNA__replication__proteins | Chromosome__and__associated__proteins | 0.1666667 |
| Drosophila | omegaNA | DNA__replication__proteins | Cytochrome__P450 | 0.9074074 |
| Drosophila | omegaNA | DNA__replication__proteins | Cytoskeleton__proteins | 0.0925926 |
| Drosophila | omegaNA | DNA__replication__proteins | DNA__repair__and__recombination__proteins | 0.7962963 |
| Drosophila | omegaNA | Exosome | Amino__acid__related__enzymes | 0.9074074 |
| Drosophila | omegaNA | Exosome | Chaperones__and__folding__catalysts | 0.5370370 |
| Drosophila | omegaNA | Exosome | Chromosome__and__associated__proteins | 0.6111111 |
| Drosophila | omegaNA | Exosome | Cytochrome__P450 | 0.7222222 |
| Drosophila | omegaNA | Exosome | Cytoskeleton__proteins | 0.8333333 |
| Drosophila | omegaNA | Exosome | DNA__repair__and__recombination__proteins | 0.2037037 |
| Drosophila | omegaNA | Exosome | DNA__replication__proteins | 0.1296296 |
| Drosophila | omegaNA | G__protein__coupled__receptors | Amino__acid__related__enzymes | 0.2777778 |
| Drosophila | omegaNA | G__protein__coupled__receptors | Chaperones__and__folding__catalysts | 0.9074074 |
| Drosophila | omegaNA | G__protein__coupled__receptors | Chromosome__and__associated__proteins | 0.4259259 |
| Drosophila | omegaNA | G__protein__coupled__receptors | Cytochrome__P450 | 0.6481481 |
| Drosophila | omegaNA | G__protein__coupled__receptors | Cytoskeleton__proteins | 0.6851852 |
| Drosophila | omegaNA | G__protein__coupled__receptors | DNA__repair__and__recombination__proteins | 0.8703704 |
| Drosophila | omegaNA | G__protein__coupled__receptors | DNA__replication__proteins | 0.8333333 |
| Drosophila | omegaNA | G__protein__coupled__receptors | Exosome | 0.7592593 |
| Drosophila | omegaNA | Glycosyltransferases | Amino__acid__related__enzymes | 0.3888889 |
| Drosophila | omegaNA | Glycosyltransferases | Chaperones__and__folding__catalysts | 0.6851852 |
| Drosophila | omegaNA | Glycosyltransferases | Chromosome__and__associated__proteins | 0.1666667 |
| Drosophila | omegaNA | Glycosyltransferases | Cytochrome__P450 | 0.9444444 |
| Drosophila | omegaNA | Glycosyltransferases | Cytoskeleton__proteins | 0.2777778 |
| Drosophila | omegaNA | Glycosyltransferases | DNA__repair__and__recombination__proteins | 0.8333333 |
| Drosophila | omegaNA | Glycosyltransferases | DNA__replication__proteins | 0.9814815 |
| Drosophila | omegaNA | Glycosyltransferases | Exosome | 0.3148148 |
| Drosophila | omegaNA | Glycosyltransferases | G__protein__coupled__receptors | 0.8333333 |
| Drosophila | omegaNA | Ion__channels | Amino__acid__related__enzymes | 0.7962963 |
| Drosophila | omegaNA | Ion__channels | Chaperones__and__folding__catalysts | 0.2037037 |
| Drosophila | omegaNA | Ion__channels | Chromosome__and__associated__proteins | 0.5000000 |
| Drosophila | omegaNA | Ion__channels | Cytochrome__P450 | 0.0925926 |
| Drosophila | omegaNA | Ion__channels | Cytoskeleton__proteins | 0.1296296 |
| Drosophila | omegaNA | Ion__channels | DNA__repair__and__recombination__proteins | 0.0185185 |
| Drosophila | omegaNA | Ion__channels | DNA__replication__proteins | 0.0185185 |
| Drosophila | omegaNA | Ion__channels | Exosome | 0.1666667 |
| Drosophila | omegaNA | Ion__channels | G__protein__coupled__receptors | 0.0555556 |
| Drosophila | omegaNA | Ion__channels | Glycosyltransferases | 0.0185185 |
| Drosophila | omegaNA | Lipid__biosynthesis__proteins | Amino__acid__related__enzymes | 0.1296296 |
| Drosophila | omegaNA | Lipid__biosynthesis__proteins | Chaperones__and__folding__catalysts | 0.9814815 |
| Drosophila | omegaNA | Lipid__biosynthesis__proteins | Chromosome__and__associated__proteins | 0.4259259 |
| Drosophila | omegaNA | Lipid__biosynthesis__proteins | Cytochrome__P450 | 0.8703704 |
| Drosophila | omegaNA | Lipid__biosynthesis__proteins | Cytoskeleton__proteins | 0.7962963 |
| Drosophila | omegaNA | Lipid__biosynthesis__proteins | DNA__repair__and__recombination__proteins | 0.9074074 |
| Drosophila | omegaNA | Lipid__biosynthesis__proteins | DNA__replication__proteins | 0.8703704 |
| Drosophila | omegaNA | Lipid__biosynthesis__proteins | Exosome | 0.8333333 |
| Drosophila | omegaNA | Lipid__biosynthesis__proteins | G__protein__coupled__receptors | 0.7592593 |
| Drosophila | omegaNA | Lipid__biosynthesis__proteins | Glycosyltransferases | 0.7962963 |
| Drosophila | omegaNA | Lipid__biosynthesis__proteins | Ion__channels | 0.1666667 |
| Drosophila | omegaNA | Membrane__trafficking | Amino__acid__related__enzymes | 0.9444444 |

*(continued)*

| species | estimate | var1 | var2 | p.value |
|---|---|---|---|---|
| Drosophila | omegaNA | Membrane__trafficking | Chaperones__and__folding__catalysts | 0.0925926 |
| Drosophila | omegaNA | Membrane__trafficking | Chromosome__and__associated__proteins | 0.0185185 |
| Drosophila | omegaNA | Membrane__trafficking | Cytochrome__P450 | 0.0925926 |
| Drosophila | omegaNA | Membrane__trafficking | Cytoskeleton__proteins | 0.0185185 |
| Drosophila | omegaNA | Membrane__trafficking | DNA__repair__and__recombination__proteins | 0.0185185 |
| Drosophila | omegaNA | Membrane__trafficking | DNA__replication__proteins | 0.0185185 |
| Drosophila | omegaNA | Membrane__trafficking | Exosome | 0.0555556 |
| Drosophila | omegaNA | Membrane__trafficking | G__protein__coupled__receptors | 0.0925926 |
| Drosophila | omegaNA | Membrane__trafficking | Glycosyltransferases | 0.0555556 |
| Drosophila | omegaNA | Membrane__trafficking | Ion__channels | 0.8333333 |
| Drosophila | omegaNA | Membrane__trafficking | Lipid__biosynthesis__proteins | 0.0925926 |
| Drosophila | omegaNA | Messenger__RNA__biogenesis | Amino__acid__related__enzymes | 0.9444444 |
| Drosophila | omegaNA | Messenger__RNA__biogenesis | Chaperones__and__folding__catalysts | 0.3148148 |
| Drosophila | omegaNA | Messenger__RNA__biogenesis | Chromosome__and__associated__proteins | 0.7962963 |
| Drosophila | omegaNA | Messenger__RNA__biogenesis | Cytochrome__P450 | 0.5740741 |
| Drosophila | omegaNA | Messenger__RNA__biogenesis | Cytoskeleton__proteins | 0.5370370 |
| Drosophila | omegaNA | Messenger__RNA__biogenesis | DNA__repair__and__recombination__proteins | 0.0555556 |
| Drosophila | omegaNA | Messenger__RNA__biogenesis | DNA__replication__proteins | 0.0555556 |
| Drosophila | omegaNA | Messenger__RNA__biogenesis | Exosome | 0.5370370 |
| Drosophila | omegaNA | Messenger__RNA__biogenesis | G__protein__coupled__receptors | 0.5000000 |
| Drosophila | omegaNA | Messenger__RNA__biogenesis | Glycosyltransferases | 0.2037037 |
| Drosophila | omegaNA | Messenger__RNA__biogenesis | Ion__channels | 0.2037037 |
| Drosophila | omegaNA | Messenger__RNA__biogenesis | Lipid__biosynthesis__proteins | 0.5370370 |
| Drosophila | omegaNA | Messenger__RNA__biogenesis | Membrane__trafficking | 0.0555556 |
| Drosophila | omegaNA | Mitochondrial__biogenesis | Amino__acid__related__enzymes | 0.7592593 |
| Drosophila | omegaNA | Mitochondrial__biogenesis | Chaperones__and__folding__catalysts | 0.6111111 |
| Drosophila | omegaNA | Mitochondrial__biogenesis | Chromosome__and__associated__proteins | 0.8333333 |
| Drosophila | omegaNA | Mitochondrial__biogenesis | Cytochrome__P450 | 0.4629630 |
| Drosophila | omegaNA | Mitochondrial__biogenesis | Cytoskeleton__proteins | 0.7962963 |
| Drosophila | omegaNA | Mitochondrial__biogenesis | DNA__repair__and__recombination__proteins | 0.3518519 |
| Drosophila | omegaNA | Mitochondrial__biogenesis | DNA__replication__proteins | 0.4629630 |
| Drosophila | omegaNA | Mitochondrial__biogenesis | Exosome | 0.6111111 |
| Drosophila | omegaNA | Mitochondrial__biogenesis | G__protein__coupled__receptors | 0.5000000 |
| Drosophila | omegaNA | Mitochondrial__biogenesis | Glycosyltransferases | 0.3888889 |
| Drosophila | omegaNA | Mitochondrial__biogenesis | Ion__channels | 0.5370370 |
| Drosophila | omegaNA | Mitochondrial__biogenesis | Lipid__biosynthesis__proteins | 0.5740741 |
| Drosophila | omegaNA | Mitochondrial__biogenesis | Membrane__trafficking | 0.1296296 |
| Drosophila | omegaNA | Mitochondrial__biogenesis | Messenger__RNA__biogenesis | 0.8703704 |
| Drosophila | omegaNA | Peptidases | Amino__acid__related__enzymes | 0.9444444 |
| Drosophila | omegaNA | Peptidases | Chaperones__and__folding__catalysts | 0.0185185 |
| Drosophila | omegaNA | Peptidases | Chromosome__and__associated__proteins | 0.0555556 |
| Drosophila | omegaNA | Peptidases | Cytochrome__P450 | 0.1296296 |
| Drosophila | omegaNA | Peptidases | Cytoskeleton__proteins | 0.0185185 |
| Drosophila | omegaNA | Peptidases | DNA__repair__and__recombination__proteins | 0.0185185 |
| Drosophila | omegaNA | Peptidases | DNA__replication__proteins | 0.0185185 |
| Drosophila | omegaNA | Peptidases | Exosome | 0.0555556 |
| Drosophila | omegaNA | Peptidases | G__protein__coupled__receptors | 0.0925926 |
| Drosophila | omegaNA | Peptidases | Glycosyltransferases | 0.0185185 |
| Drosophila | omegaNA | Peptidases | Ion__channels | 0.8703704 |
| Drosophila | omegaNA | Peptidases | Lipid__biosynthesis__proteins | 0.0185185 |
| Drosophila | omegaNA | Peptidases | Membrane__trafficking | 0.8333333 |
| Drosophila | omegaNA | Peptidases | Messenger__RNA__biogenesis | 0.0555556 |
| Drosophila | omegaNA | Peptidases | Mitochondrial__biogenesis | 0.1666667 |
| Drosophila | omegaNA | Photosynthesis__proteins | Amino__acid__related__enzymes | 0.9814815 |
| Drosophila | omegaNA | Photosynthesis__proteins | Chaperones__and__folding__catalysts | 0.3518519 |
| Drosophila | omegaNA | Photosynthesis__proteins | Chromosome__and__associated__proteins | 0.7592593 |
| Drosophila | omegaNA | Photosynthesis__proteins | Cytochrome__P450 | 0.1296296 |
| Drosophila | omegaNA | Photosynthesis__proteins | Cytoskeleton__proteins | 0.6481481 |
| Drosophila | omegaNA | Photosynthesis__proteins | DNA__repair__and__recombination__proteins | 0.2037037 |
| Drosophila | omegaNA | Photosynthesis__proteins | DNA__replication__proteins | 0.0925926 |
| Drosophila | omegaNA | Photosynthesis__proteins | Exosome | 0.7222222 |
| Drosophila | omegaNA | Photosynthesis__proteins | G__protein__coupled__receptors | 0.0925926 |

*(continued)*

| species | estimate | var1 | var2 | p.value |
|---|---|---|---|---|
| Drosophila | omegaNA | Photosynthesis__proteins | Glycosyltransferases | 0.1296296 |
| Drosophila | omegaNA | Photosynthesis__proteins | Ion__channels | 0.4629630 |
| Drosophila | omegaNA | Photosynthesis__proteins | Lipid__biosynthesis__proteins | 0.0925926 |
| Drosophila | omegaNA | Photosynthesis__proteins | Membrane__trafficking | 0.7962963 |
| Drosophila | omegaNA | Photosynthesis__proteins | Messenger__RNA__biogenesis | 0.9074074 |
| Drosophila | omegaNA | Photosynthesis__proteins | Mitochondrial__biogenesis | 0.7222222 |
| Drosophila | omegaNA | Photosynthesis__proteins | Peptidases | 0.7592593 |
| Drosophila | omegaNA | Proteasome | Amino__acid__related__enzymes | 0.9444444 |
| Drosophila | omegaNA | Proteasome | Chaperones__and__folding__catalysts | 0.3888889 |
| Drosophila | omegaNA | Proteasome | Chromosome__and__associated__proteins | 0.9074074 |
| Drosophila | omegaNA | Proteasome | Cytochrome__P450 | 0.2407407 |
| Drosophila | omegaNA | Proteasome | Cytoskeleton__proteins | 0.7592593 |
| Drosophila | omegaNA | Proteasome | DNA__repair__and__recombination__proteins | 0.2407407 |
| Drosophila | omegaNA | Proteasome | DNA__replication__proteins | 0.1296296 |
| Drosophila | omegaNA | Proteasome | Exosome | 0.7592593 |
| Drosophila | omegaNA | Proteasome | G__protein__coupled__receptors | 0.2037037 |
| Drosophila | omegaNA | Proteasome | Glycosyltransferases | 0.1666667 |
| Drosophila | omegaNA | Proteasome | Ion__channels | 0.5370370 |
| Drosophila | omegaNA | Proteasome | Lipid__biosynthesis__proteins | 0.1666667 |
| Drosophila | omegaNA | Proteasome | Membrane__trafficking | 0.6481481 |
| Drosophila | omegaNA | Proteasome | Messenger__RNA__biogenesis | 0.9814815 |
| Drosophila | omegaNA | Proteasome | Mitochondrial__biogenesis | 0.8703704 |
| Drosophila | omegaNA | Proteasome | Peptidases | 0.6481481 |
| Drosophila | omegaNA | Proteasome | Photosynthesis__proteins | 0.8703704 |
| Drosophila | omegaNA | Protein__kinases | Amino__acid__related__enzymes | 0.9814815 |
| Drosophila | omegaNA | Protein__kinases | Chaperones__and__folding__catalysts | 0.0555556 |
| Drosophila | omegaNA | Protein__kinases | Chromosome__and__associated__proteins | 0.0185185 |
| Drosophila | omegaNA | Protein__kinases | Cytochrome__P450 | 0.0925926 |
| Drosophila | omegaNA | Protein__kinases | Cytoskeleton__proteins | 0.0185185 |
| Drosophila | omegaNA | Protein__kinases | DNA__repair__and__recombination__proteins | 0.0185185 |
| Drosophila | omegaNA | Protein__kinases | DNA__replication__proteins | 0.0185185 |
| Drosophila | omegaNA | Protein__kinases | Exosome | 0.0185185 |
| Drosophila | omegaNA | Protein__kinases | G__protein__coupled__receptors | 0.0555556 |
| Drosophila | omegaNA | Protein__kinases | Glycosyltransferases | 0.0185185 |
| Drosophila | omegaNA | Protein__kinases | Ion__channels | 0.8333333 |
| Drosophila | omegaNA | Protein__kinases | Lipid__biosynthesis__proteins | 0.0925926 |
| Drosophila | omegaNA | Protein__kinases | Membrane__trafficking | 0.7962963 |
| Drosophila | omegaNA | Protein__kinases | Messenger__RNA__biogenesis | 0.0185185 |
| Drosophila | omegaNA | Protein__kinases | Mitochondrial__biogenesis | 0.2777778 |
| Drosophila | omegaNA | Protein__kinases | Peptidases | 0.6851852 |
| Drosophila | omegaNA | Protein__kinases | Photosynthesis__proteins | 0.7222222 |
| Drosophila | omegaNA | Protein__kinases | Proteasome | 0.6851852 |
| Drosophila | omegaNA | Protein__phosphatases__and__associated__proteins | Amino__acid__related__enzymes | 0.9074074 |
| Drosophila | omegaNA | Protein__phosphatases__and__associated__proteins | Chaperones__and__folding__catalysts | 0.3518519 |
| Drosophila | omegaNA | Protein__phosphatases__and__associated__proteins | Chromosome__and__associated__proteins | 0.6851852 |
| Drosophila | omegaNA | Protein__phosphatases__and__associated__proteins | Cytochrome__P450 | 0.2037037 |
| Drosophila | omegaNA | Protein__phosphatases__and__associated__proteins | Cytoskeleton__proteins | 0.3888889 |
| Drosophila | omegaNA | Protein__phosphatases__and__associated__proteins | DNA__repair__and__recombination__proteins | 0.1666667 |
| Drosophila | omegaNA | Protein__phosphatases__and__associated__proteins | DNA__replication__proteins | 0.0925926 |
| Drosophila | omegaNA | Protein__phosphatases__and__associated__proteins | Exosome | 0.4259259 |
| Drosophila | omegaNA | Protein__phosphatases__and__associated__proteins | G__protein__coupled__receptors | 0.2407407 |
| Drosophila | omegaNA | Protein__phosphatases__and__associated__proteins | Glycosyltransferases | 0.0555556 |
| Drosophila | omegaNA | Protein__phosphatases__and__associated__proteins | Ion__channels | 0.7222222 |
| Drosophila | omegaNA | Protein__phosphatases__and__associated__proteins | Lipid__biosynthesis__proteins | 0.3888889 |
| Drosophila | omegaNA | Protein__phosphatases__and__associated__proteins | Membrane__trafficking | 0.0925926 |
| Drosophila | omegaNA | Protein__phosphatases__and__associated__proteins | Messenger__RNA__biogenesis | 0.3888889 |
| Drosophila | omegaNA | Protein__phosphatases__and__associated__proteins | Mitochondrial__biogenesis | 0.9444444 |
| Drosophila | omegaNA | Protein__phosphatases__and__associated__proteins | Peptidases | 0.0925926 |
| Drosophila | omegaNA | Protein__phosphatases__and__associated__proteins | Photosynthesis__proteins | 0.8333333 |
| Drosophila | omegaNA | Protein__phosphatases__and__associated__proteins | Proteasome | 0.9444444 |
| Drosophila | omegaNA | Protein__phosphatases__and__associated__proteins | Protein__kinases | 0.0925926 |

*(continued)*

| species | estimate | var1 | var2 | p.value |
|---|---|---|---|---|
| Drosophila | omegaNA | Ribosome | Amino__acid__related__enzymes | 0.7222222 |
| Drosophila | omegaNA | Ribosome | Chaperones__and__folding__catalysts | 0.6851852 |
| Drosophila | omegaNA | Ribosome | Chromosome__and__associated__proteins | 0.9074074 |
| Drosophila | omegaNA | Ribosome | Cytochrome__P450 | 0.7592593 |
| Drosophila | omegaNA | Ribosome | Cytoskeleton__proteins | 0.9074074 |
| Drosophila | omegaNA | Ribosome | DNA__repair__and__recombination__proteins | 0.4629630 |
| Drosophila | omegaNA | Ribosome | DNA__replication__proteins | 0.3148148 |
| Drosophila | omegaNA | Ribosome | Exosome | 0.9074074 |
| Drosophila | omegaNA | Ribosome | G__protein__coupled__receptors | 0.6851852 |
| Drosophila | omegaNA | Ribosome | Glycosyltransferases | 0.3888889 |
| Drosophila | omegaNA | Ribosome | Ion__channels | 0.4629630 |
| Drosophila | omegaNA | Ribosome | Lipid__biosynthesis__proteins | 0.6851852 |
| Drosophila | omegaNA | Ribosome | Membrane__trafficking | 0.0185185 |
| Drosophila | omegaNA | Ribosome | Messenger__RNA__biogenesis | 0.9814815 |
| Drosophila | omegaNA | Ribosome | Mitochondrial__biogenesis | 0.8703704 |
| Drosophila | omegaNA | Ribosome | Peptidases | 0.0555556 |
| Drosophila | omegaNA | Ribosome | Photosynthesis__proteins | 0.5370370 |
| Drosophila | omegaNA | Ribosome | Proteasome | 0.7592593 |
| Drosophila | omegaNA | Ribosome | Protein__kinases | 0.0185185 |
| Drosophila | omegaNA | Ribosome | Protein__phosphatases__and__associated__proteins | 0.5740741 |
| Drosophila | omegaNA | Ribosome__biogenesis | Amino__acid__related__enzymes | 0.9444444 |
| Drosophila | omegaNA | Ribosome__biogenesis | Chaperones__and__folding__catalysts | 0.2037037 |
| Drosophila | omegaNA | Ribosome__biogenesis | Chromosome__and__associated__proteins | 0.4259259 |
| Drosophila | omegaNA | Ribosome__biogenesis | Cytochrome__P450 | 0.1666667 |
| Drosophila | omegaNA | Ribosome__biogenesis | Cytoskeleton__proteins | 0.1666667 |
| Drosophila | omegaNA | Ribosome__biogenesis | DNA__repair__and__recombination__proteins | 0.0555556 |
| Drosophila | omegaNA | Ribosome__biogenesis | DNA__replication__proteins | 0.0185185 |
| Drosophila | omegaNA | Ribosome__biogenesis | Exosome | 0.0925926 |
| Drosophila | omegaNA | Ribosome__biogenesis | G__protein__coupled__receptors | 0.1296296 |
| Drosophila | omegaNA | Ribosome__biogenesis | Glycosyltransferases | 0.0185185 |
| Drosophila | omegaNA | Ribosome__biogenesis | Ion__channels | 0.9074074 |
| Drosophila | omegaNA | Ribosome__biogenesis | Lipid__biosynthesis__proteins | 0.1296296 |
| Drosophila | omegaNA | Ribosome__biogenesis | Membrane__trafficking | 0.1296296 |
| Drosophila | omegaNA | Ribosome__biogenesis | Messenger__RNA__biogenesis | 0.2037037 |
| Drosophila | omegaNA | Ribosome__biogenesis | Mitochondrial__biogenesis | 0.7222222 |
| Drosophila | omegaNA | Ribosome__biogenesis | Peptidases | 0.0925926 |
| Drosophila | omegaNA | Ribosome__biogenesis | Photosynthesis__proteins | 0.9814815 |
| Drosophila | omegaNA | Ribosome__biogenesis | Proteasome | 0.7962963 |
| Drosophila | omegaNA | Ribosome__biogenesis | Protein__kinases | 0.0555556 |
| Drosophila | omegaNA | Ribosome__biogenesis | Protein__phosphatases__and__associated__proteins | 0.6111111 |
| Drosophila | omegaNA | Ribosome__biogenesis | Ribosome | 0.2777778 |
| Drosophila | omegaNA | Spliceosome | Amino__acid__related__enzymes | 0.7592593 |
| Drosophila | omegaNA | Spliceosome | Chaperones__and__folding__catalysts | 0.9814815 |
| Drosophila | omegaNA | Spliceosome | Chromosome__and__associated__proteins | 0.3518519 |
| Drosophila | omegaNA | Spliceosome | Cytochrome__P450 | 0.9444444 |
| Drosophila | omegaNA | Spliceosome | Cytoskeleton__proteins | 0.6481481 |
| Drosophila | omegaNA | Spliceosome | DNA__repair__and__recombination__proteins | 0.5000000 |
| Drosophila | omegaNA | Spliceosome | DNA__replication__proteins | 0.3148148 |
| Drosophila | omegaNA | Spliceosome | Exosome | 0.5740741 |
| Drosophila | omegaNA | Spliceosome | G__protein__coupled__receptors | 0.8333333 |
| Drosophila | omegaNA | Spliceosome | Glycosyltransferases | 0.5000000 |
| Drosophila | omegaNA | Spliceosome | Ion__channels | 0.0555556 |
| Drosophila | omegaNA | Spliceosome | Lipid__biosynthesis__proteins | 0.9444444 |
| Drosophila | omegaNA | Spliceosome | Membrane__trafficking | 0.0185185 |
| Drosophila | omegaNA | Spliceosome | Messenger__RNA__biogenesis | 0.2037037 |
| Drosophila | omegaNA | Spliceosome | Mitochondrial__biogenesis | 0.6111111 |
| Drosophila | omegaNA | Spliceosome | Peptidases | 0.0185185 |
| Drosophila | omegaNA | Spliceosome | Photosynthesis__proteins | 0.4629630 |
| Drosophila | omegaNA | Spliceosome | Proteasome | 0.5370370 |
| Drosophila | omegaNA | Spliceosome | Protein__kinases | 0.0185185 |
| Drosophila | omegaNA | Spliceosome | Protein__phosphatases__and__associated__proteins | 0.1666667 |
| Drosophila | omegaNA | Spliceosome | Ribosome | 0.5740741 |

*(continued)*

| species | estimate | var1 | var2 | p.value |
|---|---|---|---|---|
| Drosophila | omegaNA | Spliceosome | Ribosome__biogenesis | 0.0185185 |
| Drosophila | omegaNA | Transcription__factors | Amino__acid__related__enzymes | 0.7592593 |
| Drosophila | omegaNA | Transcription__factors | Chaperones__and__folding__catalysts | 0.8703704 |
| Drosophila | omegaNA | Transcription__factors | Chromosome__and__associated__proteins | 0.2037037 |
| Drosophila | omegaNA | Transcription__factors | Cytochrome__P450 | 0.9814815 |
| Drosophila | omegaNA | Transcription__factors | Cytoskeleton__proteins | 0.2407407 |
| Drosophila | omegaNA | Transcription__factors | DNA__repair__and__recombination__proteins | 0.5740741 |
| Drosophila | omegaNA | Transcription__factors | DNA__replication__proteins | 0.3148148 |
| Drosophila | omegaNA | Transcription__factors | Exosome | 0.3148148 |
| Drosophila | omegaNA | Transcription__factors | G__protein__coupled__receptors | 0.8333333 |
| Drosophila | omegaNA | Transcription__factors | Glycosyltransferases | 0.5000000 |
| Drosophila | omegaNA | Transcription__factors | Ion__channels | 0.0185185 |
| Drosophila | omegaNA | Transcription__factors | Lipid__biosynthesis__proteins | 0.9814815 |
| Drosophila | omegaNA | Transcription__factors | Membrane__trafficking | 0.0185185 |
| Drosophila | omegaNA | Transcription__factors | Messenger__RNA__biogenesis | 0.0555556 |
| Drosophila | omegaNA | Transcription__factors | Mitochondrial__biogenesis | 0.5740741 |
| Drosophila | omegaNA | Transcription__factors | Peptidases | 0.0185185 |
| Drosophila | omegaNA | Transcription__factors | Photosynthesis__proteins | 0.1666667 |
| Drosophila | omegaNA | Transcription__factors | Proteasome | 0.3888889 |
| Drosophila | omegaNA | Transcription__factors | Protein__kinases | 0.0185185 |
| Drosophila | omegaNA | Transcription__factors | Protein__phosphatases__and__associated__proteins | 0.1296296 |
| Drosophila | omegaNA | Transcription__factors | Ribosome | 0.6111111 |
| Drosophila | omegaNA | Transcription__factors | Ribosome__biogenesis | 0.0185185 |
| Drosophila | omegaNA | Transcription__factors | Spliceosome | 0.8703704 |
| Drosophila | omegaNA | Transcription__machinery | Amino__acid__related__enzymes | 0.9444444 |
| Drosophila | omegaNA | Transcription__machinery | Chaperones__and__folding__catalysts | 0.0555556 |
| Drosophila | omegaNA | Transcription__machinery | Chromosome__and__associated__proteins | 0.1666667 |
| Drosophila | omegaNA | Transcription__machinery | Cytochrome__P450 | 0.1296296 |
| Drosophila | omegaNA | Transcription__machinery | Cytoskeleton__proteins | 0.0555556 |
| Drosophila | omegaNA | Transcription__machinery | DNA__repair__and__recombination__proteins | 0.0185185 |
| Drosophila | omegaNA | Transcription__machinery | DNA__replication__proteins | 0.0185185 |
| Drosophila | omegaNA | Transcription__machinery | Exosome | 0.0185185 |
| Drosophila | omegaNA | Transcription__machinery | G__protein__coupled__receptors | 0.0555556 |
| Drosophila | omegaNA | Transcription__machinery | Glycosyltransferases | 0.0185185 |
| Drosophila | omegaNA | Transcription__machinery | Ion__channels | 0.9444444 |
| Drosophila | omegaNA | Transcription__machinery | Lipid__biosynthesis__proteins | 0.0925926 |
| Drosophila | omegaNA | Transcription__machinery | Membrane__trafficking | 0.5740741 |
| Drosophila | omegaNA | Transcription__machinery | Messenger__RNA__biogenesis | 0.0185185 |
| Drosophila | omegaNA | Transcription__machinery | Mitochondrial__biogenesis | 0.4259259 |
| Drosophila | omegaNA | Transcription__machinery | Peptidases | 0.3518519 |
| Drosophila | omegaNA | Transcription__machinery | Photosynthesis__proteins | 0.9444444 |
| Drosophila | omegaNA | Transcription__machinery | Proteasome | 0.7962963 |
| Drosophila | omegaNA | Transcription__machinery | Protein__kinases | 0.6481481 |
| Drosophila | omegaNA | Transcription__machinery | Protein__phosphatases__and__associated__proteins | 0.3148148 |
| Drosophila | omegaNA | Transcription__machinery | Ribosome | 0.0185185 |
| Drosophila | omegaNA | Transcription__machinery | Ribosome__biogenesis | 0.5370370 |
| Drosophila | omegaNA | Transcription__machinery | Spliceosome | 0.0185185 |
| Drosophila | omegaNA | Transcription__machinery | Transcription__factors | 0.0185185 |
| Drosophila | omegaNA | Transfer__RNA__biogenesis | Amino__acid__related__enzymes | 0.9444444 |
| Drosophila | omegaNA | Transfer__RNA__biogenesis | Chaperones__and__folding__catalysts | 0.4259259 |
| Drosophila | omegaNA | Transfer__RNA__biogenesis | Chromosome__and__associated__proteins | 0.5000000 |
| Drosophila | omegaNA | Transfer__RNA__biogenesis | Cytochrome__P450 | 0.5740741 |
| Drosophila | omegaNA | Transfer__RNA__biogenesis | Cytoskeleton__proteins | 0.6481481 |
| Drosophila | omegaNA | Transfer__RNA__biogenesis | DNA__repair__and__recombination__proteins | 0.0555556 |
| Drosophila | omegaNA | Transfer__RNA__biogenesis | DNA__replication__proteins | 0.0185185 |
| Drosophila | omegaNA | Transfer__RNA__biogenesis | Exosome | 0.6851852 |
| Drosophila | omegaNA | Transfer__RNA__biogenesis | G__protein__coupled__receptors | 0.6851852 |
| Drosophila | omegaNA | Transfer__RNA__biogenesis | Glycosyltransferases | 0.2037037 |
| Drosophila | omegaNA | Transfer__RNA__biogenesis | Ion__channels | 0.1296296 |
| Drosophila | omegaNA | Transfer__RNA__biogenesis | Lipid__biosynthesis__proteins | 0.5740741 |
| Drosophila | omegaNA | Transfer__RNA__biogenesis | Membrane__trafficking | 0.0185185 |
| Drosophila | omegaNA | Transfer__RNA__biogenesis | Messenger__RNA__biogenesis | 0.6851852 |

*(continued)*

| species | estimate | var1 | var2 | p.value |
|---|---|---|---|---|
| Drosophila | omegaNA | Transfer__RNA__biogenesis | Mitochondrial__biogenesis | 0.6851852 |
| Drosophila | omegaNA | Transfer__RNA__biogenesis | Peptidases | 0.0185185 |
| Drosophila | omegaNA | Transfer__RNA__biogenesis | Photosynthesis__proteins | 0.7962963 |
| Drosophila | omegaNA | Transfer__RNA__biogenesis | Proteasome | 0.9074074 |
| Drosophila | omegaNA | Transfer__RNA__biogenesis | Protein__kinases | 0.0185185 |
| Drosophila | omegaNA | Transfer__RNA__biogenesis | Protein__phosphatases__and__associated__proteins | 0.4259259 |
| Drosophila | omegaNA | Transfer__RNA__biogenesis | Ribosome | 0.9074074 |
| Drosophila | omegaNA | Transfer__RNA__biogenesis | Ribosome__biogenesis | 0.2037037 |
| Drosophila | omegaNA | Transfer__RNA__biogenesis | Spliceosome | 0.3148148 |
| Drosophila | omegaNA | Transfer__RNA__biogenesis | Transcription__factors | 0.0925926 |
| Drosophila | omegaNA | Transfer__RNA__biogenesis | Transcription__machinery | 0.0185185 |
| Drosophila | omegaNA | Translation__factors | Amino__acid__related__enzymes | 0.1666667 |
| Drosophila | omegaNA | Translation__factors | Chaperones__and__folding__catalysts | 0.9814815 |
| Drosophila | omegaNA | Translation__factors | Chromosome__and__associated__proteins | 0.6111111 |
| Drosophila | omegaNA | Translation__factors | Cytochrome__P450 | 0.7962963 |
| Drosophila | omegaNA | Translation__factors | Cytoskeleton__proteins | 0.9074074 |
| Drosophila | omegaNA | Translation__factors | DNA__repair__and__recombination__proteins | 0.7962963 |
| Drosophila | omegaNA | Translation__factors | DNA__replication__proteins | 0.8333333 |
| Drosophila | omegaNA | Translation__factors | Exosome | 0.8333333 |
| Drosophila | omegaNA | Translation__factors | G__protein__coupled__receptors | 0.7222222 |
| Drosophila | omegaNA | Translation__factors | Glycosyltransferases | 0.9814815 |
| Drosophila | omegaNA | Translation__factors | Ion__channels | 0.1666667 |
| Drosophila | omegaNA | Translation__factors | Lipid__biosynthesis__proteins | 0.7962963 |
| Drosophila | omegaNA | Translation__factors | Membrane__trafficking | 0.1296296 |
| Drosophila | omegaNA | Translation__factors | Messenger__RNA__biogenesis | 0.7592593 |
| Drosophila | omegaNA | Translation__factors | Mitochondrial__biogenesis | 0.5740741 |
| Drosophila | omegaNA | Translation__factors | Peptidases | 0.2037037 |
| Drosophila | omegaNA | Translation__factors | Photosynthesis__proteins | 0.1666667 |
| Drosophila | omegaNA | Translation__factors | Proteasome | 0.2407407 |
| Drosophila | omegaNA | Translation__factors | Protein__kinases | 0.1296296 |
| Drosophila | omegaNA | Translation__factors | Protein__phosphatases__and__associated__proteins | 0.3148148 |
| Drosophila | omegaNA | Translation__factors | Ribosome | 0.7962963 |
| Drosophila | omegaNA | Translation__factors | Ribosome__biogenesis | 0.3148148 |
| Drosophila | omegaNA | Translation__factors | Spliceosome | 0.9814815 |
| Drosophila | omegaNA | Translation__factors | Transcription__factors | 0.9444444 |
| Drosophila | omegaNA | Translation__factors | Transcription__machinery | 0.2037037 |
| Drosophila | omegaNA | Translation__factors | Transfer__RNA__biogenesis | 0.6481481 |
| Drosophila | omegaNA | Transporters | Amino__acid__related__enzymes | 0.9444444 |
| Drosophila | omegaNA | Transporters | Chaperones__and__folding__catalysts | 0.1666667 |
| Drosophila | omegaNA | Transporters | Chromosome__and__associated__proteins | 0.7962963 |
| Drosophila | omegaNA | Transporters | Cytochrome__P450 | 0.3518519 |
| Drosophila | omegaNA | Transporters | Cytoskeleton__proteins | 0.2777778 |
| Drosophila | omegaNA | Transporters | DNA__repair__and__recombination__proteins | 0.0185185 |
| Drosophila | omegaNA | Transporters | DNA__replication__proteins | 0.0185185 |
| Drosophila | omegaNA | Transporters | Exosome | 0.2407407 |
| Drosophila | omegaNA | Transporters | G__protein__coupled__receptors | 0.3148148 |
| Drosophila | omegaNA | Transporters | Glycosyltransferases | 0.0925926 |
| Drosophila | omegaNA | Transporters | Ion__channels | 0.6481481 |
| Drosophila | omegaNA | Transporters | Lipid__biosynthesis__proteins | 0.2407407 |
| Drosophila | omegaNA | Transporters | Membrane__trafficking | 0.0555556 |
| Drosophila | omegaNA | Transporters | Messenger__RNA__biogenesis | 0.3518519 |
| Drosophila | omegaNA | Transporters | Mitochondrial__biogenesis | 0.9814815 |
| Drosophila | omegaNA | Transporters | Peptidases | 0.0925926 |
| Drosophila | omegaNA | Transporters | Photosynthesis__proteins | 0.9814815 |
| Drosophila | omegaNA | Transporters | Proteasome | 0.9074074 |
| Drosophila | omegaNA | Transporters | Protein__kinases | 0.0185185 |
| Drosophila | omegaNA | Transporters | Protein__phosphatases__and__associated__proteins | 0.9074074 |
| Drosophila | omegaNA | Transporters | Ribosome | 0.6111111 |
| Drosophila | omegaNA | Transporters | Ribosome__biogenesis | 0.7222222 |
| Drosophila | omegaNA | Transporters | Spliceosome | 0.1296296 |
| Drosophila | omegaNA | Transporters | Transcription__factors | 0.0185185 |

*(continued)*

| species | estimate | var1 | var2 | p.value |
|---|---|---|---|---|
| Drosophila | omegaNA | Transporters | Transcription__machinery | 0.2777778 |
| Drosophila | omegaNA | Transporters | Transfer__RNA__biogenesis | 0.3888889 |
| Drosophila | omegaNA | Transporters | Translation__factors | 0.5370370 |
| Drosophila | omegaNA | Ubiquitin__system | Amino__acid__related__enzymes | 0.9444444 |
| Drosophila | omegaNA | Ubiquitin__system | Chaperones__and__folding__catalysts | 0.3518519 |
| Drosophila | omegaNA | Ubiquitin__system | Chromosome__and__associated__proteins | 0.9814815 |
| Drosophila | omegaNA | Ubiquitin__system | Cytochrome__P450 | 0.4259259 |
| Drosophila | omegaNA | Ubiquitin__system | Cytoskeleton__proteins | 0.5740741 |
| Drosophila | omegaNA | Ubiquitin__system | DNA__repair__and__recombination__proteins | 0.0185185 |
| Drosophila | omegaNA | Ubiquitin__system | DNA__replication__proteins | 0.0185185 |
| Drosophila | omegaNA | Ubiquitin__system | Exosome | 0.4259259 |
| Drosophila | omegaNA | Ubiquitin__system | G__protein__coupled__receptors | 0.4259259 |
| Drosophila | omegaNA | Ubiquitin__system | Glycosyltransferases | 0.1666667 |
| Drosophila | omegaNA | Ubiquitin__system | Ion__channels | 0.3518519 |
| Drosophila | omegaNA | Ubiquitin__system | Lipid__biosynthesis__proteins | 0.3888889 |
| Drosophila | omegaNA | Ubiquitin__system | Membrane__trafficking | 0.0925926 |
| Drosophila | omegaNA | Ubiquitin__system | Messenger__RNA__biogenesis | 0.9444444 |
| Drosophila | omegaNA | Ubiquitin__system | Mitochondrial__biogenesis | 0.9814815 |
| Drosophila | omegaNA | Ubiquitin__system | Peptidases | 0.0185185 |
| Drosophila | omegaNA | Ubiquitin__system | Photosynthesis__proteins | 0.8703704 |
| Drosophila | omegaNA | Ubiquitin__system | Proteasome | 0.9444444 |
| Drosophila | omegaNA | Ubiquitin__system | Protein__kinases | 0.1296296 |
| Drosophila | omegaNA | Ubiquitin__system | Protein__phosphatases__and__associated__proteins | 0.6851852 |
| Drosophila | omegaNA | Ubiquitin__system | Ribosome | 0.8333333 |
| Drosophila | omegaNA | Ubiquitin__system | Ribosome__biogenesis | 0.4629630 |
| Drosophila | omegaNA | Ubiquitin__system | Spliceosome | 0.3148148 |
| Drosophila | omegaNA | Ubiquitin__system | Transcription__factors | 0.0925926 |
| Drosophila | omegaNA | Ubiquitin__system | Transcription__machinery | 0.2037037 |
| Drosophila | omegaNA | Ubiquitin__system | Transfer__RNA__biogenesis | 0.6481481 |
| Drosophila | omegaNA | Ubiquitin__system | Translation__factors | 0.6481481 |
| Drosophila | omegaNA | Ubiquitin__system | Transporters | 0.6851852 |
| Drosophila | omegaA | Chaperones__and__folding__catalysts | Amino__acid__related__enzymes | 0.7962963 |
| Drosophila | omegaA | Chromosome__and__associated__proteins | Amino__acid__related__enzymes | 0.9814815 |
| Drosophila | omegaA | Chromosome__and__associated__proteins | Chaperones__and__folding__catalysts | 0.2037037 |
| Drosophila | omegaA | Cytochrome__P450 | Amino__acid__related__enzymes | 0.9074074 |
| Drosophila | omegaA | Cytochrome__P450 | Chaperones__and__folding__catalysts | 0.3148148 |
| Drosophila | omegaA | Cytochrome__P450 | Chromosome__and__associated__proteins | 0.8703704 |
| Drosophila | omegaA | Cytoskeleton__proteins | Amino__acid__related__enzymes | 0.5370370 |
| Drosophila | omegaA | Cytoskeleton__proteins | Chaperones__and__folding__catalysts | 0.7962963 |
| Drosophila | omegaA | Cytoskeleton__proteins | Chromosome__and__associated__proteins | 0.0185185 |
| Drosophila | omegaA | Cytoskeleton__proteins | Cytochrome__P450 | 0.0925926 |
| Drosophila | omegaA | DNA__repair__and__recombination__proteins | Amino__acid__related__enzymes | 0.8703704 |
| Drosophila | omegaA | DNA__repair__and__recombination__proteins | Chaperones__and__folding__catalysts | 0.2777778 |
| Drosophila | omegaA | DNA__repair__and__recombination__proteins | Chromosome__and__associated__proteins | 0.6481481 |
| Drosophila | omegaA | DNA__repair__and__recombination__proteins | Cytochrome__P450 | 0.9444444 |
| Drosophila | omegaA | DNA__repair__and__recombination__proteins | Cytoskeleton__proteins | 0.1296296 |
| Drosophila | omegaA | DNA__replication__proteins | Amino__acid__related__enzymes | 0.4259259 |
| Drosophila | omegaA | DNA__replication__proteins | Chaperones__and__folding__catalysts | 0.1666667 |
| Drosophila | omegaA | DNA__replication__proteins | Chromosome__and__associated__proteins | 0.0185185 |
| Drosophila | omegaA | DNA__replication__proteins | Cytochrome__P450 | 0.0555556 |
| Drosophila | omegaA | DNA__replication__proteins | Cytoskeleton__proteins | 0.1666667 |
| Drosophila | omegaA | DNA__replication__proteins | DNA__repair__and__recombination__proteins | 0.0185185 |
| Drosophila | omegaA | Exosome | Amino__acid__related__enzymes | 0.6851852 |
| Drosophila | omegaA | Exosome | Chaperones__and__folding__catalysts | 0.6111111 |
| Drosophila | omegaA | Exosome | Chromosome__and__associated__proteins | 0.0185185 |
| Drosophila | omegaA | Exosome | Cytochrome__P450 | 0.0925926 |
| Drosophila | omegaA | Exosome | Cytoskeleton__proteins | 0.8703704 |
| Drosophila | omegaA | Exosome | DNA__repair__and__recombination__proteins | 0.0925926 |
| Drosophila | omegaA | Exosome | DNA__replication__proteins | 0.0925926 |
| Drosophila | omegaA | G__protein__coupled__receptors | Amino__acid__related__enzymes | 0.6481481 |
| Drosophila | omegaA | G__protein__coupled__receptors | Chaperones__and__folding__catalysts | 0.1666667 |
| Drosophila | omegaA | G__protein__coupled__receptors | Chromosome__and__associated__proteins | 0.9444444 |

*(continued)*

| species | estimate | var1 | var2 | p.value |
|---|---|---|---|---|
| Drosophila | omegaA | G__protein__coupled__receptors | Cytochrome__P450 | 0.9444444 |
| Drosophila | omegaA | G__protein__coupled__receptors | Cytoskeleton__proteins | 0.1666667 |
| Drosophila | omegaA | G__protein__coupled__receptors | DNA__repair__and__recombination__proteins | 0.9444444 |
| Drosophila | omegaA | G__protein__coupled__receptors | DNA__replication__proteins | 0.0185185 |
| Drosophila | omegaA | G__protein__coupled__receptors | Exosome | 0.0185185 |
| Drosophila | omegaA | Glycosyltransferases | Amino__acid__related__enzymes | 0.3888889 |
| Drosophila | omegaA | Glycosyltransferases | Chaperones__and__folding__catalysts | 0.3148148 |
| Drosophila | omegaA | Glycosyltransferases | Chromosome__and__associated__proteins | 0.0555556 |
| Drosophila | omegaA | Glycosyltransferases | Cytochrome__P450 | 0.0555556 |
| Drosophila | omegaA | Glycosyltransferases | Cytoskeleton__proteins | 0.3518519 |
| Drosophila | omegaA | Glycosyltransferases | DNA__repair__and__recombination__proteins | 0.0555556 |
| Drosophila | omegaA | Glycosyltransferases | DNA__replication__proteins | 0.5740741 |
| Drosophila | omegaA | Glycosyltransferases | Exosome | 0.2777778 |
| Drosophila | omegaA | Glycosyltransferases | G__protein__coupled__receptors | 0.0185185 |
| Drosophila | omegaA | Ion__channels | Amino__acid__related__enzymes | 0.7222222 |
| Drosophila | omegaA | Ion__channels | Chaperones__and__folding__catalysts | 0.7222222 |
| Drosophila | omegaA | Ion__channels | Chromosome__and__associated__proteins | 0.0555556 |
| Drosophila | omegaA | Ion__channels | Cytochrome__P450 | 0.1666667 |
| Drosophila | omegaA | Ion__channels | Cytoskeleton__proteins | 0.3148148 |
| Drosophila | omegaA | Ion__channels | DNA__repair__and__recombination__proteins | 0.2777778 |
| Drosophila | omegaA | Ion__channels | DNA__replication__proteins | 0.0185185 |
| Drosophila | omegaA | Ion__channels | Exosome | 0.1296296 |
| Drosophila | omegaA | Ion__channels | G__protein__coupled__receptors | 0.4259259 |
| Drosophila | omegaA | Ion__channels | Glycosyltransferases | 0.0555556 |
| Drosophila | omegaA | Lipid__biosynthesis__proteins | Amino__acid__related__enzymes | 0.9074074 |
| Drosophila | omegaA | Lipid__biosynthesis__proteins | Chaperones__and__folding__catalysts | 0.2037037 |
| Drosophila | omegaA | Lipid__biosynthesis__proteins | Chromosome__and__associated__proteins | 0.7962963 |
| Drosophila | omegaA | Lipid__biosynthesis__proteins | Cytochrome__P450 | 0.9074074 |
| Drosophila | omegaA | Lipid__biosynthesis__proteins | Cytoskeleton__proteins | 0.0185185 |
| Drosophila | omegaA | Lipid__biosynthesis__proteins | DNA__repair__and__recombination__proteins | 0.9814815 |
| Drosophila | omegaA | Lipid__biosynthesis__proteins | DNA__replication__proteins | 0.0185185 |
| Drosophila | omegaA | Lipid__biosynthesis__proteins | Exosome | 0.0185185 |
| Drosophila | omegaA | Lipid__biosynthesis__proteins | G__protein__coupled__receptors | 0.9814815 |
| Drosophila | omegaA | Lipid__biosynthesis__proteins | Glycosyltransferases | 0.0185185 |
| Drosophila | omegaA | Lipid__biosynthesis__proteins | Ion__channels | 0.2407407 |
| Drosophila | omegaA | Membrane__trafficking | Amino__acid__related__enzymes | 0.7222222 |
| Drosophila | omegaA | Membrane__trafficking | Chaperones__and__folding__catalysts | 0.7592593 |
| Drosophila | omegaA | Membrane__trafficking | Chromosome__and__associated__proteins | 0.0185185 |
| Drosophila | omegaA | Membrane__trafficking | Cytochrome__P450 | 0.2037037 |
| Drosophila | omegaA | Membrane__trafficking | Cytoskeleton__proteins | 0.3888889 |
| Drosophila | omegaA | Membrane__trafficking | DNA__repair__and__recombination__proteins | 0.1296296 |
| Drosophila | omegaA | Membrane__trafficking | DNA__replication__proteins | 0.0185185 |
| Drosophila | omegaA | Membrane__trafficking | Exosome | 0.2777778 |
| Drosophila | omegaA | Membrane__trafficking | G__protein__coupled__receptors | 0.0555556 |
| Drosophila | omegaA | Membrane__trafficking | Glycosyltransferases | 0.0925926 |
| Drosophila | omegaA | Membrane__trafficking | Ion__channels | 0.8333333 |
| Drosophila | omegaA | Membrane__trafficking | Lipid__biosynthesis__proteins | 0.1296296 |
| Drosophila | omegaA | Messenger__RNA__biogenesis | Amino__acid__related__enzymes | 0.2777778 |
| Drosophila | omegaA | Messenger__RNA__biogenesis | Chaperones__and__folding__catalysts | 0.0185185 |
| Drosophila | omegaA | Messenger__RNA__biogenesis | Chromosome__and__associated__proteins | 0.0925926 |
| Drosophila | omegaA | Messenger__RNA__biogenesis | Cytochrome__P450 | 0.2777778 |
| Drosophila | omegaA | Messenger__RNA__biogenesis | Cytoskeleton__proteins | 0.0185185 |
| Drosophila | omegaA | Messenger__RNA__biogenesis | DNA__repair__and__recombination__proteins | 0.0185185 |
| Drosophila | omegaA | Messenger__RNA__biogenesis | DNA__replication__proteins | 0.0185185 |
| Drosophila | omegaA | Messenger__RNA__biogenesis | Exosome | 0.0185185 |
| Drosophila | omegaA | Messenger__RNA__biogenesis | G__protein__coupled__receptors | 0.0925926 |
| Drosophila | omegaA | Messenger__RNA__biogenesis | Glycosyltransferases | 0.0185185 |
| Drosophila | omegaA | Messenger__RNA__biogenesis | Ion__channels | 0.0185185 |
| Drosophila | omegaA | Messenger__RNA__biogenesis | Lipid__biosynthesis__proteins | 0.2037037 |
| Drosophila | omegaA | Messenger__RNA__biogenesis | Membrane__trafficking | 0.0185185 |
| Drosophila | omegaA | Mitochondrial__biogenesis | Amino__acid__related__enzymes | 0.6111111 |
| Drosophila | omegaA | Mitochondrial__biogenesis | Chaperones__and__folding__catalysts | 0.9814815 |

*(continued)*

| species | estimate | var1 | var2 | p.value |
|---|---|---|---|---|
| Drosophila | omegaA | Mitochondrial__biogenesis | Chromosome__and__associated__proteins | 0.1666667 |
| Drosophila | omegaA | Mitochondrial__biogenesis | Cytochrome__P450 | 0.3518519 |
| Drosophila | omegaA | Mitochondrial__biogenesis | Cytoskeleton__proteins | 0.8333333 |
| Drosophila | omegaA | Mitochondrial__biogenesis | DNA__repair__and__recombination__proteins | 0.3888889 |
| Drosophila | omegaA | Mitochondrial__biogenesis | DNA__replication__proteins | 0.3888889 |
| Drosophila | omegaA | Mitochondrial__biogenesis | Exosome | 0.7222222 |
| Drosophila | omegaA | Mitochondrial__biogenesis | G__protein__coupled__receptors | 0.3518519 |
| Drosophila | omegaA | Mitochondrial__biogenesis | Glycosyltransferases | 0.4259259 |
| Drosophila | omegaA | Mitochondrial__biogenesis | Ion__channels | 0.9074074 |
| Drosophila | omegaA | Mitochondrial__biogenesis | Lipid__biosynthesis__proteins | 0.3518519 |
| Drosophila | omegaA | Mitochondrial__biogenesis | Membrane__trafficking | 0.9444444 |
| Drosophila | omegaA | Mitochondrial__biogenesis | Messenger__RNA__biogenesis | 0.0185185 |
| Drosophila | omegaA | Peptidases | Amino__acid__related__enzymes | 0.4259259 |
| Drosophila | omegaA | Peptidases | Chaperones__and__folding__catalysts | 0.0555556 |
| Drosophila | omegaA | Peptidases | Chromosome__and__associated__proteins | 0.2037037 |
| Drosophila | omegaA | Peptidases | Cytochrome__P450 | 0.4259259 |
| Drosophila | omegaA | Peptidases | Cytoskeleton__proteins | 0.0185185 |
| Drosophila | omegaA | Peptidases | DNA__repair__and__recombination__proteins | 0.0555556 |
| Drosophila | omegaA | Peptidases | DNA__replication__proteins | 0.0185185 |
| Drosophila | omegaA | Peptidases | Exosome | 0.0185185 |
| Drosophila | omegaA | Peptidases | G__protein__coupled__receptors | 0.3518519 |
| Drosophila | omegaA | Peptidases | Glycosyltransferases | 0.0185185 |
| Drosophila | omegaA | Peptidases | Ion__channels | 0.0185185 |
| Drosophila | omegaA | Peptidases | Lipid__biosynthesis__proteins | 0.2037037 |
| Drosophila | omegaA | Peptidases | Membrane__trafficking | 0.0185185 |
| Drosophila | omegaA | Peptidases | Messenger__RNA__biogenesis | 0.8333333 |
| Drosophila | omegaA | Peptidases | Mitochondrial__biogenesis | 0.0555556 |
| Drosophila | omegaA | Photosynthesis__proteins | Amino__acid__related__enzymes | 0.7962963 |
| Drosophila | omegaA | Photosynthesis__proteins | Chaperones__and__folding__catalysts | 0.6481481 |
| Drosophila | omegaA | Photosynthesis__proteins | Chromosome__and__associated__proteins | 0.3518519 |
| Drosophila | omegaA | Photosynthesis__proteins | Cytochrome__P450 | 0.3888889 |
| Drosophila | omegaA | Photosynthesis__proteins | Cytoskeleton__proteins | 0.5740741 |
| Drosophila | omegaA | Photosynthesis__proteins | DNA__repair__and__recombination__proteins | 0.5370370 |
| Drosophila | omegaA | Photosynthesis__proteins | DNA__replication__proteins | 0.1296296 |
| Drosophila | omegaA | Photosynthesis__proteins | Exosome | 0.5000000 |
| Drosophila | omegaA | Photosynthesis__proteins | G__protein__coupled__receptors | 0.3888889 |
| Drosophila | omegaA | Photosynthesis__proteins | Glycosyltransferases | 0.2777778 |
| Drosophila | omegaA | Photosynthesis__proteins | Ion__channels | 0.7962963 |
| Drosophila | omegaA | Photosynthesis__proteins | Lipid__biosynthesis__proteins | 0.4629630 |
| Drosophila | omegaA | Photosynthesis__proteins | Membrane__trafficking | 0.7962963 |
| Drosophila | omegaA | Photosynthesis__proteins | Messenger__RNA__biogenesis | 0.0555556 |
| Drosophila | omegaA | Photosynthesis__proteins | Mitochondrial__biogenesis | 0.7592593 |
| Drosophila | omegaA | Photosynthesis__proteins | Peptidases | 0.0925926 |
| Drosophila | omegaA | Proteasome | Amino__acid__related__enzymes | 0.7592593 |
| Drosophila | omegaA | Proteasome | Chaperones__and__folding__catalysts | 0.5370370 |
| Drosophila | omegaA | Proteasome | Chromosome__and__associated__proteins | 0.4259259 |
| Drosophila | omegaA | Proteasome | Cytochrome__P450 | 0.6111111 |
| Drosophila | omegaA | Proteasome | Cytoskeleton__proteins | 0.5740741 |
| Drosophila | omegaA | Proteasome | DNA__repair__and__recombination__proteins | 0.5740741 |
| Drosophila | omegaA | Proteasome | DNA__replication__proteins | 0.2037037 |
| Drosophila | omegaA | Proteasome | Exosome | 0.4629630 |
| Drosophila | omegaA | Proteasome | G__protein__coupled__receptors | 0.3518519 |
| Drosophila | omegaA | Proteasome | Glycosyltransferases | 0.2037037 |
| Drosophila | omegaA | Proteasome | Ion__channels | 0.9074074 |
| Drosophila | omegaA | Proteasome | Lipid__biosynthesis__proteins | 0.4629630 |
| Drosophila | omegaA | Proteasome | Membrane__trafficking | 0.6851852 |
| Drosophila | omegaA | Proteasome | Messenger__RNA__biogenesis | 0.0555556 |
| Drosophila | omegaA | Proteasome | Mitochondrial__biogenesis | 0.7592593 |
| Drosophila | omegaA | Proteasome | Peptidases | 0.0555556 |
| Drosophila | omegaA | Proteasome | Photosynthesis__proteins | 0.9814815 |
| Drosophila | omegaA | Protein__kinases | Amino__acid__related__enzymes | 0.7222222 |

*(continued)*

| species | estimate | var1 | var2 | p.value |
|---|---|---|---|---|
| Drosophila | omegaA | Protein__kinases | Chaperones__and__folding__catalysts | 0.5370370 |
| Drosophila | omegaA | Protein__kinases | Chromosome__and__associated__proteins | 0.0185185 |
| Drosophila | omegaA | Protein__kinases | Cytochrome__P450 | 0.3148148 |
| Drosophila | omegaA | Protein__kinases | Cytoskeleton__proteins | 0.2037037 |
| Drosophila | omegaA | Protein__kinases | DNA__repair__and__recombination__proteins | 0.3148148 |
| Drosophila | omegaA | Protein__kinases | DNA__replication__proteins | 0.0185185 |
| Drosophila | omegaA | Protein__kinases | Exosome | 0.0555556 |
| Drosophila | omegaA | Protein__kinases | G__protein__coupled__receptors | 0.3888889 |
| Drosophila | omegaA | Protein__kinases | Glycosyltransferases | 0.0925926 |
| Drosophila | omegaA | Protein__kinases | Ion__channels | 0.6111111 |
| Drosophila | omegaA | Protein__kinases | Lipid__biosynthesis__proteins | 0.3518519 |
| Drosophila | omegaA | Protein__kinases | Membrane__trafficking | 0.3148148 |
| Drosophila | omegaA | Protein__kinases | Messenger__RNA__biogenesis | 0.0185185 |
| Drosophila | omegaA | Protein__kinases | Mitochondrial__biogenesis | 0.7962963 |
| Drosophila | omegaA | Protein__kinases | Peptidases | 0.0185185 |
| Drosophila | omegaA | Protein__kinases | Photosynthesis__proteins | 0.9814815 |
| Drosophila | omegaA | Protein__kinases | Proteasome | 0.9814815 |
| Drosophila | omegaA | Protein__phosphatases__and__associated__proteins | Amino__acid__related__enzymes | 0.9074074 |
| Drosophila | omegaA | Protein__phosphatases__and__associated__proteins | Chaperones__and__folding__catalysts | 0.1296296 |
| Drosophila | omegaA | Protein__phosphatases__and__associated__proteins | Chromosome__and__associated__proteins | 0.7962963 |
| Drosophila | omegaA | Protein__phosphatases__and__associated__proteins | Cytochrome__P450 | 0.7592593 |
| Drosophila | omegaA | Protein__phosphatases__and__associated__proteins | Cytoskeleton__proteins | 0.1296296 |
| Drosophila | omegaA | Protein__phosphatases__and__associated__proteins | DNA__repair__and__recombination__proteins | 0.6481481 |
| Drosophila | omegaA | Protein__phosphatases__and__associated__proteins | DNA__replication__proteins | 0.0185185 |
| Drosophila | omegaA | Protein__phosphatases__and__associated__proteins | Exosome | 0.0555556 |
| Drosophila | omegaA | Protein__phosphatases__and__associated__proteins | G__protein__coupled__receptors | 0.8333333 |
| Drosophila | omegaA | Protein__phosphatases__and__associated__proteins | Glycosyltransferases | 0.0555556 |
| Drosophila | omegaA | Protein__phosphatases__and__associated__proteins | Ion__channels | 0.2037037 |
| Drosophila | omegaA | Protein__phosphatases__and__associated__proteins | Lipid__biosynthesis__proteins | 0.7962963 |
| Drosophila | omegaA | Protein__phosphatases__and__associated__proteins | Membrane__trafficking | 0.0925926 |
| Drosophila | omegaA | Protein__phosphatases__and__associated__proteins | Messenger__RNA__biogenesis | 0.0555556 |
| Drosophila | omegaA | Protein__phosphatases__and__associated__proteins | Mitochondrial__biogenesis | 0.2777778 |
| Drosophila | omegaA | Protein__phosphatases__and__associated__proteins | Peptidases | 0.4259259 |
| Drosophila | omegaA | Protein__phosphatases__and__associated__proteins | Photosynthesis__proteins | 0.4259259 |
| Drosophila | omegaA | Protein__phosphatases__and__associated__proteins | Proteasome | 0.3518519 |
| Drosophila | omegaA | Protein__phosphatases__and__associated__proteins | Protein__kinases | 0.2407407 |
| Drosophila | omegaA | Ribosome | Amino__acid__related__enzymes | 0.6481481 |
| Drosophila | omegaA | Ribosome | Chaperones__and__folding__catalysts | 0.8333333 |
| Drosophila | omegaA | Ribosome | Chromosome__and__associated__proteins | 0.2407407 |
| Drosophila | omegaA | Ribosome | Cytochrome__P450 | 0.3148148 |
| Drosophila | omegaA | Ribosome | Cytoskeleton__proteins | 0.7222222 |
| Drosophila | omegaA | Ribosome | DNA__repair__and__recombination__proteins | 0.3518519 |
| Drosophila | omegaA | Ribosome | DNA__replication__proteins | 0.2037037 |
| Drosophila | omegaA | Ribosome | Exosome | 0.6111111 |
| Drosophila | omegaA | Ribosome | G__protein__coupled__receptors | 0.3148148 |
| Drosophila | omegaA | Ribosome | Glycosyltransferases | 0.3518519 |
| Drosophila | omegaA | Ribosome | Ion__channels | 0.8703704 |
| Drosophila | omegaA | Ribosome | Lipid__biosynthesis__proteins | 0.2407407 |
| Drosophila | omegaA | Ribosome | Membrane__trafficking | 0.9074074 |
| Drosophila | omegaA | Ribosome | Messenger__RNA__biogenesis | 0.0185185 |
| Drosophila | omegaA | Ribosome | Mitochondrial__biogenesis | 0.9444444 |
| Drosophila | omegaA | Ribosome | Peptidases | 0.0185185 |
| Drosophila | omegaA | Ribosome | Photosynthesis__proteins | 0.7592593 |
| Drosophila | omegaA | Ribosome | Proteasome | 0.7592593 |
| Drosophila | omegaA | Ribosome | Protein__kinases | 0.7962963 |
| Drosophila | omegaA | Ribosome | Protein__phosphatases__and__associated__proteins | 0.2777778 |
| Drosophila | omegaA | Ribosome__biogenesis | Amino__acid__related__enzymes | 0.6851852 |
| Drosophila | omegaA | Ribosome__biogenesis | Chaperones__and__folding__catalysts | 0.0925926 |
| Drosophila | omegaA | Ribosome__biogenesis | Chromosome__and__associated__proteins | 0.3888889 |
| Drosophila | omegaA | Ribosome__biogenesis | Cytochrome__P450 | 0.5000000 |
| Drosophila | omegaA | Ribosome__biogenesis | Cytoskeleton__proteins | 0.0185185 |
| Drosophila | omegaA | Ribosome__biogenesis | DNA__repair__and__recombination__proteins | 0.1666667 |

*(continued)*

| species | estimate | var1 | var2 | p.value |
|---|---|---|---|---|
| Drosophila | omegaA | Ribosome__biogenesis | DNA__replication__proteins | 0.0185185 |
| Drosophila | omegaA | Ribosome__biogenesis | Exosome | 0.0185185 |
| Drosophila | omegaA | Ribosome__biogenesis | G__protein__coupled__receptors | 0.7222222 |
| Drosophila | omegaA | Ribosome__biogenesis | Glycosyltransferases | 0.0185185 |
| Drosophila | omegaA | Ribosome__biogenesis | Ion__channels | 0.0555556 |
| Drosophila | omegaA | Ribosome__biogenesis | Lipid__biosynthesis__proteins | 0.3888889 |
| Drosophila | omegaA | Ribosome__biogenesis | Membrane__trafficking | 0.0185185 |
| Drosophila | omegaA | Ribosome__biogenesis | Messenger__RNA__biogenesis | 0.0555556 |
| Drosophila | omegaA | Ribosome__biogenesis | Mitochondrial__biogenesis | 0.0185185 |
| Drosophila | omegaA | Ribosome__biogenesis | Peptidases | 0.4259259 |
| Drosophila | omegaA | Ribosome__biogenesis | Photosynthesis__proteins | 0.0925926 |
| Drosophila | omegaA | Ribosome__biogenesis | Proteasome | 0.0925926 |
| Drosophila | omegaA | Ribosome__biogenesis | Protein__kinases | 0.0555556 |
| Drosophila | omegaA | Ribosome__biogenesis | Protein__phosphatases__and__associated__proteins | 0.6481481 |
| Drosophila | omegaA | Ribosome__biogenesis | Ribosome | 0.0555556 |
| Drosophila | omegaA | Spliceosome | Amino__acid__related__enzymes | 0.7222222 |
| Drosophila | omegaA | Spliceosome | Chaperones__and__folding__catalysts | 0.6111111 |
| Drosophila | omegaA | Spliceosome | Chromosome__and__associated__proteins | 0.2037037 |
| Drosophila | omegaA | Spliceosome | Cytochrome__P450 | 0.3148148 |
| Drosophila | omegaA | Spliceosome | Cytoskeleton__proteins | 0.3148148 |
| Drosophila | omegaA | Spliceosome | DNA__repair__and__recombination__proteins | 0.3888889 |
| Drosophila | omegaA | Spliceosome | DNA__replication__proteins | 0.0185185 |
| Drosophila | omegaA | Spliceosome | Exosome | 0.1666667 |
| Drosophila | omegaA | Spliceosome | G__protein__coupled__receptors | 0.5740741 |
| Drosophila | omegaA | Spliceosome | Glycosyltransferases | 0.0185185 |
| Drosophila | omegaA | Spliceosome | Ion__channels | 0.8333333 |
| Drosophila | omegaA | Spliceosome | Lipid__biosynthesis__proteins | 0.3148148 |
| Drosophila | omegaA | Spliceosome | Membrane__trafficking | 0.6111111 |
| Drosophila | omegaA | Spliceosome | Messenger__RNA__biogenesis | 0.0185185 |
| Drosophila | omegaA | Spliceosome | Mitochondrial__biogenesis | 0.8333333 |
| Drosophila | omegaA | Spliceosome | Peptidases | 0.0185185 |
| Drosophila | omegaA | Spliceosome | Photosynthesis__proteins | 0.9444444 |
| Drosophila | omegaA | Spliceosome | Proteasome | 0.9814815 |
| Drosophila | omegaA | Spliceosome | Protein__kinases | 0.9074074 |
| Drosophila | omegaA | Spliceosome | Protein__phosphatases__and__associated__proteins | 0.2037037 |
| Drosophila | omegaA | Spliceosome | Ribosome | 0.9814815 |
| Drosophila | omegaA | Spliceosome | Ribosome__biogenesis | 0.0555556 |
| Drosophila | omegaA | Transcription__factors | Amino__acid__related__enzymes | 0.7222222 |
| Drosophila | omegaA | Transcription__factors | Chaperones__and__folding__catalysts | 0.7222222 |
| Drosophila | omegaA | Transcription__factors | Chromosome__and__associated__proteins | 0.0925926 |
| Drosophila | omegaA | Transcription__factors | Cytochrome__P450 | 0.2037037 |
| Drosophila | omegaA | Transcription__factors | Cytoskeleton__proteins | 0.3888889 |
| Drosophila | omegaA | Transcription__factors | DNA__repair__and__recombination__proteins | 0.2407407 |
| Drosophila | omegaA | Transcription__factors | DNA__replication__proteins | 0.0185185 |
| Drosophila | omegaA | Transcription__factors | Exosome | 0.2407407 |
| Drosophila | omegaA | Transcription__factors | G__protein__coupled__receptors | 0.3518519 |
| Drosophila | omegaA | Transcription__factors | Glycosyltransferases | 0.1296296 |
| Drosophila | omegaA | Transcription__factors | Ion__channels | 0.9444444 |
| Drosophila | omegaA | Transcription__factors | Lipid__biosynthesis__proteins | 0.2407407 |
| Drosophila | omegaA | Transcription__factors | Membrane__trafficking | 0.7592593 |
| Drosophila | omegaA | Transcription__factors | Messenger__RNA__biogenesis | 0.0185185 |
| Drosophila | omegaA | Transcription__factors | Mitochondrial__biogenesis | 0.9444444 |
| Drosophila | omegaA | Transcription__factors | Peptidases | 0.0185185 |
| Drosophila | omegaA | Transcription__factors | Photosynthesis__proteins | 0.8333333 |
| Drosophila | omegaA | Transcription__factors | Proteasome | 0.8333333 |
| Drosophila | omegaA | Transcription__factors | Protein__kinases | 0.4629630 |
| Drosophila | omegaA | Transcription__factors | Protein__phosphatases__and__associated__proteins | 0.2037037 |
| Drosophila | omegaA | Transcription__factors | Ribosome | 0.9444444 |
| Drosophila | omegaA | Transcription__factors | Ribosome__biogenesis | 0.0185185 |
| Drosophila | omegaA | Transcription__factors | Spliceosome | 0.9444444 |
| Drosophila | omegaA | Transcription__machinery | Amino__acid__related__enzymes | 0.7592593 |
| Drosophila | omegaA | Transcription__machinery | Chaperones__and__folding__catalysts | 0.4259259 |

*(continued)*

| species | estimate | var1 | var2 | p.value |
|---|---|---|---|---|
| Drosophila | omegaA | Transcription__machinery | Chromosome__and__associated__proteins | 0.3888889 |
| Drosophila | omegaA | Transcription__machinery | Cytochrome__P450 | 0.5740741 |
| Drosophila | omegaA | Transcription__machinery | Cytoskeleton__proteins | 0.2037037 |
| Drosophila | omegaA | Transcription__machinery | DNA__repair__and__recombination__proteins | 0.6481481 |
| Drosophila | omegaA | Transcription__machinery | DNA__replication__proteins | 0.0185185 |
| Drosophila | omegaA | Transcription__machinery | Exosome | 0.0185185 |
| Drosophila | omegaA | Transcription__machinery | G__protein__coupled__receptors | 0.5370370 |
| Drosophila | omegaA | Transcription__machinery | Glycosyltransferases | 0.0185185 |
| Drosophila | omegaA | Transcription__machinery | Ion__channels | 0.3888889 |
| Drosophila | omegaA | Transcription__machinery | Lipid__biosynthesis__proteins | 0.6111111 |
| Drosophila | omegaA | Transcription__machinery | Membrane__trafficking | 0.1666667 |
| Drosophila | omegaA | Transcription__machinery | Messenger__RNA__biogenesis | 0.0185185 |
| Drosophila | omegaA | Transcription__machinery | Mitochondrial__biogenesis | 0.4629630 |
| Drosophila | omegaA | Transcription__machinery | Peptidases | 0.0925926 |
| Drosophila | omegaA | Transcription__machinery | Photosynthesis__proteins | 0.8333333 |
| Drosophila | omegaA | Transcription__machinery | Proteasome | 0.8703704 |
| Drosophila | omegaA | Transcription__machinery | Protein__kinases | 0.5000000 |
| Drosophila | omegaA | Transcription__machinery | Protein__phosphatases__and__associated__proteins | 0.4629630 |
| Drosophila | omegaA | Transcription__machinery | Ribosome | 0.6111111 |
| Drosophila | omegaA | Transcription__machinery | Ribosome__biogenesis | 0.0925926 |
| Drosophila | omegaA | Transcription__machinery | Spliceosome | 0.6851852 |
| Drosophila | omegaA | Transcription__machinery | Transcription__factors | 0.2777778 |
| Drosophila | omegaA | Transfer__RNA__biogenesis | Amino__acid__related__enzymes | 0.7222222 |
| Drosophila | omegaA | Transfer__RNA__biogenesis | Chaperones__and__folding__catalysts | 0.8703704 |
| Drosophila | omegaA | Transfer__RNA__biogenesis | Chromosome__and__associated__proteins | 0.0555556 |
| Drosophila | omegaA | Transfer__RNA__biogenesis | Cytochrome__P450 | 0.1666667 |
| Drosophila | omegaA | Transfer__RNA__biogenesis | Cytoskeleton__proteins | 0.5000000 |
| Drosophila | omegaA | Transfer__RNA__biogenesis | DNA__repair__and__recombination__proteins | 0.2407407 |
| Drosophila | omegaA | Transfer__RNA__biogenesis | DNA__replication__proteins | 0.0925926 |
| Drosophila | omegaA | Transfer__RNA__biogenesis | Exosome | 0.2407407 |
| Drosophila | omegaA | Transfer__RNA__biogenesis | G__protein__coupled__receptors | 0.2037037 |
| Drosophila | omegaA | Transfer__RNA__biogenesis | Glycosyltransferases | 0.1666667 |
| Drosophila | omegaA | Transfer__RNA__biogenesis | Ion__channels | 0.6851852 |
| Drosophila | omegaA | Transfer__RNA__biogenesis | Lipid__biosynthesis__proteins | 0.1666667 |
| Drosophila | omegaA | Transfer__RNA__biogenesis | Membrane__trafficking | 0.9074074 |
| Drosophila | omegaA | Transfer__RNA__biogenesis | Messenger__RNA__biogenesis | 0.0185185 |
| Drosophila | omegaA | Transfer__RNA__biogenesis | Mitochondrial__biogenesis | 0.9444444 |
| Drosophila | omegaA | Transfer__RNA__biogenesis | Peptidases | 0.0185185 |
| Drosophila | omegaA | Transfer__RNA__biogenesis | Photosynthesis__proteins | 0.7962963 |
| Drosophila | omegaA | Transfer__RNA__biogenesis | Proteasome | 0.6481481 |
| Drosophila | omegaA | Transfer__RNA__biogenesis | Protein__kinases | 0.3148148 |
| Drosophila | omegaA | Transfer__RNA__biogenesis | Protein__phosphatases__and__associated__proteins | 0.2037037 |
| Drosophila | omegaA | Transfer__RNA__biogenesis | Ribosome | 0.9814815 |
| Drosophila | omegaA | Transfer__RNA__biogenesis | Ribosome__biogenesis | 0.0185185 |
| Drosophila | omegaA | Transfer__RNA__biogenesis | Spliceosome | 0.6481481 |
| Drosophila | omegaA | Transfer__RNA__biogenesis | Transcription__factors | 0.7962963 |
| Drosophila | omegaA | Transfer__RNA__biogenesis | Transcription__machinery | 0.2407407 |
| Drosophila | omegaA | Translation__factors | Amino__acid__related__enzymes | 0.5740741 |
| Drosophila | omegaA | Translation__factors | Chaperones__and__folding__catalysts | 0.3148148 |
| Drosophila | omegaA | Translation__factors | Chromosome__and__associated__proteins | 0.9444444 |
| Drosophila | omegaA | Translation__factors | Cytochrome__P450 | 0.8333333 |
| Drosophila | omegaA | Translation__factors | Cytoskeleton__proteins | 0.0925926 |
| Drosophila | omegaA | Translation__factors | DNA__repair__and__recombination__proteins | 0.7592593 |
| Drosophila | omegaA | Translation__factors | DNA__replication__proteins | 0.0185185 |
| Drosophila | omegaA | Translation__factors | Exosome | 0.0925926 |
| Drosophila | omegaA | Translation__factors | G__protein__coupled__receptors | 0.8703704 |
| Drosophila | omegaA | Translation__factors | Glycosyltransferases | 0.0555556 |
| Drosophila | omegaA | Translation__factors | Ion__channels | 0.3518519 |
| Drosophila | omegaA | Translation__factors | Lipid__biosynthesis__proteins | 0.6851852 |
| Drosophila | omegaA | Translation__factors | Membrane__trafficking | 0.2407407 |
| Drosophila | omegaA | Translation__factors | Messenger__RNA__biogenesis | 0.3888889 |

*(continued)*

| species | estimate | var1 | var2 | p.value |
|---|---|---|---|---|
| Drosophila | omegaA | Translation__factors | Mitochondrial__biogenesis | 0.2777778 |
| Drosophila | omegaA | Translation__factors | Peptidases | 0.5000000 |
| Drosophila | omegaA | Translation__factors | Photosynthesis__proteins | 0.2777778 |
| Drosophila | omegaA | Translation__factors | Proteasome | 0.3148148 |
| Drosophila | omegaA | Translation__factors | Protein__kinases | 0.5000000 |
| Drosophila | omegaA | Translation__factors | Protein__phosphatases__and__associated__proteins | 0.9074074 |
| Drosophila | omegaA | Translation__factors | Ribosome | 0.2407407 |
| Drosophila | omegaA | Translation__factors | Ribosome__biogenesis | 0.8703704 |
| Drosophila | omegaA | Translation__factors | Spliceosome | 0.4259259 |
| Drosophila | omegaA | Translation__factors | Transcription__factors | 0.3888889 |
| Drosophila | omegaA | Translation__factors | Transcription__machinery | 0.6111111 |
| Drosophila | omegaA | Translation__factors | Transfer__RNA__biogenesis | 0.2777778 |
| Drosophila | omegaA | Transporters | Amino__acid__related__enzymes | 0.7222222 |
| Drosophila | omegaA | Transporters | Chaperones__and__folding__catalysts | 0.8333333 |
| Drosophila | omegaA | Transporters | Chromosome__and__associated__proteins | 0.0925926 |
| Drosophila | omegaA | Transporters | Cytochrome__P450 | 0.2407407 |
| Drosophila | omegaA | Transporters | Cytoskeleton__proteins | 0.4629630 |
| Drosophila | omegaA | Transporters | DNA__repair__and__recombination__proteins | 0.2037037 |
| Drosophila | omegaA | Transporters | DNA__replication__proteins | 0.0185185 |
| Drosophila | omegaA | Transporters | Exosome | 0.2407407 |
| Drosophila | omegaA | Transporters | G__protein__coupled__receptors | 0.2407407 |
| Drosophila | omegaA | Transporters | Glycosyltransferases | 0.0925926 |
| Drosophila | omegaA | Transporters | Ion__channels | 0.7592593 |
| Drosophila | omegaA | Transporters | Lipid__biosynthesis__proteins | 0.1666667 |
| Drosophila | omegaA | Transporters | Membrane__trafficking | 0.9814815 |
| Drosophila | omegaA | Transporters | Messenger__RNA__biogenesis | 0.0185185 |
| Drosophila | omegaA | Transporters | Mitochondrial__biogenesis | 0.9444444 |
| Drosophila | omegaA | Transporters | Peptidases | 0.0185185 |
| Drosophila | omegaA | Transporters | Photosynthesis__proteins | 0.7222222 |
| Drosophila | omegaA | Transporters | Proteasome | 0.7592593 |
| Drosophila | omegaA | Transporters | Protein__kinases | 0.4259259 |
| Drosophila | omegaA | Transporters | Protein__phosphatases__and__associated__proteins | 0.0555556 |
| Drosophila | omegaA | Transporters | Ribosome | 0.9444444 |
| Drosophila | omegaA | Transporters | Ribosome__biogenesis | 0.0555556 |
| Drosophila | omegaA | Transporters | Spliceosome | 0.6851852 |
| Drosophila | omegaA | Transporters | Transcription__factors | 0.6111111 |
| Drosophila | omegaA | Transporters | Transcription__machinery | 0.2777778 |
| Drosophila | omegaA | Transporters | Transfer__RNA__biogenesis | 0.9814815 |
| Drosophila | omegaA | Transporters | Translation__factors | 0.2407407 |
| Drosophila | omegaA | Ubiquitin__system | Amino__acid__related__enzymes | 0.8333333 |
| Drosophila | omegaA | Ubiquitin__system | Chaperones__and__folding__catalysts | 0.3148148 |
| Drosophila | omegaA | Ubiquitin__system | Chromosome__and__associated__proteins | 0.5370370 |
| Drosophila | omegaA | Ubiquitin__system | Cytochrome__P450 | 0.7592593 |
| Drosophila | omegaA | Ubiquitin__system | Cytoskeleton__proteins | 0.1296296 |
| Drosophila | omegaA | Ubiquitin__system | DNA__repair__and__recombination__proteins | 0.7592593 |
| Drosophila | omegaA | Ubiquitin__system | DNA__replication__proteins | 0.0185185 |
| Drosophila | omegaA | Ubiquitin__system | Exosome | 0.0185185 |
| Drosophila | omegaA | Ubiquitin__system | G__protein__coupled__receptors | 0.9814815 |
| Drosophila | omegaA | Ubiquitin__system | Glycosyltransferases | 0.0185185 |
| Drosophila | omegaA | Ubiquitin__system | Ion__channels | 0.1296296 |
| Drosophila | omegaA | Ubiquitin__system | Lipid__biosynthesis__proteins | 0.9444444 |
| Drosophila | omegaA | Ubiquitin__system | Membrane__trafficking | 0.0555556 |
| Drosophila | omegaA | Ubiquitin__system | Messenger__RNA__biogenesis | 0.0185185 |
| Drosophila | omegaA | Ubiquitin__system | Mitochondrial__biogenesis | 0.4629630 |
| Drosophila | omegaA | Ubiquitin__system | Peptidases | 0.1296296 |
| Drosophila | omegaA | Ubiquitin__system | Photosynthesis__proteins | 0.7222222 |
| Drosophila | omegaA | Ubiquitin__system | Proteasome | 0.6481481 |
| Drosophila | omegaA | Ubiquitin__system | Protein__kinases | 0.1666667 |
| Drosophila | omegaA | Ubiquitin__system | Protein__phosphatases__and__associated__proteins | 0.5740741 |
| Drosophila | omegaA | Ubiquitin__system | Ribosome | 0.3148148 |
| Drosophila | omegaA | Ubiquitin__system | Ribosome__biogenesis | 0.2037037 |
| Drosophila | omegaA | Ubiquitin__system | Spliceosome | 0.3148148 |

*(continued)*

| species | estimate | var1 | var2 | p.value |
|---|---|---|---|---|
| Drosophila | omegaA | Ubiquitin_system | Transcription_factors | 0.1296296 |
| Drosophila | omegaA | Ubiquitin_system | Transcription_machinery | 0.5000000 |
| Drosophila | omegaA | Ubiquitin_system | Transfer_RNA_biogenesis | 0.1666667 |
| Drosophila | omegaA | Ubiquitin_system | Translation_factors | 0.7222222 |
| Drosophila | omegaA | Ubiquitin_system | Transporters | 0.0185185 |

The next chunk presents the correlation performed between the categories of the two species.

```
# Will do this correlation for values of omegaA only
omegaA.arab <- subset(stat.arab, stat.arab$variable == "omega[a]")
omegaA.arab <- omegaA.arab[,-3] # removing the 'species' column
colnames(omegaA.arab) <- c("variable", "FunctionalClass",
                           "value.mean.arab", "value.sd.arab")

omegaA.dmel <- subset(stat.dmel, stat.dmel$variable == "omega[a]")
omegaA.dmel <- omegaA.dmel[,-3] # removing the 'species' column
colnames(omegaA.dmel) <- c("variable", "FunctionalClass",
                           "value.mean.dmel", "value.sd.dmel")

cor.kegg <- merge(omegaA.arab, omegaA.dmel, by = c("variable",
                                                   "FunctionalClass")) # 21 categories

plot.cor <- ggplot(cor.kegg, aes(value.mean.arab, value.mean.dmel,
                                 label = FunctionalClass)) +
  geom_point() +
  geom_smooth(method = "lm", se = TRUE) +
  xlab(expression(paste("Mean",~omega[a],~italic(A.thaliana), sep = ""))) +
  ylab(expression(paste("Mean",~omega[a],~italic(D.melanogaster), sep = ""))) +
  theme_bw() +
  theme.plot()
plot.cor
```

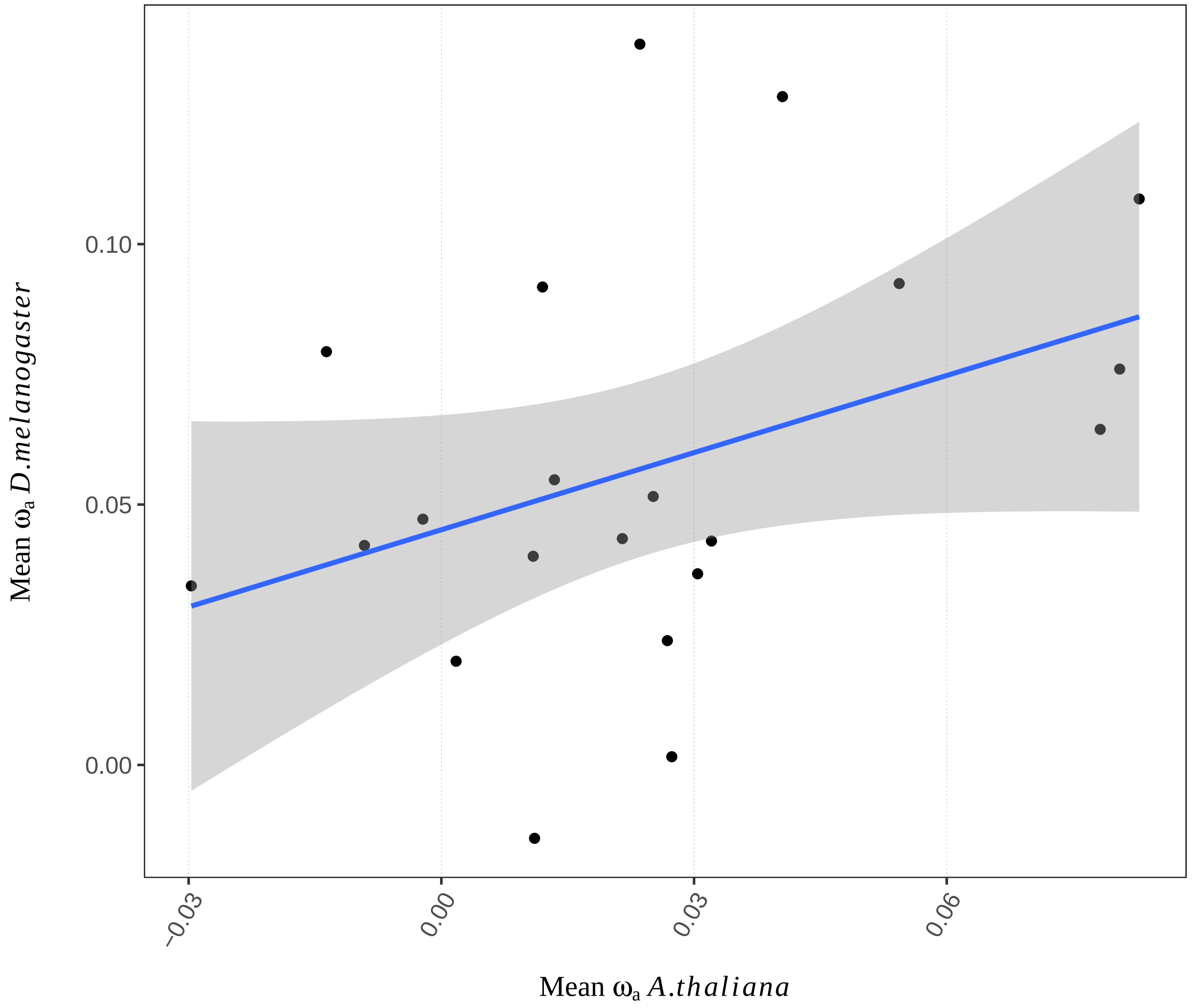


```
# outliers: mRNA biogenesis (row 9) and Glycositransferases (row 7)
sub.cor <- cor.kegg[-c(7,9),]
plot.subcor <- ggplot(sub.cor, aes(value.mean.arab, value.mean.dmel,
                                   label = FunctionalClass)) +
  geom_point() +
  geom_smooth(method = "lm", se = TRUE) +
  xlab(expression(paste("Mean",~omega[a],~italic(A.thaliana), sep = ""))) +
  ylab(expression(paste("Mean",~omega[a],~italic(D.melanogaster), sep = ""))) +
  theme_bw() +
  theme.plot()
plot.subcor
```

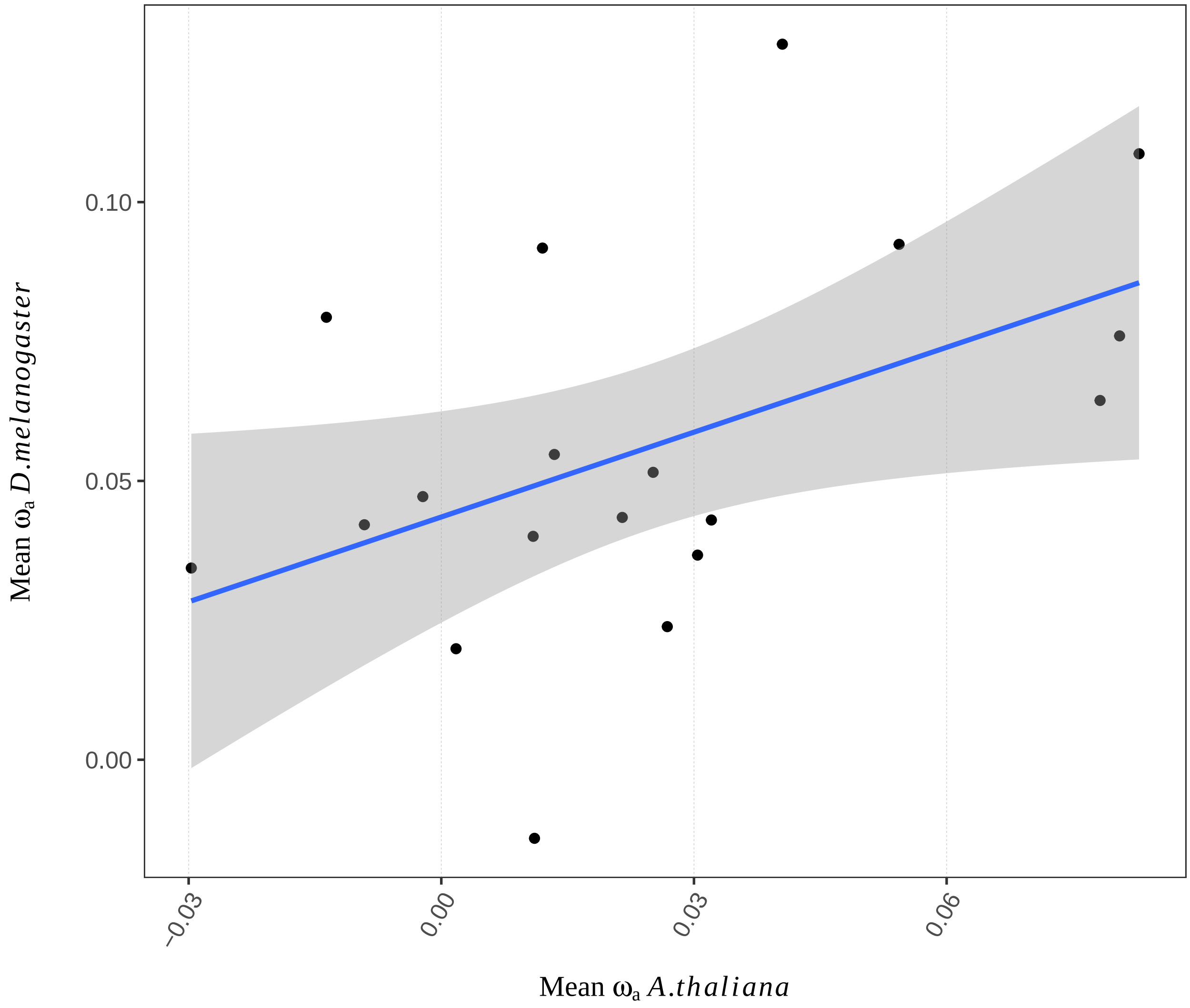


Statistical tests performed on both correlations:

```
# full data:
cor.all <- cor.test(~value.mean.arab+value.mean.dmel,cor.kegg, method = "kendall")
Kendall.tau <- cor.all$estimate
p.value <- cor.all$p.value
stat.all <- data.frame(Kendall.tau, p.value)
stat.all
```

| | Kendall.tau | p.value |
|---|---|---|
| tau | 0.2571429 | 0.1101277 |

```
# without outliers
cor.sub <- cor.test(~value.mean.arab+value.mean.dmel,sub.cor, method = "kendall")
Kendall.tau <- cor.sub$estimate
p.value <- cor.sub$p.value
sub.stat <- data.frame(Kendall.tau, p.value)
sub.stat
```

| | Kendall.tau | p.value |
|---|---|---|
| tau | 0.3333333 | 0.0490493 |
