## Supplemental Data 1 for "The impact of protein architecture on adaptive evolution"

### Supplementary Figures

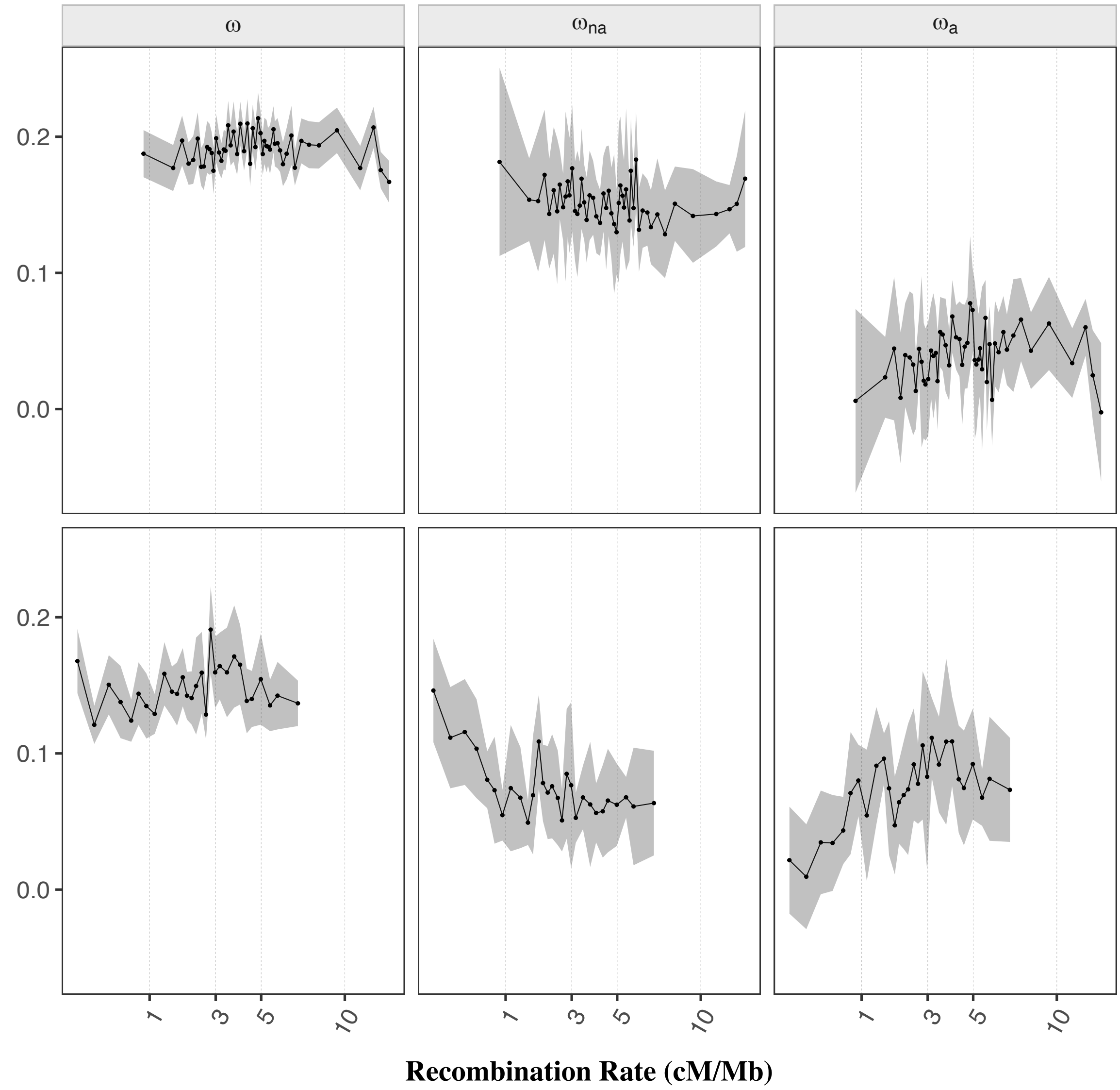


**Figure S1.** Correlation of ω, $\omega_{na}$ and $\omega_a$ with mean values of recombination rate for *A. thaliana* (on top) and *D. melanogaster* (in the bottom). Mean values of each estimate for each category are represented with the connected black dots. The shaded area represents the 95% confidence interval.

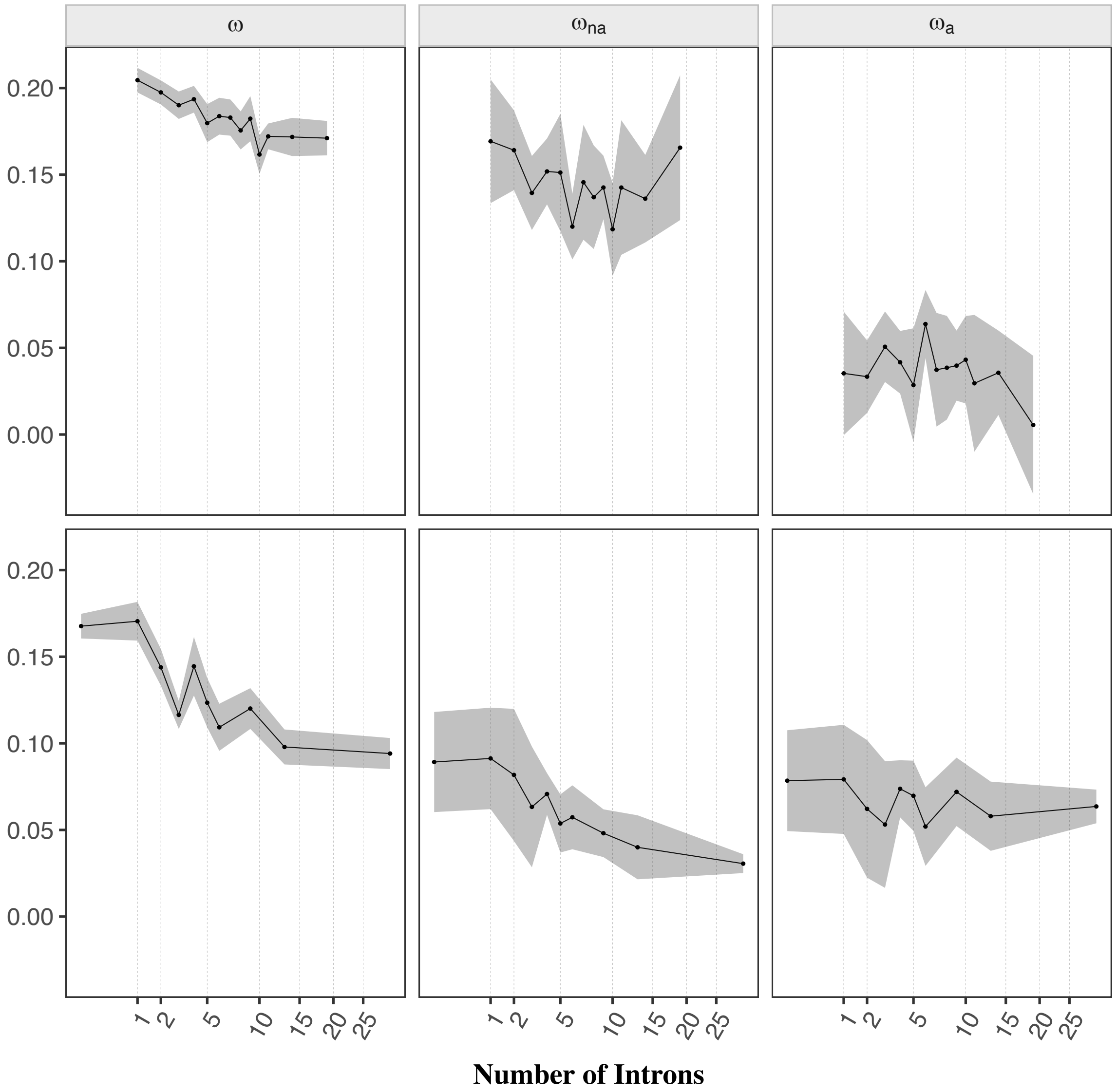


**Figure S2.** Estimates of ω, $\omega_{na}$ and $\omega_a$ for each category of intron number for *A. thaliana* (on top) and *D. melanogaster* (in the bottom). The x axis is scaled using a squared root function. Mean values of each estimate for each category are represented with the connected black dots. The shaded area represents the 95% confidence interval.

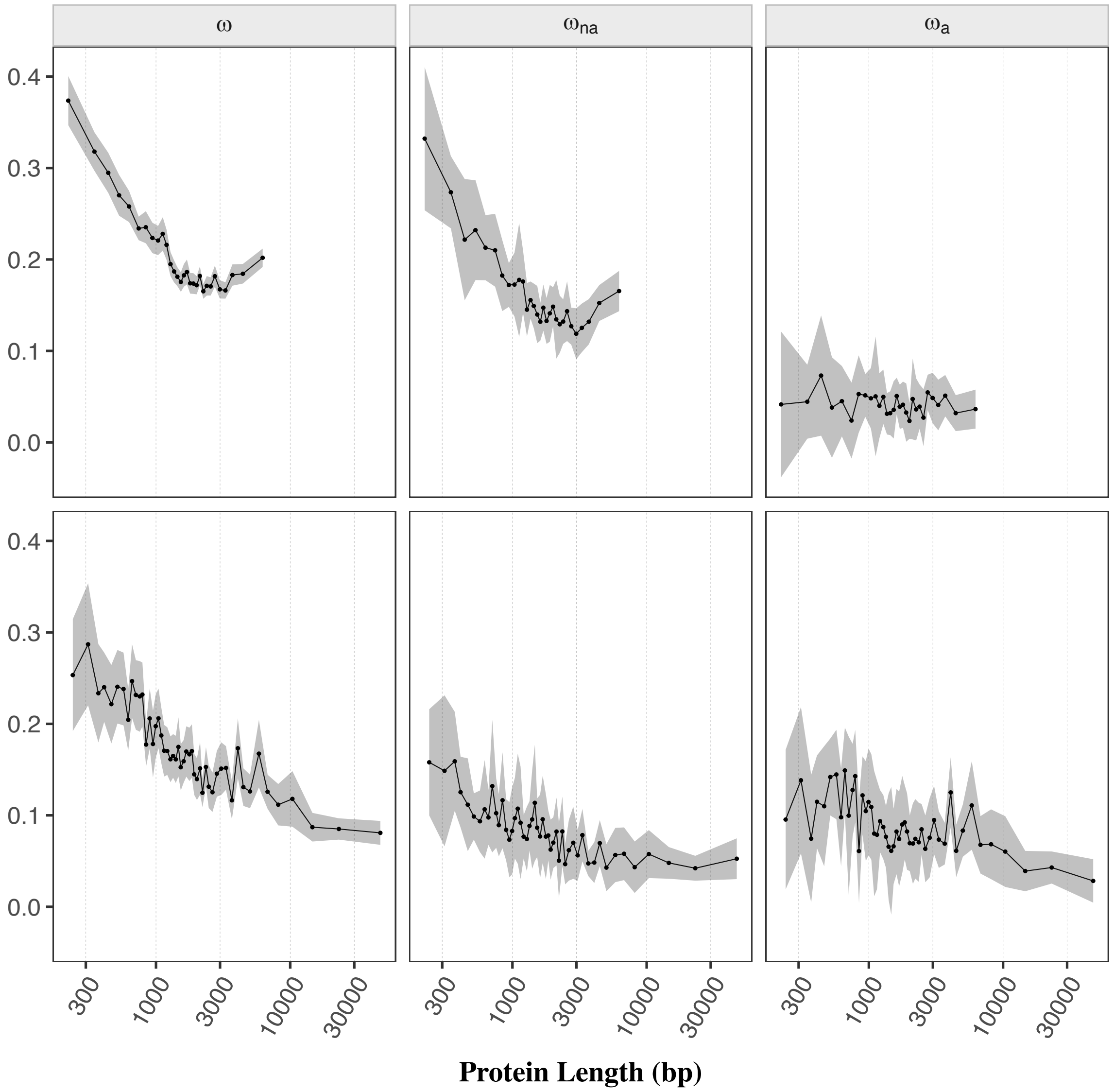


**Figure S3.** Correlation of ω, $\omega_{na}$ and $\omega_a$ with protein length for *A. thaliana* (on top) and *D. melanogaster* (in the bottom). The x axis is log scaled. Mean values of each estimate for each category are represented with the connected black dots. The shaded area represents the 95% confidence interval.

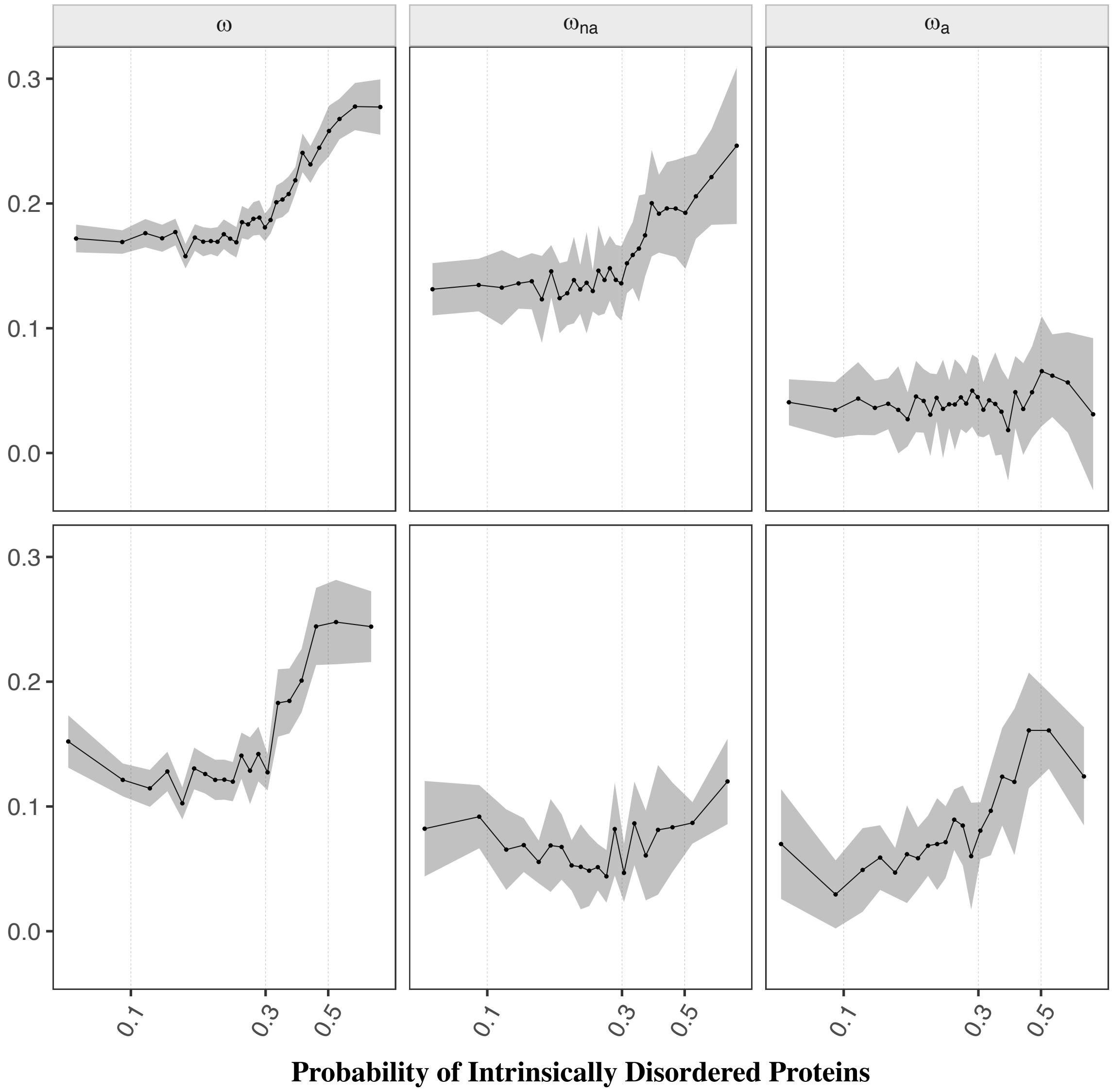


**Figure S4.** Correlation between ω, $\omega_{na}$ and $\omega_{a}$ with the probability of intrinsically disordered proteins for *A. thaliana* (on top) and *D. melanogaster* (in the bottom). The x axis is log scaled. Mean values of each estimate for each category are represented with the connected black dots. The shaded area represents the 95% confidence interval.

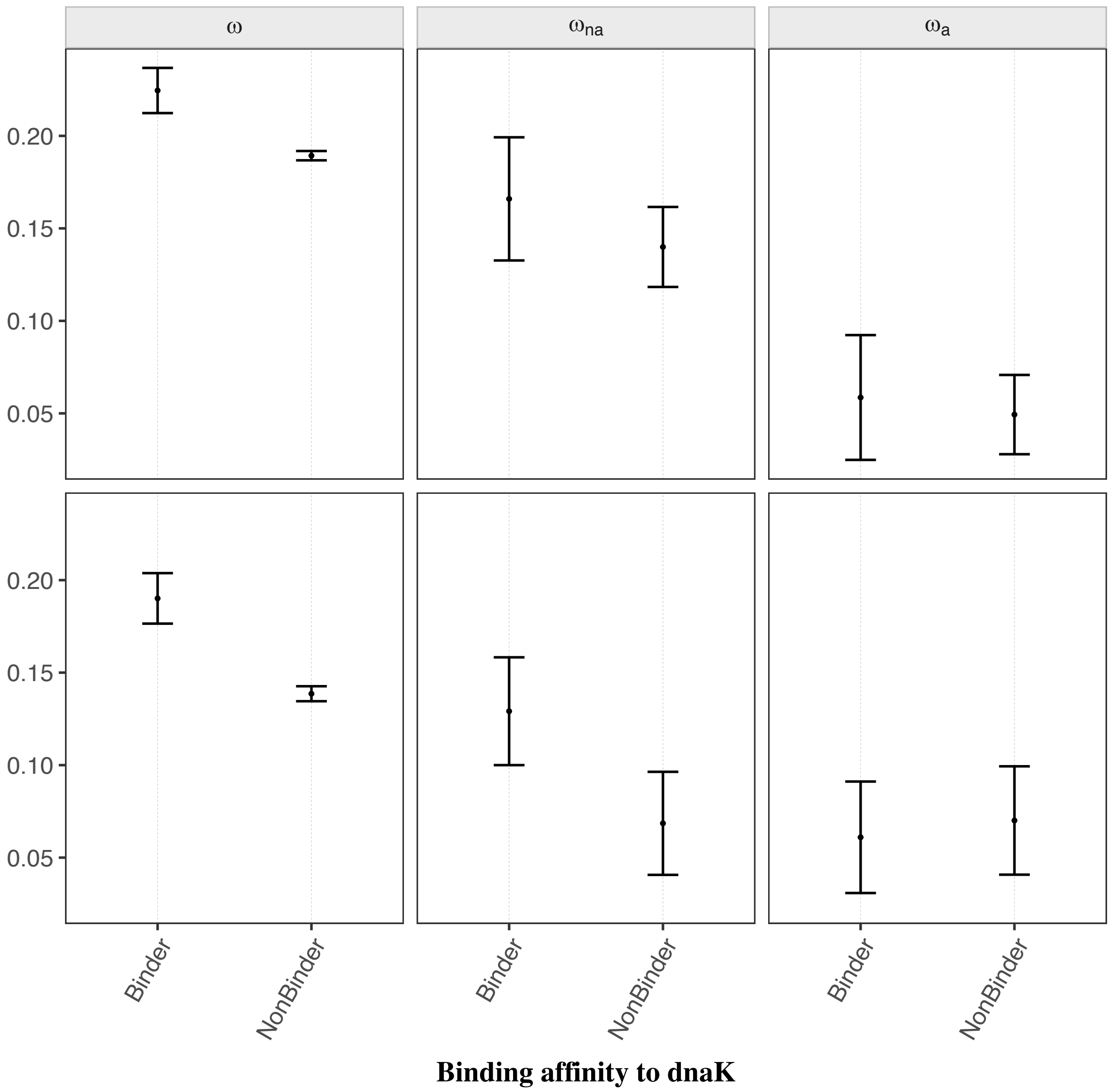


**Figure S5.** Estimates of ω, $\omega_{na}$ and $\omega_a$ for binders and non-binders to the molecular chaperone *DnaK*, in *A. thaliana* (top) and *D. melanogaster* (bottom). Mean values of each estimate for each category are represented with the black points. The error bar denotes for the 95% confidence interval.

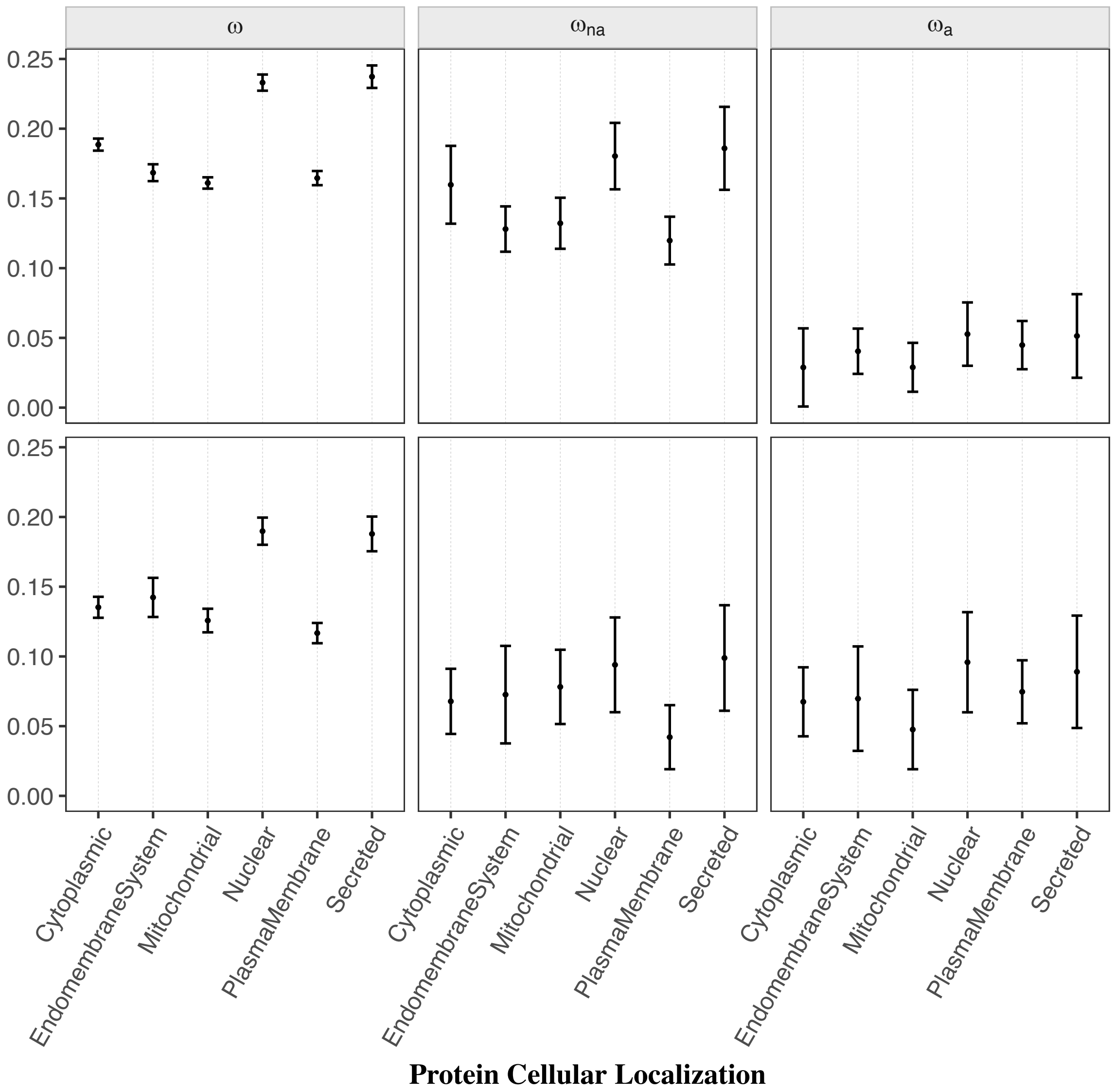


**Figure S6.** Estimates of the rate of ω, $\omega_{na}$ and $\omega_a$ for each category of cellular localization of proteins in *A. thaliana* (top) and *D. melanogaster* (bottom). Mean values of each estimate for each class are represented with the black points. The error bar denotes for the 95% confidence interval for each category.

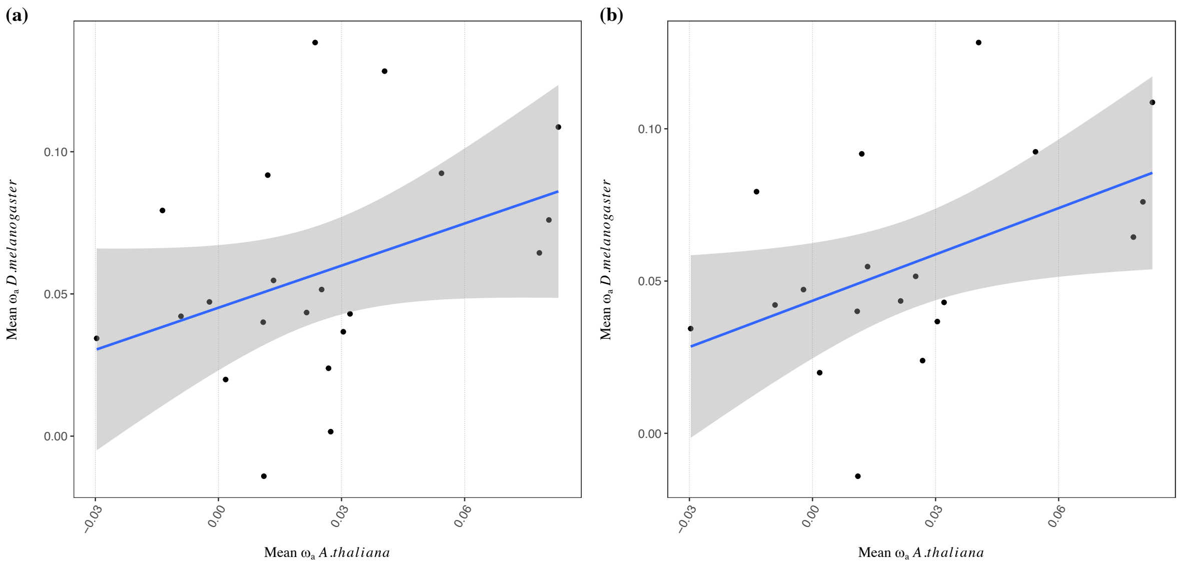


**Figure S7.** Relationship between mean $\omega_a$ values of the common categories of protein functional class among species (total of 21 categories). (a) A linear model was fitted between $\omega_a$ values of the common functional classes, the standard error is represented. (b) After removing the outliers from the correlation in (a), a new linear model was fitted with the respective standard error.

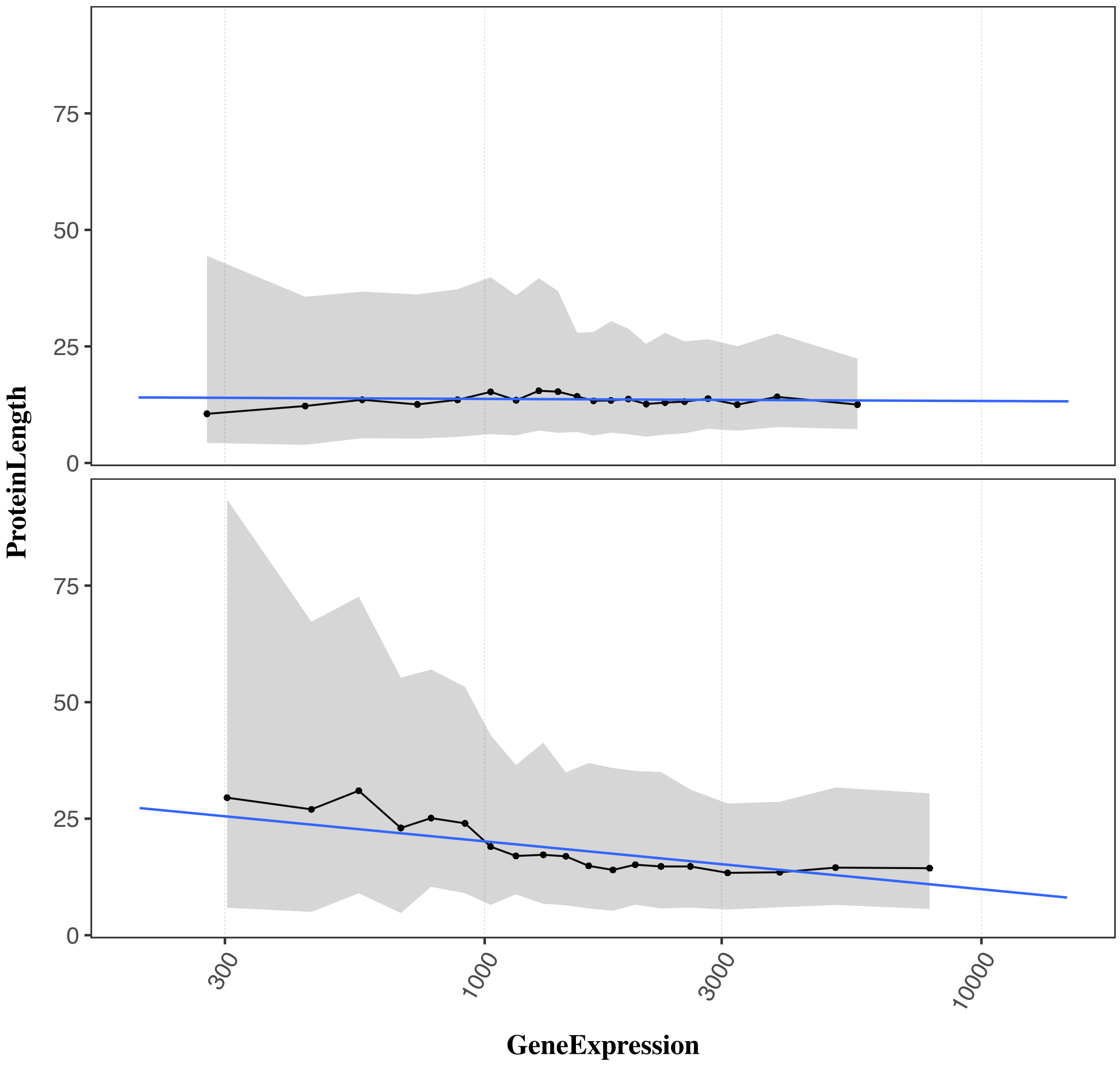


**Figure S8.** Relationship between protein length and mean expression levels for *A. thaliana* (top) and *D. melanogaster* (bottom). The x axis is log scaled. This analysis was accomplished by categorizing protein length in 20 categories in each species. For each category of protein length, the median value of gene expression is depicted with the black dot. The shaded area represents the values of mean expression levels within the 1$^{st}$ and 3$^{rd}$ quartile. The blue line represents the 50% quantile regression fitted among the values of protein length and gene expression for each gene.

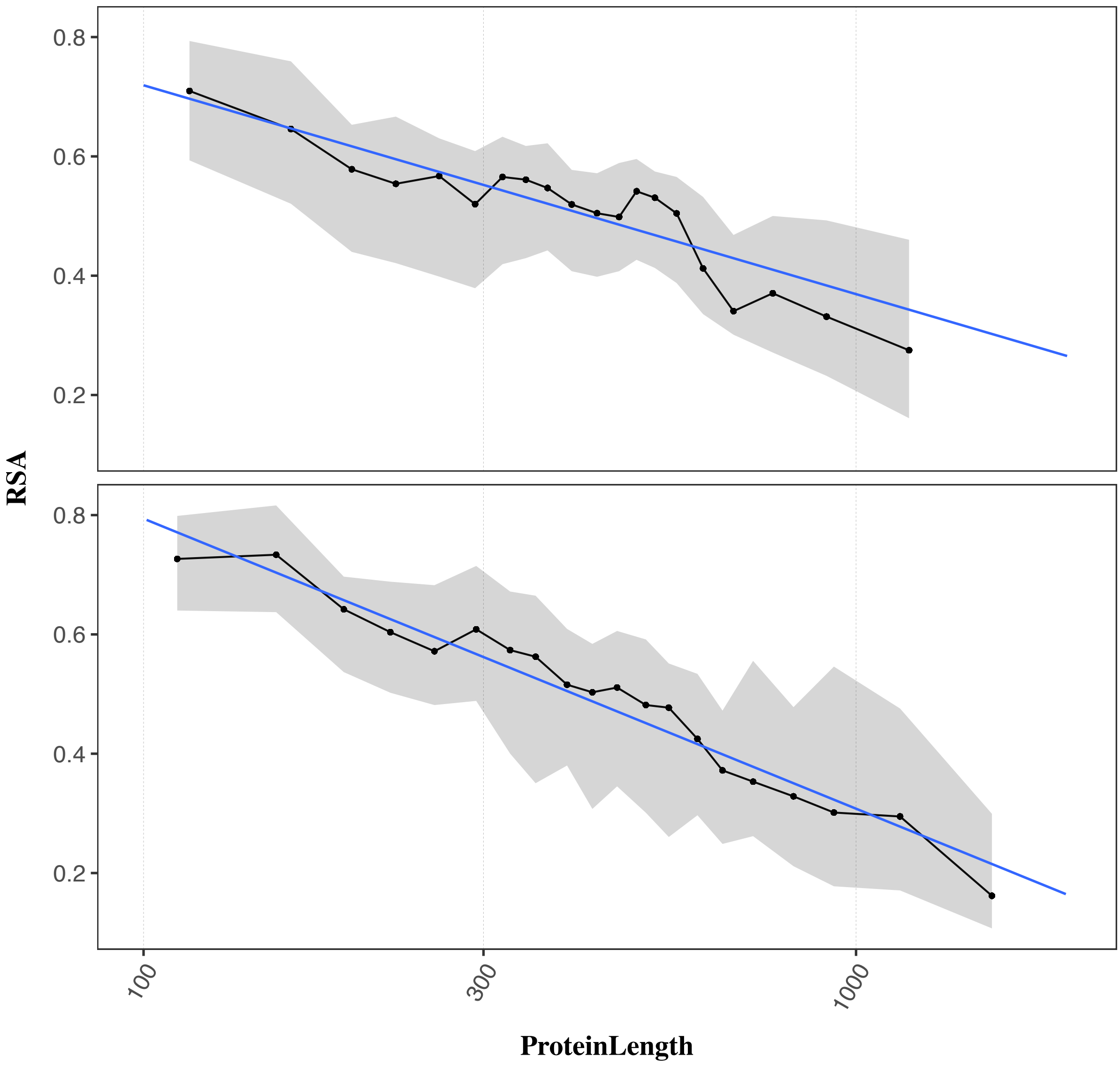


**Figure S9.** Relationship between protein length and proportion of exposed residues per protein for *A. thaliana* (top) and *D. melanogaster* (bottom). The x axis is log scaled. For the purpose of this analysis, a residue was considered exposed if RSA >= 0.05. The proportion of exposed sites per protein was then calculated by counting the number of amino-acid residues. This analysis was accomplished by categorizing protein length in 20 categories in each species. For each category of protein length, the median value of the proportion of exposed sites is depicted with the black dot. The shaded area represents the values for the proportion of exposed residues within the 1st and 3rd quartile. The blue line represents the 50% quantile regression fitted among the values of protein length of each gene with the respective estimate of the proportion of residue exposure.

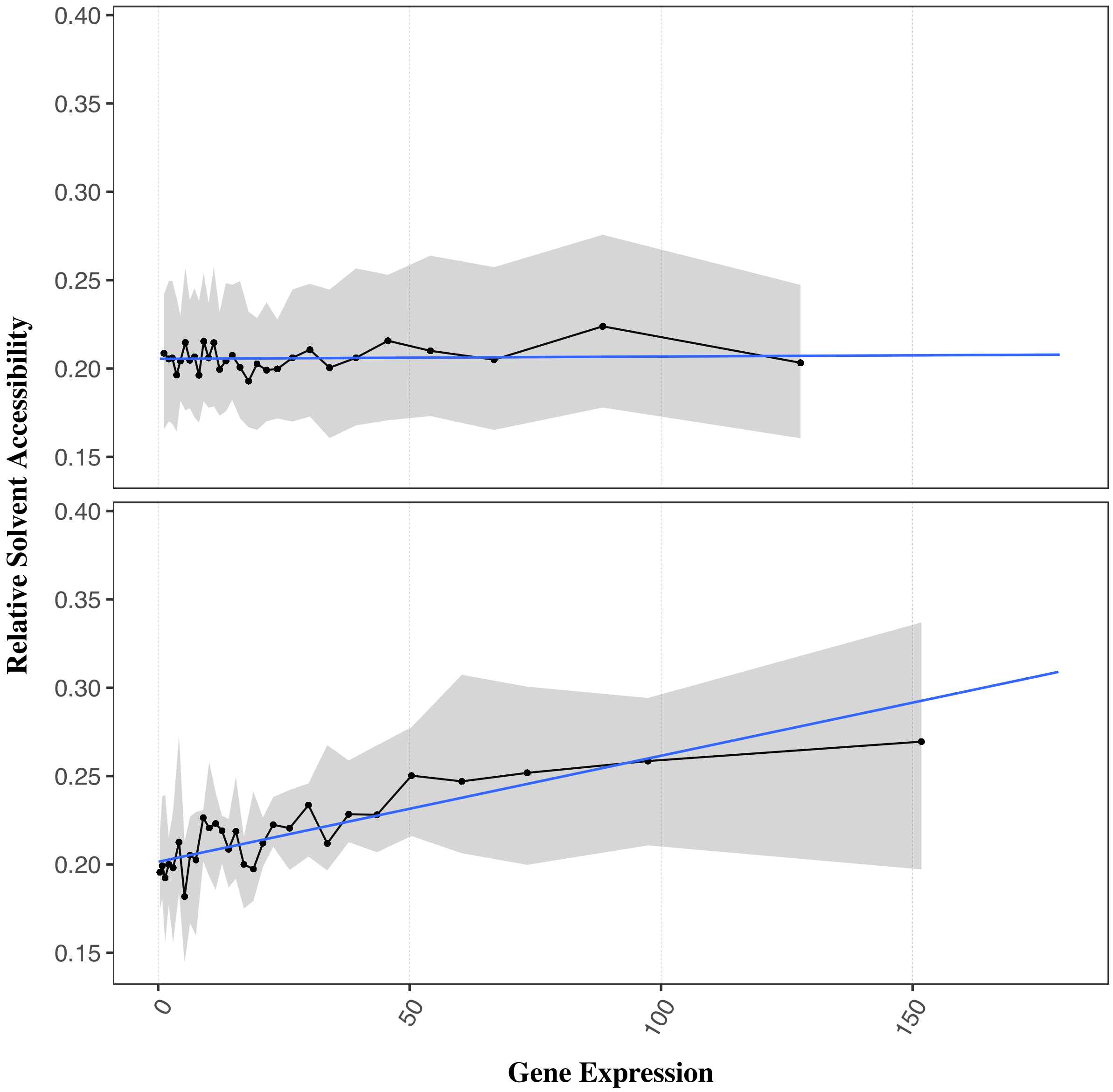


**Figure S10.** Relationship between mean gene expression and solvent exposure for *A. thaliana* (top) and *D. melanogaster* (bottom). For each value of mean gene expression, the median value of RSA is represented. This analysis was accomplished by categorizing gene expression in 30 categories in each species. For each category of gene expression, the median value of RSA is depicted with the black dot. The shaded area represents the values of RSA within the 1st and 3rd quartile. The blue line represents the 50% quantile regression fitted among the values of gene expression of each gene with the respective estimate of RSA.

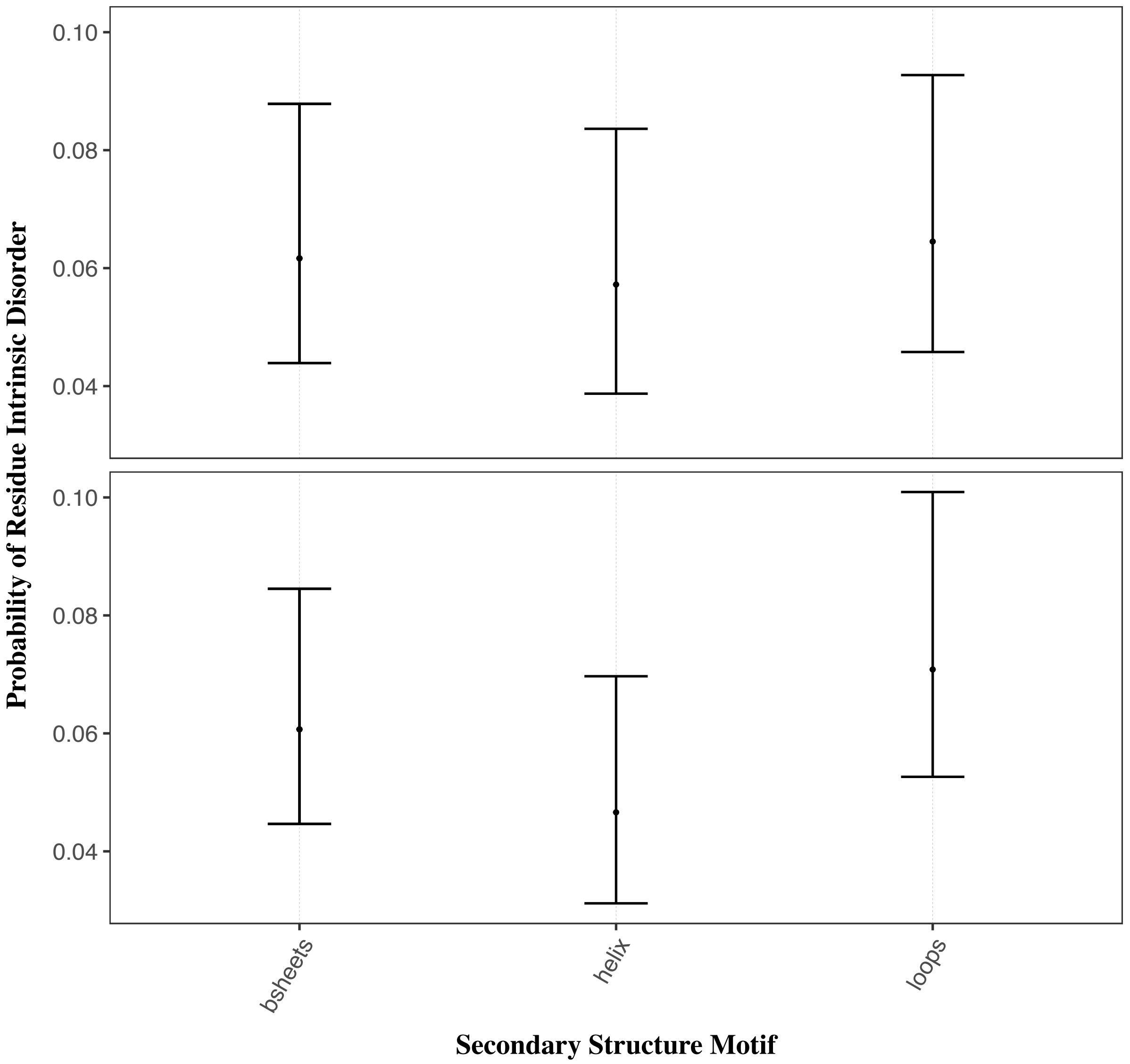


**Figure S11.** Relationship between the secondary structural motif (β-sheets, α-helices and loops) and probability of residue intrinsic disorder, for *A. thaliana* (top) and *D. melanogaster* (bottom). For each category of secondary structure, the median value of disorder probability is depicted with the black dot. The error bar represents the values of the probability of intrinsic disorder within the 1$^{st}$ and 3$^{rd}$ quartile.

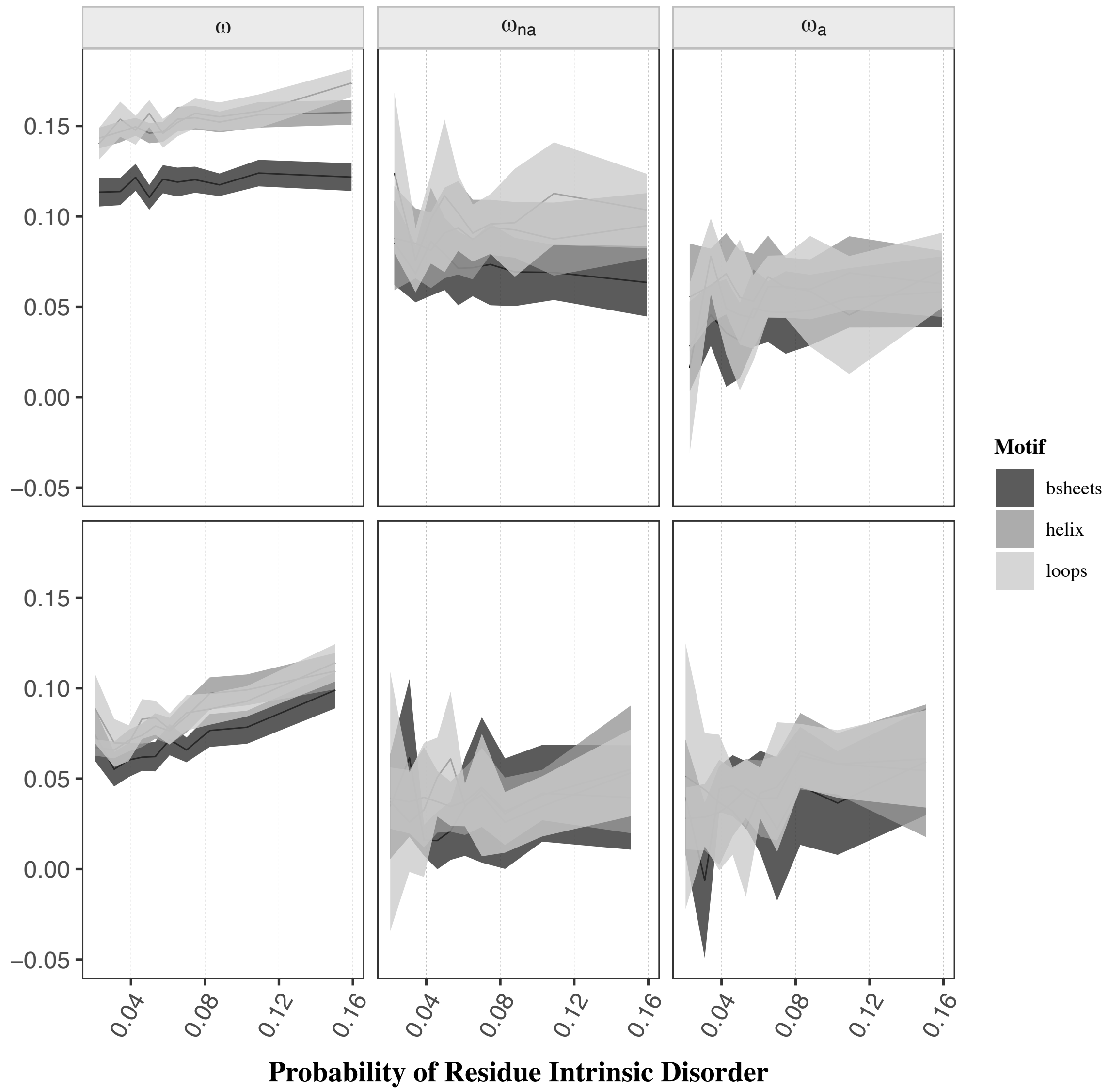


**Figure S12.** Estimates of ω, $\omega_{na}$ and $\omega_a$ for the correlation between the secondary structural motif and probability of residue intrinsic disorder in *A. thaliana* (top) and *D. melanogaster* (bottom). Analyses were performed by comparing residues present in β-sheets, α-helices and loops across 10 categories of intrinsic disorder for both species. Mean values of each estimate for each category are represented with the connected black dots. The shaded area represents the 95% confidence interval.

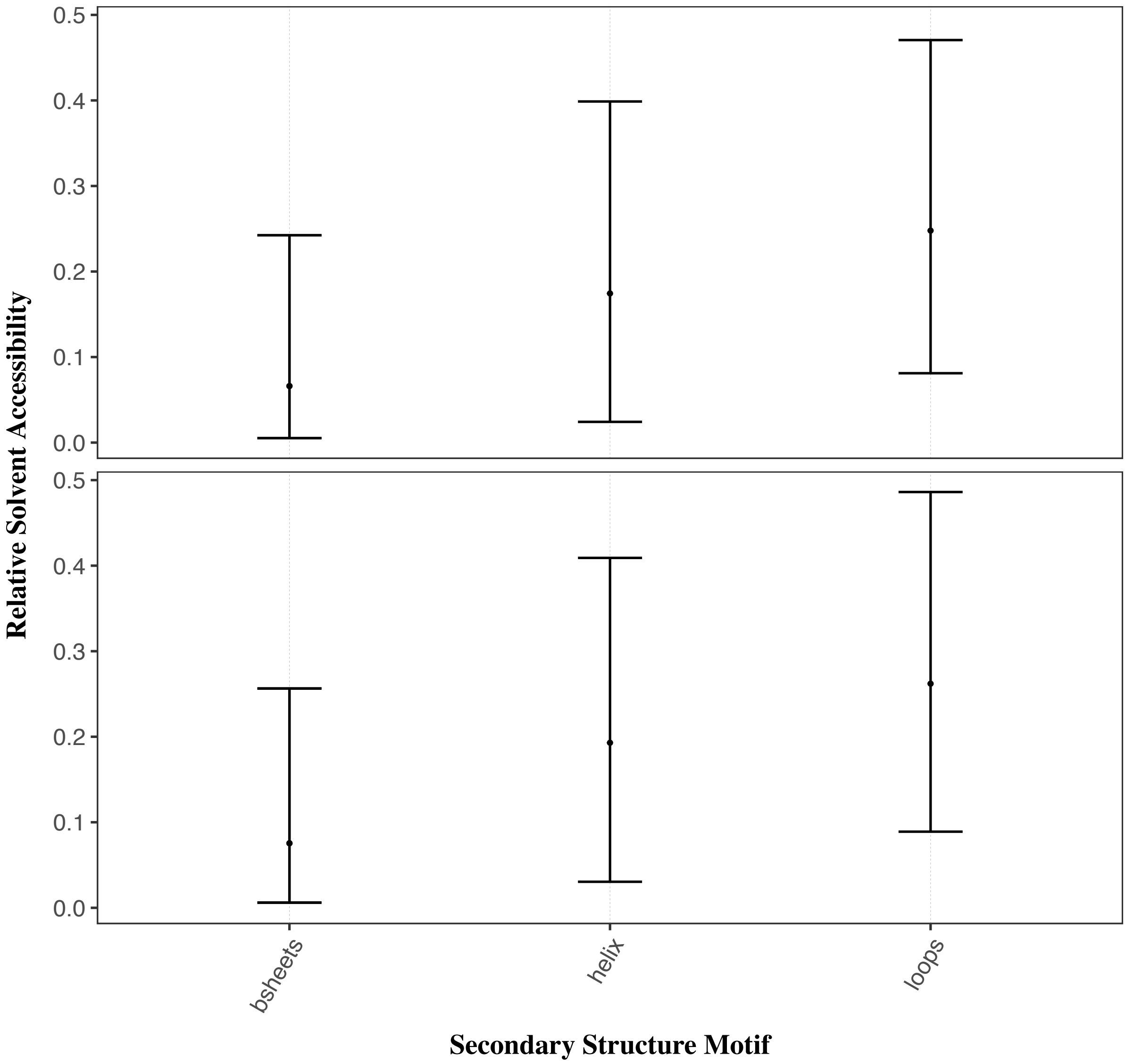


**Figure S13.** Relationship between the secondary structural motif (β-sheets, α-helices and loops) and solvent exposure, for *A. thaliana* (top) and *D. melanogaster* (bottom). For each category of secondary structure, the median value of solvent accessibility is depicted with the black dots. The error bar represents the values of RSA within the 1$^{st}$ and 3$^{rd}$ quartile.

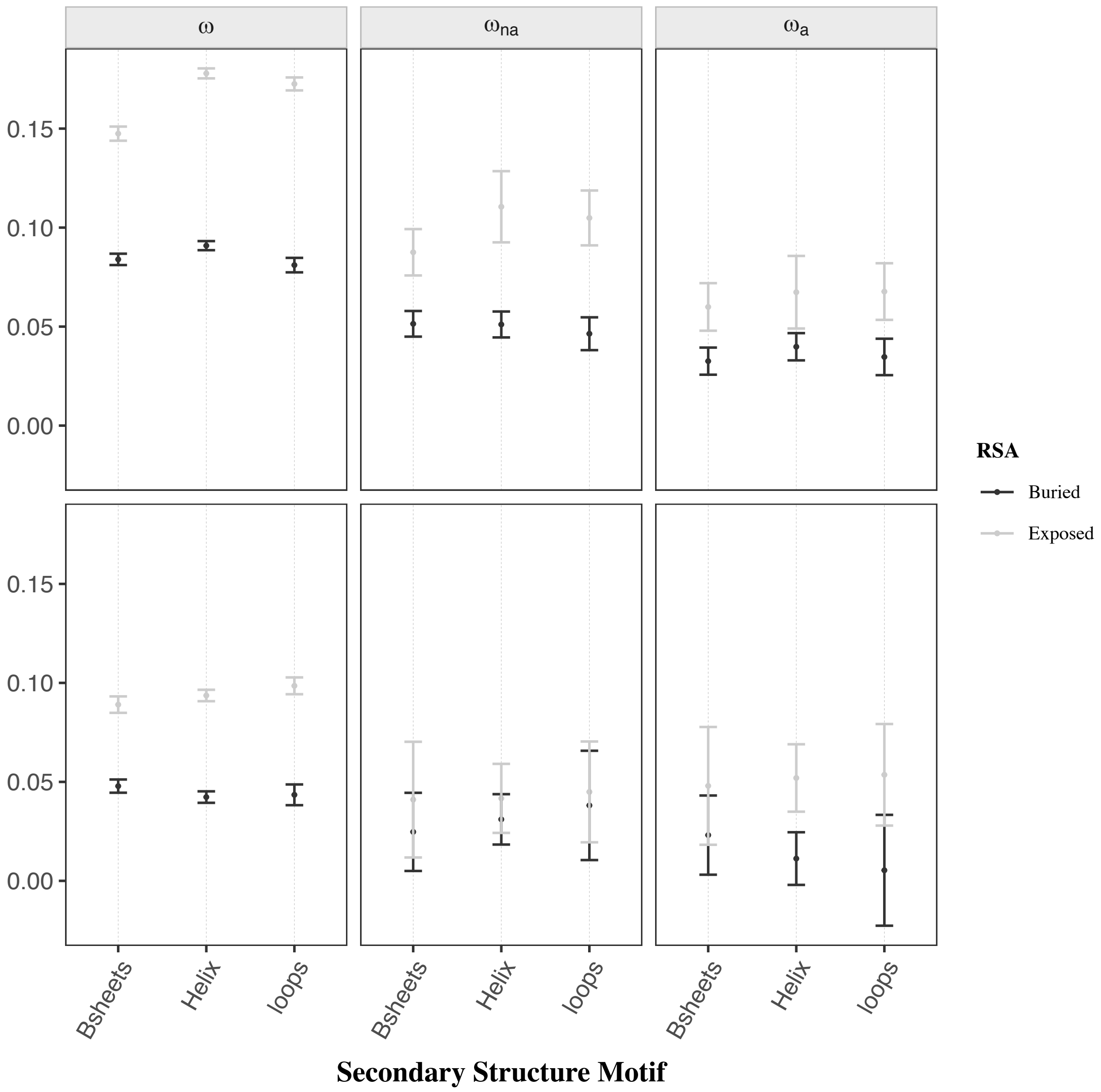


**Figure S14.** Estimates of ω, $\omega_{na}$ and $\omega_a$ for the correlation between the relative solvent accessibility and the secondary structural motif (β-sheets, α-helices and loops) in *A. thaliana* (top) and *D. melanogaster* (bottom). Analyses were performed by comparing buried (RSA < 0.05) and exposed (RSA >= 0.05) residues across the three structural motifs. Mean values of each estimate for each category are represented with the black dots. The error bar represents the 95% confidence interval.

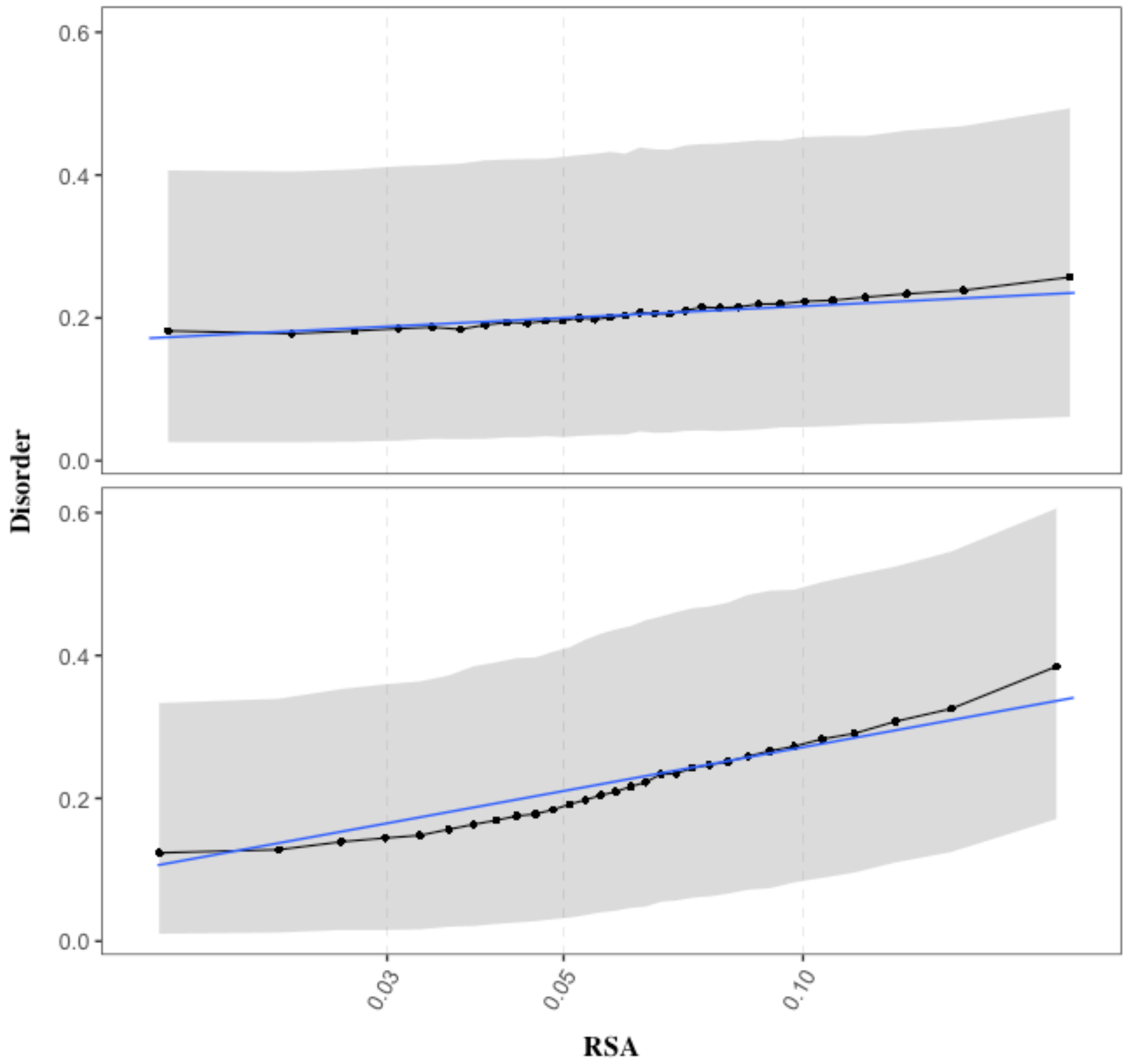


**Figure S15.** Relationship between probability of intrinsic disorder and the relative solvent accessibility of a residue. For the purpose of this analysis, a residue was considered exposed if RSA >= 0.05. This analysis was accomplished by categorizing the probability of residue intrinsic disorder in 30 categories in each species. For each category of disorder, the median value of solvent exposure is depicted with the black dot. The shaded area represents the values of RSA within the 1st and 3rd quartile. The blue line represents the 50% quantile regression fitted among 10,000 randomly sampled residues for each category, since we were dealing with big amounts of data.

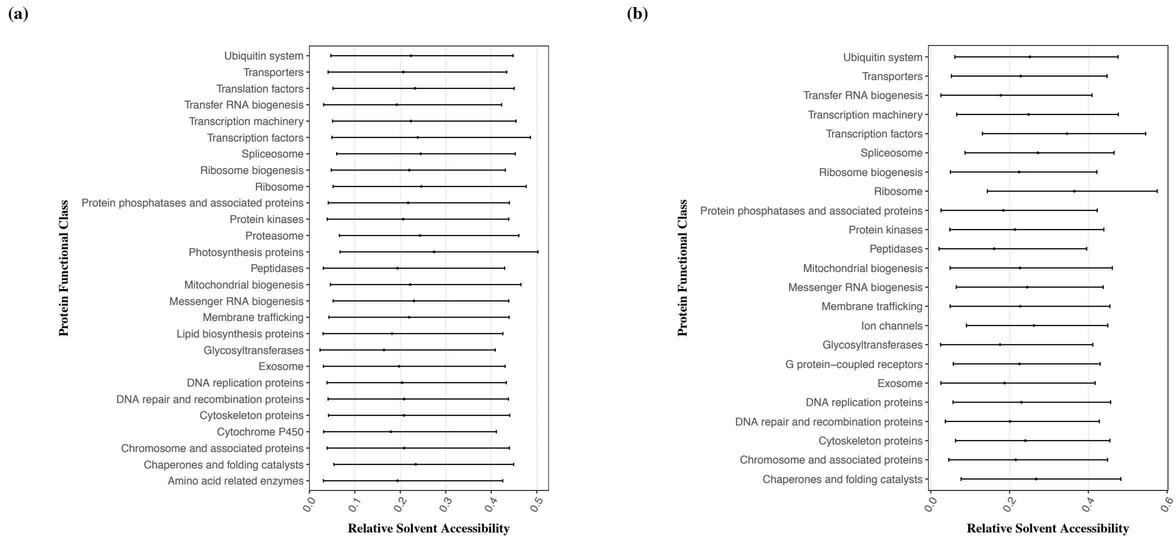


**Figure S16.** Relationship between the protein functional class and solvent exposure, for *A. thaliana* (a) and *D. melanogaster* (b). For each category of protein function, the median value of solvent accessibility is depicted with the black dot. The error bar represents the values of RSA within the 1st and 3rd quartile.

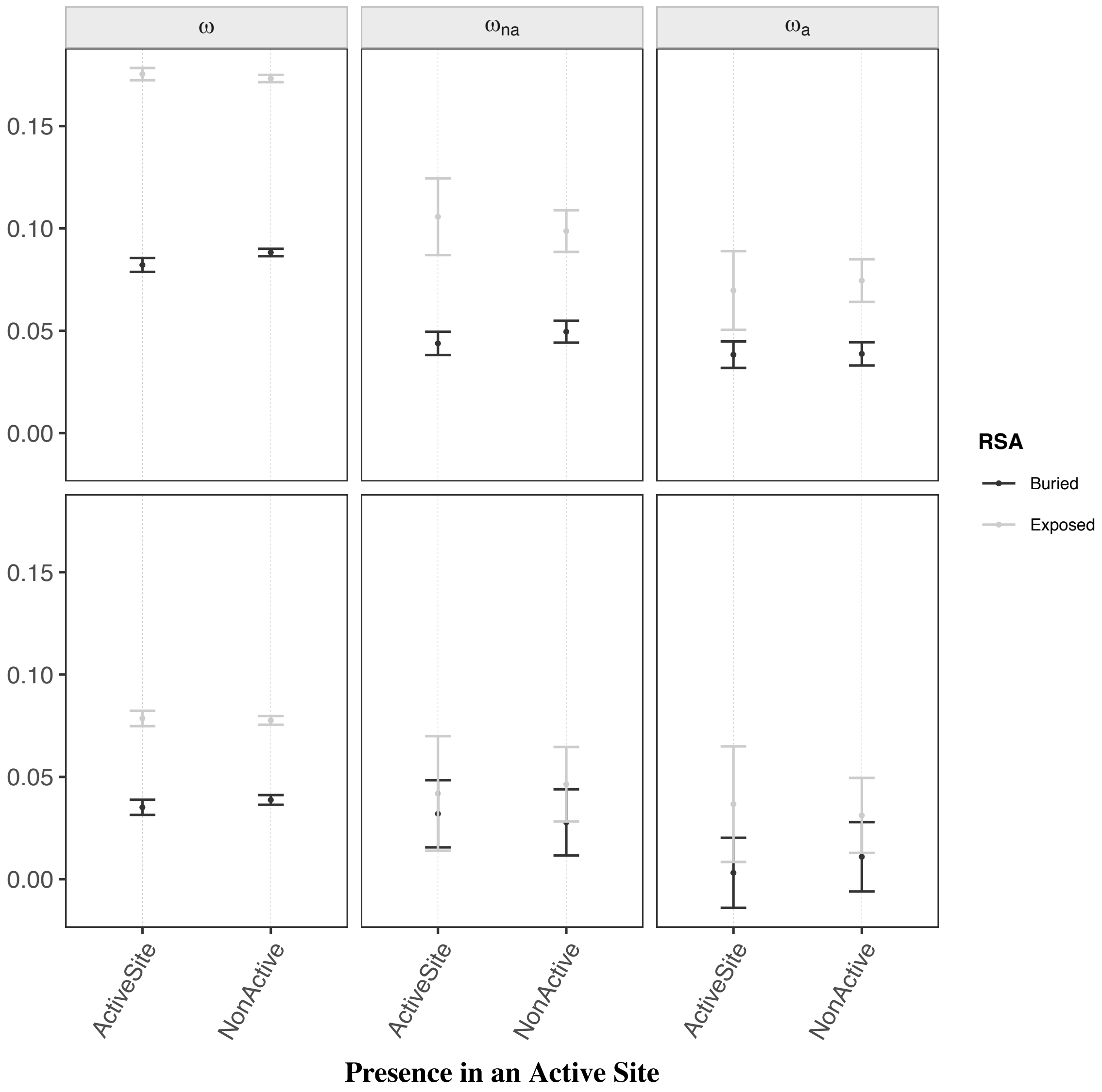


**Figure S17.** Estimates of ω, $\omega_{na}$ and $\omega_a$ for active and non-active sites, correcting for solvent exposure in *A. thaliana* (on top) and *D. melanogaster* (bottom). Analyses were performed by comparing buried (RSA < 0.05) and exposed (RSA >= 0.05) sites in residues present, and not present, in an active site. Mean values of each estimate for each category are represented with the black dots. The error bar represents the 95% confidence interval.

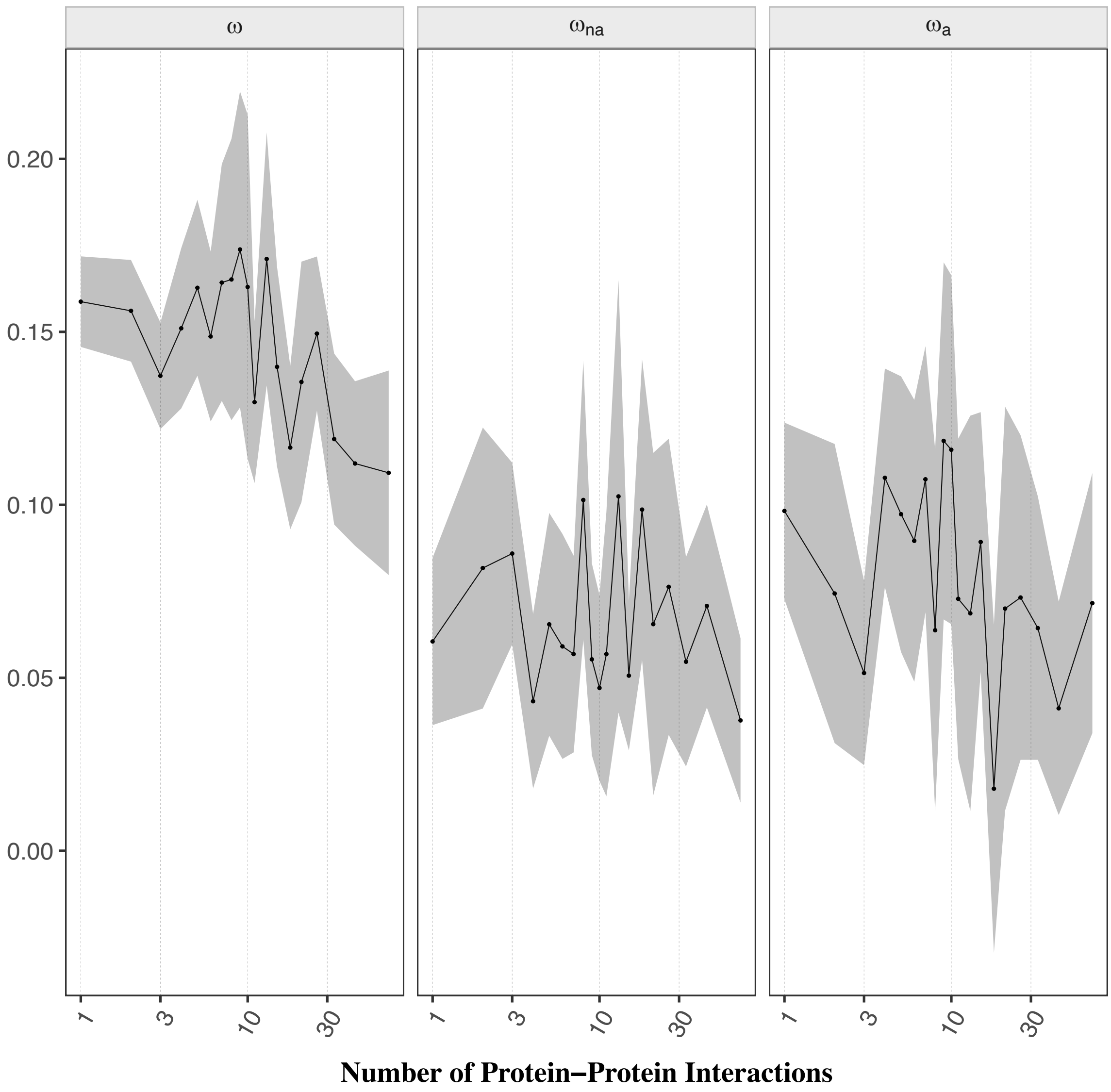


**Figure S18.** Relationship between ω, $\omega_{na}$ and $\omega_a$ with the number of protein-protein interactions for *D. melanogaster*. The x axis is log scaled. Mean values of each estimate for each category are represented with the connected black dots. The shaded area represents the 95% confidence interval for each category.
